# Marker genes for predicting cytokine release syndrome in vitro before CAR T cell infusion

**DOI:** 10.1101/2025.05.26.655341

**Authors:** Mengxiang Chen, Tao Wu, Yunfei Hu, Qiulin Liu, Mengfei Chen, Jing Zhang, Yan Du, Fu Ouyang, Yunhong Huang

## Abstract

Cytokine release syndrome (CRS) and neurotoxicity are common adverse events of the Chimeric antigen receptor (CAR) T cell therapy. Assessing the cytotoxicity associated biomarkers would be essential for therapy design to avoid developing severe toxicities. In this study, we re-analyzed previously published RNAseq results of CAR T cells before infusion and combined it with the clinical response post infusion. We observed that CAR T cells from patients who developed severe CRS displayed a higher expression of TCL6, HPCAL4, CCDC144B, and SIRPG, but lower levels of IL2, IL21, and HSPA1B when stimulated with anti-CAR19 idiotypic antibody in vitro. Interestingly, the upregulated gene SIRPG is positively correlated with CRS severity. In addition, without stimulation, CAR T cells from CRS group showed a higher levels of IFNAR1, IL7R, ZNF69, and USP32P1 but lower levels of CCL3, IL4, IL17A, IL23R, IL13, CD70, and IFNGR2. These results provided insights to evaluate the adverse events of CAR T products before treatments, which could be beneficial for designing therapy plans.

## Introduction

Chimeric antigen receptor (CAR) T cell therapy involves the infusion of genetically modified autologous T cells from the patient to mediate antitumor effects. The CAR includes an antigen-recognition domain that can specifically recognize CD19 on B lymphocytes and trigger T cell activation[1, 2]. It has shown remarkable activity in the treatment of B cell malignancies including pre-B cell acute lymphoblastic leukemia and diffuse large B cell lymphoma [1]. Currently, FDA has approved six CART cell therapies including Kymriah, and there are hundreds in clinical development for other hematological and solid tumors[3].

Even though many patients achieve complete remission after CAR T cell therapy and the genetically modified T cell represents a therapeutic strategy that changing the drug development, life-threatening toxicities such as cytokine release syndrome (CRS) and immune effector cell-associated neurotoxicity syndrome (ICANS) are common side effects post CAR T cell infusion[1, 4]. CRS is a systemic inflammatory response commonly occurring 2-3 days after the first infusion of CAR T cells. CRS of any grade was observed in 73.4% of patients, this syndrome can be more severe than the influenza-like syndrome with 27.4% of patients developing severe CRS[5]. The neurologic complications can be mild and are largely reversible, however, Life-threatening neurologic toxicities including cerebral edema have also been reported across different clinical studies of CD19-specific CAR T therapies [1, 5].

Although CAR-T cell therapy has been a revolutionary treatment for B cell malignancies, a high percentage of toxicities has prevented it from becoming the first-line therapy[6]. The mechanisms underlying T cell immunotherapy associated CRS and cerebral edema are poorly understood. The observed evidence of endothelial injury suggests that the inflammatory cytokines contribute to the onset of neurotoxicity[7]. Recent studies have identified factors that could predict toxicities after CAR T cell infusion, such as tumor burden, platelet count, and blood cytokines[5]. These biomarkers are essential for evaluating CAR T treatments before developing life threatening complications. However, it is unknown whether the prediction could be performed before CAR T cell infusion using an in vitro model.

In this study, we combined the bulk RNAseq results of stimulation CAR T cells in vitro and clinical data in a previously published trial [8] to evaluate the potential relationship between cytokine gene expression in CAR T cell cultures and severe CRS post infusion. Our results showed that stimulating CAR T cells with anti-CAR19 idiotypic antibody in vitro activated a broad spectrum of cytokine gene expression including IL3, CCL1, IL8, IL13, CCL3, and CCL4. Compared to without severe CRS, the patients who developed severe CRS displayed higher expression of TCL6, HPCAL4, CCDC144B, and SIRPG, while lower expression of IL2, IL21, and HSPA1B in T cells products. In addition, we also analyzed the differentially expressed genes prior to stimulation, and identified the intrinsic differences in gene expression between severe CRS and non-severe CRS patients. These data suggest that stimulating CAR T cells in vitro with anti-CAR19 idiotypic antibody could model clinical response post infusion, which could serve as a screening tool for CAR T therapy.

## Results

### Expression profile of CAR T cell activation in vitro

According to the original experiment design, CD19-targeted T cells (CTL019) were stimulated with a bead-bound anti-CAR19 idiotypic antibody in vitro, which serves as a surrogate for congnate CD19 antigen[8, 9]. This treatment mimics the recognition of CD19 and activates CTL019 cells. Because the original study was designed to identify the determinants of different responses between complete remission (CR) and nonresponding (NR) patients derived CAR-T cells, a direct comparison of transcriptome between untreated and treated CTL019 cells would provide a gene signature for evaluating CAR T cell activation before infusion. By using DESeq2 paired comparison workflow, we reanalyzed this dataset which included CTL019 cells from 13 patients.

When compared to the untreated control, a total of 3165 differentially expressed genes (p<0.01) were identified in anti-CAR19 antibody treated CTL019 cells (Table S1). As shown in the heatmap (Figure 1A), the majority of the up or down-regulated genes are generally consistent among each stimulated sample, and the gene expression profiles of stimulated CTL019 cells are markedly different from those of untreated cells. As shown in the volcano plot (Figure 1B), many of the differentially expressed genes are inflammatory cytokines including IL3, CCL1, IL8, IL13, CCL3, CCL4, etc, suggesting the activation of immune reaction by anti-CAR19 antibody treatment. We then performed GSEA (gene set enrichment analysis) and demonstrated that the differentially expressed genes are predominantly involved in TNF-NFKB, IL2-Stat5, IL6-JAK-Stat3, and interferon alpha and gamma pathways (Figure 1C). In addition, genes related to hypoxia and angiogenesis were also stimulated.

**Fig. 1.**
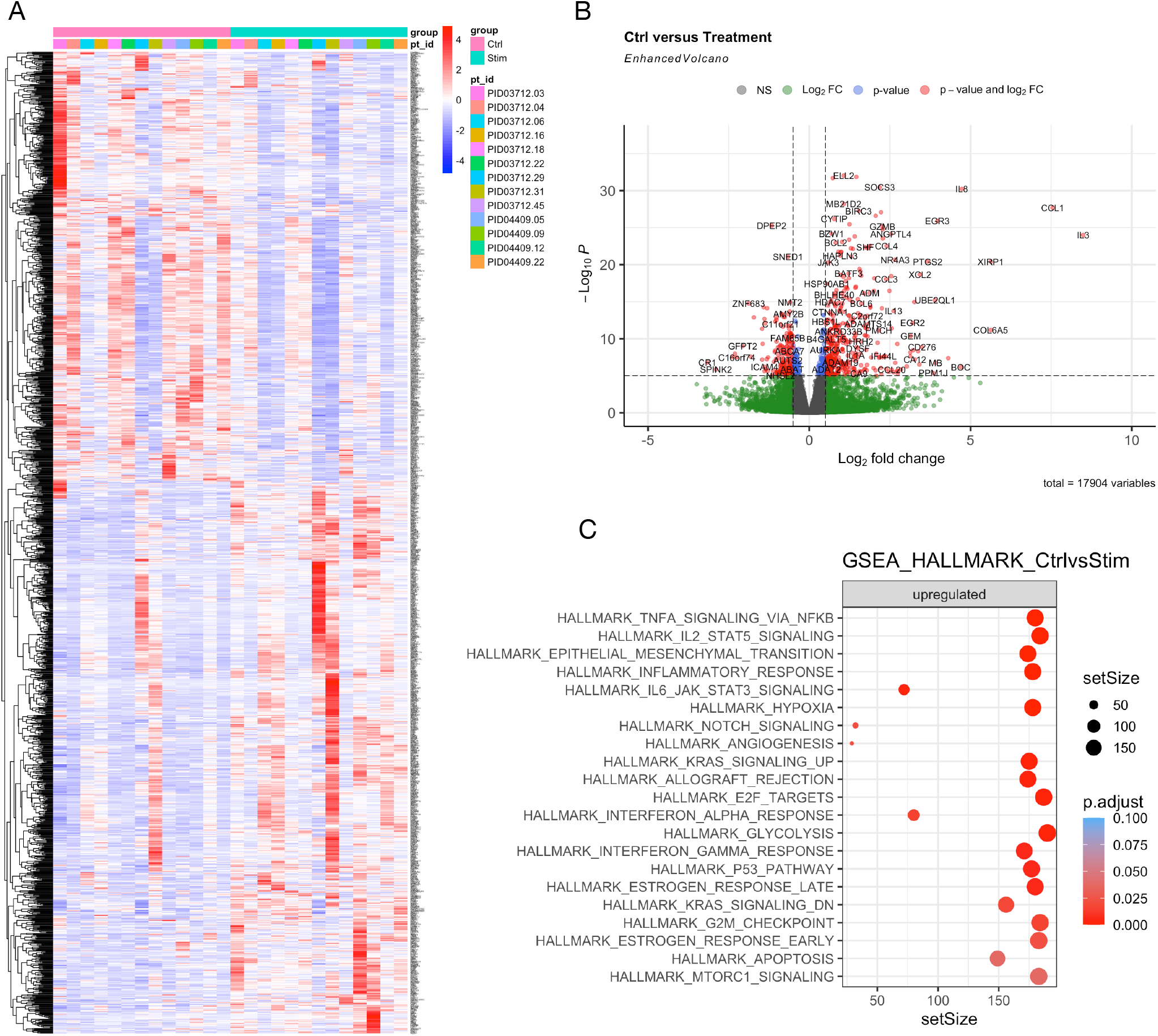
Expression profile of CAR T cell activation in vitro. (A) Heatmap shows differentially expressed genes in CTL019 cells before infusion. Differentially expressed genes between Control (Ctrl) and anti-idiotypic antibody stimulated CTL019 cells (Stim) were presented. (B) Volcano plot shows activated genes (red dots on right) in anti-idiotypic antibody stimulated CTL019 cells. (C) Gene Set Enrichment Analysis shows the activated genes in stimulated CTL019 cells enriched in signal pathways.

### Biomarkers of CAR T cell activation

To visualize the results at the sample level, we next present the top 20 differentially expressed genes according to the adjusted p values (Figure 2A). Except for up-regulating inflammatory response (CCL1, IL3, IL8, TNFRSF18, JUNB, and NR4A1), other stimulated genes by anti-CAR19 antibody including GZMB, a maker for CD8T and NK cells, which is crucial for the rapid induction of target cell apoptosis. ANGPTL4 is induced under hypoxic condition and promotes vascular inflammation and increases vascular permeability[10]. CRTAM (MHC class I– restricted T cell–associated molecule) is predominantly expressed on activated CD8+ T cells and NK/NKT cells[11]. We further analyzed the up-regulated individual cytokine, and found that the expressions of classic pro-inflammatory cytokines, IL1B, TNF, and IL6 are comparable between untreated and stimulated groups (Figure 2B). Significantly increased cytokines in the stimulated group are shown in Figure 2C. Together, these data suggested that anti-CAR19 antibody treatment in vitro stimulates a broad-spectrum cytokine gene expression in CTL019 cells. These activated gene signatures could serve as a reference standard for assessing the function of new CAR-T cell products.

**Fig. 2.**
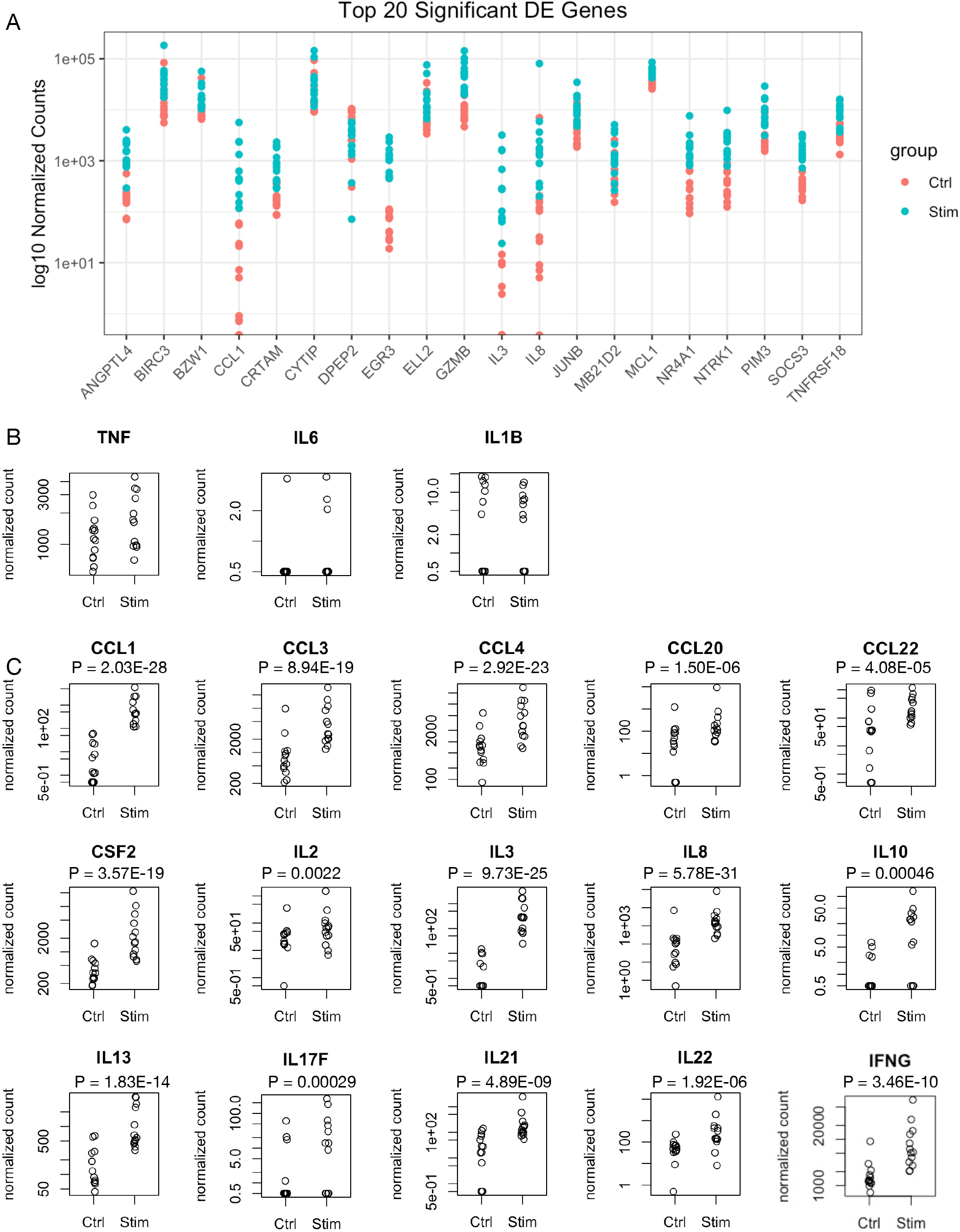
Biomarkers of CAR T cell activation. (A) Top 20 differentially expressed genes after anti-idiotypic antibody stimulation. Data was presented at individual sample level. (B) The expression of cytokine genes TNF, IL6, or IL1B were comparable between control and anti-idiotypic antibody CTL019 cells. The Log2fold change, P value, and adjust P value (padj) for each gene can be found in Table S1. (C) The upregulated cytokine genes in CTL019 cells after anti-idiotypic antibody stimulation.

### Signaling pathways related to severe CRS after in vitro stimulation

CAR T cell therapy is associated with cytokine release syndrome (CRS). The molecular mechanisms that underlying the severity of CRS are not fully understood[1]. According to the original article in Supplementary Table 2[8], the clinical characteristics of responding patients who received CTL019 cell infusion were collected, and the serious adverse events were recorded as grades 0-4. Seven of 13 patients that received CTL019 cell infusion developed moderate to severe CRS with grade 2 to 4 reaction. We next asked whether severe CRS could be predicted before infusion by evaluating the differentially expressed genes in anti-CAR19 antibody treated CTL019 cells in vitro. We re-analyzed the raw RNAseq data using DESeq2 and exported the differentially expressed genes of stimulated CTL019 cells between patients with or without severe CRS.

We identified about 509 differentially expressed genes (Log2fold change > 0.5 and p< 0.01) between severe CRS and without CRS groups (Table S2), suggesting the different characteristics of the CTL019 cells after anti-CAR19 antibody stimulation in vitro (Figure 3A). When compared to patients without CRS, the significantly unregulated genes in the severe CRS group include TCL6 (T cell leukemia/lymphoma 6), HPCAL4 (Hippocalcin-like 4), CCDC144B (coiled-coil domain containing 144B) and SIRPG (signal-regulatory proteins gamma) (Figure 3C). In addition, IL2, IL21, and HSPA1B were downregulated in the severe CRS group. SIRPG is expressed by T cells and may function as an accessory protein in T cell responses[12]. It has been reported that TCL6, a long non-coding RNA, is associated with clinical outcomes in pediatric B-cell acute lymphoblastic leukemia[13]. In addition, a recent study has identified a unique stress response state in tumor-infiltrating T cells, which was characterized by the expression of stress-related heat shock genes including HSPA1B. HSPA1B expression in intratumoral T cells was upregulated following immune checkpoint blockade treatment, especially in nonresponsive tumors, suggesting a role of immunotherapy resistance[14]. GSEA analysis revealed that the differentially expressed genes were enriched in TNF-NFKB, DNA repair, Glycolysis, and MTORC1 signaling pathways (Figure 3B). We further analyzed the correlation between differentially expressed genes and CRS severity. As shown in Figure 3D, the upregulated gene SIRPG is positively correlated with CRS severity. The correlation between down-regulated genes and CRS severity requires further validation with a larger sample size.

**Fig. 3.**
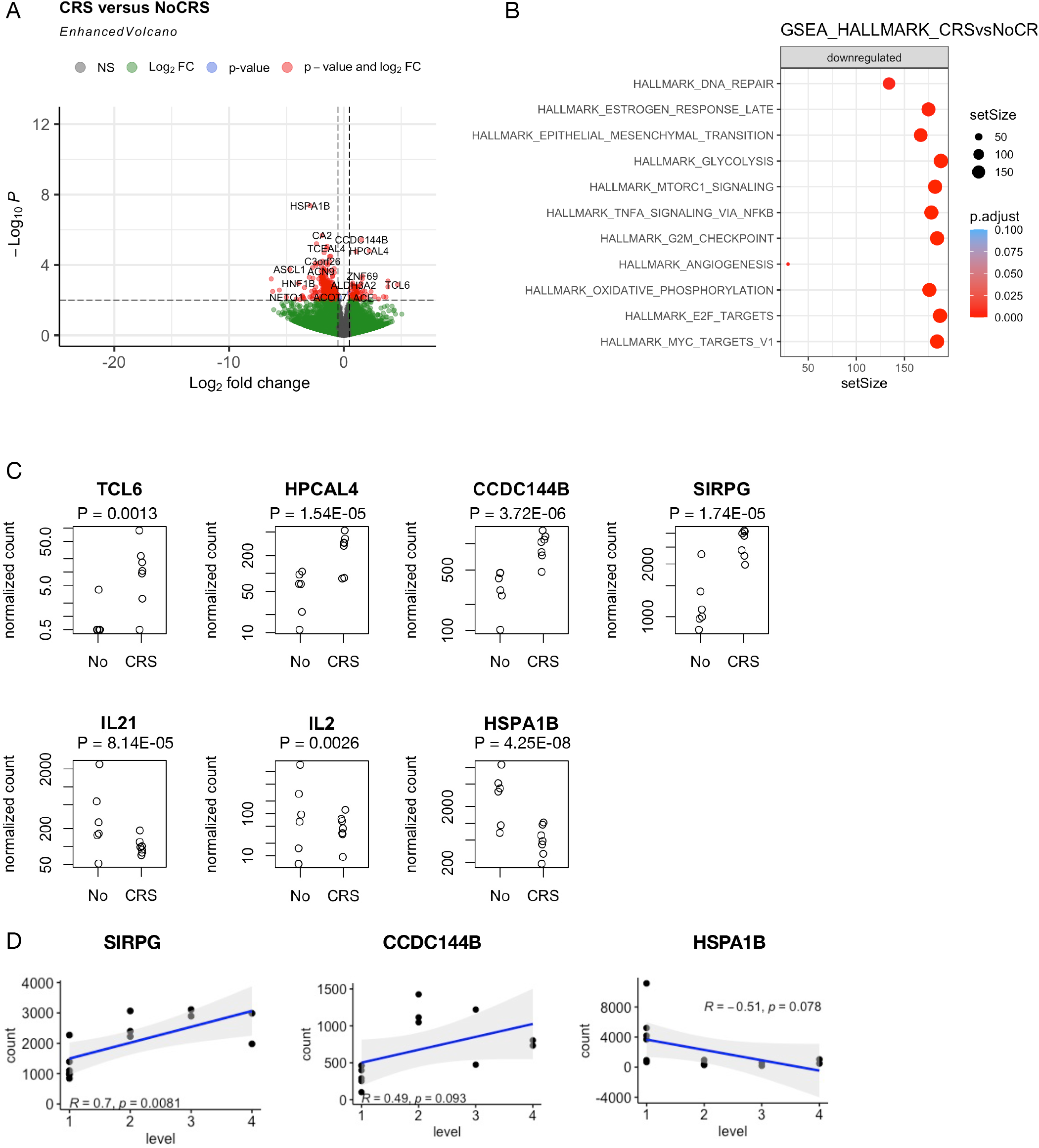
Signaling pathways related to severe CRS after in vitro stimulation. (A) Volcano plot shows differentially expressed genes in CTL019 cells between severe CRS and non-severe CRS patients. (B) Dot plot shows the activated genes in CTL019 cells of severe CRS enriched in signal pathways. (C) Differentially expressed genes in CTL019 cells between severe CRS (CRS) and non-severe CRS (No). Each data point represents an individual sample. (D) Scatter plots show the normalized count and CRS severity levels of differentially expressed genes.

### Intrinsic transcriptomic difference between severe CRS and without CRS

Furthermore, we analyzed the gene expression profile in the control group (without stimulation) to clarify the intrinsic difference between severe CRS and without CRS before CTL019 antibody treatment. We identified about 1509 differentially expressed genes (Log2fold change > 0.5 and p< 0.01), including 247 genes that were expressed higher, and 1262 genes that were lower expressed in the severe CRS group when compared to those without CRS (Figure 4A, Table S3). The significantly higher expressed genes in the severe CRS group included IFNAR1, IL7R, ZNF69 (zinc finger protein 69), and USP32P1 (ubiquitin specific peptidase 32 pseudogene 1). In addition, the expressions of CCL3, IL4, IL17A, IL23R, IL13, CD70, and IFNGR2 were significantly lower (Figure 4B). These results suggested an intrinsic difference in T cells between the patients who develop severe CRS and non-severe CRS. Further study using gene set enrichment analysis didn’t identify an obvious enriched signal pathway except a few genes involved in estrogen response, oxidative phosphorylation, E2F, and MYC pathways (Figure 4C).

**Fig. 4.**
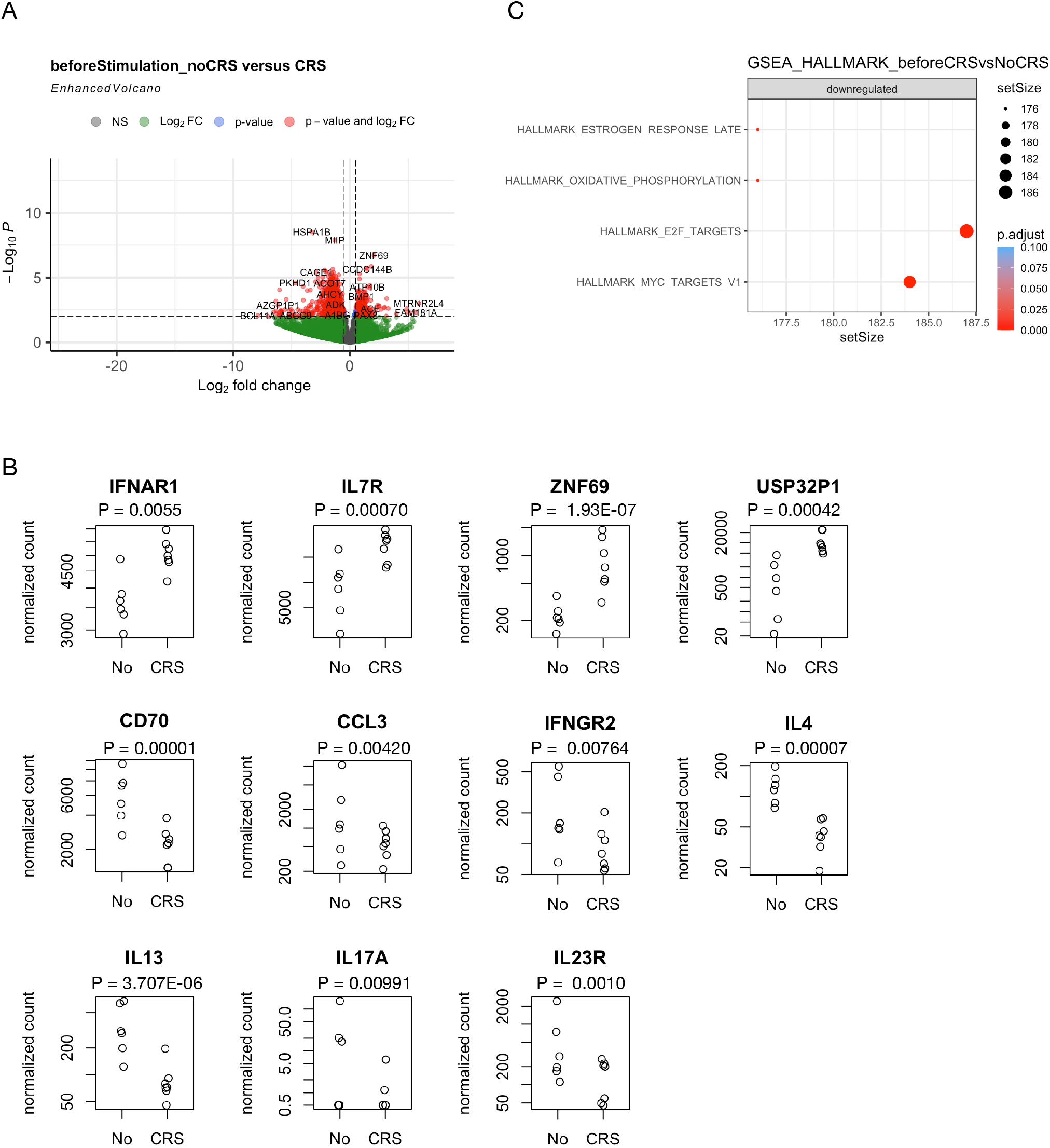
Intrinsic transcriptomic difference between severe CRS and without CRS. (A) Volcano plot shows differentially expressed genes in CTL019 cells prior to stimulation. (B) Plots show the differentially expressed genes in CTL019 cells without stimulation, suggesting the intrinsic difference between CTL019 cells of non-severe CRS (No) and severe CRS (CRS) patients. (C) Dot plot shows the enriched signal pathways of differentially expressed genes in CTL019 cells without stimulation. Each data point represents an individual sample.

## Discussion

Chimeric antigen receptor (CAR) T cell therapy has achieved astonishing remission in patients with B cell malignancies[1]. Since CAR T cells are engineered to express fusion proteins that can be activated by CD19 present on malignant B cells to generate an anti-tumor immune response, all approved CAR T cell products share the common adverse effects including cytokine-release syndrome (CRS) and immune effector cell-associated neurotoxicity syndrome (ICANS)[3, 6].

Approximately 30%–50% of the patients require special intensive care units (ICUs), additional diagnostics, and medications to manage toxicities[15]. Previous studies have shown biomarkers that could predict life-threatening events post infusion, these factors include CAR T cell dose, fever, tumor burden, and platelet count. Patients who progress to severe CRS typically show symptoms within 36 hours post infusion, and fever often be reported as the first symptom[5]. Overactivation of the immune system is the direct consequence of CRS, which leads to elevated serum cytokines. The blood level of cytokines on day 1 post infusion, such as IFN-γ, IL-6, IL-8, IL-10, etc., correlates with the severity of both CRS and ICANS[5]. While these biomarkers are sensitive and specific, the results are not available until day 1 or later post infusion. To support the design of CAR T cell therapy plan, the ideal predicting biomarkers would be applied before T cell infusion.

By re-analyzing the RNAseq dataset, we identified a gene expression signature that was activated by anti-CAR19 idiotypic antibody stimulation. The genes activated by in vitro stimulation are predominantly involved in TNF-NFKB, IL2-Stat5, IL6-JAK-Stat3, and interferon alpha and gamma pathways, which in consistent with findings from previous clinical data[16].

Recent review papers by Tedesco et al[5]. and Larson et al[16]. have summarized cytokine activation patterns in clinical settings, which largely overlap with those observed in in vitro stimulated CAR-T cells in Figure 2. Therefore, this set of upregulated genes, including IFNG, IL8, IL10 etc., could serve as a reference for evaluating the function of CAR-T cell products. In addition, elevated levels of cytokines such as IL6, CCL2, and CXCL10 in the cerebrospinal fluid (CSF) are correlated with severe neurotoxicity[16].

We observed that anti-CAR19 idiotypic antibody stimulated a higher expression of TCL6, HPCAL4, CCDC144B, and SIRPG in T cells of severe CRS patients. Even though these genes do not belong to the inflammatory cytokine families, they are directly involved in the T cell immune function. The interaction between SIRPγ and CD47 plays a key role in T-cell transendothelial migration[17]. SIRPγ-CD47 interaction positively regulates the activation of human T cells in situation of chronic stimulation[18]. In human lung adenocarcinoma, SIRPG was upregulated and its overexpression predicted poor survival outcome. SIRPγ bridged MST1 and PP2A to activate Hippo/YAP signal, and lead to cytokine release by CSLCs, which stimulated CD47 expression in lung adenocarcinoma cells and consequently inhibited tumor cell phagocytosis[19]. We also observed an intrinsic difference in T cell gene expression between the patients who develop severe CRS and non-severe CRS.

Together, our study provided new insights into evaluating CAR-T cell product function in vitro, and the gene expression signature present here could support the research of predicting severe CRS events before infusion. One of the limitations of our study is the relatively low number of patients included in the comparison between severe CRS and non-CRS (7 vs 6), further studies need to combine more samples and multi-module data to validate these observations.

## Methods

Patient samples. Patient characteristics were obtained from the original article [8] Supplementary Table 2 “Treatment and clinical characteristics of responding patients”. CTL019 cell products from 13 chronic lymphocytic leukemia patients were analyzed in this study [8]. For bulk RNAseq analysis, the cell samples were cultured overnight with an anti-idiotypic antibody or isotype-control antibody as described.

RNAseq data analysis. The gene expression count table was acquired from the original article [8] Supplementary Table 5(a and b). Data analysis was carried out using R programming. Differentially expressed genes between groups were analyzed using DEseq2. Gene Set Enrichment Analysis (GSEA) was conducted using the Molecular Signatures Database (MSigDB). Correlation analysis was conducted using ggpubr package.

## Abbreviations

CRS: Cytokine release syndrome
CAR-T: Chimeric antigen receptor T cell
ICANS: Immune effector cell-associated neurotoxicity syndrome
CR: Complete remission
NR: nonresponding
CTL019: CD19-targeted T cells
GSEA: Gene Set Enrichment Analysis
MSigDB: Signatures Database

## Declarations

### Ethics approval and consent to participate

This study was conducted in accordance with the principles of the Declaration of Helsinki. The data related to human participants is from previously published work, so the consent to participate and ethics approval were waived by the ethics committee of Guizhou Medical University.

### Consent for publication

All authors consent to the publication of this work

### Availability of data and material

The original files including the transcriptomic profiling of mock-stimulated (control) and CAR-stimulated CTL019 infusion products are available for public access in paper [8]. The processed data set will be made available upon request.

### Author contributions

MX.C, F.OY, and YH.H designed research. MX.C and F.OY performed the RNA-Seq data analysis. MX.C, FO.Y, and MF.C prepared the manuscript, MX. C, F. OY, MF. C, TW, YF.H, QL.L, JZ, YD, reviewed and edited the manuscript. F. OY and YH. H contributed equally to supervising the work. All authors read and approved the final manuscript

## Acknowledgments

We appreciate Dr. Carl H. June and Dr. J. Joseph Melenhorst for their pioneer work and for making this data set publicly available.

## Table captions

Table S1. Differentially expressed genes between anti-CAR19 antibody treated CTL019 cells and untreated control.

Table S2. Differentially expressed genes between severe CRS and without CRS groups after CTL019 antibody treatment.

Table S3. Differentially expressed genes between severe CRS group and without CRS before CTL019 antibody treatment.

