## Supplementary material for "Marker genes for predicting cytokine release syndrome in vitro before CAR T cell infusion": Table S1

| gene | baseMean | log2FoldChange | lfcSE | stat | pvalue | padj |
| --- | --- | --- | --- | --- | --- | --- |
| IL3 | 330.7218 | 8.4918 | 0.8269 | 10.2689 | 9.73E-25 | 7.74E-22 |
| CCL1 | 568.8872 | 7.5469 | 0.6825 | 11.0571 | 2.03E-28 | 4.38E-25 |
| XIRP1 | 71.0775 | 5.6278 | 0.5965 | 9.4345 | 3.93E-21 | 1.52E-18 |
| COL6A5 | 21.4818 | 5.6256 | 0.8191 | 6.8680 | 6.51E-12 | 6.00E-10 |
| ECEL1 | 8.9485 | 5.3015 | 1.3576 | 3.9049 | 9.43E-05 | 0.00120113 |
| SPRY4 | 12.7890 | 4.9439 | 1.1574 | 4.2717 | 1.94E-05 | 0.000325394 |
| IL8 | 4214.3111 | 4.7298 | 0.4088 | 11.5711 | 5.78E-31 | 1.75E-27 |
| BOC | 19.0712 | 4.6969 | 0.9472 | 4.9588 | 7.09E-07 | 1.97E-05 |
| SNCB | 10.0069 | 4.4829 | 1.0417 | 4.3034 | 1.68E-05 | 0.000290123 |
| PNOC | 46.0436 | 4.3069 | 0.7853 | 5.4841 | 4.15E-08 | 1.71E-06 |
| DACT2 | 15.5407 | 4.1134 | 1.2742 | 3.2281 | 0.001245945 | 0.009523343 |
| EGR3 | 622.1643 | 3.9702 | 0.3717 | 10.6803 | 1.26E-26 | 1.59E-23 |
| SEPT5-GP1BB | 7.7572 | 3.9598 | 1.0594 | 3.7378 | 0.000185626 | 0.00208615 |
| MB | 34.0692 | 3.9174 | 0.7508 | 5.2175 | 1.81E-07 | 6.08E-06 |
| UBE2QL1 | 322.5610 | 3.9002 | 0.4825 | 8.0828 | 6.33E-16 | 1.29E-13 |
| ACSM2A | 15.5527 | 3.8810 | 0.9844 | 3.9425 | 8.06E-05 | 0.001059567 |
| CEACAM16 | 7.3331 | 3.8474 | 1.2564 | 3.0623 | 0.002196495 | 0.014900797 |
| PPM1J | 17.3538 | 3.8411 | 0.8352 | 4.5991 | 4.24E-06 | 9.12E-05 |
| SH3RF3 | 7.7556 | 3.8340 | 1.0330 | 3.7114 | 0.000206112 | 0.00226588 |
| FAM13C | 7.8737 | 3.6909 | 1.2825 | 2.8780 | 0.00400197 | 0.02383439 |
| PTGS2 | 822.2153 | 3.6700 | 0.3889 | 9.4360 | 3.87E-21 | 1.52E-18 |
| FAM131C | 9.6239 | 3.6314 | 1.3071 | 2.7782 | 0.005466372 | 0.030338773 |
| LOC100128993 | 20.2459 | 3.6174 | 0.9679 | 3.7375 | 0.000185852 | 0.002087142 |
| HEYL | 13.6077 | 3.5968 | 1.1644 | 3.0889 | 0.002009075 | 0.013896329 |
| IL10 | 13.4477 | 3.5750 | 1.0206 | 3.5028 | 0.000460383 | 0.004312211 |
| CD276 | 81.2366 | 3.4940 | 0.5759 | 6.0668 | 1.31E-09 | 7.62E-08 |
| MST1R | 16.0303 | 3.4703 | 0.9059 | 3.8308 | 0.000127734 | 0.001548265 |
| DMBX1 | 15.3741 | 3.4540 | 0.7822 | 4.4156 | 1.01E-05 | 0.000187912 |
| XCL2 | 151.8488 | 3.4312 | 0.3809 | 9.0069 | 2.12E-19 | 7.29E-17 |
| RIMS1 | 16.6200 | 3.4293 | 0.9908 | 3.4611 | 0.000537885 | 0.004866383 |
| RASAL1 | 142.3409 | 3.4154 | 0.6660 | 5.1284 | 2.92E-07 | 9.25E-06 |
| IL21 | 193.8827 | 3.3734 | 0.5766 | 5.8509 | 4.89E-09 | 2.51E-07 |
| GLI2 | 8.8738 | 3.3574 | 1.2999 | 2.5828 | 0.009798859 | 0.047146098 |
| INSM1 | 16.4976 | 3.2884 | 0.9425 | 3.4890 | 0.00048486 | 0.004488897 |
| CA12 | 76.9792 | 3.2741 | 0.6056 | 5.4062 | 6.44E-08 | 2.49E-06 |
| XCL1 | 398.7239 | 3.2592 | 0.4066 | 8.0166 | 1.09E-15 | 2.16E-13 |
| IL17F | 27.5197 | 3.2302 | 0.8927 | 3.6185 | 0.000296273 | 0.003030242 |
| SNAP25 | 367.1717 | 3.2190 | 0.5455 | 5.9012 | 3.61E-09 | 1.91E-07 |
| EGR2 | 759.0368 | 3.2033 | 0.4467 | 7.1717 | 7.41E-13 | 8.49E-11 |
| GEM | 967.8461 | 3.1455 | 0.4770 | 6.5938 | 4.29E-11 | 3.43E-09 |
| WFDC5 | 70.5517 | 3.1285 | 0.5404 | 5.7890 | 7.08E-09 | 3.56E-07 |
| GNAI1 | 87.9315 | 3.1281 | 0.5599 | 5.5872 | 2.31E-08 | 1.04E-06 |
| COL2A1 | 10.6481 | 3.1078 | 1.0986 | 2.8288 | 0.004672314 | 0.026976369 |
| MPP2 | 13.3677 | 3.1037 | 0.9007 | 3.4460 | 0.000569 | 0.005096068 |
| C9orf131 | 8.5139 | 3.0739 | 1.0915 | 2.8162 | 0.004860127 | 0.027861803 |
| IFITM5 | 27.2378 | 3.0509 | 0.7083 | 4.3074 | 1.65E-05 | 0.000286202 |
| LOC286002 | 17.7282 | 3.0361 | 0.7934 | 3.8268 | 0.000129838 | 0.001571242 |
| STC1 | 77.2523 | 3.0226 | 0.6037 | 5.0070 | 5.53E-07 | 1.60E-05 |
| GUCY1B3 | 16.2967 | 3.0130 | 0.7084 | 4.2532 | 2.11E-05 | 0.000346839 |
| GRB7 | 5.6874 | 2.9961 | 1.0371 | 2.8888 | 0.003866646 | NA |
| C2CD4A | 32.7952 | 2.9911 | 0.8818 | 3.3921 | 0.000693503 | 0.005960577 |
| FLJ41278 | 19.0215 | 2.9833 | 0.7226 | 4.1285 | 3.65E-05 | 0.000548562 |
| OLIG1 | 9.6401 | 2.9560 | 0.9337 | 3.1658 | 0.001546478 | 0.01127401 |
| CCL4L1 | 14.9139 | 2.9221 | 0.8922 | 3.2753 | 0.001055451 | 0.008345955 |
| OVOL1 | 17.7514 | 2.8987 | 1.0305 | 2.8129 | 0.004910221 | 0.028035971 |
| CLDN1 | 12.8645 | 2.8983 | 1.0670 | 2.7164 | 0.006600498 | 0.035021303 |
| IL19 | 12.7605 | 2.8440 | 0.9812 | 2.8983 | 0.003751353 | 0.022635306 |
| IL17REL | 66.8156 | 2.8413 | 0.5883 | 4.8298 | 1.37E-06 | 3.43E-05 |
| FAM155B | 8.1872 | 2.8256 | 1.0826 | 2.6099 | 0.009056297 | 0.04432048 |
| ACSM2B | 10.1957 | 2.7940 | 1.0727 | 2.6047 | 0.009195131 | 0.044854805 |
| HS6ST2 | 27.4739 | 2.7446 | 0.5836 | 4.7027 | 2.57E-06 | 5.89E-05 |
| DACT3 | 36.6122 | 2.7427 | 0.6036 | 4.5435 | 5.53E-06 | 0.000115423 |
| BTNL8 | 57.0721 | 2.7425 | 0.7689 | 3.5666 | 0.000361594 | 0.003545272 |
| HES4 | 25.2018 | 2.7392 | 0.7443 | 3.6802 | 0.000233052 | 0.002487915 |
| IL22 | 694.9836 | 2.7318 | 0.5737 | 4.7614 | 1.92E-06 | 4.57E-05 |
| TNS3 | 359.8852 | 2.7314 | 0.4395 | 6.2155 | 5.12E-10 | 3.21E-08 |
| SSTR2 | 83.3756 | 2.6642 | 0.5102 | 5.2220 | 1.77E-07 | 5.96E-06 |
| NR4A3 | 1376.2701 | 2.6638 | 0.2806 | 9.4947 | 2.21E-21 | 9.03E-19 |
| FREM2 | 14.3475 | 2.6547 | 0.8339 | 3.1833 | 0.001455887 | 0.010753759 |
| MUC16 | 20.3849 | 2.6415 | 0.8590 | 3.0752 | 0.002103565 | 0.014391959 |
| CCL22 | 253.3632 | 2.6366 | 0.6426 | 4.1029 | 4.08E-05 | 0.000603317 |
| ACHE | 16.0823 | 2.6226 | 0.8342 | 3.1438 | 0.00166765 | 0.011961377 |
| TMEM145 | 39.6331 | 2.6146 | 0.5627 | 4.6468 | 3.37E-06 | 7.41E-05 |
| IL13 | 816.5152 | 2.6123 | 0.3409 | 7.6620 | 1.83E-14 | 2.92E-12 |
| BIRC7 | 31.6071 | 2.5919 | 0.6851 | 3.7834 | 0.000154698 | 0.001786348 |
| CCL20 | 512.7334 | 2.5558 | 0.5313 | 4.8107 | 1.50E-06 | 3.71E-05 |
| TWIST1 | 127.8445 | 2.5540 | 0.3682 | 6.9372 | 4.00E-12 | 3.95E-10 |
| ISM2 | 12.5951 | 2.5457 | 0.8307 | 3.0646 | 0.002179286 | 0.014842888 |
| FGF12 | 25.0466 | 2.5434 | 0.7658 | 3.3212 | 0.000896339 | 0.007337079 |
| CSF2 | 2360.8950 | 2.5399 | 0.2838 | 8.9494 | 3.57E-19 | 1.15E-16 |
| ANGPTL4 | 1007.0446 | 2.5356 | 0.2463 | 10.2944 | 7.47E-25 | 6.28E-22 |
| ADRA2B | 14.0922 | 2.5353 | 0.7679 | 3.3018 | 0.000960585 | 0.007741489 |
| FMOD | 24.7728 | 2.5329 | 0.8019 | 3.1585 | 0.001585613 | 0.011492842 |
| MFSD2A | 134.8053 | 2.4830 | 0.2944 | 8.4333 | 3.36E-17 | 8.48E-15 |
| NOTUM | 34.4543 | 2.4748 | 0.7515 | 3.2933 | 0.000990073 | 0.007936348 |
| PAX5 | 60.4703 | 2.4630 | 0.5264 | 4.6788 | 2.89E-06 | 6.53E-05 |
| HES2 | 25.3209 | 2.4305 | 0.7843 | 3.0988 | 0.001943237 | 0.013521318 |
| KHDRBS3 | 29.2262 | 2.4093 | 0.8609 | 2.7985 | 0.005134083 | 0.029043857 |
| HPCA | 19.2725 | 2.4088 | 0.8225 | 2.9286 | 0.003404929 | 0.020912044 |
| DTNA | 39.0073 | 2.4027 | 0.6682 | 3.5955 | 0.000323744 | 0.003259574 |
| CCL4 | 3812.0372 | 2.3946 | 0.2410 | 9.9355 | 2.92E-23 | 1.70E-20 |
| THSD7A | 79.9321 | 2.3859 | 0.4910 | 4.8588 | 1.18E-06 | 3.04E-05 |
| CCL3 | 4143.0280 | 2.3841 | 0.2695 | 8.8476 | 8.94E-19 | 2.65E-16 |
| LRRC32 | 94.7383 | 2.3829 | 0.4271 | 5.5787 | 2.42E-08 | 1.08E-06 |
| CACNA1E | 15.3664 | 2.3811 | 0.7340 | 3.2439 | 0.001178986 | 0.009089971 |
| FLJ22184 | 10.2979 | 2.3642 | 0.9131 | 2.5891 | 0.009623158 | 0.04656734 |
| SH3GL3 | 27.7219 | 2.3183 | 0.7014 | 3.3052 | 0.000949127 | 0.007665477 |
| IFI44L | 1743.6363 | 2.3126 | 0.4139 | 5.5874 | 2.31E-08 | 1.04E-06 |
| CDH1 | 25.6003 | 2.3083 | 0.6627 | 3.4835 | 0.000494937 | 0.004551313 |
| NR4A2 | 483.8495 | 2.2791 | 0.2242 | 10.1636 | 2.88E-24 | 2.08E-21 |
| ERC2 | 36.2965 | 2.2784 | 0.6741 | 3.3799 | 0.000725163 | 0.006186991 |
| TEAD4 | 98.6107 | 2.2746 | 0.5999 | 3.7918 | 0.000149531 | 0.001735963 |
| TSKU | 169.4111 | 2.2741 | 0.4115 | 5.5264 | 3.27E-08 | 1.39E-06 |
| NR4A1 | 1316.7425 | 2.2729 | 0.2178 | 10.4338 | 1.74E-25 | 1.64E-22 |
| GZMB | 32600.5804 | 2.2708 | 0.2159 | 10.5164 | 7.26E-26 | 7.32E-23 |
| FLT1 | 417.0517 | 2.2518 | 0.2897 | 7.7728 | 7.68E-15 | 1.25E-12 |
| SLC30A3 | 42.4307 | 2.2364 | 0.6372 | 3.5099 | 0.000448341 | 0.004225581 |
| IRX3 | 67.2689 | 2.2333 | 0.4937 | 4.5233 | 6.09E-06 | 0.000123594 |
| NTRK1 | 1608.8697 | 2.2230 | 0.2034 | 10.9308 | 8.22E-28 | 1.38E-24 |
| COL5A1 | 22.9509 | 2.2210 | 0.7195 | 3.0870 | 0.002022212 | 0.01396804 |
| IGLON5 | 118.7976 | 2.2165 | 0.4503 | 4.9223 | 8.55E-07 | 2.32E-05 |
| LAMC1 | 35.9131 | 2.1932 | 0.6036 | 3.6333 | 0.000279789 | 0.002890959 |
| PVRL4 | 10.5277 | 2.1918 | 0.8151 | 2.6891 | 0.007163918 | 0.037137967 |
| SOCS3 | 1077.3809 | 2.1872 | 0.1884 | 11.6113 | 3.61E-31 | 1.37E-27 |
| TNFRSF8 | 933.0784 | 2.1752 | 0.3919 | 5.5512 | 2.84E-08 | 1.24E-06 |
| ZNF860 | 41.1082 | 2.1635 | 0.7589 | 2.8506 | 0.004363265 | 0.025483825 |
| IRF8 | 92.4741 | 2.1615 | 0.3923 | 5.5092 | 3.61E-08 | 1.51E-06 |
| PMCH | 374.2960 | 2.1577 | 0.3149 | 6.8512 | 7.33E-12 | 6.72E-10 |
| NUDT10 | 22.0992 | 2.1534 | 0.8171 | 2.6355 | 0.00840159 | 0.041861281 |
| IFNG | 7372.0087 | 2.1153 | 0.3370 | 6.2764 | 3.47E-10 | 2.26E-08 |
| CCDC3 | 27.5399 | 2.1078 | 0.6986 | 3.0170 | 0.00255315 | 0.016654377 |
| BARX1 | 91.4735 | 2.1032 | 0.4812 | 4.3708 | 1.24E-05 | 0.000222916 |
| LOC286442 | 234.0303 | 2.0767 | 0.3633 | 5.7169 | 1.08E-08 | 5.18E-07 |
| MET | 85.3245 | 2.0680 | 0.4679 | 4.4196 | 9.89E-06 | 0.00018561 |
| CRTAM | 642.5276 | 2.0584 | 0.1901 | 10.8284 | 2.52E-27 | 3.82E-24 |
| MYH6 | 59.9442 | 2.0557 | 0.6052 | 3.3969 | 0.000681571 | 0.005871368 |
| LRP2 | 86.9750 | 2.0532 | 0.4305 | 4.7696 | 1.85E-06 | 4.43E-05 |
| ARHGAP39 | 28.6475 | 2.0391 | 0.7053 | 2.8912 | 0.003837629 | 0.023036435 |
| PDE4C | 43.2505 | 2.0321 | 0.6129 | 3.3155 | 0.000914913 | 0.007436805 |
| RGS16 | 325.0099 | 2.0223 | 0.4684 | 4.3180 | 1.57E-05 | 0.00027476 |
| FAIM2 | 27.2494 | 2.0195 | 0.6688 | 3.0198 | 0.002529439 | 0.016535358 |
| PLK2 | 405.4245 | 2.0164 | 0.2272 | 8.8763 | 6.91E-19 | 2.09E-16 |
| APBB2 | 211.7673 | 2.0047 | 0.2378 | 8.4299 | 3.46E-17 | 8.58E-15 |
| IL5 | 94.1723 | 1.9934 | 0.4063 | 4.9058 | 9.31E-07 | 2.49E-05 |
| OSMR | 45.9373 | 1.9633 | 0.4888 | 4.0169 | 5.90E-05 | 0.000822203 |
| TMEM200B | 30.5527 | 1.9623 | 0.6647 | 2.9521 | 0.003156431 | 0.019755502 |
| LRRC43 | 25.9726 | 1.9518 | 0.7502 | 2.6019 | 0.009271941 | 0.045101427 |
| ATP12A | 34.1243 | 1.9404 | 0.5783 | 3.3552 | 0.000793084 | 0.00663917 |
| KIAA0226L | 87.4750 | 1.9149 | 0.3469 | 5.5206 | 3.38E-08 | 1.43E-06 |
| GJC1 | 31.9964 | 1.9052 | 0.7222 | 2.6379 | 0.008341282 | 0.041637833 |
| PROS1 | 81.1822 | 1.9012 | 0.4597 | 4.1355 | 3.54E-05 | 0.000535774 |
| TSPAN13 | 612.2294 | 1.8947 | 0.2417 | 7.8401 | 4.50E-15 | 7.74E-13 |
| HLX | 109.3616 | 1.8936 | 0.4324 | 4.3792 | 1.19E-05 | 0.000216056 |
| EMP1 | 377.6925 | 1.8917 | 0.2426 | 7.7979 | 6.29E-15 | 1.06E-12 |
| EGR1 | 989.4871 | 1.8850 | 0.1897 | 9.9352 | 2.92E-23 | 1.70E-20 |
| ATRNL1 | 53.6315 | 1.8822 | 0.5993 | 3.1403 | 0.001687489 | 0.012097939 |
| ADM | 998.1913 | 1.8688 | 0.2239 | 8.3477 | 6.96E-17 | 1.64E-14 |
| ADAMTS14 | 494.3775 | 1.8303 | 0.2565 | 7.1355 | 9.65E-13 | 1.08E-10 |
| CABP1 | 85.1049 | 1.8252 | 0.4417 | 4.1320 | 3.60E-05 | 0.000542346 |
| GLDC | 365.6525 | 1.8190 | 0.4356 | 4.1753 | 2.98E-05 | 0.000461643 |
| DOCK6 | 59.0834 | 1.8183 | 0.4601 | 3.9516 | 7.76E-05 | 0.001028987 |
| EHD2 | 37.8772 | 1.8167 | 0.5582 | 3.2547 | 0.001134978 | 0.008813558 |
| ATP9A | 2082.8113 | 1.8069 | 0.2749 | 6.5735 | 4.91E-11 | 3.83E-09 |
| OR9Q1 | 46.7432 | 1.8066 | 0.5638 | 3.2045 | 0.001352983 | 0.010131964 |
| TNFRSF9 | 2308.5095 | 1.8065 | 0.1832 | 9.8621 | 6.07E-23 | 3.27E-20 |
| CAMK2B | 50.2501 | 1.8045 | 0.5240 | 3.4438 | 0.000573543 | 0.005122081 |
| AQPEP | 54.4863 | 1.8006 | 0.4465 | 4.0327 | 5.51E-05 | 0.000775277 |
| C2orf72 | 375.6624 | 1.7950 | 0.2406 | 7.4618 | 8.53E-14 | 1.16E-11 |
| DOK5 | 176.4151 | 1.7938 | 0.6317 | 2.8396 | 0.004517379 | 0.02621321 |
| GALNTL4 | 260.0937 | 1.7884 | 0.3506 | 5.1008 | 3.38E-07 | 1.05E-05 |
| NCS1 | 225.1177 | 1.7798 | 0.2570 | 6.9254 | 4.35E-12 | 4.27E-10 |
| GNG8 | 86.1576 | 1.7775 | 0.3730 | 4.7662 | 1.88E-06 | 4.48E-05 |
| SP5 | 24.3265 | 1.7745 | 0.6580 | 2.6967 | 0.00700391 | 0.036521252 |
| LGALS17A | 54.9697 | 1.7736 | 0.5119 | 3.4646 | 0.000530963 | 0.00481253 |
| LYPD6B | 24.3616 | 1.7692 | 0.5816 | 3.0417 | 0.002352182 | 0.015716194 |
| TMPRSS6 | 4389.7480 | 1.7655 | 0.2982 | 5.9198 | 3.22E-09 | 1.72E-07 |
| C19orf26 | 37.9073 | 1.7654 | 0.6067 | 2.9099 | 0.003615246 | 0.021982649 |
| COL6A3 | 1758.5480 | 1.7580 | 0.4540 | 3.8723 | 0.00010783 | 0.001334812 |
| ADD2 | 54.0654 | 1.7528 | 0.5244 | 3.3422 | 0.000831191 | 0.006878239 |
| IL4I1 | 595.9405 | 1.7483 | 0.2642 | 6.6173 | 3.66E-11 | 3.01E-09 |
| PHF21B | 26.6411 | 1.7451 | 0.6105 | 2.8586 | 0.004254932 | 0.025025024 |
| CTTN | 247.7484 | 1.7399 | 0.5119 | 3.3986 | 0.000677362 | 0.005848433 |
| SHF | 782.8268 | 1.7369 | 0.1755 | 9.8947 | 4.39E-23 | 2.46E-20 |
| GPR133 | 344.9090 | 1.7365 | 0.2496 | 6.9583 | 3.44E-12 | 3.43E-10 |
| IL1RN | 307.6102 | 1.7324 | 0.3557 | 4.8700 | 1.12E-06 | 2.89E-05 |
| PDGFA | 407.7123 | 1.7262 | 0.2346 | 7.3589 | 1.85E-13 | 2.38E-11 |
| PLOD2 | 97.1748 | 1.7259 | 0.4315 | 3.9995 | 6.35E-05 | 0.000871475 |
| P2RY1 | 103.7492 | 1.7183 | 0.4154 | 4.1362 | 3.53E-05 | 0.000534707 |
| PTGIS | 353.9530 | 1.7182 | 0.2593 | 6.6265 | 3.44E-11 | 2.84E-09 |
| LHFP | 490.9294 | 1.7157 | 0.2435 | 7.0470 | 1.83E-12 | 1.88E-10 |
| RASGEF1B | 263.6927 | 1.7002 | 0.2039 | 8.3395 | 7.46E-17 | 1.71E-14 |
| RRAD | 149.5589 | 1.6977 | 0.3519 | 4.8240 | 1.41E-06 | 3.51E-05 |
| MPO | 39.5691 | 1.6898 | 0.5228 | 3.2325 | 0.001226967 | 0.009402395 |
| IL11 | 18.9335 | 1.6860 | 0.5555 | 3.0348 | 0.002407005 | 0.015983653 |
| ARVCF | 132.3277 | 1.6524 | 0.4630 | 3.5694 | 0.000357851 | 0.003519648 |
| KCNG1 | 90.9838 | 1.6482 | 0.4651 | 3.5435 | 0.000394875 | 0.003802208 |
| FAM81A | 28.1857 | 1.6400 | 0.5587 | 2.9356 | 0.003328736 | 0.020560959 |
| GPRC5C | 74.4700 | 1.6243 | 0.3466 | 4.6868 | 2.78E-06 | 6.31E-05 |
| BCL6 | 2080.8053 | 1.6147 | 0.2030 | 7.9553 | 1.79E-15 | 3.26E-13 |
| DGKI | 1690.2734 | 1.6087 | 0.1793 | 8.9706 | 2.95E-19 | 9.70E-17 |
| HRH2 | 610.9297 | 1.6060 | 0.2533 | 6.3408 | 2.29E-10 | 1.55E-08 |
| OSM | 402.2684 | 1.6026 | 0.3154 | 5.0806 | 3.76E-07 | 1.15E-05 |
| SHC4 | 776.7775 | 1.5995 | 0.2136 | 7.4892 | 6.93E-14 | 9.80E-12 |
| GBP1P1 | 37.1019 | 1.5989 | 0.6066 | 2.6359 | 0.008390917 | 0.04183566 |
| LOC116437 | 60.7594 | 1.5845 | 0.5354 | 2.9593 | 0.003082895 | 0.019390831 |
| KCNK13 | 18.6565 | 1.5839 | 0.5731 | 2.7639 | 0.005711164 | 0.031290393 |
| IL1RAP | 3567.8546 | 1.5835 | 0.1738 | 9.1129 | 8.02E-20 | 2.89E-17 |
| DTX1 | 36.3148 | 1.5739 | 0.5976 | 2.6335 | 0.008450688 | 0.042011869 |
| COL27A1 | 217.5287 | 1.5686 | 0.3329 | 4.7116 | 2.46E-06 | 5.66E-05 |
| CACNA1D | 390.4368 | 1.5667 | 0.4393 | 3.5660 | 0.00036249 | 0.003551419 |
| LAYN | 442.5612 | 1.5597 | 0.2734 | 5.7050 | 1.16E-08 | 5.52E-07 |
| IER3 | 2072.6639 | 1.5555 | 0.1692 | 9.1941 | 3.78E-20 | 1.39E-17 |
| IL2RA | 23429.0765 | 1.5525 | 0.2087 | 7.4391 | 1.01E-13 | 1.37E-11 |
| CA9 | 153.5002 | 1.5383 | 0.3368 | 4.5675 | 4.94E-06 | 0.000104023 |
| NDNF | 204.0099 | 1.5340 | 0.4307 | 3.5613 | 0.00036908 | 0.003596247 |
| PRODH | 147.5583 | 1.5340 | 0.4167 | 3.6812 | 0.000232097 | 0.002479473 |
| BIRC3 | 32882.8253 | 1.5293 | 0.1395 | 10.9658 | 5.58E-28 | 1.06E-24 |
| TEAD1 | 37.5274 | 1.5250 | 0.5373 | 2.8384 | 0.00453428 | 0.02629987 |
| LTA | 1756.0297 | 1.5105 | 0.1993 | 7.5784 | 3.50E-14 | 5.29E-12 |
| DYSF | 151.6987 | 1.5100 | 0.2518 | 5.9964 | 2.02E-09 | 1.13E-07 |
| EEF1A2 | 149.9582 | 1.5097 | 0.3846 | 3.9250 | 8.67E-05 | 0.001116425 |
| SECTM1 | 979.9687 | 1.5060 | 0.1854 | 8.1223 | 4.57E-16 | 9.74E-14 |
| BVES | 118.4602 | 1.5036 | 0.2716 | 5.5356 | 3.10E-08 | 1.34E-06 |
| MAP7D2 | 168.2186 | 1.5032 | 0.3818 | 3.9369 | 8.26E-05 | 0.001075654 |
| PTPRU | 40.5016 | 1.5013 | 0.4678 | 3.2091 | 0.001331464 | 0.010020427 |
| NEURL3 | 237.3128 | 1.4984 | 0.3484 | 4.3003 | 1.71E-05 | 0.000293261 |
| BCL6B | 42.6421 | 1.4864 | 0.4011 | 3.7059 | 0.00021066 | 0.002305823 |
| RAP1GAP | 56.5598 | 1.4801 | 0.3869 | 3.8256 | 0.000130444 | 0.001575958 |
| C15orf48 | 236.7961 | 1.4647 | 0.3513 | 4.1694 | 3.05E-05 | 0.00047245 |
| NRP2 | 455.3753 | 1.4624 | 0.2411 | 6.0643 | 1.33E-09 | 7.71E-08 |
| LOC253573 | 249.1533 | 1.4610 | 0.3292 | 4.4377 | 9.09E-06 | 0.000172585 |
| JUNB | 8318.1573 | 1.4594 | 0.1228 | 11.8857 | 1.40E-32 | 1.02E-28 |
| KIAA1244 | 324.6795 | 1.4542 | 0.2705 | 5.3752 | 7.65E-08 | 2.93E-06 |
| SIK1 | 387.4583 | 1.4485 | 0.2270 | 6.3803 | 1.77E-10 | 1.24E-08 |
| NAB2 | 1597.2882 | 1.4440 | 0.1683 | 8.5789 | 9.58E-18 | 2.68E-15 |
| DUSP4 | 11229.2660 | 1.4435 | 0.2510 | 5.7505 | 8.90E-09 | 4.33E-07 |
| MT3 | 22.5133 | 1.4427 | 0.5221 | 2.7634 | 0.005719435 | 0.0312974 |
| KIAA1274 | 89.1912 | 1.4415 | 0.3553 | 4.0570 | 4.97E-05 | 0.000709725 |
| NEBL | 137.2702 | 1.4406 | 0.4782 | 3.0125 | 0.002590991 | 0.016857599 |
| POU2AF1 | 1382.0035 | 1.4382 | 0.3266 | 4.4037 | 1.06E-05 | 0.000197533 |
| IL1A | 693.8695 | 1.4352 | 0.2543 | 5.6432 | 1.67E-08 | 7.72E-07 |
| CTTNBP2NL | 185.9731 | 1.4316 | 0.4138 | 3.4593 | 0.000541653 | 0.004892923 |
| LAMP3 | 206.8964 | 1.4289 | 0.3039 | 4.7023 | 2.57E-06 | 5.90E-05 |
| FAM40B | 5354.2157 | 1.4286 | 0.1672 | 8.5424 | 1.31E-17 | 3.49E-15 |
| G0S2 | 922.0306 | 1.4251 | 0.2155 | 6.6116 | 3.80E-11 | 3.09E-09 |
| DHRS2 | 39.4457 | 1.4226 | 0.4263 | 3.3370 | 0.000846816 | 0.006984617 |
| UTS2 | 316.0608 | 1.4128 | 0.2100 | 6.7273 | 1.73E-11 | 1.52E-09 |
| TNFRSF18 | 7186.2360 | 1.4073 | 0.1376 | 10.2243 | 1.54E-24 | 1.17E-21 |
| HNF1B | 572.0406 | 1.3991 | 0.3964 | 3.5293 | 0.000416594 | 0.003988573 |
| TNIP3 | 435.2111 | 1.3939 | 0.2138 | 6.5184 | 7.11E-11 | 5.30E-09 |
| USP18 | 205.0305 | 1.3928 | 0.3356 | 4.1505 | 3.32E-05 | 0.000505443 |
| EPB41L4B | 301.9110 | 1.3897 | 0.2946 | 4.7173 | 2.39E-06 | 5.54E-05 |
| PLA2G4C | 658.4101 | 1.3894 | 0.2494 | 5.5705 | 2.54E-08 | 1.13E-06 |
| CASKIN1 | 142.8469 | 1.3892 | 0.2837 | 4.8963 | 9.76E-07 | 2.58E-05 |
| HSPG2 | 376.9741 | 1.3861 | 0.2834 | 4.8911 | 1.00E-06 | 2.64E-05 |
| IGSF23 | 41.1853 | 1.3833 | 0.4343 | 3.1847 | 0.001449055 | 0.010724002 |
| HDAC9 | 187.8621 | 1.3800 | 0.2482 | 5.5606 | 2.69E-08 | 1.18E-06 |
| SPRY1 | 974.1832 | 1.3652 | 0.1855 | 7.3602 | 1.84E-13 | 2.37E-11 |
| IGF2BP3 | 57.1493 | 1.3608 | 0.5257 | 2.5886 | 0.009637896 | 0.046608839 |
| IL23R | 492.1918 | 1.3572 | 0.1882 | 7.2118 | 5.52E-13 | 6.43E-11 |
| PTN | 82.6547 | 1.3547 | 0.4333 | 3.1265 | 0.001768818 | 0.012514926 |
| DUSP5 | 5057.2961 | 1.3542 | 0.1376 | 9.8430 | 7.35E-23 | 3.71E-20 |
| PTGR1 | 94.7288 | 1.3490 | 0.3032 | 4.4489 | 8.63E-06 | 0.000165274 |
| ULBP2 | 75.3359 | 1.3478 | 0.5040 | 2.6742 | 0.007490602 | 0.03850091 |
| RSAD2 | 406.0041 | 1.3474 | 0.2737 | 4.9224 | 8.55E-07 | 2.32E-05 |
| MAGI1 | 327.7444 | 1.3471 | 0.2499 | 5.3895 | 7.07E-08 | 2.71E-06 |
| SLC41A2 | 2257.6565 | 1.3435 | 0.1862 | 7.2160 | 5.36E-13 | 6.28E-11 |
| ENOX1 | 202.3703 | 1.3422 | 0.3308 | 4.0579 | 4.95E-05 | 0.000709174 |
| DPF3 | 60.7975 | 1.3369 | 0.4788 | 2.7919 | 0.005239847 | 0.029433035 |
| HILPDA | 15136.2782 | 1.3359 | 0.2316 | 5.7670 | 8.07E-09 | 3.95E-07 |
| TNFRSF12A | 299.1691 | 1.3355 | 0.1666 | 8.0149 | 1.10E-15 | 2.16E-13 |
| APOD | 78.7527 | 1.3244 | 0.4986 | 2.6563 | 0.007900774 | 0.039958211 |
| SLC26A4 | 126.2486 | 1.3176 | 0.3076 | 4.2842 | 1.83E-05 | 0.000310728 |
| MX1 | 2637.3745 | 1.3125 | 0.3176 | 4.1322 | 3.59E-05 | 0.000542346 |
| CSRP2 | 64.4586 | 1.3079 | 0.4318 | 3.0293 | 0.002451154 | 0.016205686 |
| NYAP1 | 27.1539 | 1.3045 | 0.5039 | 2.5889 | 0.009627891 | 0.046575348 |
| CAV1 | 108.4568 | 1.3034 | 0.4565 | 2.8554 | 0.00429887 | 0.025210704 |
| NAMPT | 25928.6026 | 1.3007 | 0.1355 | 9.6016 | 7.87E-22 | 3.40E-19 |
| PLA2G4A | 298.3637 | 1.2977 | 0.4765 | 2.7235 | 0.006459791 | 0.034468166 |
| C7orf57 | 70.5563 | 1.2961 | 0.3673 | 3.5284 | 0.000418119 | 0.003995503 |
| BCL3 | 2936.2501 | 1.2911 | 0.1310 | 9.8590 | 6.26E-23 | 3.27E-20 |
| ZBTB32 | 1892.2436 | 1.2821 | 0.1750 | 7.3271 | 2.35E-13 | 2.96E-11 |
| ZNF215 | 179.1175 | 1.2818 | 0.3756 | 3.4128 | 0.000642888 | 0.005611123 |
| TTBK1 | 240.4995 | 1.2672 | 0.3161 | 4.0089 | 6.10E-05 | 0.000843448 |
| C3orf52 | 169.0822 | 1.2670 | 0.2003 | 6.3251 | 2.53E-10 | 1.70E-08 |
| LOC653075 | 31.7689 | 1.2664 | 0.4287 | 2.9539 | 0.003138002 | 0.019674539 |
| CD200 | 784.4098 | 1.2636 | 0.4288 | 2.9467 | 0.003211681 | 0.020009513 |
| SLC12A8 | 143.3264 | 1.2614 | 0.2979 | 4.2336 | 2.30E-05 | 0.000371229 |
| DLL1 | 30.5723 | 1.2565 | 0.4506 | 2.7885 | 0.005295195 | 0.029677811 |
| PLXNB3 | 129.5643 | 1.2557 | 0.3309 | 3.7945 | 0.000147958 | 0.001722991 |
| EVI5 | 3826.8217 | 1.2515 | 0.1544 | 8.1071 | 5.18E-16 | 1.09E-13 |
| OSBPL6 | 88.0478 | 1.2495 | 0.3642 | 3.4311 | 0.000601216 | 0.00530297 |
| AKAP12 | 129.1634 | 1.2453 | 0.2375 | 5.2428 | 1.58E-07 | 5.51E-06 |
| FLJ44511 | 25.5374 | 1.2434 | 0.4612 | 2.6958 | 0.007021786 | 0.036601846 |
| PIM3 | 7375.8084 | 1.2434 | 0.1175 | 10.5849 | 3.50E-26 | 4.08E-23 |
| RGS6 | 271.6821 | 1.2379 | 0.2581 | 4.7957 | 1.62E-06 | 3.96E-05 |
| KIRREL2 | 66.2039 | 1.2368 | 0.3536 | 3.4973 | 0.000469919 | 0.004377131 |
| NINJ1 | 4261.8222 | 1.2337 | 0.1218 | 10.1273 | 4.18E-24 | 2.87E-21 |
| PTPRN | 301.8764 | 1.2293 | 0.2434 | 5.0512 | 4.39E-07 | 1.31E-05 |
| LAG3 | 20716.4242 | 1.2275 | 0.1382 | 8.8851 | 6.39E-19 | 1.97E-16 |
| SDC4 | 3114.7033 | 1.2263 | 0.1639 | 7.4796 | 7.45E-14 | 1.03E-11 |
| MYO1B | 1038.6368 | 1.2257 | 0.1638 | 7.4840 | 7.21E-14 | 1.01E-11 |
| SPTBN4 | 124.2888 | 1.2244 | 0.3419 | 3.5809 | 0.000342374 | 0.003411788 |
| BATF3 | 1582.9889 | 1.2175 | 0.1347 | 9.0413 | 1.55E-19 | 5.44E-17 |
| DPYSL4 | 300.5929 | 1.2168 | 0.2614 | 4.6551 | 3.24E-06 | 7.15E-05 |
| METRNL | 1363.2655 | 1.2145 | 0.1723 | 7.0471 | 1.83E-12 | 1.88E-10 |
| PER2 | 604.7714 | 1.2010 | 0.1694 | 7.0887 | 1.35E-12 | 1.45E-10 |
| RGS1 | 21234.1685 | 1.2005 | 0.2263 | 5.3042 | 1.13E-07 | 4.09E-06 |
| SPRED2 | 109.1070 | 1.1985 | 0.2811 | 4.2638 | 2.01E-05 | 0.00033468 |
| HIF1A-AS2 | 86.5260 | 1.1963 | 0.3474 | 3.4438 | 0.000573597 | 0.005122081 |
| PLEKHA7 | 1359.5237 | 1.1922 | 0.2121 | 5.6217 | 1.89E-08 | 8.69E-07 |
| IL2 | 119.1134 | 1.1907 | 0.3891 | 3.0597 | 0.002215798 | 0.014953948 |
| SAMD14 | 122.5880 | 1.1840 | 0.2615 | 4.5269 | 5.98E-06 | 0.000122 |
| EBI3 | 970.3506 | 1.1839 | 0.1873 | 6.3196 | 2.62E-10 | 1.76E-08 |
| RELB | 2466.5156 | 1.1834 | 0.1392 | 8.5038 | 1.84E-17 | 4.79E-15 |
| PTPN3 | 3115.5898 | 1.1827 | 0.2387 | 4.9542 | 7.26E-07 | 2.01E-05 |
| VPS37D | 76.9071 | 1.1806 | 0.3317 | 3.5596 | 0.000371441 | 0.003613367 |
| TMTC1 | 99.2548 | 1.1681 | 0.2651 | 4.4062 | 1.05E-05 | 0.0001955 |
| MEST | 61.5565 | 1.1668 | 0.3868 | 3.0164 | 0.002557891 | 0.016670926 |
| SEMA3B | 94.5257 | 1.1660 | 0.3743 | 3.1150 | 0.001839352 | 0.012929312 |
| ARID5A | 5644.6228 | 1.1633 | 0.1422 | 8.1788 | 2.87E-16 | 6.19E-14 |
| TNFRSF4 | 8102.6381 | 1.1620 | 0.1358 | 8.5562 | 1.17E-17 | 3.18E-15 |
| LMCD1 | 750.3477 | 1.1589 | 0.3257 | 3.5580 | 0.000373637 | 0.00363006 |
| BAI2 | 117.0559 | 1.1565 | 0.2928 | 3.9500 | 7.82E-05 | 0.001034284 |
| TCF4 | 207.0831 | 1.1520 | 0.2484 | 4.6378 | 3.52E-06 | 7.70E-05 |
| IL26 | 1397.9339 | 1.1494 | 0.1850 | 6.2123 | 5.22E-10 | 3.26E-08 |
| CCDC144C | 45.4648 | 1.1477 | 0.3266 | 3.5144 | 0.00044077 | 0.004162 |
| CDC42EP1 | 108.9937 | 1.1421 | 0.3500 | 3.2628 | 0.001103176 | 0.008610809 |
| PLS3 | 561.3750 | 1.1408 | 0.2218 | 5.1441 | 2.69E-07 | 8.65E-06 |
| RCAN1 | 591.7299 | 1.1400 | 0.1780 | 6.4028 | 1.53E-10 | 1.09E-08 |
| ADORA2B | 173.7243 | 1.1391 | 0.3771 | 3.0208 | 0.002520688 | 0.016506688 |
| STAT3 | 19765.6127 | 1.1363 | 0.2181 | 5.2091 | 1.90E-07 | 6.34E-06 |
| DMD | 781.8435 | 1.1352 | 0.1591 | 7.1349 | 9.69E-13 | 1.08E-10 |
| SNHG15 | 6612.5394 | 1.1285 | 0.1319 | 8.5551 | 1.18E-17 | 3.18E-15 |
| VDR | 10617.1679 | 1.1280 | 0.1285 | 8.7759 | 1.69E-18 | 4.93E-16 |
| CPM | 402.5749 | 1.1144 | 0.2309 | 4.8261 | 1.39E-06 | 3.48E-05 |
| NFKBID | 1112.3851 | 1.1105 | 0.1733 | 6.4095 | 1.46E-10 | 1.05E-08 |
| GJB2 | 466.7067 | 1.1104 | 0.2858 | 3.8845 | 0.000102522 | 0.001282758 |
| SLC7A5 | 10164.2759 | 1.1093 | 0.2186 | 5.0744 | 3.89E-07 | 1.17E-05 |
| RCAN2 | 580.8067 | 1.0977 | 0.2250 | 4.8780 | 1.07E-06 | 2.79E-05 |
| LIF | 3558.1507 | 1.0973 | 0.2908 | 3.7735 | 0.000161005 | 0.001846496 |
| ZC3H12C | 6573.9308 | 1.0923 | 0.2426 | 4.5031 | 6.70E-06 | 0.000134516 |
| AXL | 149.7810 | 1.0826 | 0.3903 | 2.7742 | 0.00553345 | 0.030605504 |
| ATF3 | 2391.6819 | 1.0824 | 0.1658 | 6.5273 | 6.70E-11 | 5.07E-09 |
| CSF1 | 4539.6048 | 1.0805 | 0.2462 | 4.3886 | 1.14E-05 | 0.000207962 |
| PPP1R3B | 4530.0808 | 1.0752 | 0.1276 | 8.4242 | 3.63E-17 | 8.86E-15 |
| ID3 | 97.6249 | 1.0750 | 0.3786 | 2.8396 | 0.004517607 | 0.02621321 |
| NDFIP2 | 8005.6901 | 1.0747 | 0.1409 | 7.6297 | 2.35E-14 | 3.63E-12 |
| LIN7A | 342.6455 | 1.0734 | 0.2676 | 4.0114 | 6.04E-05 | 0.000836165 |
| FZD7 | 93.7025 | 1.0714 | 0.3007 | 3.5625 | 0.000367336 | 0.003587272 |
| MB21D2 | 1277.5099 | 1.0691 | 0.0958 | 11.1604 | 6.37E-29 | 1.61E-25 |
| MTSS1 | 1144.0719 | 1.0682 | 0.2109 | 5.0636 | 4.11E-07 | 1.23E-05 |
| CXADR | 148.3829 | 1.0680 | 0.2521 | 4.2365 | 2.27E-05 | 0.000367271 |
| ELL2 | 15173.0395 | 1.0664 | 0.0895 | 11.9185 | 9.48E-33 | 1.02E-28 |
| NEU4 | 74.1154 | 1.0653 | 0.3303 | 3.2250 | 0.001259693 | 0.009590023 |
| MAL | 4187.4598 | 1.0641 | 0.1293 | 8.2308 | 1.86E-16 | 4.14E-14 |
| SLC8A3 | 137.8620 | 1.0610 | 0.3670 | 2.8908 | 0.003842507 | 0.023051837 |
| ITPKA | 231.1727 | 1.0597 | 0.2708 | 3.9136 | 9.09E-05 | 0.001163651 |
| SGPP2 | 689.1459 | 1.0575 | 0.2218 | 4.7668 | 1.87E-06 | 4.47E-05 |
| DOT1L | 8531.8449 | 1.0565 | 0.1183 | 8.9283 | 4.32E-19 | 1.36E-16 |
| SEMA7A | 1324.8658 | 1.0535 | 0.2020 | 5.2158 | 1.83E-07 | 6.12E-06 |
| EXOC3L1 | 57.6042 | 1.0524 | 0.3271 | 3.2175 | 0.001293197 | 0.009795791 |
| ZNF282 | 3865.6365 | 1.0485 | 0.1598 | 6.5624 | 5.30E-11 | 4.07E-09 |
| CAMK1 | 401.3468 | 1.0477 | 0.2039 | 5.1378 | 2.78E-07 | 8.87E-06 |
| CDKN1A | 6686.3650 | 1.0447 | 0.1664 | 6.2801 | 3.38E-10 | 2.22E-08 |
| NFKBIZ | 2425.7427 | 1.0426 | 0.1456 | 7.1624 | 7.93E-13 | 9.02E-11 |
| PLAT | 623.5099 | 1.0421 | 0.1789 | 5.8252 | 5.71E-09 | 2.90E-07 |
| IGFBP2 | 340.3177 | 1.0366 | 0.2520 | 4.1130 | 3.91E-05 | 0.000581557 |
| TMOD1 | 475.3079 | 1.0354 | 0.2835 | 3.6519 | 0.000260272 | 0.002717139 |
| PPME1 | 3137.2032 | 1.0329 | 0.1735 | 5.9546 | 2.61E-09 | 1.41E-07 |
| CDHR1 | 93.8359 | 1.0291 | 0.3855 | 2.6697 | 0.007592926 | 0.038829681 |
| PTGIR | 593.0309 | 1.0284 | 0.2017 | 5.0990 | 3.41E-07 | 1.05E-05 |
| CSF2RB | 1040.5058 | 1.0257 | 0.2065 | 4.9671 | 6.80E-07 | 1.92E-05 |
| HAVCR2 | 6339.9892 | 1.0256 | 0.1289 | 7.9582 | 1.75E-15 | 3.23E-13 |
| WBP5 | 83.1671 | 1.0254 | 0.3946 | 2.5988 | 0.009355252 | 0.045474579 |
| FES | 815.8929 | 1.0229 | 0.1714 | 5.9680 | 2.40E-09 | 1.32E-07 |
| SOX5 | 126.5612 | 1.0222 | 0.3426 | 2.9836 | 0.002849253 | 0.018286233 |
| HMSD | 715.9840 | 1.0189 | 0.1711 | 5.9552 | 2.60E-09 | 1.41E-07 |
| PMAIP1 | 6376.8357 | 1.0174 | 0.1576 | 6.4542 | 1.09E-10 | 7.95E-09 |
| FURIN | 31774.6249 | 1.0161 | 0.1010 | 10.0586 | 8.42E-24 | 5.53E-21 |
| LINC00239 | 40.2191 | 1.0123 | 0.3650 | 2.7730 | 0.005553813 | 0.030636243 |
| CSRNP1 | 4480.1323 | 1.0097 | 0.1210 | 8.3443 | 7.16E-17 | 1.67E-14 |
| MTHFD2 | 7261.6723 | 1.0091 | 0.1542 | 6.5458 | 5.92E-11 | 4.52E-09 |
| GAL3ST2 | 135.9889 | 1.0046 | 0.2826 | 3.5550 | 0.000377955 | 0.003660256 |
| CHRNA6 | 241.1207 | 1.0001 | 0.3374 | 2.9637 | 0.003039887 | 0.019200158 |
| MIR155HG | 2767.1208 | 0.9974 | 0.1268 | 7.8665 | 3.65E-15 | 6.41E-13 |
| ART3 | 161.1052 | 0.9958 | 0.3228 | 3.0843 | 0.002040073 | 0.014052905 |
| PIM1 | 16513.2598 | 0.9945 | 0.1020 | 9.7478 | 1.88E-22 | 8.91E-20 |
| VWA1 | 146.3728 | 0.9941 | 0.2699 | 3.6829 | 0.000230587 | 0.002468568 |
| NPTX1 | 763.7366 | 0.9923 | 0.1874 | 5.2941 | 1.20E-07 | 4.27E-06 |
| SERPINB9 | 245.7308 | 0.9883 | 0.2353 | 4.1996 | 2.67E-05 | 0.000420381 |
| CISH | 5287.2228 | 0.9795 | 0.1453 | 6.7438 | 1.54E-11 | 1.37E-09 |
| NFKB2 | 12232.3361 | 0.9792 | 0.1005 | 9.7421 | 1.99E-22 | 9.14E-20 |
| IRF4 | 13208.2818 | 0.9767 | 0.1253 | 7.7921 | 6.59E-15 | 1.10E-12 |
| NUPL1 | 13839.6095 | 0.9763 | 0.1085 | 8.9942 | 2.38E-19 | 8.00E-17 |
| SALL4 | 244.4334 | 0.9751 | 0.3026 | 3.2226 | 0.001270337 | 0.009651626 |
| ADAM19 | 8720.8618 | 0.9629 | 0.1839 | 5.2372 | 1.63E-07 | 5.64E-06 |
| TDRD9 | 59.5974 | 0.9610 | 0.3723 | 2.5810 | 0.009852694 | 0.047299812 |
| HAPLN3 | 2152.7148 | 0.9565 | 0.0993 | 9.6281 | 6.08E-22 | 2.71E-19 |
| AGRN | 6748.9744 | 0.9537 | 0.2160 | 4.4143 | 1.01E-05 | 0.000188798 |
| SBNO2 | 7939.9419 | 0.9525 | 0.1236 | 7.7053 | 1.31E-14 | 2.10E-12 |
| RBKS | 277.5270 | 0.9523 | 0.1951 | 4.8818 | 1.05E-06 | 2.74E-05 |
| OSBPL10 | 958.7070 | 0.9519 | 0.1783 | 5.3399 | 9.30E-08 | 3.43E-06 |
| NSUN7 | 531.2895 | 0.9518 | 0.2446 | 3.8918 | 9.95E-05 | 0.001251244 |
| PER1 | 10550.7855 | 0.9473 | 0.1484 | 6.3828 | 1.74E-10 | 1.22E-08 |
| ETS2 | 1449.9694 | 0.9447 | 0.1331 | 7.0951 | 1.29E-12 | 1.40E-10 |
| DAPK1 | 667.4895 | 0.9324 | 0.1904 | 4.8979 | 9.69E-07 | 2.57E-05 |
| NFKBIA | 4850.1519 | 0.9283 | 0.1212 | 7.6591 | 1.87E-14 | 2.95E-12 |
| C17orf96 | 868.9163 | 0.9256 | 0.1778 | 5.2049 | 1.94E-07 | 6.47E-06 |
| PDCD1 | 2289.6004 | 0.9231 | 0.1450 | 6.3684 | 1.91E-10 | 1.32E-08 |
| HLF | 234.8006 | 0.9223 | 0.2931 | 3.1470 | 0.001649624 | 0.011877132 |
| FASLG | 832.6597 | 0.9151 | 0.2197 | 4.1657 | 3.10E-05 | 0.00047772 |
| ANKRD33B | 3433.8006 | 0.9130 | 0.1343 | 6.7982 | 1.06E-11 | 9.54E-10 |
| HEG1 | 1184.0051 | 0.9126 | 0.1652 | 5.5250 | 3.29E-08 | 1.40E-06 |
| HOXB9 | 365.7986 | 0.9125 | 0.2350 | 3.8832 | 0.000103071 | 0.001287491 |
| PPARG | 997.2388 | 0.9117 | 0.1300 | 7.0133 | 2.33E-12 | 2.35E-10 |
| PLAUR | 830.4361 | 0.9084 | 0.1137 | 7.9918 | 1.33E-15 | 2.55E-13 |
| VSIG10L | 430.2091 | 0.9080 | 0.2093 | 4.3381 | 1.44E-05 | 0.000253077 |
| SERPINE1 | 963.2922 | 0.9045 | 0.1862 | 4.8578 | 1.19E-06 | 3.04E-05 |
| BAIAP2 | 158.7668 | 0.9039 | 0.2056 | 4.3973 | 1.10E-05 | 0.000202166 |
| ZNRF1 | 5014.5426 | 0.9036 | 0.0927 | 9.7518 | 1.81E-22 | 8.84E-20 |
| LOC646329 | 410.4463 | 0.9030 | 0.1930 | 4.6779 | 2.90E-06 | 6.53E-05 |
| SHB | 190.2703 | 0.9028 | 0.2445 | 3.6925 | 0.00022202 | 0.002404082 |
| PHEX | 467.6188 | 0.8992 | 0.1719 | 5.2304 | 1.69E-07 | 5.74E-06 |
| F5 | 2824.7887 | 0.8984 | 0.1884 | 4.7678 | 1.86E-06 | 4.46E-05 |
| SGK1 | 317.1372 | 0.8962 | 0.2617 | 3.4253 | 0.000614143 | 0.005391841 |
| EPAS1 | 2105.5201 | 0.8962 | 0.1909 | 4.6957 | 2.66E-06 | 6.07E-05 |
| TERT | 112.5767 | 0.8959 | 0.3230 | 2.7741 | 0.005535576 | 0.030605504 |
| ATP8B4 | 10731.5650 | 0.8925 | 0.1299 | 6.8707 | 6.39E-12 | 5.93E-10 |
| PNCK | 105.4026 | 0.8904 | 0.3169 | 2.8092 | 0.004966335 | 0.028282343 |
| C6orf105 | 350.1086 | 0.8896 | 0.2695 | 3.3007 | 0.00096452 | 0.007764928 |
| GAS2L3 | 154.7860 | 0.8891 | 0.3008 | 2.9558 | 0.003118439 | 0.019581828 |
| GADD45G | 730.5048 | 0.8887 | 0.1963 | 4.5279 | 5.96E-06 | 0.000121793 |
| RFX2 | 387.0909 | 0.8878 | 0.1520 | 5.8423 | 5.15E-09 | 2.63E-07 |
| SLC43A3 | 1798.8889 | 0.8853 | 0.2762 | 3.2052 | 0.001349807 | 0.010123218 |
| KCNMA1 | 76.8377 | 0.8853 | 0.3254 | 2.7203 | 0.006521494 | 0.034723912 |
| JUN | 6491.4696 | 0.8789 | 0.1409 | 6.2363 | 4.48E-10 | 2.86E-08 |
| HDC | 95.6885 | 0.8782 | 0.2720 | 3.2282 | 0.001245785 | 0.009523343 |
| SNX9 | 4885.0232 | 0.8740 | 0.1329 | 6.5748 | 4.87E-11 | 3.82E-09 |
| MYO1E | 2964.6831 | 0.8717 | 0.1429 | 6.0986 | 1.07E-09 | 6.40E-08 |
| CABLES1 | 379.6180 | 0.8684 | 0.1504 | 5.7759 | 7.65E-09 | 3.78E-07 |
| GPR153 | 64.7660 | 0.8672 | 0.3102 | 2.7957 | 0.005179489 | 0.029202436 |
| GNA15 | 14790.6736 | 0.8614 | 0.1382 | 6.2318 | 4.61E-10 | 2.93E-08 |
| OAS3 | 5849.1218 | 0.8613 | 0.2554 | 3.3726 | 0.000744532 | 0.006320163 |
| EBF4 | 322.1154 | 0.8604 | 0.2137 | 4.0262 | 5.67E-05 | 0.00079248 |
| RHOB | 769.6414 | 0.8587 | 0.2450 | 3.5051 | 0.000456364 | 0.004285177 |
| CCR7 | 3415.7045 | 0.8563 | 0.1863 | 4.5956 | 4.32E-06 | 9.26E-05 |
| NTRK2 | 1430.8549 | 0.8535 | 0.1485 | 5.7484 | 9.01E-09 | 4.37E-07 |
| TRIB1 | 480.7597 | 0.8521 | 0.2692 | 3.1651 | 0.001550387 | 0.011286194 |
| GNG4 | 1043.0426 | 0.8518 | 0.2196 | 3.8790 | 0.000104871 | 0.001305662 |
| STK3 | 276.4340 | 0.8514 | 0.2276 | 3.7406 | 0.000183601 | 0.002071093 |
| SERPINE2 | 2006.1833 | 0.8494 | 0.1204 | 7.0542 | 1.74E-12 | 1.84E-10 |
| HIVEP1 | 3796.9022 | 0.8478 | 0.1817 | 4.6668 | 3.06E-06 | 6.84E-05 |
| HOMER2 | 582.2082 | 0.8408 | 0.1983 | 4.2395 | 2.24E-05 | 0.000363661 |
| UPRT | 2174.5980 | 0.8407 | 0.0994 | 8.4607 | 2.66E-17 | 6.81E-15 |
| SH2B2 | 205.2902 | 0.8402 | 0.2738 | 3.0690 | 0.00214803 | 0.014649795 |
| SEC24A | 12846.0907 | 0.8396 | 0.1409 | 5.9610 | 2.51E-09 | 1.36E-07 |
| FAM114A1 | 78.0844 | 0.8306 | 0.3090 | 2.6886 | 0.007175911 | 0.037178191 |
| RIMKLA | 1090.0406 | 0.8273 | 0.1544 | 5.3579 | 8.42E-08 | 3.17E-06 |
| MAFF | 1351.3227 | 0.8270 | 0.1360 | 6.0790 | 1.21E-09 | 7.12E-08 |
| HERC3 | 6316.5542 | 0.8268 | 0.1519 | 5.4432 | 5.23E-08 | 2.09E-06 |
| ADCY1 | 1099.1613 | 0.8267 | 0.2980 | 2.7739 | 0.005538174 | 0.03060868 |
| LOC100506274 | 492.1382 | 0.8243 | 0.2018 | 4.0849 | 4.41E-05 | 0.000643175 |
| GPT2 | 1574.9523 | 0.8234 | 0.1452 | 5.6693 | 1.43E-08 | 6.67E-07 |
| NIM1 | 292.9924 | 0.8227 | 0.2063 | 3.9881 | 6.66E-05 | 0.00090438 |
| LRRC8B | 4305.6262 | 0.8226 | 0.1172 | 7.0162 | 2.28E-12 | 2.31E-10 |
| RAMP1 | 342.5158 | 0.8206 | 0.2950 | 2.7820 | 0.005403111 | 0.030082023 |
| BCL2 | 11033.7871 | 0.8205 | 0.0818 | 10.0318 | 1.10E-23 | 6.96E-21 |
| ETV7 | 674.0253 | 0.8202 | 0.1658 | 4.9458 | 7.59E-07 | 2.09E-05 |
| PYCR1 | 1082.9602 | 0.8196 | 0.2105 | 3.8944 | 9.84E-05 | 0.001240987 |
| GADD45A | 5361.1751 | 0.8186 | 0.1442 | 5.6760 | 1.38E-08 | 6.44E-07 |
| ICAM1 | 3221.9920 | 0.8174 | 0.1237 | 6.6090 | 3.87E-11 | 3.13E-09 |
| AFAP1 | 695.5326 | 0.8171 | 0.1184 | 6.9000 | 5.20E-12 | 4.92E-10 |
| B4GALNT1 | 131.5792 | 0.8167 | 0.2452 | 3.3314 | 0.000864197 | 0.007108592 |
| CD83 | 578.9482 | 0.8162 | 0.1587 | 5.1430 | 2.70E-07 | 8.69E-06 |
| INSIG2 | 15858.0527 | 0.8158 | 0.1360 | 6.0004 | 1.97E-09 | 1.11E-07 |
| PBX4 | 3737.5888 | 0.8117 | 0.1035 | 7.8460 | 4.30E-15 | 7.47E-13 |
| C1orf198 | 319.4574 | 0.8110 | 0.2084 | 3.8913 | 9.97E-05 | 0.001252905 |
| VMP1 | 7786.7465 | 0.8104 | 0.0971 | 8.3506 | 6.79E-17 | 1.63E-14 |
| RAB38 | 413.2808 | 0.8066 | 0.2431 | 3.3183 | 0.000905726 | 0.007396464 |
| PTPRF | 761.3232 | 0.8065 | 0.1990 | 4.0532 | 5.05E-05 | 0.000718869 |
| SPOCK1 | 269.6784 | 0.8053 | 0.1972 | 4.0829 | 4.45E-05 | 0.000648064 |
| SLC29A1 | 1094.1828 | 0.8047 | 0.1518 | 5.2998 | 1.16E-07 | 4.18E-06 |
| HLA-L | 589.5144 | 0.8033 | 0.2647 | 3.0352 | 0.002404077 | 0.015983653 |
| RHPN2 | 108.1407 | 0.8025 | 0.2657 | 3.0203 | 0.002524943 | 0.016520251 |
| PHLDA1 | 15182.3566 | 0.8009 | 0.1469 | 5.4524 | 4.97E-08 | 1.99E-06 |
| FNDC3B | 1609.2191 | 0.7979 | 0.0972 | 8.2095 | 2.22E-16 | 4.87E-14 |
| PIM2 | 18119.1479 | 0.7978 | 0.1008 | 7.9169 | 2.43E-15 | 4.38E-13 |
| PRDM1 | 14746.3005 | 0.7967 | 0.1377 | 5.7863 | 7.19E-09 | 3.59E-07 |
| SETBP1 | 1305.7497 | 0.7964 | 0.1775 | 4.4859 | 7.26E-06 | 0.000142807 |
| CDK6 | 15205.7567 | 0.7964 | 0.1129 | 7.0527 | 1.75E-12 | 1.84E-10 |
| WNT10A | 115.0032 | 0.7883 | 0.2930 | 2.6907 | 0.007129476 | 0.036988897 |
| TNIP2 | 4947.9782 | 0.7863 | 0.0987 | 7.9651 | 1.65E-15 | 3.12E-13 |
| KIAA0895L | 2531.4214 | 0.7855 | 0.1124 | 6.9865 | 2.82E-12 | 2.82E-10 |
| CTH | 1473.6789 | 0.7824 | 0.1855 | 4.2177 | 2.47E-05 | 0.00039428 |
| USP37 | 5301.5819 | 0.7820 | 0.1171 | 6.6784 | 2.42E-11 | 2.04E-09 |
| GBE1 | 9936.1723 | 0.7813 | 0.1100 | 7.1000 | 1.25E-12 | 1.36E-10 |
| REL | 1634.1897 | 0.7806 | 0.2008 | 3.8885 | 0.000100886 | 0.001264377 |
| ARL5B | 2623.5774 | 0.7805 | 0.1834 | 4.2551 | 2.09E-05 | 0.000345421 |
| PANX2 | 456.2423 | 0.7770 | 0.1518 | 5.1184 | 3.08E-07 | 9.65E-06 |
| BHLHE40 | 49440.1701 | 0.7768 | 0.0937 | 8.2899 | 1.13E-16 | 2.56E-14 |
| C15orf42 | 1928.1128 | 0.7763 | 0.1243 | 6.2466 | 4.19E-10 | 2.69E-08 |
| CYTIP | 32256.2707 | 0.7751 | 0.0722 | 10.7385 | 6.71E-27 | 9.23E-24 |
| KISS1R | 428.1540 | 0.7736 | 0.2575 | 3.0041 | 0.002664007 | 0.017243658 |
| GPR125 | 262.8483 | 0.7732 | 0.2049 | 3.7738 | 0.000160768 | 0.001845173 |
| PTRF | 262.3566 | 0.7730 | 0.2379 | 3.2492 | 0.001157114 | 0.008948705 |
| NXT1 | 813.0751 | 0.7724 | 0.1820 | 4.2431 | 2.20E-05 | 0.000359357 |
| KIAA1324 | 2330.6782 | 0.7719 | 0.1266 | 6.0964 | 1.08E-09 | 6.46E-08 |
| GPR56 | 9688.6418 | 0.7698 | 0.1712 | 4.4972 | 6.89E-06 | 0.000136682 |
| KIF7 | 165.2014 | 0.7668 | 0.2610 | 2.9378 | 0.003305604 | 0.020459849 |
| RAB3A | 295.6370 | 0.7624 | 0.1694 | 4.5008 | 6.77E-06 | 0.000134997 |
| AK2 | 1168.0146 | 0.7623 | 0.1380 | 5.5226 | 3.34E-08 | 1.42E-06 |
| KIAA1467 | 995.9807 | 0.7609 | 0.1425 | 5.3393 | 9.33E-08 | 3.43E-06 |
| IL17RB | 3426.5335 | 0.7601 | 0.2717 | 2.7978 | 0.005144448 | 0.029080743 |
| TMPRSS3 | 8057.2309 | 0.7601 | 0.2555 | 2.9751 | 0.002928615 | 0.018684585 |
| IQGAP3 | 1600.0309 | 0.7595 | 0.1292 | 5.8795 | 4.11E-09 | 2.13E-07 |
| IQCG | 3665.5785 | 0.7569 | 0.1094 | 6.9218 | 4.46E-12 | 4.35E-10 |
| SREBF1 | 3670.8139 | 0.7567 | 0.1419 | 5.3330 | 9.66E-08 | 3.54E-06 |
| IRF7 | 2294.5310 | 0.7559 | 0.1824 | 4.1453 | 3.39E-05 | 0.000516455 |
| EHD4 | 1991.9937 | 0.7463 | 0.1642 | 4.5459 | 5.47E-06 | 0.000114274 |
| PDGFRB | 462.0476 | 0.7451 | 0.2254 | 3.3052 | 0.000948998 | 0.007665477 |
| APBA1 | 767.9176 | 0.7437 | 0.2350 | 3.1652 | 0.001549655 | 0.011286194 |
| SLC27A2 | 4797.2386 | 0.7428 | 0.1165 | 6.3753 | 1.83E-10 | 1.27E-08 |
| DST | 2676.9442 | 0.7393 | 0.1743 | 4.2414 | 2.22E-05 | 0.000361266 |
| GAN | 480.9846 | 0.7393 | 0.1402 | 5.2748 | 1.33E-07 | 4.69E-06 |
| P4HA2 | 7004.7767 | 0.7389 | 0.0990 | 7.4646 | 8.36E-14 | 1.15E-11 |
| PIK3AP1 | 859.1709 | 0.7347 | 0.1346 | 5.4591 | 4.78E-08 | 1.93E-06 |
| SLC6A8 | 531.7286 | 0.7310 | 0.1004 | 7.2827 | 3.27E-13 | 3.99E-11 |
| LAPTM4B | 418.3441 | 0.7309 | 0.2738 | 2.6698 | 0.007590395 | 0.038829681 |
| JAM3 | 946.7145 | 0.7293 | 0.1946 | 3.7479 | 0.000178347 | 0.00201633 |
| DUSP2 | 1795.4692 | 0.7285 | 0.1471 | 4.9512 | 7.37E-07 | 2.04E-05 |
| COL1A1 | 177.2759 | 0.7283 | 0.2393 | 3.0441 | 0.002333654 | 0.015626908 |
| 10-Sep | 67.3511 | 0.7282 | 0.2776 | 2.6231 | 0.008713195 | 0.043053312 |
| MCL1 | 46812.6720 | 0.7281 | 0.0614 | 11.8555 | 2.02E-32 | 1.02E-28 |
| PIP5K1B | 1091.2644 | 0.7246 | 0.1408 | 5.1477 | 2.64E-07 | 8.54E-06 |
| MAPT | 92.3479 | 0.7228 | 0.2601 | 2.7793 | 0.005447817 | 0.030252984 |
| CREM | 2859.3150 | 0.7211 | 0.1249 | 5.7716 | 7.85E-09 | 3.86E-07 |
| TRAF4 | 2155.6408 | 0.7198 | 0.0996 | 7.2239 | 5.05E-13 | 6.02E-11 |
| GK | 7533.3741 | 0.7192 | 0.1317 | 5.4619 | 4.71E-08 | 1.91E-06 |
| ENPP2 | 819.6837 | 0.7185 | 0.1655 | 4.3420 | 1.41E-05 | 0.00025037 |
| TUBB6 | 609.0021 | 0.7182 | 0.1651 | 4.3507 | 1.36E-05 | 0.000241514 |
| EPB41L2 | 1951.8163 | 0.7168 | 0.1142 | 6.2754 | 3.49E-10 | 2.26E-08 |
| CHSY1 | 7826.8739 | 0.7121 | 0.1245 | 5.7208 | 1.06E-08 | 5.08E-07 |
| GRAMD1B | 5427.0088 | 0.7100 | 0.1538 | 4.6173 | 3.89E-06 | 8.43E-05 |
| ZBTB17 | 3418.8331 | 0.7079 | 0.1070 | 6.6137 | 3.75E-11 | 3.07E-09 |
| CBLN3 | 849.9206 | 0.7057 | 0.1754 | 4.0229 | 5.75E-05 | 0.000803062 |
| DIXDC1 | 3245.9265 | 0.7055 | 0.1445 | 4.8842 | 1.04E-06 | 2.72E-05 |
| PPTC7 | 6206.6340 | 0.7034 | 0.2323 | 3.0281 | 0.002461174 | 0.016236448 |
| ZDHHC9 | 1582.0444 | 0.7029 | 0.1078 | 6.5190 | 7.08E-11 | 5.30E-09 |
| SFXN2 | 584.5624 | 0.7023 | 0.1732 | 4.0541 | 5.03E-05 | 0.000716981 |
| SPHK1 | 188.1996 | 0.7023 | 0.2417 | 2.9053 | 0.003668968 | 0.022217968 |
| FOSL1 | 202.0934 | 0.7017 | 0.2230 | 3.1465 | 0.001652274 | 0.01188402 |
| MAPK11 | 1494.2214 | 0.6996 | 0.1309 | 5.3431 | 9.14E-08 | 3.40E-06 |
| MAOA | 364.0576 | 0.6995 | 0.1774 | 3.9426 | 8.06E-05 | 0.001059567 |
| IL4R | 14188.1857 | 0.6993 | 0.1097 | 6.3749 | 1.83E-10 | 1.27E-08 |
| RNF19A | 23348.4372 | 0.6992 | 0.2077 | 3.3662 | 0.00076207 | 0.006436535 |
| CTPS | 988.0741 | 0.6987 | 0.1229 | 5.6867 | 1.30E-08 | 6.09E-07 |
| ICAM5 | 523.1947 | 0.6979 | 0.2181 | 3.2004 | 0.001372292 | 0.010236378 |
| SLC22A4 | 213.2950 | 0.6956 | 0.2187 | 3.1808 | 0.00146848 | 0.010825387 |
| CCDC64 | 6043.9983 | 0.6954 | 0.1121 | 6.2020 | 5.57E-10 | 3.46E-08 |
| BZW1 | 16307.7202 | 0.6917 | 0.0671 | 10.3033 | 6.81E-25 | 6.06E-22 |
| FABP5 | 1483.0516 | 0.6905 | 0.1536 | 4.4947 | 6.97E-06 | 0.000137949 |
| FAM46A | 336.6082 | 0.6898 | 0.1897 | 3.6358 | 0.000277154 | 0.002870169 |
| FAAH2 | 1185.4743 | 0.6895 | 0.1521 | 4.5323 | 5.84E-06 | 0.000120392 |
| TMEM74B | 276.0696 | 0.6879 | 0.2040 | 3.3721 | 0.000746015 | 0.006326129 |
| CMAHP | 15171.5028 | 0.6872 | 0.1246 | 5.5145 | 3.50E-08 | 1.47E-06 |
| STX6 | 2389.6274 | 0.6870 | 0.1332 | 5.1568 | 2.51E-07 | 8.16E-06 |
| WDR45L | 11968.3339 | 0.6864 | 0.0911 | 7.5303 | 5.06E-14 | 7.36E-12 |
| KLF10 | 4545.8321 | 0.6838 | 0.0846 | 8.0845 | 6.24E-16 | 1.29E-13 |
| TRIM9 | 76.8256 | 0.6834 | 0.2531 | 2.6997 | 0.006940888 | 0.036283412 |
| LINC00263 | 1336.6134 | 0.6824 | 0.1803 | 3.7859 | 0.000153161 | 0.001771309 |
| MXD1 | 3994.8891 | 0.6823 | 0.1099 | 6.2104 | 5.29E-10 | 3.29E-08 |
| GOLIM4 | 1163.7009 | 0.6815 | 0.2164 | 3.1500 | 0.001632649 | 0.011771728 |
| IGSF3 | 186.1110 | 0.6815 | 0.2298 | 2.9655 | 0.003022159 | 0.019125334 |
| GINS4 | 1167.8104 | 0.6800 | 0.1401 | 4.8531 | 1.22E-06 | 3.10E-05 |
| STX1A | 325.2553 | 0.6764 | 0.2404 | 2.8139 | 0.004893729 | 0.027987692 |
| ZC3H12D | 10304.9420 | 0.6763 | 0.0979 | 6.9051 | 5.02E-12 | 4.77E-10 |
| RQCD1 | 3678.8918 | 0.6751 | 0.1599 | 4.2213 | 2.43E-05 | 0.000389716 |
| DEPDC7 | 900.9480 | 0.6742 | 0.1578 | 4.2727 | 1.93E-05 | 0.000324648 |
| ETV3 | 1154.8384 | 0.6732 | 0.2029 | 3.3178 | 0.000907162 | 0.007397652 |
| ANKRD37 | 2490.5368 | 0.6731 | 0.1474 | 4.5674 | 4.94E-06 | 0.000104023 |
| TMEM217 | 235.7092 | 0.6710 | 0.2383 | 2.8151 | 0.004876272 | 0.027919519 |
| EPSTI1 | 1958.5319 | 0.6652 | 0.1209 | 5.5017 | 3.76E-08 | 1.57E-06 |
| WARS | 13515.6785 | 0.6639 | 0.1332 | 4.9839 | 6.23E-07 | 1.78E-05 |
| ANO7 | 166.5927 | 0.6622 | 0.2569 | 2.5779 | 0.009939809 | 0.047642424 |
| TNFSF4 | 369.0149 | 0.6614 | 0.1825 | 3.6250 | 0.000288965 | 0.002967526 |
| VWCE | 334.4406 | 0.6588 | 0.2263 | 2.9112 | 0.003600009 | 0.021905604 |
| ZNF267 | 4344.6145 | 0.6580 | 0.0951 | 6.9182 | 4.57E-12 | 4.38E-10 |
| BLM | 4311.5941 | 0.6574 | 0.1080 | 6.0858 | 1.16E-09 | 6.85E-08 |
| PEX3 | 4200.4926 | 0.6561 | 0.1469 | 4.4662 | 7.96E-06 | 0.000154186 |
| CASZ1 | 364.3638 | 0.6546 | 0.1763 | 3.7139 | 0.000204082 | 0.002248465 |
| HDAC7 | 18815.7539 | 0.6523 | 0.0813 | 8.0192 | 1.06E-15 | 2.15E-13 |
| HSD3B7 | 353.8404 | 0.6517 | 0.2333 | 2.7930 | 0.005222413 | 0.029353579 |
| RYBP | 11213.2484 | 0.6510 | 0.1015 | 6.4135 | 1.42E-10 | 1.02E-08 |
| CCNG2 | 9632.3910 | 0.6509 | 0.1045 | 6.2297 | 4.67E-10 | 2.96E-08 |
| PRKCE | 926.1734 | 0.6506 | 0.1574 | 4.1329 | 3.58E-05 | 0.000541249 |
| PPP1CB | 3989.0530 | 0.6465 | 0.2500 | 2.5862 | 0.009704486 | 0.046813247 |
| PRKCDBP | 430.3179 | 0.6465 | 0.2173 | 2.9748 | 0.002931729 | 0.018696017 |
| CTNNA1 | 5909.7670 | 0.6440 | 0.0845 | 7.6170 | 2.60E-14 | 3.97E-12 |
| C3orf26 | 698.6598 | 0.6439 | 0.1756 | 3.6660 | 0.000246404 | 0.002593847 |
| FGF11 | 3976.7722 | 0.6431 | 0.1145 | 5.6171 | 1.94E-08 | 8.87E-07 |
| SERTAD1 | 844.3469 | 0.6425 | 0.1798 | 3.5735 | 0.00035229 | 0.00348307 |
| CLEC4A | 641.3438 | 0.6418 | 0.1367 | 4.6962 | 2.65E-06 | 6.07E-05 |
| UST | 1896.0412 | 0.6414 | 0.1160 | 5.5305 | 3.19E-08 | 1.37E-06 |
| HOMER1 | 302.5186 | 0.6390 | 0.1921 | 3.3269 | 0.00087809 | 0.007199383 |
| IFI44 | 2387.4186 | 0.6386 | 0.1903 | 3.3556 | 0.000791954 | 0.006633382 |
| ESPL1 | 3148.7183 | 0.6381 | 0.1255 | 5.0832 | 3.71E-07 | 1.13E-05 |
| CXorf21 | 931.8702 | 0.6374 | 0.1340 | 4.7561 | 1.97E-06 | 4.67E-05 |
| CCNT1 | 2039.5378 | 0.6371 | 0.1957 | 3.2552 | 0.001133139 | 0.008808325 |
| IKZF3 | 1206.8104 | 0.6355 | 0.1779 | 3.5724 | 0.000353662 | 0.003494344 |
| FANCB | 462.2755 | 0.6333 | 0.1454 | 4.3562 | 1.32E-05 | 0.000236622 |
| SH2D2A | 11577.7522 | 0.6325 | 0.0977 | 6.4715 | 9.70E-11 | 7.12E-09 |
| MCM8 | 2071.8496 | 0.6321 | 0.1366 | 4.6278 | 3.70E-06 | 8.03E-05 |
| RAB20 | 473.3939 | 0.6315 | 0.1794 | 3.5209 | 0.000430145 | 0.00408719 |
| MIR17HG | 158.2817 | 0.6308 | 0.2016 | 3.1294 | 0.001751508 | 0.012444837 |
| GNLY | 104625.0054 | 0.6273 | 0.1116 | 5.6234 | 1.87E-08 | 8.63E-07 |
| DENND3 | 9735.1966 | 0.6269 | 0.1294 | 4.8451 | 1.27E-06 | 3.22E-05 |
| ZC3H12A | 2381.1462 | 0.6252 | 0.1195 | 5.2329 | 1.67E-07 | 5.69E-06 |
| TP53INP2 | 561.6951 | 0.6244 | 0.2223 | 2.8081 | 0.004983562 | 0.028351389 |
| HSD17B12 | 2211.4680 | 0.6241 | 0.1282 | 4.8687 | 1.12E-06 | 2.90E-05 |
| ZMIZ1 | 9651.4720 | 0.6238 | 0.1037 | 6.0138 | 1.81E-09 | 1.03E-07 |
| RBMS3 | 346.5447 | 0.6203 | 0.2216 | 2.7997 | 0.005114798 | 0.028967259 |
| CCDC74B | 146.0024 | 0.6198 | 0.1882 | 3.2932 | 0.000990535 | 0.007936348 |
| BOLA3 | 748.9615 | 0.6192 | 0.1792 | 3.4553 | 0.000549703 | 0.004955515 |
| SLC48A1 | 1059.7497 | 0.6188 | 0.1511 | 4.0953 | 4.22E-05 | 0.000620448 |
| PELI1 | 5987.2043 | 0.6186 | 0.1390 | 4.4503 | 8.58E-06 | 0.000164481 |
| RNFT2 | 104.6507 | 0.6185 | 0.2318 | 2.6680 | 0.007630303 | 0.038963057 |
| SCD | 26788.5546 | 0.6185 | 0.1151 | 5.3746 | 7.68E-08 | 2.93E-06 |
| KCNK1 | 583.4689 | 0.6182 | 0.1638 | 3.7753 | 0.000159835 | 0.001835855 |
| JAKMIP1 | 4634.2123 | 0.6174 | 0.0907 | 6.8056 | 1.01E-11 | 9.17E-10 |
| IL21R | 1590.5116 | 0.6147 | 0.1405 | 4.3763 | 1.21E-05 | 0.000218479 |
| FOXP3 | 3028.3328 | 0.6144 | 0.1599 | 3.8425 | 0.000121798 | 0.001483439 |
| NFIL3 | 20174.6284 | 0.6117 | 0.0929 | 6.5841 | 4.58E-11 | 3.63E-09 |
| MMD | 1199.6000 | 0.6111 | 0.1334 | 4.5804 | 4.64E-06 | 9.89E-05 |
| PTPN14 | 710.4825 | 0.6108 | 0.2032 | 3.0054 | 0.002652501 | 0.017183891 |
| FAM3C | 12347.9259 | 0.6091 | 0.1268 | 4.8040 | 1.56E-06 | 3.82E-05 |
| HBEGF | 303.7537 | 0.6072 | 0.2194 | 2.7679 | 0.005641254 | 0.030994053 |
| SDHAP2 | 3918.4407 | 0.6061 | 0.1679 | 3.6100 | 0.000306233 | 0.003113167 |
| COL6A2 | 1349.6390 | 0.6058 | 0.2198 | 2.7563 | 0.005846161 | 0.031845472 |
| PARP9 | 4588.9118 | 0.6049 | 0.1268 | 4.7705 | 1.84E-06 | 4.42E-05 |
| EED | 6494.7429 | 0.6034 | 0.0897 | 6.7237 | 1.77E-11 | 1.55E-09 |
| IL18RAP | 521.5161 | 0.6023 | 0.2337 | 2.5769 | 0.009970492 | 0.0477139 |
| BATF | 4269.4347 | 0.6021 | 0.1445 | 4.1669 | 3.09E-05 | 0.00047619 |
| FSCN1 | 3170.0263 | 0.6021 | 0.1075 | 5.6022 | 2.12E-08 | 9.64E-07 |
| OTUD7B | 2014.8084 | 0.5993 | 0.0951 | 6.3024 | 2.93E-10 | 1.95E-08 |
| KIF2A | 19994.4746 | 0.5978 | 0.1257 | 4.7562 | 1.97E-06 | 4.67E-05 |
| DUSP6 | 745.5015 | 0.5974 | 0.1394 | 4.2844 | 1.83E-05 | 0.000310673 |
| MAMDC4 | 2325.9711 | 0.5963 | 0.1374 | 4.3396 | 1.43E-05 | 0.000252236 |
| LMAN1 | 27620.5439 | 0.5956 | 0.1449 | 4.1118 | 3.93E-05 | 0.000583451 |
| CRYBB2P1 | 3694.3557 | 0.5949 | 0.1160 | 5.1308 | 2.88E-07 | 9.17E-06 |
| JAK3 | 29392.0531 | 0.5935 | 0.0631 | 9.4110 | 4.92E-21 | 1.86E-18 |
| NPM3 | 678.7342 | 0.5933 | 0.1973 | 3.0066 | 0.002642199 | 0.01713917 |
| PSMD14 | 5856.4543 | 0.5930 | 0.0811 | 7.3113 | 2.65E-13 | 3.31E-11 |
| TIAM1 | 3834.1456 | 0.5928 | 0.1132 | 5.2375 | 1.63E-07 | 5.64E-06 |
| DFNB31 | 5760.1364 | 0.5910 | 0.1472 | 4.0139 | 5.97E-05 | 0.000828854 |
| PUS7 | 1106.3759 | 0.5896 | 0.1321 | 4.4639 | 8.05E-06 | 0.000155663 |
| ZEB2 | 3223.2785 | 0.5892 | 0.1300 | 4.5312 | 5.87E-06 | 0.000120725 |
| LOC729737 | 319.8860 | 0.5888 | 0.2167 | 2.7171 | 0.006584865 | 0.034962883 |
| GPR174 | 1981.4376 | 0.5877 | 0.1575 | 3.7312 | 0.000190546 | 0.002128642 |
| CXorf40A | 2524.4718 | 0.5857 | 0.1067 | 5.4902 | 4.01E-08 | 1.66E-06 |
| MICAL2 | 13390.7250 | 0.5835 | 0.1374 | 4.2468 | 2.17E-05 | 0.000354689 |
| TRAF3 | 8314.2132 | 0.5807 | 0.0737 | 7.8813 | 3.24E-15 | 5.77E-13 |
| BUB1B | 2676.7983 | 0.5797 | 0.1504 | 3.8533 | 0.00011654 | 0.001423989 |
| UNC119 | 5075.7468 | 0.5794 | 0.1126 | 5.1469 | 2.65E-07 | 8.56E-06 |
| CTLA4 | 5613.5363 | 0.5794 | 0.1138 | 5.0910 | 3.56E-07 | 1.10E-05 |
| SLC9A7 | 1589.4616 | 0.5781 | 0.1170 | 4.9408 | 7.78E-07 | 2.14E-05 |
| HPGD | 1842.9807 | 0.5775 | 0.1374 | 4.2041 | 2.62E-05 | 0.000413795 |
| WSB1 | 31775.9427 | 0.5768 | 0.1199 | 4.8107 | 1.50E-06 | 3.71E-05 |
| SYTL3 | 24508.7313 | 0.5765 | 0.1376 | 4.1895 | 2.80E-05 | 0.000436965 |
| IL12RB2 | 3333.5806 | 0.5764 | 0.1224 | 4.7093 | 2.49E-06 | 5.72E-05 |
| MAMLD1 | 1943.0577 | 0.5753 | 0.1316 | 4.3719 | 1.23E-05 | 0.000222357 |
| C10orf128 | 4723.3341 | 0.5753 | 0.1758 | 3.2724 | 0.001066274 | 0.008403476 |
| TANC2 | 1304.5546 | 0.5752 | 0.1190 | 4.8353 | 1.33E-06 | 3.36E-05 |
| PARVB | 3208.9652 | 0.5748 | 0.1146 | 5.0150 | 5.30E-07 | 1.55E-05 |
| KSR1 | 1820.6408 | 0.5726 | 0.1641 | 3.4904 | 0.000482331 | 0.004473462 |
| C14orf49 | 3186.2406 | 0.5716 | 0.1799 | 3.1769 | 0.001488781 | 0.010953694 |
| JARID2 | 6711.5421 | 0.5712 | 0.0954 | 5.9873 | 2.13E-09 | 1.19E-07 |
| TOMM34 | 146.6077 | 0.5699 | 0.1941 | 2.9358 | 0.003326805 | 0.020557427 |
| UHRF1BP1L | 5706.2899 | 0.5699 | 0.0906 | 6.2885 | 3.21E-10 | 2.11E-08 |
| BCL2A1 | 367.6797 | 0.5685 | 0.2188 | 2.5980 | 0.009375955 | 0.045545946 |
| LMNB2 | 4410.5933 | 0.5679 | 0.1108 | 5.1272 | 2.94E-07 | 9.26E-06 |
| RELT | 5406.1080 | 0.5673 | 0.1292 | 4.3892 | 1.14E-05 | 0.000207586 |
| MAP3K6 | 792.6590 | 0.5665 | 0.1234 | 4.5892 | 4.45E-06 | 9.52E-05 |
| TIAM2 | 1139.6918 | 0.5663 | 0.1258 | 4.5016 | 6.75E-06 | 0.000134798 |
| NCDN | 5256.2987 | 0.5662 | 0.1625 | 3.4833 | 0.000495298 | 0.004551867 |
| SLC2A13 | 356.6756 | 0.5653 | 0.1898 | 2.9784 | 0.002898044 | 0.018544173 |
| SCARB1 | 1332.6952 | 0.5648 | 0.1341 | 4.2110 | 2.54E-05 | 0.000403561 |
| GPRIN3 | 11173.1238 | 0.5644 | 0.2190 | 2.5772 | 0.009960671 | 0.047701797 |
| PDCD2L | 211.9409 | 0.5640 | 0.1971 | 2.8620 | 0.004209584 | 0.024796878 |
| RNF122 | 851.7995 | 0.5623 | 0.1311 | 4.2899 | 1.79E-05 | 0.000305218 |
| ZFP36L1 | 8142.5180 | 0.5611 | 0.1159 | 4.8415 | 1.29E-06 | 3.27E-05 |
| SEC23B | 9566.0477 | 0.5610 | 0.1063 | 5.2767 | 1.32E-07 | 4.65E-06 |
| METTL21D | 1210.9139 | 0.5609 | 0.1438 | 3.9009 | 9.58E-05 | 0.001217009 |
| NAGS | 102.8831 | 0.5608 | 0.2005 | 2.7967 | 0.005163043 | 0.029131426 |
| SLC2A3 | 110455.2363 | 0.5602 | 0.0931 | 6.0155 | 1.79E-09 | 1.03E-07 |
| SEMA4A | 1164.7688 | 0.5597 | 0.1443 | 3.8784 | 0.000105154 | 0.001307036 |
| ATP13A3 | 9100.9765 | 0.5596 | 0.1277 | 4.3833 | 1.17E-05 | 0.000212302 |
| PRSS23 | 523.7066 | 0.5580 | 0.1647 | 3.3882 | 0.00070351 | 0.006032877 |
| SNAPC1 | 2083.9379 | 0.5546 | 0.1232 | 4.5021 | 6.73E-06 | 0.00013478 |
| ZFP36 | 8230.3572 | 0.5541 | 0.0927 | 5.9766 | 2.28E-09 | 1.25E-07 |
| FIGNL1 | 3325.6915 | 0.5538 | 0.1334 | 4.1505 | 3.32E-05 | 0.000505443 |
| IPMK | 4093.7322 | 0.5516 | 0.1055 | 5.2261 | 1.73E-07 | 5.86E-06 |
| EXT1 | 1979.6253 | 0.5503 | 0.1299 | 4.2374 | 2.26E-05 | 0.000366517 |
| PLTP | 273.7804 | 0.5495 | 0.2102 | 2.6144 | 0.008938334 | 0.043842469 |
| HSP90AB1 | 40441.9010 | 0.5492 | 0.0632 | 8.6851 | 3.78E-18 | 1.08E-15 |
| THOP1 | 2281.3495 | 0.5488 | 0.1162 | 4.7233 | 2.32E-06 | 5.40E-05 |
| STARD4 | 8001.2917 | 0.5476 | 0.1493 | 3.6682 | 0.000244261 | 0.002577098 |
| SLC19A1 | 1153.7770 | 0.5460 | 0.1023 | 5.3389 | 9.35E-08 | 3.43E-06 |
| FRMD4B | 4575.9258 | 0.5454 | 0.1100 | 4.9598 | 7.06E-07 | 1.97E-05 |
| GPN3 | 913.0601 | 0.5454 | 0.1175 | 4.6435 | 3.43E-06 | 7.51E-05 |
| RBM19 | 5791.7645 | 0.5448 | 0.0852 | 6.3924 | 1.63E-10 | 1.16E-08 |
| TNFSF9 | 532.6092 | 0.5443 | 0.1560 | 3.4887 | 0.00048545 | 0.004488897 |
| PIK3C2B | 5729.5881 | 0.5421 | 0.0920 | 5.8949 | 3.75E-09 | 1.96E-07 |
| CRIM1 | 656.9555 | 0.5411 | 0.2065 | 2.6205 | 0.008778849 | 0.043256562 |
| TNFAIP3 | 33395.3960 | 0.5400 | 0.1406 | 3.8410 | 0.000122536 | 0.001490032 |
| UNQ6494 | 1460.1522 | 0.5400 | 0.1263 | 4.2764 | 1.90E-05 | 0.000320602 |
| SIAH2 | 7174.8537 | 0.5398 | 0.0813 | 6.6404 | 3.13E-11 | 2.60E-09 |
| FAM210A | 5077.0035 | 0.5397 | 0.0900 | 5.9946 | 2.04E-09 | 1.14E-07 |
| LACTB | 2689.6587 | 0.5396 | 0.0739 | 7.3033 | 2.81E-13 | 3.45E-11 |
| RAB11FIP5 | 2870.2084 | 0.5396 | 0.1026 | 5.2611 | 1.43E-07 | 5.02E-06 |
| CADM1 | 331.1460 | 0.5391 | 0.1879 | 2.8684 | 0.004125134 | 0.024421546 |
| PTGS1 | 586.4023 | 0.5388 | 0.1759 | 3.0625 | 0.002194807 | 0.014900797 |
| PVR | 456.1523 | 0.5384 | 0.2085 | 2.5824 | 0.009812955 | 0.04719891 |
| LOC100507582 | 3996.8017 | 0.5382 | 0.1587 | 3.3902 | 0.000698307 | 0.005998457 |
| NPHP4 | 3005.0166 | 0.5376 | 0.1435 | 3.7467 | 0.000179204 | 0.002024506 |
| EGFL6 | 212.1164 | 0.5370 | 0.1810 | 2.9673 | 0.003004603 | 0.019032925 |
| MAP3K8 | 3924.9715 | 0.5363 | 0.1090 | 4.9193 | 8.68E-07 | 2.35E-05 |
| NOL3 | 5662.8027 | 0.5348 | 0.1117 | 4.7877 | 1.69E-06 | 4.10E-05 |
| PLEKHA8 | 3704.8497 | 0.5347 | 0.1232 | 4.3408 | 1.42E-05 | 0.000251486 |
| SPAG4 | 463.7867 | 0.5347 | 0.1451 | 3.6850 | 0.000228668 | 0.002454973 |
| LPAL2 | 370.4623 | 0.5346 | 0.1674 | 3.1937 | 0.001404643 | 0.010444482 |
| RAD51AP1 | 2460.4010 | 0.5343 | 0.1158 | 4.6159 | 3.91E-06 | 8.46E-05 |
| ADAT2 | 2790.5332 | 0.5337 | 0.1104 | 4.8345 | 1.34E-06 | 3.37E-05 |
| STAM | 4271.1625 | 0.5330 | 0.1006 | 5.2985 | 1.17E-07 | 4.19E-06 |
| LRMP | 6436.3878 | 0.5326 | 0.1200 | 4.4382 | 9.07E-06 | 0.000172418 |
| SAP30 | 5526.3936 | 0.5322 | 0.1196 | 4.4481 | 8.66E-06 | 0.000165647 |
| MTHFD1L | 3147.3945 | 0.5317 | 0.0901 | 5.8983 | 3.67E-09 | 1.94E-07 |
| HACE1 | 3606.4322 | 0.5304 | 0.1446 | 3.6692 | 0.000243356 | 0.0025743 |
| IL2RB | 38769.7227 | 0.5301 | 0.1467 | 3.6132 | 0.000302497 | 0.003083471 |
| WDR36 | 2312.5569 | 0.5295 | 0.0886 | 5.9795 | 2.24E-09 | 1.24E-07 |
| HEATR5A | 1752.3970 | 0.5287 | 0.1371 | 3.8564 | 0.000115061 | 0.001409328 |
| B4GALT5 | 4125.9113 | 0.5283 | 0.0820 | 6.4434 | 1.17E-10 | 8.49E-09 |
| SPAG1 | 1080.5457 | 0.5283 | 0.1308 | 4.0388 | 5.37E-05 | 0.000759554 |
| RNF19B | 3234.5284 | 0.5276 | 0.1232 | 4.2806 | 1.86E-05 | 0.000315365 |
| SRGN | 28624.3108 | 0.5273 | 0.0960 | 5.4935 | 3.94E-08 | 1.64E-06 |
| IMPAD1 | 6966.2856 | 0.5272 | 0.1134 | 4.6472 | 3.36E-06 | 7.41E-05 |
| SLC39A14 | 2178.0766 | 0.5271 | 0.1417 | 3.7203 | 0.000198952 | 0.002201569 |
| NGLY1 | 11010.3901 | 0.5267 | 0.0984 | 5.3554 | 8.54E-08 | 3.20E-06 |
| IPPK | 1805.3725 | 0.5264 | 0.1162 | 4.5301 | 5.90E-06 | 0.000120785 |
| C1orf55 | 2383.7760 | 0.5263 | 0.1060 | 4.9636 | 6.92E-07 | 1.95E-05 |
| NRN1 | 275.1832 | 0.5257 | 0.1990 | 2.6414 | 0.008255266 | 0.041377535 |
| PLXNA1 | 705.4295 | 0.5253 | 0.1735 | 3.0269 | 0.002470523 | 0.01628392 |
| FAM13A | 4979.7112 | 0.5252 | 0.1781 | 2.9486 | 0.003191788 | 0.019922041 |
| ZNF331 | 2061.3815 | 0.5249 | 0.1567 | 3.3508 | 0.000805797 | 0.006723267 |
| ZMIZ2 | 8882.0291 | 0.5248 | 0.0737 | 7.1171 | 1.10E-12 | 1.21E-10 |
| CENPO | 642.7531 | 0.5241 | 0.1405 | 3.7309 | 0.000190814 | 0.002128642 |
| LONP1 | 12591.5362 | 0.5240 | 0.1148 | 4.5643 | 5.01E-06 | 0.000105455 |
| TIMD4 | 388.8436 | 0.5237 | 0.1943 | 2.6951 | 0.007037594 | 0.036634955 |
| RSPH3 | 668.0499 | 0.5236 | 0.1571 | 3.3316 | 0.000863552 | 0.007107159 |
| DLGAP4 | 3936.5768 | 0.5230 | 0.1537 | 3.4024 | 0.000667867 | 0.005786264 |
| SIX4 | 236.8060 | 0.5228 | 0.1978 | 2.6424 | 0.008231721 | 0.041286884 |
| CDYL2 | 1337.6382 | 0.5226 | 0.0962 | 5.4345 | 5.50E-08 | 2.18E-06 |
| TFRC | 20953.3747 | 0.5222 | 0.1160 | 4.5017 | 6.74E-06 | 0.000134798 |
| KCTD17 | 1855.5857 | 0.5220 | 0.0955 | 5.4682 | 4.55E-08 | 1.85E-06 |
| RTTN | 1747.8910 | 0.5201 | 0.1300 | 4.0010 | 6.31E-05 | 0.000868209 |
| FEM1C | 4837.4673 | 0.5197 | 0.1037 | 5.0097 | 5.45E-07 | 1.58E-05 |
| HBS1L | 7889.8970 | 0.5193 | 0.0719 | 7.2227 | 5.10E-13 | 6.02E-11 |
| STX11 | 1297.0162 | 0.5191 | 0.1140 | 4.5547 | 5.25E-06 | 0.000109906 |
| UBR5 | 33886.9079 | 0.5177 | 0.1412 | 3.6678 | 0.000244613 | 0.002578583 |
| C16orf87 | 2567.6579 | 0.5171 | 0.0925 | 5.5920 | 2.24E-08 | 1.01E-06 |
| PPP1R15A | 10220.6451 | 0.5169 | 0.1262 | 4.0943 | 4.23E-05 | 0.000621837 |
| TFB1M | 1598.0729 | 0.5164 | 0.0951 | 5.4324 | 5.56E-08 | 2.20E-06 |
| PPFIBP1 | 1404.7438 | 0.5158 | 0.1749 | 2.9484 | 0.003193989 | 0.019923907 |
| PKIA | 3045.4394 | 0.5157 | 0.0974 | 5.2945 | 1.19E-07 | 4.27E-06 |
| MCM10 | 958.1397 | 0.5151 | 0.1356 | 3.7984 | 0.000145656 | 0.001704056 |
| RNF144A | 3515.8288 | 0.5149 | 0.1014 | 5.0782 | 3.81E-07 | 1.16E-05 |
| LOC100630923 | 224.8851 | 0.5147 | 0.1994 | 2.5811 | 0.009849093 | 0.047297531 |
| POLR3G | 485.2347 | 0.5144 | 0.1706 | 3.0156 | 0.002565101 | 0.016710717 |
| DNAJB5 | 341.4798 | 0.5134 | 0.1776 | 2.8912 | 0.003837281 | 0.023036435 |
| TSR1 | 1887.4856 | 0.5122 | 0.1314 | 3.8971 | 9.74E-05 | 0.001230273 |
| ITPR1 | 7185.7195 | 0.5121 | 0.1130 | 4.5328 | 5.82E-06 | 0.000120269 |
| TOP1 | 2775.2231 | 0.5117 | 0.1717 | 2.9804 | 0.002878355 | 0.018449525 |
| GARS | 16707.7916 | 0.5106 | 0.1095 | 4.6613 | 3.14E-06 | 6.99E-05 |
| TSHZ2 | 2063.2336 | 0.5095 | 0.1576 | 3.2336 | 0.001222616 | 0.009383314 |
| AURKA | 2754.9522 | 0.5090 | 0.0859 | 5.9225 | 3.17E-09 | 1.70E-07 |
| KLHL15 | 1691.9400 | 0.5082 | 0.1205 | 4.2162 | 2.49E-05 | 0.000396446 |
| ANK3 | 3657.2731 | 0.5078 | 0.1156 | 4.3937 | 1.11E-05 | 0.000204604 |
| CD9 | 1596.9369 | 0.5075 | 0.1334 | 3.8040 | 0.000142366 | 0.001678371 |
| SPIRE1 | 1027.3788 | 0.5074 | 0.1222 | 4.1538 | 3.27E-05 | 0.000500699 |
| BTAF1 | 15848.8088 | 0.5073 | 0.1367 | 3.7105 | 0.000206842 | 0.002272254 |
| WDHD1 | 1596.3769 | 0.5067 | 0.1186 | 4.2721 | 1.94E-05 | 0.000325105 |
| JUND | 7468.2327 | 0.5053 | 0.1494 | 3.3815 | 0.000720859 | 0.0061607 |
| MPP1 | 2992.9159 | 0.5048 | 0.1112 | 4.5394 | 5.64E-06 | 0.000117406 |
| IGFLR1 | 4040.8848 | 0.5046 | 0.1177 | 4.2860 | 1.82E-05 | 0.000309615 |
| NCAPD3 | 5677.9992 | 0.5033 | 0.1146 | 4.3933 | 1.12E-05 | 0.00020471 |
| ATN1 | 5055.2331 | 0.5025 | 0.1077 | 4.6672 | 3.05E-06 | 6.84E-05 |
| DIAPH3 | 1509.3748 | 0.5025 | 0.1325 | 3.7920 | 0.000149464 | 0.001735963 |
| TAF13 | 318.9423 | 0.5020 | 0.1856 | 2.7045 | 0.006839889 | 0.035976009 |
| FANCI | 8062.3526 | 0.5012 | 0.1139 | 4.4019 | 1.07E-05 | 0.000198672 |
| ARHGAP31 | 1065.0059 | 0.5012 | 0.1511 | 3.3165 | 0.000911468 | 0.007412785 |
| KIAA0247 | 7058.4240 | 0.5003 | 0.1117 | 4.4777 | 7.54E-06 | 0.000147268 |
| CLK1 | 20389.5951 | 0.5000 | 0.1244 | 4.0185 | 5.86E-05 | 0.000817494 |
| CLPB | 902.2029 | 0.4998 | 0.0878 | 5.6953 | 1.23E-08 | 5.82E-07 |
| ENO2 | 49796.3070 | 0.4988 | 0.1116 | 4.4686 | 7.87E-06 | 0.00015272 |
| KDM6B | 4952.6627 | 0.4979 | 0.1582 | 3.1473 | 0.001647813 | 0.01186975 |
| RAB21 | 4632.8957 | 0.4974 | 0.1011 | 4.9216 | 8.59E-07 | 2.32E-05 |
| KDM5C | 11822.1376 | 0.4960 | 0.1030 | 4.8162 | 1.46E-06 | 3.64E-05 |
| ZDHHC14 | 2410.3590 | 0.4957 | 0.1261 | 3.9302 | 8.49E-05 | 0.001098154 |
| BCAT1 | 12962.0620 | 0.4954 | 0.1325 | 3.7385 | 0.000185157 | 0.002082426 |
| PAPD7 | 8985.2539 | 0.4951 | 0.1060 | 4.6693 | 3.02E-06 | 6.78E-05 |
| PEMT | 666.8413 | 0.4949 | 0.1450 | 3.4125 | 0.000643595 | 0.005611336 |
| PRIC285 | 3853.1135 | 0.4939 | 0.1720 | 2.8715 | 0.004084877 | 0.024201618 |
| LONRF1 | 999.2918 | 0.4937 | 0.1510 | 3.2692 | 0.001078665 | 0.008476343 |
| STAT1 | 37483.9967 | 0.4937 | 0.1237 | 3.9915 | 6.57E-05 | 0.000893933 |
| RANBP1 | 2861.7285 | 0.4934 | 0.1425 | 3.4622 | 0.00053578 | 0.004850232 |
| FASN | 6414.9025 | 0.4925 | 0.1706 | 2.8865 | 0.00389532 | 0.023336441 |
| INPP5F | 2868.6825 | 0.4917 | 0.1288 | 3.8166 | 0.000135289 | 0.001619075 |
| TUBE1 | 2454.8770 | 0.4911 | 0.1334 | 3.6825 | 0.000230918 | 0.00246994 |
| CYCS | 5811.2155 | 0.4906 | 0.0773 | 6.3438 | 2.24E-10 | 1.53E-08 |
| PPP1R14B | 3055.9527 | 0.4902 | 0.1416 | 3.4606 | 0.000539034 | 0.00487386 |
| PHF6 | 4421.7492 | 0.4901 | 0.1018 | 4.8140 | 1.48E-06 | 3.67E-05 |
| PLCXD2 | 1864.3429 | 0.4895 | 0.1735 | 2.8210 | 0.004787553 | 0.027526158 |
| VDAC3 | 15424.4371 | 0.4894 | 0.1214 | 4.0308 | 5.56E-05 | 0.000779433 |
| PSD | 585.6188 | 0.4883 | 0.1498 | 3.2605 | 0.001112172 | 0.00866761 |
| EIF2C2 | 4094.7270 | 0.4873 | 0.1306 | 3.7310 | 0.000190696 | 0.002128642 |
| CD80 | 1144.2050 | 0.4872 | 0.1717 | 2.8381 | 0.004537769 | 0.026310013 |
| TRIO | 1040.3240 | 0.4867 | 0.1555 | 3.1311 | 0.001741505 | 0.012385402 |
| HSF4 | 4516.8187 | 0.4867 | 0.0985 | 4.9399 | 7.82E-07 | 2.14E-05 |
| KDM5B | 11613.0006 | 0.4865 | 0.0898 | 5.4191 | 5.99E-08 | 2.35E-06 |
| CXorf40B | 4869.9670 | 0.4865 | 0.1113 | 4.3689 | 1.25E-05 | 0.000224108 |
| USP12 | 6938.2154 | 0.4862 | 0.0961 | 5.0586 | 4.22E-07 | 1.26E-05 |
| ZNF148 | 1152.5305 | 0.4858 | 0.1279 | 3.7985 | 0.000145575 | 0.001704056 |
| DRAM1 | 964.2841 | 0.4856 | 0.1197 | 4.0569 | 4.97E-05 | 0.000709725 |
| ERO1L | 31135.8009 | 0.4855 | 0.1270 | 3.8228 | 0.00013193 | 0.001591476 |
| RPL22L1 | 2331.1880 | 0.4851 | 0.1494 | 3.2460 | 0.001170546 | 0.009029498 |
| TNF | 1610.6782 | 0.4836 | 0.1499 | 3.2254 | 0.001257972 | 0.009581746 |
| AHI1 | 9092.6759 | 0.4824 | 0.1024 | 4.7132 | 2.44E-06 | 5.62E-05 |
| BUB1 | 5280.3711 | 0.4814 | 0.1064 | 4.5246 | 6.05E-06 | 0.000123006 |
| RBBP9 | 2398.3480 | 0.4812 | 0.1312 | 3.6682 | 0.000244302 | 0.002577098 |
| CEP135 | 1897.0901 | 0.4809 | 0.1071 | 4.4878 | 7.20E-06 | 0.000141908 |
| SSH1 | 10783.7084 | 0.4805 | 0.1629 | 2.9495 | 0.003182519 | 0.019893373 |
| SPAG9 | 9148.2005 | 0.4800 | 0.1027 | 4.6719 | 2.98E-06 | 6.71E-05 |
| C1orf96 | 2502.4934 | 0.4797 | 0.0967 | 4.9582 | 7.11E-07 | 1.97E-05 |
| MYH10 | 2606.3997 | 0.4795 | 0.1290 | 3.7170 | 0.000201579 | 0.002222514 |
| UBA6 | 8993.0418 | 0.4795 | 0.1181 | 4.0612 | 4.88E-05 | 0.000700773 |
| OTUD4 | 6578.1198 | 0.4794 | 0.1200 | 3.9942 | 6.49E-05 | 0.000886932 |
| C12orf48 | 1243.4214 | 0.4794 | 0.1326 | 3.6142 | 0.000301283 | 0.003075246 |
| PIK3R6 | 1038.8173 | 0.4792 | 0.1640 | 2.9222 | 0.003476131 | 0.021271618 |
| GPR107 | 2378.1514 | 0.4770 | 0.1420 | 3.3592 | 0.000781701 | 0.006572977 |
| SDHAP1 | 4810.9928 | 0.4765 | 0.1109 | 4.2982 | 1.72E-05 | 0.000295603 |
| PCBP3 | 1145.3543 | 0.4762 | 0.1637 | 2.9084 | 0.00363228 | 0.022066465 |
| YEATS2 | 11407.8206 | 0.4749 | 0.1527 | 3.1101 | 0.001870185 | 0.013103419 |
| SYNGR2 | 5402.0564 | 0.4742 | 0.1116 | 4.2487 | 2.15E-05 | 0.000352028 |
| SMAP2 | 30578.2897 | 0.4737 | 0.0707 | 6.7042 | 2.03E-11 | 1.73E-09 |
| DTL | 2545.0537 | 0.4736 | 0.1290 | 3.6716 | 0.000241061 | 0.002551806 |
| GRPEL1 | 1897.3382 | 0.4721 | 0.1051 | 4.4913 | 7.08E-06 | 0.000139976 |
| ZCCHC24 | 602.3287 | 0.4715 | 0.1776 | 2.6546 | 0.00793944 | 0.040113532 |
| DHX35 | 1183.4641 | 0.4715 | 0.1187 | 3.9733 | 7.09E-05 | 0.000953177 |
| ACTR1B | 11837.5948 | 0.4714 | 0.0894 | 5.2742 | 1.33E-07 | 4.69E-06 |
| GMNN | 3157.5027 | 0.4707 | 0.0962 | 4.8931 | 9.92E-07 | 2.62E-05 |
| TAF1A | 480.1394 | 0.4707 | 0.1486 | 3.1674 | 0.001537896 | 0.011227679 |
| UGCG | 982.8296 | 0.4706 | 0.1769 | 2.6597 | 0.007821365 | 0.039650907 |
| LIMD1 | 4088.3357 | 0.4706 | 0.1318 | 3.5705 | 0.000356362 | 0.00350956 |
| AHR | 3966.7246 | 0.4705 | 0.1075 | 4.3762 | 1.21E-05 | 0.000218479 |
| BCL2L1 | 3203.8520 | 0.4703 | 0.1002 | 4.6940 | 2.68E-06 | 6.11E-05 |
| ASCC3 | 7693.8381 | 0.4680 | 0.0990 | 4.7277 | 2.27E-06 | 5.30E-05 |
| RDX | 8000.3736 | 0.4676 | 0.0688 | 6.7931 | 1.10E-11 | 9.82E-10 |
| DENND4A | 5067.6300 | 0.4668 | 0.0854 | 5.4645 | 4.64E-08 | 1.88E-06 |
| UBAC2 | 250.7395 | 0.4661 | 0.1376 | 3.3884 | 0.000702999 | 0.006031916 |
| MAST1 | 476.1674 | 0.4658 | 0.1291 | 3.6082 | 0.000308311 | 0.003132183 |
| FLVCR2 | 618.0370 | 0.4656 | 0.1584 | 2.9391 | 0.003291971 | 0.020394504 |
| GMDS | 1378.6848 | 0.4655 | 0.1509 | 3.0844 | 0.002039627 | 0.014052905 |
| XAF1 | 5316.7735 | 0.4653 | 0.1549 | 3.0050 | 0.002655806 | 0.017197933 |
| RNMT | 11465.4151 | 0.4650 | 0.1206 | 3.8555 | 0.00011549 | 0.00141344 |
| CEP72 | 843.2889 | 0.4648 | 0.1167 | 3.9840 | 6.78E-05 | 0.000917007 |
| TXNDC17 | 1236.4882 | 0.4641 | 0.1667 | 2.7838 | 0.005372488 | 0.030009875 |
| CCDC6 | 4999.5578 | 0.4625 | 0.0770 | 6.0072 | 1.89E-09 | 1.07E-07 |
| LINC00271 | 3702.9870 | 0.4624 | 0.1591 | 2.9066 | 0.003653452 | 0.022146367 |
| TRIP10 | 2377.2547 | 0.4617 | 0.0689 | 6.6980 | 2.11E-11 | 1.80E-09 |
| BANP | 2386.7302 | 0.4617 | 0.0945 | 4.8843 | 1.04E-06 | 2.72E-05 |
| PHKA1 | 2188.2110 | 0.4616 | 0.1618 | 2.8539 | 0.004318532 | 0.025300708 |
| ATP6V1B2 | 4295.3410 | 0.4607 | 0.1601 | 2.8773 | 0.004010909 | 0.023868223 |
| IFIH1 | 4218.6087 | 0.4601 | 0.0966 | 4.7626 | 1.91E-06 | 4.55E-05 |
| TTC7A | 3292.6963 | 0.4599 | 0.1247 | 3.6870 | 0.00022692 | 0.002446627 |
| NOTCH1 | 3664.0822 | 0.4589 | 0.1375 | 3.3377 | 0.000844869 | 0.006974939 |
| KIF18A | 1641.7376 | 0.4589 | 0.1165 | 3.9379 | 8.22E-05 | 0.001073681 |
| JMY | 5504.0139 | 0.4586 | 0.1138 | 4.0309 | 5.56E-05 | 0.000779433 |
| LYST | 11947.6501 | 0.4582 | 0.1369 | 3.3477 | 0.000814998 | 0.00677761 |
| NUP35 | 591.3932 | 0.4577 | 0.0979 | 4.6751 | 2.94E-06 | 6.61E-05 |
| MRTO4 | 991.2776 | 0.4576 | 0.1181 | 3.8748 | 0.000106698 | 0.001322962 |
| SLFN12L | 14643.3880 | 0.4574 | 0.1060 | 4.3154 | 1.59E-05 | 0.000277682 |
| CD82 | 14446.8123 | 0.4569 | 0.0738 | 6.1944 | 5.85E-10 | 3.61E-08 |
| GNL3 | 4960.7055 | 0.4567 | 0.0834 | 5.4783 | 4.29E-08 | 1.76E-06 |
| NDUFV2 | 13340.1069 | 0.4564 | 0.1029 | 4.4353 | 9.20E-06 | 0.000174318 |
| PPP2R5B | 3227.1639 | 0.4564 | 0.0937 | 4.8733 | 1.10E-06 | 2.85E-05 |
| CCDC41 | 978.4085 | 0.4562 | 0.1198 | 3.8073 | 0.000140515 | 0.001666906 |
| SASS6 | 2412.2077 | 0.4559 | 0.1083 | 4.2110 | 2.54E-05 | 0.000403561 |
| E2F7 | 857.3288 | 0.4558 | 0.1620 | 2.8129 | 0.004910024 | 0.028035971 |
| MAP2K1 | 7821.2789 | 0.4558 | 0.1044 | 4.3675 | 1.26E-05 | 0.000225212 |
| CCDC47 | 7189.0348 | 0.4558 | 0.0973 | 4.6842 | 2.81E-06 | 6.36E-05 |
| CENPI | 590.9368 | 0.4552 | 0.1633 | 2.7882 | 0.005299613 | 0.029691573 |
| B4GALT2 | 625.4770 | 0.4551 | 0.1196 | 3.8060 | 0.000141234 | 0.001670399 |
| CDC25A | 458.0877 | 0.4551 | 0.1728 | 2.6340 | 0.008438837 | 0.042005358 |
| ATP1B1 | 361.7715 | 0.4547 | 0.1688 | 2.6937 | 0.007065914 | 0.036755874 |
| HIPK2 | 10595.2528 | 0.4547 | 0.1334 | 3.4091 | 0.000651776 | 0.005676114 |
| SDF4 | 13521.3778 | 0.4545 | 0.0888 | 5.1173 | 3.10E-07 | 9.68E-06 |
| TMEM194A | 6704.8954 | 0.4544 | 0.1195 | 3.8012 | 0.000143985 | 0.001689734 |
| DUSP11 | 2384.7757 | 0.4516 | 0.1005 | 4.4953 | 6.95E-06 | 0.000137751 |
| SLC4A10 | 348.3369 | 0.4506 | 0.1398 | 3.2223 | 0.001271671 | 0.009656911 |
| SERTAD2 | 4680.4306 | 0.4502 | 0.1064 | 4.2306 | 2.33E-05 | 0.000375883 |
| SQLE | 5512.7647 | 0.4501 | 0.0822 | 5.4782 | 4.30E-08 | 1.76E-06 |
| MAP3K2 | 8508.1041 | 0.4490 | 0.1217 | 3.6899 | 0.000224331 | 0.002424272 |
| MTHFD1 | 5384.7543 | 0.4488 | 0.1280 | 3.5073 | 0.000452703 | 0.004258733 |
| PPIP5K1 | 668.1152 | 0.4487 | 0.1151 | 3.8989 | 9.66E-05 | 0.001224364 |
| ORC1 | 2377.5941 | 0.4487 | 0.1206 | 3.7220 | 0.000197613 | 0.002189956 |
| PLAGL1 | 2116.6202 | 0.4486 | 0.1676 | 2.6761 | 0.007447735 | 0.038359512 |
| EIF4ENIF1 | 3214.0959 | 0.4481 | 0.0598 | 7.4935 | 6.71E-14 | 9.57E-12 |
| PDIA5 | 562.8014 | 0.4477 | 0.1630 | 2.7465 | 0.006023804 | 0.032526001 |
| NAF1 | 930.3380 | 0.4473 | 0.1246 | 3.5910 | 0.000329359 | 0.003306049 |
| PATL1 | 10098.4856 | 0.4469 | 0.1132 | 3.9479 | 7.88E-05 | 0.001040508 |
| OAS1 | 2227.5051 | 0.4465 | 0.1659 | 2.6914 | 0.007115837 | 0.03693935 |
| TRIT1 | 3379.7120 | 0.4465 | 0.0836 | 5.3407 | 9.26E-08 | 3.42E-06 |
| SLC17A9 | 1540.7984 | 0.4461 | 0.1128 | 3.9541 | 7.68E-05 | 0.001021253 |
| SH3D21 | 6418.3320 | 0.4452 | 0.1248 | 3.5675 | 0.00036037 | 0.003539819 |
| TIGIT | 6509.8997 | 0.4451 | 0.1207 | 3.6867 | 0.000227191 | 0.002447805 |
| PAICS | 15306.3324 | 0.4450 | 0.0916 | 4.8578 | 1.19E-06 | 3.04E-05 |
| POU2F1 | 2531.3960 | 0.4448 | 0.0810 | 5.4919 | 3.98E-08 | 1.65E-06 |
| SLC39A8 | 6838.0579 | 0.4447 | 0.0999 | 4.4502 | 8.58E-06 | 0.000164481 |
| BTG3 | 7456.7860 | 0.4431 | 0.0918 | 4.8295 | 1.37E-06 | 3.43E-05 |
| MED12 | 9367.6941 | 0.4423 | 0.1273 | 3.4743 | 0.000512243 | 0.004687661 |
| TRIP13 | 765.2196 | 0.4413 | 0.1187 | 3.7192 | 0.000199863 | 0.002208194 |
| PGAM1 | 27207.4956 | 0.4410 | 0.0588 | 7.4972 | 6.52E-14 | 9.40E-12 |
| PBK | 916.0468 | 0.4396 | 0.1409 | 3.1201 | 0.001807707 | 0.012754282 |
| PFKFB3 | 13359.7393 | 0.4392 | 0.1061 | 4.1394 | 3.48E-05 | 0.000528235 |
| SPATS2L | 1977.6045 | 0.4387 | 0.1284 | 3.4169 | 0.000633297 | 0.005543918 |
| BTG2 | 6083.1799 | 0.4383 | 0.0967 | 4.5320 | 5.84E-06 | 0.000120392 |
| POM121 | 10200.2999 | 0.4380 | 0.0837 | 5.2352 | 1.65E-07 | 5.67E-06 |
| GPD2 | 1467.7080 | 0.4377 | 0.1091 | 4.0119 | 6.02E-05 | 0.000835218 |
| LSM12 | 2060.7263 | 0.4376 | 0.0966 | 4.5300 | 5.90E-06 | 0.000120785 |
| MRPL17 | 2475.8011 | 0.4375 | 0.1404 | 3.1165 | 0.001830212 | 0.012887534 |
| C14orf182 | 1723.6569 | 0.4374 | 0.1491 | 2.9346 | 0.003340283 | 0.020623864 |
| SLC7A1 | 5968.2864 | 0.4373 | 0.1327 | 3.2948 | 0.000985011 | 0.007908841 |
| SLCO4A1 | 5662.3545 | 0.4372 | 0.0759 | 5.7636 | 8.23E-09 | 4.02E-07 |
| TMEM185A | 1640.0465 | 0.4370 | 0.1259 | 3.4718 | 0.000516916 | 0.004716153 |
| TCTN3 | 3876.5391 | 0.4369 | 0.1110 | 3.9354 | 8.31E-05 | 0.001078899 |
| SEC14L1 | 5997.6422 | 0.4368 | 0.0910 | 4.8021 | 1.57E-06 | 3.85E-05 |
| HK1 | 14346.6349 | 0.4364 | 0.0844 | 5.1713 | 2.32E-07 | 7.61E-06 |
| MTF2 | 5880.9392 | 0.4363 | 0.0825 | 5.2887 | 1.23E-07 | 4.37E-06 |
| MAPK8 | 3863.8774 | 0.4362 | 0.1093 | 3.9928 | 6.53E-05 | 0.000889693 |
| SMOX | 1542.3175 | 0.4358 | 0.0968 | 4.5043 | 6.66E-06 | 0.000133951 |
| BTN2A2 | 1908.5179 | 0.4346 | 0.1089 | 3.9913 | 6.57E-05 | 0.000893933 |
| HDGFRP3 | 4889.7691 | 0.4345 | 0.0836 | 5.1983 | 2.01E-07 | 6.64E-06 |
| IKZF4 | 1415.7335 | 0.4342 | 0.1400 | 3.1018 | 0.001923305 | 0.01341348 |
| MED10 | 6281.5431 | 0.4341 | 0.1457 | 2.9784 | 0.002897359 | 0.018544173 |
| RRP12 | 649.7608 | 0.4330 | 0.1454 | 2.9781 | 0.002900477 | 0.018544173 |
| INSIG1 | 24823.7557 | 0.4329 | 0.0882 | 4.9102 | 9.10E-07 | 2.45E-05 |
| NUF2 | 3420.2738 | 0.4327 | 0.1159 | 3.7328 | 0.00018933 | 0.002119976 |
| HSPH1 | 5234.8265 | 0.4320 | 0.0830 | 5.2028 | 1.96E-07 | 6.53E-06 |
| AGK | 3755.9578 | 0.4320 | 0.0851 | 5.0746 | 3.88E-07 | 1.17E-05 |
| ISCU | 5862.9098 | 0.4319 | 0.0823 | 5.2512 | 1.51E-07 | 5.29E-06 |
| CXCR4 | 13972.0792 | 0.4319 | 0.1137 | 3.7984 | 0.000145615 | 0.001704056 |
| PPP4R1L | 2389.3187 | 0.4315 | 0.1481 | 2.9131 | 0.003579096 | 0.021804666 |
| RBM7 | 3012.0436 | 0.4313 | 0.0852 | 5.0596 | 4.20E-07 | 1.26E-05 |
| GTPBP1 | 5920.3130 | 0.4312 | 0.0626 | 6.8840 | 5.82E-12 | 5.47E-10 |
| EIF4E | 2673.8779 | 0.4312 | 0.0834 | 5.1714 | 2.32E-07 | 7.61E-06 |
| TAX1BP1 | 12580.0858 | 0.4302 | 0.0626 | 6.8731 | 6.28E-12 | 5.86E-10 |
| HJURP | 3641.7921 | 0.4301 | 0.1094 | 3.9304 | 8.48E-05 | 0.001098154 |
| N4BP3 | 1337.8530 | 0.4299 | 0.0788 | 5.4540 | 4.92E-08 | 1.98E-06 |
| C5orf30 | 1045.3445 | 0.4296 | 0.1090 | 3.9397 | 8.16E-05 | 0.001069267 |
| ITGAX | 13239.3823 | 0.4292 | 0.1160 | 3.6992 | 0.000216243 | 0.002355013 |
| PSAT1 | 3399.0478 | 0.4291 | 0.1466 | 2.9262 | 0.003431613 | 0.021053648 |
| IRGQ | 2135.2576 | 0.4285 | 0.0898 | 4.7733 | 1.81E-06 | 4.36E-05 |
| GLUL | 13444.6867 | 0.4285 | 0.1188 | 3.6054 | 0.000311648 | 0.0031576 |
| ZC3H15 | 4851.5128 | 0.4280 | 0.1219 | 3.5122 | 0.00044449 | 0.004194509 |
| ZSCAN2 | 1152.5103 | 0.4280 | 0.0790 | 5.4159 | 6.10E-08 | 2.38E-06 |
| CYB5A | 1122.5574 | 0.4276 | 0.0858 | 4.9822 | 6.29E-07 | 1.79E-05 |
| CASC3 | 9842.8717 | 0.4271 | 0.0831 | 5.1398 | 2.75E-07 | 8.80E-06 |
| TRAF2 | 3974.3951 | 0.4261 | 0.0863 | 4.9354 | 8.00E-07 | 2.19E-05 |
| EME1 | 931.8203 | 0.4258 | 0.1418 | 3.0021 | 0.002680995 | 0.017323967 |
| GTPBP4 | 2592.5682 | 0.4252 | 0.0945 | 4.4986 | 6.84E-06 | 0.000136123 |
| SLC41A1 | 3358.3625 | 0.4249 | 0.1277 | 3.3273 | 0.000876766 | 0.007196333 |
| PIGA | 2186.5914 | 0.4246 | 0.1096 | 3.8745 | 0.000106832 | 0.001323539 |
| PGAP1 | 4682.2220 | 0.4240 | 0.1186 | 3.5736 | 0.000352096 | 0.00348307 |
| MAD2L2 | 1691.1390 | 0.4223 | 0.1419 | 2.9765 | 0.00291563 | 0.018617451 |
| SLC3A2 | 14581.2503 | 0.4220 | 0.1114 | 3.7887 | 0.000151427 | 0.001752789 |
| FAM89B | 2417.4732 | 0.4207 | 0.1213 | 3.4675 | 0.000525285 | 0.004769496 |
| MTX3 | 7814.3578 | 0.4204 | 0.1201 | 3.5015 | 0.000462703 | 0.004325899 |
| FAH | 2689.6720 | 0.4201 | 0.0936 | 4.4860 | 7.26E-06 | 0.000142807 |
| VPS37A | 3236.6653 | 0.4200 | 0.1036 | 4.0546 | 5.02E-05 | 0.000716092 |
| DIEXF | 1786.2055 | 0.4196 | 0.1077 | 3.8957 | 9.79E-05 | 0.001236345 |
| PIF1 | 2555.6433 | 0.4190 | 0.1086 | 3.8579 | 0.000114365 | 0.001404224 |
| UPP1 | 5341.1858 | 0.4190 | 0.1005 | 4.1709 | 3.03E-05 | 0.000469839 |
| EXO1 | 1182.6297 | 0.4188 | 0.1278 | 3.2766 | 0.001050537 | 0.008315786 |
| QSOX2 | 3935.0738 | 0.4187 | 0.0966 | 4.3367 | 1.45E-05 | 0.000254119 |
| JMJD6 | 5032.5350 | 0.4185 | 0.0898 | 4.6599 | 3.16E-06 | 7.03E-05 |
| E2F4 | 7309.0693 | 0.4182 | 0.0952 | 4.3917 | 1.12E-05 | 0.000205709 |
| TTK | 2582.1926 | 0.4181 | 0.1460 | 2.8639 | 0.004184789 | 0.024704197 |
| SLC1A5 | 11119.1711 | 0.4171 | 0.1226 | 3.4030 | 0.000666524 | 0.005783004 |
| TMEM106C | 4020.4069 | 0.4170 | 0.0926 | 4.5007 | 6.77E-06 | 0.000134997 |
| OSBPL3 | 9195.3668 | 0.4164 | 0.1059 | 3.9318 | 8.43E-05 | 0.001092785 |
| MORF4L2 | 10873.1360 | 0.4163 | 0.0922 | 4.5127 | 6.40E-06 | 0.000129431 |
| NFKBIE | 1113.3357 | 0.4161 | 0.1360 | 3.0595 | 0.002216946 | 0.014953948 |
| FAM174A | 485.2590 | 0.4160 | 0.1486 | 2.8000 | 0.005109628 | 0.028948815 |
| FAS | 9224.5531 | 0.4158 | 0.1002 | 4.1510 | 3.31E-05 | 0.000505295 |
| CBS | 897.7141 | 0.4156 | 0.1488 | 2.7942 | 0.005203525 | 0.029294277 |
| SH3GL1 | 4560.5522 | 0.4155 | 0.0833 | 4.9859 | 6.17E-07 | 1.77E-05 |
| TESK1 | 2428.7317 | 0.4155 | 0.0971 | 4.2795 | 1.87E-05 | 0.000316531 |
| CCND2 | 17121.1455 | 0.4154 | 0.1158 | 3.5885 | 0.000332579 | 0.003327328 |
| IKBKB | 13561.8887 | 0.4147 | 0.1355 | 3.0609 | 0.002206966 | 0.014937257 |
| BRI3BP | 430.9808 | 0.4147 | 0.1558 | 2.6609 | 0.00779262 | 0.039579457 |
| PPP1R15B | 12301.0073 | 0.4141 | 0.0936 | 4.4243 | 9.68E-06 | 0.000182521 |
| PDE12 | 2464.6719 | 0.4134 | 0.0873 | 4.7373 | 2.17E-06 | 5.07E-05 |
| XYLT2 | 4801.8603 | 0.4120 | 0.0965 | 4.2692 | 1.96E-05 | 0.000328497 |
| POLD2 | 2818.2284 | 0.4118 | 0.1055 | 3.9016 | 9.56E-05 | 0.001214825 |
| RAN | 15515.3431 | 0.4116 | 0.1051 | 3.9169 | 8.97E-05 | 0.001150695 |
| ANKRD10 | 18464.4023 | 0.4113 | 0.1142 | 3.6023 | 0.000315405 | 0.003191392 |
| STARD8 | 675.7511 | 0.4105 | 0.1400 | 2.9330 | 0.003357392 | 0.020712591 |
| DPP4 | 13553.7965 | 0.4093 | 0.0707 | 5.7875 | 7.14E-09 | 3.58E-07 |
| C3orf18 | 1819.3663 | 0.4088 | 0.1126 | 3.6312 | 0.000282055 | 0.002906896 |
| MIS18A | 665.2313 | 0.4087 | 0.1336 | 3.0595 | 0.00221734 | 0.014953948 |
| LOC100133091 | 1741.1718 | 0.4084 | 0.1524 | 2.6800 | 0.007362884 | 0.038000119 |
| ZAK | 309.8696 | 0.4080 | 0.1486 | 2.7463 | 0.006027586 | 0.032529182 |
| AIM2 | 2735.2680 | 0.4078 | 0.1333 | 3.0596 | 0.002216398 | 0.014953948 |
| INPP4B | 9417.7224 | 0.4076 | 0.0849 | 4.8004 | 1.58E-06 | 3.88E-05 |
| CERS6 | 1898.3266 | 0.4075 | 0.1135 | 3.5899 | 0.000330813 | 0.00331405 |
| CDK2AP1 | 2559.4903 | 0.4067 | 0.0897 | 4.5358 | 5.74E-06 | 0.000118895 |
| FAM162A | 11708.1051 | 0.4066 | 0.1216 | 3.3430 | 0.000828885 | 0.006870438 |
| DDIT4 | 48148.9399 | 0.4064 | 0.1009 | 4.0266 | 5.66E-05 | 0.000791933 |
| MIIP | 2489.2156 | 0.4062 | 0.1309 | 3.1033 | 0.001913577 | 0.013370288 |
| ALDH18A1 | 5134.4718 | 0.4061 | 0.0833 | 4.8731 | 1.10E-06 | 2.85E-05 |
| GBP5 | 23290.1198 | 0.4052 | 0.1067 | 3.7969 | 0.000146525 | 0.001710775 |
| HUWE1 | 32717.9472 | 0.4051 | 0.1313 | 3.0842 | 0.002041043 | 0.014053193 |
| CD38 | 4093.4625 | 0.4047 | 0.1553 | 2.6051 | 0.009185442 | 0.044821992 |
| CCDC58 | 1274.0561 | 0.4043 | 0.1042 | 3.8810 | 0.00010403 | 0.001297331 |
| TEC | 559.4509 | 0.4030 | 0.1367 | 2.9486 | 0.003192373 | 0.019922041 |
| EHBP1L1 | 12718.4178 | 0.4028 | 0.0889 | 4.5309 | 5.87E-06 | 0.000120735 |
| FAM129B | 2613.2869 | 0.4027 | 0.1011 | 3.9811 | 6.86E-05 | 0.000926318 |
| DNAJB2 | 7575.3587 | 0.4027 | 0.1085 | 3.7115 | 0.000206055 | 0.00226588 |
| SMYD5 | 1233.3628 | 0.4025 | 0.0928 | 4.3386 | 1.43E-05 | 0.000252827 |
| ZNF295 | 2361.4102 | 0.4023 | 0.1107 | 3.6326 | 0.00028062 | 0.002895589 |
| GPR114 | 3774.6845 | 0.4016 | 0.1069 | 3.7561 | 0.000172561 | 0.001958239 |
| TBC1D4 | 4751.6320 | 0.4015 | 0.1084 | 3.7050 | 0.000211418 | 0.002311682 |
| EIF4A3 | 3340.7184 | 0.4015 | 0.0984 | 4.0808 | 4.49E-05 | 0.000653337 |
| GPX4 | 6382.0673 | 0.4015 | 0.1360 | 2.9521 | 0.003156544 | 0.019755502 |
| DDX41 | 5976.7344 | 0.4014 | 0.0895 | 4.4852 | 7.28E-06 | 0.000143014 |
| SLC35F2 | 1491.9256 | 0.4013 | 0.0912 | 4.4026 | 1.07E-05 | 0.000198246 |
| KCTD11 | 2683.1624 | 0.4012 | 0.1125 | 3.5669 | 0.000361251 | 0.003545272 |
| PDE4D | 21925.2298 | 0.4007 | 0.1075 | 3.7286 | 0.000192529 | 0.002146194 |
| PGK1 | 135697.2308 | 0.4004 | 0.0781 | 5.1289 | 2.91E-07 | 9.24E-06 |
| GPR137B | 1956.0567 | 0.4003 | 0.1143 | 3.5020 | 0.000461784 | 0.004322656 |
| TTI2 | 625.8678 | 0.4001 | 0.1257 | 3.1839 | 0.001453006 | 0.010742725 |
| PTPN13 | 1939.7979 | 0.3993 | 0.1512 | 2.6410 | 0.008266463 | 0.041406219 |
| CORO1C | 3830.3710 | 0.3992 | 0.0779 | 5.1241 | 2.99E-07 | 9.38E-06 |
| GUK1 | 14474.3011 | 0.3985 | 0.1399 | 2.8479 | 0.004400667 | 0.025652752 |
| SPEN | 7735.3890 | 0.3979 | 0.1488 | 2.6737 | 0.007502439 | 0.038523216 |
| SLFN13 | 5938.6573 | 0.3975 | 0.0861 | 4.6171 | 3.89E-06 | 8.43E-05 |
| FAM153C | 1497.0781 | 0.3975 | 0.1301 | 3.0564 | 0.002240016 | 0.015093416 |
| EMD | 4316.2640 | 0.3965 | 0.1051 | 3.7713 | 0.000162428 | 0.00185836 |
| ZFPM1 | 969.2296 | 0.3959 | 0.1308 | 3.0264 | 0.002474932 | 0.016295106 |
| LRRC61 | 1362.6236 | 0.3952 | 0.1455 | 2.7169 | 0.006590599 | 0.03498105 |
| SFI1 | 14335.2101 | 0.3950 | 0.0606 | 6.5137 | 7.33E-11 | 5.44E-09 |
| TNFSF10 | 5325.0513 | 0.3942 | 0.1081 | 3.6446 | 0.000267824 | 0.002788285 |
| DOCK3 | 668.7915 | 0.3937 | 0.1046 | 3.7647 | 0.000166729 | 0.001897751 |
| TTF2 | 2348.4367 | 0.3933 | 0.1256 | 3.1317 | 0.001737751 | 0.012364517 |
| NDFIP1 | 10239.1275 | 0.3932 | 0.0857 | 4.5872 | 4.49E-06 | 9.60E-05 |
| ABLIM2 | 2266.5002 | 0.3928 | 0.1124 | 3.4963 | 0.000471823 | 0.004390573 |
| NCOA3 | 10688.1813 | 0.3922 | 0.1049 | 3.7401 | 0.000183961 | 0.002071944 |
| ARG2 | 6445.3980 | 0.3915 | 0.1205 | 3.2499 | 0.001154378 | 0.008936682 |
| MAPK6 | 5014.6263 | 0.3913 | 0.1055 | 3.7089 | 0.000208152 | 0.002280027 |
| RSRC2 | 10435.6044 | 0.3912 | 0.0860 | 4.5492 | 5.38E-06 | 0.000112662 |
| SAMSN1 | 9602.4427 | 0.3910 | 0.1095 | 3.5699 | 0.000357148 | 0.003515015 |
| GABPB1 | 2564.1847 | 0.3909 | 0.0762 | 5.1330 | 2.85E-07 | 9.08E-06 |
| WDR89 | 1161.9010 | 0.3906 | 0.1110 | 3.5200 | 0.000431496 | 0.004097451 |
| FAM83D | 926.7433 | 0.3900 | 0.1354 | 2.8812 | 0.003961597 | 0.023694453 |
| KIAA1715 | 4850.7157 | 0.3900 | 0.1026 | 3.8017 | 0.000143699 | 0.001687684 |
| UBE2C | 2780.2628 | 0.3898 | 0.1277 | 3.0532 | 0.002264426 | 0.015223989 |
| MDM2 | 19774.4796 | 0.3897 | 0.0943 | 4.1316 | 3.60E-05 | 0.000542874 |
| PLXNB2 | 3903.2557 | 0.3897 | 0.1286 | 3.0302 | 0.002443545 | 0.01616951 |
| PHPT1 | 4699.8160 | 0.3886 | 0.1286 | 3.0213 | 0.002516917 | 0.016503423 |
| AEN | 2834.2619 | 0.3886 | 0.0987 | 3.9353 | 8.31E-05 | 0.001078899 |
| FYCO1 | 5680.8350 | 0.3884 | 0.1146 | 3.3887 | 0.000702147 | 0.006028021 |
| PTGER2 | 3738.0067 | 0.3882 | 0.0954 | 4.0682 | 4.74E-05 | 0.000683179 |
| HLA-F | 9432.6453 | 0.3877 | 0.1128 | 3.4362 | 0.000589967 | 0.005232345 |
| GRAMD3 | 719.9933 | 0.3874 | 0.1087 | 3.5632 | 0.000366293 | 0.003581711 |
| C2orf47 | 898.3179 | 0.3874 | 0.1059 | 3.6587 | 0.000253501 | 0.002659304 |
| MRC2 | 1530.2354 | 0.3872 | 0.1189 | 3.2563 | 0.001128581 | 0.008781912 |
| PSMD12 | 2054.2983 | 0.3871 | 0.0983 | 3.9389 | 8.18E-05 | 0.001070933 |
| NCKIPSD | 6624.4363 | 0.3869 | 0.0723 | 5.3535 | 8.63E-08 | 3.23E-06 |
| ERI1 | 4497.1683 | 0.3866 | 0.0705 | 5.4824 | 4.20E-08 | 1.73E-06 |
| DNAJA1 | 5723.6914 | 0.3865 | 0.0805 | 4.8043 | 1.55E-06 | 3.82E-05 |
| KIF3A | 3096.1060 | 0.3860 | 0.1102 | 3.5030 | 0.000460094 | 0.00431217 |
| NOD2 | 3786.3459 | 0.3854 | 0.1020 | 3.7769 | 0.00015882 | 0.001826972 |
| YES1 | 1339.4816 | 0.3850 | 0.1424 | 2.7033 | 0.006865859 | 0.036049932 |
| PMS2CL | 2265.5716 | 0.3850 | 0.1242 | 3.0982 | 0.00194672 | 0.01352597 |
| SHROOM1 | 593.6055 | 0.3847 | 0.1222 | 3.1484 | 0.001641549 | 0.011830258 |
| SURF4 | 23504.0625 | 0.3840 | 0.0565 | 6.7986 | 1.06E-11 | 9.54E-10 |
| CDK2 | 3488.5801 | 0.3837 | 0.0973 | 3.9444 | 8.00E-05 | 0.001054071 |
| MFSD10 | 15191.7422 | 0.3834 | 0.0969 | 3.9563 | 7.61E-05 | 0.001013672 |
| ABHD10 | 3343.3624 | 0.3828 | 0.0899 | 4.2579 | 2.06E-05 | 0.0003419 |
| SLC36A4 | 1256.7129 | 0.3827 | 0.1331 | 2.8751 | 0.004038399 | 0.023975222 |
| ZC3H3 | 3639.2035 | 0.3826 | 0.0765 | 5.0030 | 5.65E-07 | 1.63E-05 |
| YWHAG | 10398.4988 | 0.3823 | 0.0713 | 5.3611 | 8.27E-08 | 3.12E-06 |
| PLAGL2 | 3539.3434 | 0.3818 | 0.1190 | 3.2088 | 0.00133276 | 0.010025188 |
| MAD2L1 | 4358.9681 | 0.3817 | 0.1075 | 3.5505 | 0.000384566 | 0.003721899 |
| OTUD6B | 1372.3830 | 0.3814 | 0.1191 | 3.2032 | 0.001358887 | 0.010157363 |
| KIAA1958 | 2296.3855 | 0.3814 | 0.1068 | 3.5719 | 0.000354372 | 0.003499079 |
| FOSL2 | 12860.6999 | 0.3807 | 0.1088 | 3.4999 | 0.000465418 | 0.004347778 |
| DDHD1 | 15099.7507 | 0.3806 | 0.1120 | 3.3975 | 0.000680005 | 0.005867904 |
| ITPRIPL1 | 1257.2541 | 0.3799 | 0.1422 | 2.6717 | 0.007545904 | 0.038706982 |
| SLC25A37 | 956.9442 | 0.3798 | 0.1119 | 3.3940 | 0.00068887 | 0.005924126 |
| ING2 | 1403.0012 | 0.3796 | 0.1329 | 2.8557 | 0.004294402 | 0.025198376 |
| LRP5 | 859.9079 | 0.3793 | 0.1457 | 2.6035 | 0.009227598 | 0.044998672 |
| MPZL1 | 1589.8639 | 0.3786 | 0.0947 | 3.9995 | 6.35E-05 | 0.000871475 |
| EAF1 | 4385.1316 | 0.3784 | 0.0945 | 4.0045 | 6.21E-05 | 0.000857773 |
| PPIL4 | 3367.3801 | 0.3781 | 0.0896 | 4.2183 | 2.46E-05 | 0.000393584 |
| CIC | 9261.2691 | 0.3779 | 0.1249 | 3.0257 | 0.002480923 | 0.01631692 |
| BCOR | 7181.6572 | 0.3777 | 0.1193 | 3.1662 | 0.001544696 | 0.011266448 |
| MOB1A | 7504.4996 | 0.3771 | 0.1137 | 3.3179 | 0.00090701 | 0.007397652 |
| BRIX1 | 1588.5370 | 0.3771 | 0.0932 | 4.0460 | 5.21E-05 | 0.000739332 |
| TMEM237 | 1073.4921 | 0.3768 | 0.1116 | 3.3768 | 0.000733338 | 0.006242659 |
| COPG2 | 804.4130 | 0.3767 | 0.1322 | 2.8498 | 0.004374364 | 0.025523271 |
| ARHGAP10 | 2880.9453 | 0.3764 | 0.0911 | 4.1300 | 3.63E-05 | 0.000545466 |
| FAM126B | 4932.6329 | 0.3762 | 0.0957 | 3.9300 | 8.49E-05 | 0.001098188 |
| KIF3B | 3965.8130 | 0.3760 | 0.0851 | 4.4167 | 1.00E-05 | 0.000187428 |
| MAN2A2 | 9862.6924 | 0.3757 | 0.1145 | 3.2816 | 0.001032358 | 0.008206245 |
| PSMD11 | 4868.1905 | 0.3757 | 0.0746 | 5.0372 | 4.72E-07 | 1.39E-05 |
| C11orf30 | 3577.5828 | 0.3754 | 0.0997 | 3.7665 | 0.000165564 | 0.00188859 |
| C14orf118 | 2140.4226 | 0.3752 | 0.1367 | 2.7442 | 0.00606668 | 0.032693506 |
| KLC2 | 1971.2152 | 0.3749 | 0.1043 | 3.5952 | 0.000324083 | 0.003259574 |
| FKBP4 | 3456.4679 | 0.3748 | 0.0787 | 4.7648 | 1.89E-06 | 4.50E-05 |
| SEC24D | 9614.5304 | 0.3746 | 0.1416 | 2.6459 | 0.008146923 | 0.040929428 |
| ARID3A | 2929.7106 | 0.3731 | 0.0957 | 3.8976 | 9.71E-05 | 0.001228728 |
| GOLGA4 | 7075.8724 | 0.3729 | 0.1044 | 3.5713 | 0.000355264 | 0.003503917 |
| TTC21B | 2235.5397 | 0.3728 | 0.1208 | 3.0863 | 0.002026672 | 0.013977466 |
| ETV6 | 7777.6404 | 0.3722 | 0.0569 | 6.5415 | 6.09E-11 | 4.63E-09 |
| NFKB1 | 8787.7935 | 0.3721 | 0.0798 | 4.6630 | 3.12E-06 | 6.94E-05 |
| MARS | 6288.3709 | 0.3719 | 0.0675 | 5.5121 | 3.55E-08 | 1.49E-06 |
| FADS3 | 853.1554 | 0.3718 | 0.1028 | 3.6176 | 0.000297342 | 0.003039115 |
| PSMC2 | 1412.1922 | 0.3712 | 0.1318 | 2.8161 | 0.004860666 | 0.027861803 |
| LARS2 | 1069.5637 | 0.3691 | 0.0991 | 3.7258 | 0.000194661 | 0.002166766 |
| GBP2 | 36169.6551 | 0.3687 | 0.0928 | 3.9729 | 7.10E-05 | 0.000954 |
| PPIF | 5048.7158 | 0.3685 | 0.0790 | 4.6633 | 3.11E-06 | 6.94E-05 |
| HSD17B7P2 | 1941.5880 | 0.3678 | 0.1041 | 3.5327 | 0.000411329 | 0.003943079 |
| SFXN1 | 17495.4766 | 0.3677 | 0.0816 | 4.5075 | 6.56E-06 | 0.000132494 |
| HNRPLL | 6929.5913 | 0.3675 | 0.0618 | 5.9465 | 2.74E-09 | 1.47E-07 |
| GNPTAB | 11142.5762 | 0.3672 | 0.1103 | 3.3289 | 0.000871848 | 0.007167631 |
| NUDCD1 | 1324.1124 | 0.3670 | 0.1291 | 2.8427 | 0.004473453 | 0.025996897 |
| SLC16A1 | 4987.2093 | 0.3666 | 0.0889 | 4.1257 | 3.70E-05 | 0.000553667 |
| NACC1 | 4631.6855 | 0.3666 | 0.1179 | 3.1099 | 0.001871471 | 0.013106359 |
| PAK4 | 790.4659 | 0.3665 | 0.1136 | 3.2264 | 0.001253766 | 0.009564122 |
| DUSP3 | 2049.5927 | 0.3665 | 0.1145 | 3.2019 | 0.001365199 | 0.010198205 |
| PIK3C3 | 4315.8241 | 0.3664 | 0.0933 | 3.9266 | 8.61E-05 | 0.001110972 |
| SNRPA1 | 6607.2905 | 0.3664 | 0.1121 | 3.2694 | 0.001077895 | 0.008474695 |
| NOP58 | 8912.3116 | 0.3661 | 0.0929 | 3.9413 | 8.11E-05 | 0.001064305 |
| SBF2 | 1183.1128 | 0.3660 | 0.1170 | 3.1274 | 0.001763341 | 0.012499557 |
| XPO5 | 3314.5067 | 0.3658 | 0.1244 | 2.9394 | 0.003288038 | 0.020389942 |
| RAB5A | 2609.1098 | 0.3655 | 0.1265 | 2.8891 | 0.003862984 | 0.023151885 |
| C7orf30 | 1329.7594 | 0.3651 | 0.1203 | 3.0349 | 0.002406483 | 0.015983653 |
| ENOPH1 | 3675.1345 | 0.3651 | 0.0619 | 5.8965 | 3.71E-09 | 1.95E-07 |
| ELAVL1 | 6353.8778 | 0.3647 | 0.0814 | 4.4800 | 7.46E-06 | 0.000146055 |
| CFLAR | 21257.4054 | 0.3645 | 0.0926 | 3.9357 | 8.30E-05 | 0.001078899 |
| MCTP2 | 5133.4600 | 0.3638 | 0.1287 | 2.8270 | 0.004698763 | 0.02711873 |
| ZNF395 | 23382.3290 | 0.3636 | 0.1172 | 3.1025 | 0.001919203 | 0.013391041 |
| BCKDK | 2843.3163 | 0.3631 | 0.1107 | 3.2800 | 0.001037939 | 0.008241942 |
| PFDN2 | 2367.4732 | 0.3628 | 0.1297 | 2.7970 | 0.005157522 | 0.029122001 |
| TAF1D | 10273.5421 | 0.3627 | 0.1006 | 3.6055 | 0.00031155 | 0.0031576 |
| MAP2K3 | 5328.4120 | 0.3626 | 0.0668 | 5.4291 | 5.66E-08 | 2.24E-06 |
| PDIA6 | 12174.9362 | 0.3622 | 0.0984 | 3.6791 | 0.000234035 | 0.002496653 |
| FAM105B | 7409.3002 | 0.3620 | 0.0832 | 4.3514 | 1.35E-05 | 0.000241068 |
| CD7 | 11000.3938 | 0.3616 | 0.1167 | 3.0991 | 0.001940875 | 0.013511096 |
| TFG | 9976.4445 | 0.3614 | 0.0823 | 4.3896 | 1.14E-05 | 0.000207427 |
| SGOL2 | 3505.2044 | 0.3612 | 0.1073 | 3.3676 | 0.000758186 | 0.006414477 |
| RBM15 | 5937.8891 | 0.3609 | 0.0848 | 4.2540 | 2.10E-05 | 0.00034599 |
| IVNS1ABP | 21255.3953 | 0.3609 | 0.0806 | 4.4801 | 7.46E-06 | 0.000146055 |
| NAA15 | 2652.9921 | 0.3600 | 0.0882 | 4.0799 | 4.51E-05 | 0.000654635 |
| FMNL3 | 2819.2288 | 0.3600 | 0.1328 | 2.7117 | 0.00669334 | 0.035414535 |
| TYW5 | 3062.7113 | 0.3599 | 0.1169 | 3.0785 | 0.002080256 | 0.014264751 |
| DTYMK | 1567.4181 | 0.3594 | 0.0918 | 3.9167 | 8.98E-05 | 0.001150762 |
| ACSL1 | 2138.0380 | 0.3593 | 0.1088 | 3.3023 | 0.000959019 | 0.007732989 |
| RFC5 | 2225.7477 | 0.3592 | 0.0837 | 4.2911 | 1.78E-05 | 0.000303904 |
| SUV420H2 | 1339.9121 | 0.3590 | 0.1006 | 3.5687 | 0.00035869 | 0.003525606 |
| SQRDL | 10649.0728 | 0.3583 | 0.1044 | 3.4330 | 0.000596999 | 0.005278089 |
| TMEM39A | 4909.0818 | 0.3583 | 0.1211 | 2.9579 | 0.003097316 | 0.019473444 |
| FAM98B | 1504.3241 | 0.3583 | 0.1027 | 3.4895 | 0.000484012 | 0.004486302 |
| AUH | 10377.8021 | 0.3583 | 0.1139 | 3.1443 | 0.001665077 | 0.011948583 |
| CTDP1 | 4018.2855 | 0.3579 | 0.0932 | 3.8397 | 0.000123174 | 0.001496583 |
| MLEC | 13190.5850 | 0.3572 | 0.0934 | 3.8256 | 0.000130455 | 0.001575958 |
| ADRBK2 | 6417.2862 | 0.3570 | 0.1230 | 2.9026 | 0.003700739 | 0.022365591 |
| STK35 | 3113.0053 | 0.3568 | 0.0964 | 3.7016 | 0.000214218 | 0.002336994 |
| GAD1 | 1164.3131 | 0.3566 | 0.1153 | 3.0921 | 0.001987645 | 0.013773296 |
| HMGCS1 | 8341.7602 | 0.3566 | 0.0659 | 5.4095 | 6.32E-08 | 2.45E-06 |
| ATAD5 | 3422.2259 | 0.3566 | 0.0955 | 3.7328 | 0.000189336 | 0.002119976 |
| ADAR | 26371.6319 | 0.3553 | 0.0891 | 3.9860 | 6.72E-05 | 0.00091147 |
| TANK | 9908.7214 | 0.3550 | 0.1087 | 3.2665 | 0.001088936 | 0.008532375 |
| GPR35 | 1669.4377 | 0.3549 | 0.1208 | 2.9374 | 0.003310278 | 0.020480398 |
| SPATA5L1 | 1171.0701 | 0.3548 | 0.0824 | 4.3040 | 1.68E-05 | 0.000289624 |
| RBBP8 | 3442.4713 | 0.3536 | 0.0869 | 4.0710 | 4.68E-05 | 0.000676965 |
| STAMBP | 7212.4110 | 0.3536 | 0.0823 | 4.2963 | 1.74E-05 | 0.000297919 |
| LOC613037 | 498.1949 | 0.3536 | 0.1207 | 2.9304 | 0.003385756 | 0.020836586 |
| CEP152 | 5823.6884 | 0.3532 | 0.1218 | 2.8989 | 0.003744476 | 0.022602829 |
| BNIP3 | 11928.1062 | 0.3529 | 0.1072 | 3.2921 | 0.00099442 | 0.007956601 |
| BOD1 | 1223.5110 | 0.3521 | 0.1203 | 2.9261 | 0.003432161 | 0.021053648 |
| TFE3 | 5148.0201 | 0.3517 | 0.0694 | 5.0717 | 3.94E-07 | 1.19E-05 |
| CNTROB | 7021.0168 | 0.3513 | 0.0788 | 4.4589 | 8.24E-06 | 0.000158564 |
| UBASH3B | 2110.8212 | 0.3511 | 0.1113 | 3.1543 | 0.001608563 | 0.011648027 |
| TIFA | 4390.2872 | 0.3510 | 0.0995 | 3.5276 | 0.000419303 | 0.004001765 |
| C17orf53 | 729.3661 | 0.3508 | 0.1160 | 3.0246 | 0.002489771 | 0.016360891 |
| SIPA1L1 | 6666.7150 | 0.3507 | 0.0869 | 4.0372 | 5.41E-05 | 0.000762448 |
| PPAT | 2195.5356 | 0.3506 | 0.0989 | 3.5455 | 0.000391919 | 0.003785794 |
| AFF4 | 8716.2640 | 0.3502 | 0.1011 | 3.4646 | 0.000530979 | 0.00481253 |
| MBD2 | 12202.4080 | 0.3496 | 0.0625 | 5.5926 | 2.24E-08 | 1.01E-06 |
| PTMS | 4603.2012 | 0.3491 | 0.1322 | 2.6411 | 0.008263169 | 0.041403429 |
| ADAM9 | 2142.0650 | 0.3491 | 0.1127 | 3.0981 | 0.001947482 | 0.01352597 |
| PRC1 | 4501.1602 | 0.3486 | 0.0975 | 3.5771 | 0.00034737 | 0.003447947 |
| FAM98A | 1579.6263 | 0.3484 | 0.0882 | 3.9480 | 7.88E-05 | 0.001040508 |
| PFKFB4 | 22588.9844 | 0.3479 | 0.1137 | 3.0596 | 0.002216028 | 0.014953948 |
| SART3 | 5552.3963 | 0.3478 | 0.0751 | 4.6324 | 3.62E-06 | 7.87E-05 |
| NCAPG2 | 5726.7386 | 0.3471 | 0.1031 | 3.3657 | 0.000763389 | 0.006440479 |
| KIF2C | 4130.5584 | 0.3469 | 0.1148 | 3.0209 | 0.002520456 | 0.016506688 |
| CSTF2 | 1441.7097 | 0.3468 | 0.0980 | 3.5399 | 0.00040025 | 0.003846618 |
| KIAA1586 | 1191.1830 | 0.3466 | 0.1110 | 3.1221 | 0.00179575 | 0.012677979 |
| MSI2 | 6722.0740 | 0.3464 | 0.1025 | 3.3813 | 0.000721472 | 0.006162458 |
| ERF | 2207.2932 | 0.3452 | 0.1162 | 2.9712 | 0.002966346 | 0.018877539 |
| ABCB7 | 3465.7304 | 0.3452 | 0.0830 | 4.1568 | 3.23E-05 | 0.000495183 |
| TMEM106A | 499.4108 | 0.3451 | 0.1249 | 2.7640 | 0.005710401 | 0.031290393 |
| STIP1 | 7239.9047 | 0.3451 | 0.0703 | 4.9058 | 9.30E-07 | 2.49E-05 |
| CARS | 9193.4633 | 0.3448 | 0.1139 | 3.0266 | 0.002473363 | 0.016295106 |
| LAMB3 | 1101.7304 | 0.3447 | 0.1293 | 2.6656 | 0.007685036 | 0.039155114 |
| CD226 | 8074.1238 | 0.3446 | 0.1176 | 2.9311 | 0.003377545 | 0.020811457 |
| FUT8 | 5878.0233 | 0.3443 | 0.0788 | 4.3689 | 1.25E-05 | 0.000224108 |
| PEA15 | 2551.2886 | 0.3440 | 0.1140 | 3.0178 | 0.002546121 | 0.016622864 |
| PSMC3IP | 377.2337 | 0.3436 | 0.1333 | 2.5778 | 0.009943113 | 0.047643167 |
| IER5 | 4204.0794 | 0.3434 | 0.1131 | 3.0369 | 0.002390268 | 0.01592144 |
| RAI1 | 2076.1340 | 0.3429 | 0.1267 | 2.7065 | 0.006798937 | 0.035860361 |
| ACP5 | 2209.2549 | 0.3428 | 0.1082 | 3.1672 | 0.001539098 | 0.011231038 |
| C11orf24 | 821.0603 | 0.3428 | 0.1041 | 3.2925 | 0.000993155 | 0.007953128 |
| DKFZp761E198 | 1789.9698 | 0.3416 | 0.1308 | 2.6126 | 0.008985012 | 0.044015315 |
| TTLL4 | 1144.2433 | 0.3412 | 0.0912 | 3.7399 | 0.000184088 | 0.002071944 |
| UFM1 | 9806.3160 | 0.3412 | 0.0917 | 3.7223 | 0.000197454 | 0.002189805 |
| PSMC6 | 5953.6954 | 0.3406 | 0.1039 | 3.2799 | 0.001038546 | 0.008242437 |
| EIF2AK3 | 4168.2932 | 0.3405 | 0.0940 | 3.6222 | 0.000292127 | 0.002993907 |
| DLEU2 | 2176.6171 | 0.3402 | 0.0919 | 3.7016 | 0.00021428 | 0.002336994 |
| ARL3 | 2535.9982 | 0.3392 | 0.0861 | 3.9372 | 8.24E-05 | 0.001075654 |
| AASDHPPT | 1747.8676 | 0.3389 | 0.1267 | 2.6757 | 0.007457956 | 0.038386018 |
| CDK1 | 4804.8026 | 0.3389 | 0.1117 | 3.0342 | 0.002411991 | 0.016009735 |
| MBIP | 2643.0055 | 0.3388 | 0.0870 | 3.8920 | 9.94E-05 | 0.001251244 |
| SERPINB8 | 1257.0417 | 0.3387 | 0.1114 | 3.0389 | 0.002374175 | 0.015835159 |
| PSMD5 | 5387.0769 | 0.3385 | 0.0918 | 3.6880 | 0.000226051 | 0.002438997 |
| SMC2 | 6189.3548 | 0.3381 | 0.1150 | 2.9390 | 0.00329235 | 0.020394504 |
| SEH1L | 2398.1801 | 0.3375 | 0.0747 | 4.5175 | 6.26E-06 | 0.000126881 |
| DNMT3A | 3376.5304 | 0.3365 | 0.0907 | 3.7091 | 0.000208034 | 0.002280027 |
| AGPAT5 | 4336.6610 | 0.3364 | 0.0885 | 3.8035 | 0.000142684 | 0.001678371 |
| IRAK2 | 1126.5435 | 0.3363 | 0.0949 | 3.5441 | 0.000393938 | 0.003798027 |
| FAM133B | 3437.1434 | 0.3361 | 0.1210 | 2.7781 | 0.005467355 | 0.030338773 |
| NFE2L1 | 10004.4351 | 0.3361 | 0.0896 | 3.7494 | 0.00017723 | 0.002005209 |
| CHORDC1 | 4315.0519 | 0.3353 | 0.1090 | 3.0767 | 0.002093142 | 0.014342782 |
| GFI1 | 10458.4456 | 0.3350 | 0.1099 | 3.0485 | 0.002300086 | 0.015428867 |
| WEE1 | 2677.8246 | 0.3346 | 0.0853 | 3.9209 | 8.82E-05 | 0.001133611 |
| ACBD3 | 4297.8673 | 0.3338 | 0.0822 | 4.0613 | 4.88E-05 | 0.000700773 |
| SCAPER | 1863.9999 | 0.3337 | 0.0803 | 4.1545 | 3.26E-05 | 0.000499574 |
| NEDD9 | 3640.8233 | 0.3335 | 0.0995 | 3.3526 | 0.000800437 | 0.006693321 |
| POLD1 | 3000.2126 | 0.3334 | 0.1041 | 3.2032 | 0.00135906 | 0.010157363 |
| PTP4A1 | 11532.8311 | 0.3333 | 0.0857 | 3.8907 | 1.00E-04 | 0.001253964 |
| PPIL3 | 5899.5509 | 0.3331 | 0.0815 | 4.0877 | 4.36E-05 | 0.000636714 |
| POLA2 | 1678.3730 | 0.3330 | 0.1083 | 3.0739 | 0.002112972 | 0.01444326 |
| ZNF627 | 574.1132 | 0.3329 | 0.1053 | 3.1627 | 0.001563267 | 0.011369006 |
| HPS5 | 2345.6516 | 0.3319 | 0.0932 | 3.5622 | 0.000367695 | 0.003588464 |
| CHD1L | 6830.1999 | 0.3312 | 0.0912 | 3.6312 | 0.0002821 | 0.002906896 |
| RAB1A | 9669.9794 | 0.3310 | 0.1052 | 3.1450 | 0.001660985 | 0.011924881 |
| MAP3K14 | 2125.3388 | 0.3310 | 0.1043 | 3.1741 | 0.001503073 | 0.011037374 |
| SMG9 | 3935.1517 | 0.3310 | 0.0977 | 3.3859 | 0.000709534 | 0.006074202 |
| CPD | 4115.8475 | 0.3308 | 0.0929 | 3.5612 | 0.000369206 | 0.003596247 |
| TGIF1 | 3040.1188 | 0.3307 | 0.0950 | 3.4800 | 0.000501494 | 0.004603215 |
| CAB39L | 1806.5142 | 0.3307 | 0.0875 | 3.7781 | 0.000158017 | 0.00181912 |
| ZUFSP | 2025.3166 | 0.3305 | 0.1101 | 3.0026 | 0.002676588 | 0.017302881 |
| MED20 | 1418.4240 | 0.3305 | 0.1105 | 2.9919 | 0.00277286 | 0.017865804 |
| KLHL5 | 3052.3550 | 0.3305 | 0.0786 | 4.2038 | 2.63E-05 | 0.000413795 |
| TRIM25 | 13140.4964 | 0.3303 | 0.0844 | 3.9125 | 9.13E-05 | 0.001168021 |
| ACOX3 | 2877.9413 | 0.3298 | 0.1000 | 3.2967 | 0.000978215 | 0.007866808 |
| POM121C | 11718.4995 | 0.3297 | 0.0826 | 3.9936 | 6.51E-05 | 0.000888324 |
| FNBP1 | 23532.5061 | 0.3285 | 0.1109 | 2.9628 | 0.003048452 | 0.019230166 |
| ARID3B | 4809.6975 | 0.3282 | 0.0895 | 3.6657 | 0.000246688 | 0.002595025 |
| TAP1 | 21228.8622 | 0.3281 | 0.0661 | 4.9621 | 6.97E-07 | 1.96E-05 |
| SLC2A1 | 20417.3328 | 0.3280 | 0.0830 | 3.9538 | 7.69E-05 | 0.001021404 |
| DSCC1 | 775.3413 | 0.3278 | 0.1248 | 2.6262 | 0.008635231 | 0.042771821 |
| MCM2 | 5360.4471 | 0.3275 | 0.1161 | 2.8197 | 0.004806618 | 0.027625268 |
| C13orf15 | 5720.8020 | 0.3270 | 0.1110 | 2.9469 | 0.003209721 | 0.020005545 |
| SRP54 | 9107.1337 | 0.3270 | 0.0986 | 3.3182 | 0.000906039 | 0.007396464 |
| TSPAN17 | 2583.8085 | 0.3266 | 0.1114 | 2.9322 | 0.003366081 | 0.020749269 |
| PLXNA3 | 8875.6574 | 0.3260 | 0.1235 | 2.6407 | 0.008274471 | 0.041420957 |
| PHGDH | 3602.4306 | 0.3260 | 0.1263 | 2.5804 | 0.009868181 | 0.047344111 |
| TAB2 | 19532.6683 | 0.3257 | 0.0775 | 4.2044 | 2.62E-05 | 0.000413795 |
| CD70 | 4301.0578 | 0.3255 | 0.1132 | 2.8763 | 0.004023383 | 0.023914229 |
| PNRC1 | 12391.0764 | 0.3252 | 0.0782 | 4.1582 | 3.21E-05 | 0.000492554 |
| PSMB5 | 2009.2642 | 0.3250 | 0.0944 | 3.4424 | 0.000576588 | 0.005142719 |
| SMNDC1 | 5364.9867 | 0.3246 | 0.0850 | 3.8204 | 0.000133248 | 0.001604814 |
| DHX58 | 1999.1834 | 0.3242 | 0.1192 | 2.7188 | 0.006551486 | 0.034846811 |
| FAM213B | 1870.7318 | 0.3241 | 0.1048 | 3.0939 | 0.001975746 | 0.013697116 |
| ING3 | 3824.4451 | 0.3241 | 0.0919 | 3.5279 | 0.000418899 | 0.004000429 |
| MAP3K9 | 648.4938 | 0.3240 | 0.1198 | 2.7053 | 0.006824218 | 0.035926522 |
| PRSS53 | 2085.2822 | 0.3238 | 0.0992 | 3.2653 | 0.001093629 | 0.008558371 |
| SLC15A4 | 3887.7654 | 0.3236 | 0.0768 | 4.2160 | 2.49E-05 | 0.000396446 |
| C1orf144 | 2264.1666 | 0.3234 | 0.0988 | 3.2723 | 0.001066614 | 0.008403476 |
| ZFYVE20 | 2857.9585 | 0.3233 | 0.0838 | 3.8594 | 0.000113655 | 0.001397769 |
| PPP1R16B | 17026.5815 | 0.3224 | 0.0852 | 3.7853 | 0.000153546 | 0.001774401 |
| DCTN4 | 3702.8447 | 0.3224 | 0.0872 | 3.6981 | 0.000217185 | 0.002360173 |
| DEPDC1 | 1501.1548 | 0.3223 | 0.1152 | 2.7980 | 0.005142278 | 0.029079344 |
| MASTL | 3311.8537 | 0.3222 | 0.0876 | 3.6778 | 0.000235246 | 0.002506032 |
| RBM8A | 3380.8611 | 0.3221 | 0.1002 | 3.2150 | 0.001304242 | 0.009864631 |
| MYB | 6077.6335 | 0.3208 | 0.0965 | 3.3246 | 0.000885509 | 0.007256279 |
| DCP1A | 3949.4543 | 0.3200 | 0.0926 | 3.4556 | 0.000549093 | 0.00495297 |
| SATB1 | 16837.8765 | 0.3198 | 0.1077 | 2.9682 | 0.002995307 | 0.01902489 |
| NCAPH | 6234.4095 | 0.3197 | 0.0926 | 3.4525 | 0.000555498 | 0.005001793 |
| LRRC42 | 3533.4221 | 0.3195 | 0.0930 | 3.4347 | 0.000593317 | 0.005251667 |
| GNA12 | 2168.8768 | 0.3194 | 0.1212 | 2.6356 | 0.008397851 | 0.041856438 |
| RFC3 | 2369.2304 | 0.3194 | 0.1051 | 3.0383 | 0.002379003 | 0.015853384 |
| SERAC1 | 1223.0134 | 0.3193 | 0.1073 | 2.9770 | 0.002911257 | 0.018597377 |
| ZNF438 | 1306.9080 | 0.3192 | 0.0996 | 3.2047 | 0.001352078 | 0.010131964 |
| NARS2 | 1125.1077 | 0.3191 | 0.0846 | 3.7711 | 0.000162542 | 0.00185836 |
| GEMIN4 | 1729.3673 | 0.3189 | 0.1008 | 3.1652 | 0.001549513 | 0.011286194 |
| CD58 | 10044.2053 | 0.3186 | 0.1004 | 3.1738 | 0.001504364 | 0.011041489 |
| F11R | 3904.9329 | 0.3186 | 0.1233 | 2.5840 | 0.009765211 | 0.047014116 |
| CCNB1 | 2916.9288 | 0.3184 | 0.1189 | 2.6771 | 0.007425891 | 0.038286112 |
| PIP5K1A | 5068.3306 | 0.3182 | 0.0860 | 3.7009 | 0.00021481 | 0.002341083 |
| NAT10 | 6434.6637 | 0.3180 | 0.0718 | 4.4257 | 9.61E-06 | 0.000181517 |
| ATP7A | 5918.2446 | 0.3177 | 0.1189 | 2.6708 | 0.007566062 | 0.038784078 |
| AKAP10 | 4382.2022 | 0.3177 | 0.1145 | 2.7735 | 0.005546087 | 0.030630033 |
| PTPN1 | 8089.3248 | 0.3176 | 0.0805 | 3.9444 | 8.00E-05 | 0.001054071 |
| GAB2 | 5374.1832 | 0.3170 | 0.1022 | 3.1028 | 0.001917017 | 0.013381966 |
| SEC23A | 6424.7901 | 0.3165 | 0.0965 | 3.2810 | 0.001034264 | 0.008217073 |
| LRP5L | 2323.1282 | 0.3165 | 0.1164 | 2.7181 | 0.006565196 | 0.034895195 |
| TET2 | 5826.1109 | 0.3162 | 0.1204 | 2.6264 | 0.008630242 | 0.042761112 |
| GSTCD | 925.7108 | 0.3159 | 0.1085 | 2.9119 | 0.003591871 | 0.021864884 |
| IKBIP | 2121.7211 | 0.3151 | 0.1178 | 2.6745 | 0.007485064 | 0.038486255 |
| HMGCR | 5453.6433 | 0.3150 | 0.0740 | 4.2546 | 2.09E-05 | 0.000345421 |
| DHRS13 | 1055.4729 | 0.3150 | 0.0980 | 3.2138 | 0.001310023 | 0.009898457 |
| RNF24 | 2234.8041 | 0.3147 | 0.1163 | 2.7063 | 0.006804449 | 0.035864424 |
| TIMM21 | 1092.7521 | 0.3133 | 0.1014 | 3.0887 | 0.002010092 | 0.013897011 |
| IL2RG | 63004.0042 | 0.3132 | 0.0687 | 4.5560 | 5.21E-06 | 0.000109392 |
| ZC3H7A | 9037.8262 | 0.3119 | 0.0877 | 3.5556 | 0.000377086 | 0.003658527 |
| SLC39A1 | 5052.7023 | 0.3118 | 0.0731 | 4.2670 | 1.98E-05 | 0.000330384 |
| TDP1 | 3917.2107 | 0.3116 | 0.0816 | 3.8188 | 0.000134093 | 0.001609861 |
| EIF4H | 25776.2018 | 0.3114 | 0.0665 | 4.6858 | 2.79E-06 | 6.33E-05 |
| TARS | 8180.2595 | 0.3113 | 0.0752 | 4.1367 | 3.52E-05 | 0.000534058 |
| INTS10 | 5936.3301 | 0.3113 | 0.0786 | 3.9614 | 7.45E-05 | 0.000993868 |
| ORC6 | 1047.6396 | 0.3113 | 0.1100 | 2.8292 | 0.004666912 | 0.026955471 |
| ANP32E | 13892.9348 | 0.3109 | 0.0734 | 4.2337 | 2.30E-05 | 0.000371229 |
| AARSD1 | 1337.2176 | 0.3108 | 0.1197 | 2.5971 | 0.00940097 | 0.045638148 |
| RPF2 | 1218.5733 | 0.3107 | 0.1167 | 2.6627 | 0.007751702 | 0.039401881 |
| PARP2 | 1980.0753 | 0.3091 | 0.1064 | 2.9066 | 0.003654216 | 0.022146367 |
| ZEB1 | 7799.0680 | 0.3090 | 0.0852 | 3.6286 | 0.00028494 | 0.002930177 |
| CYP4F35P | 665.5970 | 0.3090 | 0.1164 | 2.6541 | 0.007953077 | 0.040144804 |
| ODC1 | 5514.4056 | 0.3087 | 0.0867 | 3.5612 | 0.000369126 | 0.003596247 |
| INTS2 | 2603.8074 | 0.3083 | 0.1088 | 2.8339 | 0.004598372 | 0.02660022 |
| KIF23 | 1999.3687 | 0.3082 | 0.1122 | 2.7454 | 0.006043282 | 0.032602257 |
| SLC25A36 | 13576.3340 | 0.3079 | 0.1095 | 2.8109 | 0.004939865 | 0.028166356 |
| HPRT1 | 4725.7949 | 0.3078 | 0.0992 | 3.1014 | 0.001926244 | 0.013427787 |
| MIER2 | 1313.7638 | 0.3073 | 0.0936 | 3.2816 | 0.001032098 | 0.008206245 |
| VEGFA | 9611.8490 | 0.3072 | 0.0900 | 3.4115 | 0.000646164 | 0.005630488 |
| ADO | 1412.1591 | 0.3070 | 0.1148 | 2.6741 | 0.007493004 | 0.03850091 |
| HSPD1 | 17145.1480 | 0.3068 | 0.0586 | 5.2331 | 1.67E-07 | 5.69E-06 |
| ST3GAL3 | 1193.2134 | 0.3067 | 0.1007 | 3.0449 | 0.00232794 | 0.01560246 |
| CDK7 | 2221.1867 | 0.3067 | 0.0930 | 3.2968 | 0.000977927 | 0.007866808 |
| TXLNA | 8962.1459 | 0.3066 | 0.0601 | 5.1045 | 3.32E-07 | 1.03E-05 |
| ATAD2 | 5602.2646 | 0.3065 | 0.1038 | 2.9515 | 0.003162723 | 0.019785987 |
| RFC1 | 7827.6653 | 0.3056 | 0.0832 | 3.6733 | 0.000239465 | 0.002536684 |
| PJA2 | 14227.6611 | 0.3055 | 0.1130 | 2.7023 | 0.00688538 | 0.036127347 |
| ZC3H18 | 6576.3608 | 0.3052 | 0.0932 | 3.2755 | 0.001054704 | 0.008344411 |
| JMJD1C | 6746.1307 | 0.3051 | 0.1000 | 3.0512 | 0.002279325 | 0.015317351 |
| STIL | 1787.5478 | 0.3046 | 0.0995 | 3.0626 | 0.002194436 | 0.014900797 |
| ANKLE2 | 8761.0312 | 0.3046 | 0.0770 | 3.9578 | 7.56E-05 | 0.001007994 |
| ELK1 | 4180.4359 | 0.3041 | 0.0802 | 3.7919 | 0.000149478 | 0.001735963 |
| PEX26 | 2172.3334 | 0.3038 | 0.1089 | 2.7910 | 0.005253997 | 0.029479677 |
| NOC3L | 1551.3401 | 0.3036 | 0.0934 | 3.2494 | 0.001156359 | 0.008947437 |
| FBRS | 11115.9458 | 0.3031 | 0.0697 | 4.3493 | 1.37E-05 | 0.000242794 |
| ATHL1 | 21077.7099 | 0.3027 | 0.0933 | 3.2434 | 0.001181259 | 0.009102853 |
| GBP4 | 6511.2775 | 0.3024 | 0.1081 | 2.7976 | 0.00514885 | 0.029094754 |
| NAA35 | 2763.5942 | 0.3012 | 0.0907 | 3.3201 | 0.000899829 | 0.00735768 |
| RELA | 8469.5999 | 0.3012 | 0.0601 | 5.0120 | 5.39E-07 | 1.57E-05 |
| GPCPD1 | 7281.8917 | 0.3012 | 0.1109 | 2.7150 | 0.006626585 | 0.035122757 |
| VDAC1 | 22482.3890 | 0.3009 | 0.0597 | 5.0408 | 4.64E-07 | 1.37E-05 |
| PHF3 | 15332.6413 | 0.3008 | 0.1052 | 2.8583 | 0.004259583 | 0.025042641 |
| SPCS2 | 5610.5939 | 0.3008 | 0.0934 | 3.2214 | 0.001275774 | 0.00968011 |
| NUP205 | 5875.1898 | 0.3003 | 0.1022 | 2.9389 | 0.003293912 | 0.02039583 |
| SAMD10 | 2567.2909 | 0.3000 | 0.1025 | 2.9267 | 0.003425986 | 0.021032828 |
| NSFL1C | 2689.6683 | 0.2992 | 0.0941 | 3.1786 | 0.00147978 | 0.010895531 |
| MED8 | 2360.2078 | 0.2992 | 0.1039 | 2.8805 | 0.003970476 | 0.023730302 |
| RRM2B | 7485.8334 | 0.2989 | 0.1158 | 2.5812 | 0.009846693 | 0.047297531 |
| RPAP3 | 3858.3410 | 0.2977 | 0.0813 | 3.6629 | 0.000249401 | 0.002619186 |
| NAA25 | 2864.0980 | 0.2975 | 0.0997 | 2.9849 | 0.002836834 | 0.018214254 |
| SEC13 | 5316.5753 | 0.2974 | 0.0844 | 3.5240 | 0.000425034 | 0.004046244 |
| C12orf23 | 7024.0969 | 0.2962 | 0.0859 | 3.4496 | 0.000561434 | 0.005043235 |
| PRPF39 | 6964.8008 | 0.2961 | 0.0885 | 3.3467 | 0.00081793 | 0.006798253 |
| GGT1 | 3947.7990 | 0.2960 | 0.0763 | 3.8784 | 0.000105149 | 0.001307036 |
| AMPD3 | 4339.9105 | 0.2959 | 0.0899 | 3.2910 | 0.000998457 | 0.007982908 |
| NUDT4 | 2929.4972 | 0.2959 | 0.1067 | 2.7731 | 0.005552754 | 0.030636243 |
| EPT1 | 2383.3380 | 0.2958 | 0.1001 | 2.9552 | 0.003124917 | 0.019611263 |
| KIF20B | 4031.2033 | 0.2956 | 0.1128 | 2.6210 | 0.008766849 | 0.04323969 |
| NDUFB4 | 6820.6336 | 0.2954 | 0.0894 | 3.3058 | 0.000947107 | 0.007657343 |
| ARAP1 | 17979.6948 | 0.2952 | 0.0806 | 3.6628 | 0.000249504 | 0.002619186 |
| LCLAT1 | 1376.3179 | 0.2946 | 0.1023 | 2.8802 | 0.003974273 | 0.023743615 |
| RAPGEF6 | 5596.4007 | 0.2941 | 0.0954 | 3.0834 | 0.002046509 | 0.014071613 |
| MRPL39 | 2807.1520 | 0.2939 | 0.1105 | 2.6598 | 0.007818838 | 0.039650907 |
| ERLIN1 | 1617.5949 | 0.2935 | 0.0990 | 2.9641 | 0.003036072 | 0.019192083 |
| METTL13 | 2546.6498 | 0.2935 | 0.0698 | 4.2064 | 2.59E-05 | 0.000410535 |
| ALG2 | 3631.1722 | 0.2930 | 0.0961 | 3.0483 | 0.00230102 | 0.015428867 |
| GPR180 | 1587.1878 | 0.2925 | 0.1021 | 2.8653 | 0.004166743 | 0.02462122 |
| APOBEC3C | 5873.6749 | 0.2923 | 0.1069 | 2.7342 | 0.006253375 | 0.033568065 |
| TMTC2 | 1370.0932 | 0.2923 | 0.1009 | 2.8957 | 0.003782722 | 0.022788225 |
| B3GNT2 | 1799.5386 | 0.2921 | 0.0965 | 3.0263 | 0.002475451 | 0.016295106 |
| FHOD1 | 5823.3548 | 0.2919 | 0.0954 | 3.0598 | 0.002214744 | 0.014953948 |
| DHX15 | 13326.1651 | 0.2918 | 0.0893 | 3.2685 | 0.001081343 | 0.008492977 |
| YIPF6 | 2754.2037 | 0.2909 | 0.1034 | 2.8141 | 0.004891875 | 0.027987667 |
| LMNA | 10596.1144 | 0.2902 | 0.1035 | 2.8038 | 0.005049972 | 0.028675271 |
| STAT5A | 8987.3520 | 0.2901 | 0.0902 | 3.2140 | 0.001308769 | 0.009893929 |
| CYLD | 20716.1418 | 0.2900 | 0.0743 | 3.9059 | 9.39E-05 | 0.001197544 |
| SMARCC1 | 8919.2629 | 0.2898 | 0.0798 | 3.6331 | 0.000280035 | 0.002891525 |
| LOC150776 | 3476.0245 | 0.2896 | 0.0801 | 3.6146 | 0.00030077 | 0.003072078 |
| PSMD3 | 4911.2038 | 0.2894 | 0.0732 | 3.9514 | 7.77E-05 | 0.001029206 |
| CENPC1 | 4053.4175 | 0.2890 | 0.1113 | 2.5968 | 0.009408612 | 0.045660597 |
| CHEK1 | 2737.6763 | 0.2888 | 0.0978 | 2.9526 | 0.003151203 | 0.019738407 |
| HNRPDL | 19428.0313 | 0.2886 | 0.0809 | 3.5657 | 0.000362854 | 0.003552681 |
| POLR2E | 5865.4723 | 0.2883 | 0.1043 | 2.7634 | 0.005720719 | 0.0312974 |
| FDPS | 7204.7353 | 0.2878 | 0.0999 | 2.8793 | 0.003985064 | 0.023780268 |
| ARMCX3 | 5801.0785 | 0.2875 | 0.0860 | 3.3437 | 0.000826666 | 0.006855798 |
| FAM60A | 10594.8321 | 0.2870 | 0.1037 | 2.7684 | 0.005633083 | 0.030963534 |
| MAPRE2 | 5884.9471 | 0.2869 | 0.0776 | 3.6973 | 0.000217898 | 0.002366216 |
| KIF21A | 2119.1612 | 0.2866 | 0.0996 | 2.8759 | 0.004028401 | 0.023934651 |
| THOC1 | 4264.4269 | 0.2866 | 0.0866 | 3.3105 | 0.000931206 | 0.00754491 |
| STRN4 | 4397.5126 | 0.2865 | 0.0822 | 3.4858 | 0.000490685 | 0.004520459 |
| C4orf46 | 1240.6488 | 0.2864 | 0.0987 | 2.9002 | 0.003729766 | 0.022523024 |
| IWS1 | 4337.4916 | 0.2862 | 0.0879 | 3.2548 | 0.00113463 | 0.008813558 |
| FAM54A | 928.7451 | 0.2856 | 0.1070 | 2.6700 | 0.007584783 | 0.038829681 |
| ATP6V0A2 | 3732.6182 | 0.2854 | 0.0945 | 3.0205 | 0.002523685 | 0.01651916 |
| ABCC1 | 11528.1299 | 0.2850 | 0.0821 | 3.4707 | 0.000519053 | 0.004724258 |
| TMOD3 | 5341.3698 | 0.2850 | 0.0888 | 3.2106 | 0.001324422 | 0.009985458 |
| HCFC1 | 10111.9460 | 0.2848 | 0.0755 | 3.7709 | 0.000162654 | 0.00185836 |
| PTGER4 | 6913.0789 | 0.2845 | 0.0795 | 3.5797 | 0.000344006 | 0.003423535 |
| CDC73 | 5989.6411 | 0.2843 | 0.0913 | 3.1135 | 0.00184908 | 0.012979599 |
| MOB1B | 3569.1389 | 0.2838 | 0.0864 | 3.2832 | 0.001026538 | 0.008181474 |
| STAMBPL1 | 6888.0790 | 0.2838 | 0.0841 | 3.3744 | 0.000739751 | 0.006286633 |
| VAMP1 | 1030.4320 | 0.2831 | 0.1093 | 2.5910 | 0.009570541 | 0.046372063 |
| USP22 | 11708.4805 | 0.2830 | 0.0587 | 4.8255 | 1.40E-06 | 3.49E-05 |
| NFE2L2 | 4853.8044 | 0.2826 | 0.0674 | 4.1928 | 2.75E-05 | 0.000431394 |
| MLF1IP | 3254.4396 | 0.2824 | 0.1002 | 2.8188 | 0.004821019 | 0.027697514 |
| MAST2 | 1510.7696 | 0.2824 | 0.1005 | 2.8102 | 0.004950632 | 0.028217111 |
| TOMM40 | 1415.8105 | 0.2820 | 0.1024 | 2.7548 | 0.005872757 | 0.031922043 |
| C15orf23 | 960.9386 | 0.2819 | 0.0979 | 2.8784 | 0.003997139 | 0.02383439 |
| NUP153 | 8245.1872 | 0.2818 | 0.0815 | 3.4592 | 0.000541789 | 0.004892923 |
| PKM2 | 158864.2991 | 0.2817 | 0.0562 | 5.0127 | 5.37E-07 | 1.56E-05 |
| ABCE1 | 4152.9568 | 0.2813 | 0.0950 | 2.9618 | 0.003058056 | 0.019266645 |
| AHCYL1 | 8033.4948 | 0.2809 | 0.0878 | 3.2006 | 0.001371524 | 0.010236378 |
| YBX1 | 32886.1638 | 0.2809 | 0.0875 | 3.2108 | 0.001323723 | 0.009985458 |
| SLC35A2 | 972.2934 | 0.2809 | 0.0933 | 3.0108 | 0.002605455 | 0.01693714 |
| RLIM | 4891.7595 | 0.2808 | 0.1030 | 2.7266 | 0.006399899 | 0.034208932 |
| ANKFY1 | 6645.7610 | 0.2807 | 0.0792 | 3.5460 | 0.000391186 | 0.003781135 |
| MLF2 | 5814.8006 | 0.2801 | 0.0838 | 3.3442 | 0.000825214 | 0.006847509 |
| KIFC1 | 4569.5087 | 0.2797 | 0.1059 | 2.6406 | 0.008274882 | 0.041420957 |
| SNX10 | 4226.5021 | 0.2797 | 0.0944 | 2.9619 | 0.003057366 | 0.019266645 |
| POLRMT | 2855.1406 | 0.2795 | 0.0880 | 3.1761 | 0.001492532 | 0.010975954 |
| IPO11 | 927.0815 | 0.2795 | 0.1077 | 2.5939 | 0.009488422 | 0.04601839 |
| TRPS1 | 1890.5291 | 0.2792 | 0.0998 | 2.7987 | 0.005130305 | 0.029043857 |
| FGFR1OP2 | 11147.1024 | 0.2790 | 0.0879 | 3.1750 | 0.001498581 | 0.011009727 |
| VARS | 3516.0009 | 0.2787 | 0.0739 | 3.7719 | 0.000162026 | 0.00185539 |
| MCM6 | 6966.6139 | 0.2786 | 0.0978 | 2.8498 | 0.004374312 | 0.025523271 |
| USP10 | 7088.1854 | 0.2786 | 0.0684 | 4.0716 | 4.67E-05 | 0.000676434 |
| DENND1A | 1973.8641 | 0.2776 | 0.0997 | 2.7857 | 0.00534158 | 0.029893483 |
| TRMU | 5075.7550 | 0.2775 | 0.0916 | 3.0284 | 0.002458358 | 0.016229996 |
| CBLB | 13170.9060 | 0.2771 | 0.1029 | 2.6941 | 0.007057545 | 0.036724967 |
| CCDC138 | 875.1620 | 0.2771 | 0.0911 | 3.0420 | 0.002350221 | 0.015710029 |
| RFWD3 | 3387.1525 | 0.2771 | 0.0963 | 2.8782 | 0.003999909 | 0.02383439 |
| CLASP1 | 10495.2227 | 0.2770 | 0.0997 | 2.7766 | 0.005493748 | 0.030452153 |
| TTI1 | 2627.9286 | 0.2764 | 0.0918 | 3.0120 | 0.002595486 | 0.016879588 |
| ZFAND5 | 11807.4296 | 0.2764 | 0.0798 | 3.4647 | 0.00053087 | 0.00481253 |
| NDE1 | 3062.6633 | 0.2757 | 0.0879 | 3.1360 | 0.001712658 | 0.012220458 |
| WDR43 | 2966.0304 | 0.2751 | 0.0879 | 3.1285 | 0.001756812 | 0.012470808 |
| NUP98 | 11191.9903 | 0.2737 | 0.0930 | 2.9427 | 0.003253392 | 0.020227726 |
| SGOL1 | 2049.6921 | 0.2734 | 0.1041 | 2.6252 | 0.008659458 | 0.042863753 |
| RNF111 | 4069.6059 | 0.2733 | 0.1017 | 2.6882 | 0.007184042 | 0.037204041 |
| IMPDH1 | 6081.0397 | 0.2730 | 0.0850 | 3.2136 | 0.001310869 | 0.009899904 |
| HK2 | 14066.6719 | 0.2728 | 0.0845 | 3.2290 | 0.001242314 | 0.009510368 |
| FKBP15 | 6273.1774 | 0.2724 | 0.0957 | 2.8460 | 0.004427842 | 0.025761523 |
| HMGXB4 | 3376.1163 | 0.2722 | 0.0988 | 2.7546 | 0.00587698 | 0.03193286 |
| TULP3 | 1224.0192 | 0.2717 | 0.1042 | 2.6084 | 0.009095558 | 0.04447299 |
| RAD54B | 1471.5113 | 0.2714 | 0.0890 | 3.0507 | 0.002283111 | 0.015335979 |
| PDIA4 | 9317.6594 | 0.2713 | 0.1018 | 2.6657 | 0.007681833 | 0.039155114 |
| SMS | 5142.1157 | 0.2709 | 0.0973 | 2.7830 | 0.005385958 | 0.030019673 |
| LARP4B | 8196.6359 | 0.2708 | 0.0667 | 4.0575 | 4.96E-05 | 0.000709725 |
| TMEM214 | 5655.6196 | 0.2704 | 0.0583 | 4.6375 | 3.53E-06 | 7.70E-05 |
| VEZT | 4039.5343 | 0.2701 | 0.0704 | 3.8372 | 0.000124441 | 0.001510765 |
| PNPLA8 | 2959.8010 | 0.2700 | 0.0902 | 2.9918 | 0.002773115 | 0.017865804 |
| CASP9 | 1899.5328 | 0.2695 | 0.0766 | 3.5171 | 0.000436216 | 0.004127917 |
| CABIN1 | 12730.7436 | 0.2694 | 0.0979 | 2.7515 | 0.005932468 | 0.032130483 |
| RBPJ | 44808.2168 | 0.2691 | 0.0807 | 3.3325 | 0.000860728 | 0.007087771 |
| SHCBP1 | 1965.8783 | 0.2686 | 0.0964 | 2.7861 | 0.005334517 | 0.029865004 |
| LIG1 | 4841.2155 | 0.2677 | 0.0853 | 3.1401 | 0.001689066 | 0.01210351 |
| CDK16 | 3421.5796 | 0.2673 | 0.0648 | 4.1262 | 3.69E-05 | 0.00055292 |
| MZT1 | 3715.6086 | 0.2672 | 0.0969 | 2.7578 | 0.005819093 | 0.031743753 |
| SAMD8 | 3294.0532 | 0.2670 | 0.0916 | 2.9158 | 0.003547682 | 0.021665636 |
| RCOR1 | 3667.3670 | 0.2669 | 0.0790 | 3.3790 | 0.000727627 | 0.006204516 |
| MLLT6 | 25690.8385 | 0.2666 | 0.0815 | 3.2700 | 0.001075636 | 0.008461338 |
| STRBP | 1932.0292 | 0.2660 | 0.0873 | 3.0491 | 0.002295078 | 0.015402679 |
| IMPA1 | 2886.8897 | 0.2660 | 0.0886 | 3.0033 | 0.002670515 | 0.017271002 |
| PLOD1 | 2503.9755 | 0.2660 | 0.0721 | 3.6891 | 0.000225053 | 0.002429962 |
| ASXL1 | 15899.1450 | 0.2655 | 0.0647 | 4.1017 | 4.10E-05 | 0.000605258 |
| PITPNM1 | 6118.8897 | 0.2652 | 0.0841 | 3.1523 | 0.001620119 | 0.011698109 |
| ALKBH5 | 13558.9199 | 0.2650 | 0.0606 | 4.3712 | 1.24E-05 | 0.000222835 |
| ZCCHC8 | 3971.5284 | 0.2644 | 0.0859 | 3.0765 | 0.002094131 | 0.014342782 |
| DDX47 | 5060.2873 | 0.2644 | 0.0859 | 3.0785 | 0.002080233 | 0.014264751 |
| MLH1 | 3011.0352 | 0.2633 | 0.0804 | 3.2735 | 0.001062303 | 0.008386985 |
| AZIN1 | 12198.8923 | 0.2631 | 0.0698 | 3.7705 | 0.000162922 | 0.001860017 |
| C18orf25 | 4654.3339 | 0.2631 | 0.0884 | 2.9757 | 0.002922766 | 0.018655139 |
| FBXO42 | 3200.7757 | 0.2629 | 0.0714 | 3.6838 | 0.000229742 | 0.002461262 |
| PHF11 | 6513.3872 | 0.2628 | 0.0579 | 4.5370 | 5.71E-06 | 0.0001184 |
| DKC1 | 3187.6827 | 0.2626 | 0.0804 | 3.2644 | 0.001096799 | 0.008574308 |
| PHLPP1 | 1475.5299 | 0.2625 | 0.0976 | 2.6903 | 0.007139155 | 0.037022281 |
| TRA2B | 10399.4471 | 0.2624 | 0.0680 | 3.8567 | 0.000114937 | 0.001408954 |
| RFTN1 | 6911.6337 | 0.2623 | 0.0971 | 2.7009 | 0.006916218 | 0.036238874 |
| MEF2D | 7615.1550 | 0.2617 | 0.0826 | 3.1699 | 0.001524932 | 0.011165363 |
| CSNK2A1 | 6409.9302 | 0.2616 | 0.0841 | 3.1094 | 0.001874821 | 0.01311768 |
| CNOT4 | 3581.5409 | 0.2612 | 0.0824 | 3.1696 | 0.001526643 | 0.011172483 |
| C11orf75 | 3490.0243 | 0.2609 | 0.0888 | 2.9401 | 0.003281366 | 0.020359812 |
| PGD | 3281.3688 | 0.2608 | 0.0813 | 3.2081 | 0.001336314 | 0.010041938 |
| SETD5 | 16114.3625 | 0.2607 | 0.0878 | 2.9677 | 0.003000029 | 0.01902489 |
| RAB3GAP1 | 3973.0841 | 0.2598 | 0.0716 | 3.6258 | 0.000288054 | 0.002960185 |
| TMEM57 | 3739.5634 | 0.2593 | 0.0905 | 2.8661 | 0.004155555 | 0.024564704 |
| SCAF4 | 4927.8701 | 0.2591 | 0.0948 | 2.7323 | 0.006289293 | 0.033681656 |
| SCLT1 | 2603.8816 | 0.2591 | 0.0965 | 2.6858 | 0.007236306 | 0.037415594 |
| DCUN1D5 | 3104.0030 | 0.2590 | 0.0900 | 2.8793 | 0.003985124 | 0.023780268 |
| UBE2O | 4710.4814 | 0.2588 | 0.0681 | 3.7978 | 0.000145973 | 0.001706445 |
| ORC3 | 2360.8180 | 0.2585 | 0.0788 | 3.2817 | 0.001031966 | 0.008206245 |
| XPNPEP1 | 9019.4808 | 0.2584 | 0.0823 | 3.1381 | 0.001700267 | 0.012166479 |
| NET1 | 2254.3199 | 0.2581 | 0.0776 | 3.3271 | 0.000877441 | 0.007197965 |
| AGPAT6 | 8208.8904 | 0.2581 | 0.0708 | 3.6430 | 0.00026945 | 0.002803281 |
| SGTB | 3748.4227 | 0.2577 | 0.0935 | 2.7567 | 0.005838246 | 0.031825278 |
| PLK4 | 2103.2316 | 0.2574 | 0.0979 | 2.6295 | 0.008550075 | 0.042449922 |
| CLEC16A | 3248.7943 | 0.2573 | 0.0631 | 4.0801 | 4.50E-05 | 0.000654635 |
| UBC | 62731.7887 | 0.2566 | 0.0850 | 3.0198 | 0.002529243 | 0.016535358 |
| NBAS | 4165.5610 | 0.2562 | 0.0734 | 3.4886 | 0.000485479 | 0.004488897 |
| MELK | 1999.2646 | 0.2557 | 0.0960 | 2.6623 | 0.007760147 | 0.039431555 |
| POLA1 | 2030.2653 | 0.2556 | 0.0975 | 2.6206 | 0.008778279 | 0.043256562 |
| STAU1 | 8307.3702 | 0.2552 | 0.0534 | 4.7811 | 1.74E-06 | 4.21E-05 |
| BPGM | 1396.8041 | 0.2548 | 0.0785 | 3.2463 | 0.001169356 | 0.009024926 |
| REXO2 | 4005.7011 | 0.2547 | 0.0783 | 3.2513 | 0.001148787 | 0.008910567 |
| C2orf48 | 1521.2132 | 0.2537 | 0.0941 | 2.6974 | 0.006989014 | 0.036456141 |
| SRPK1 | 6778.7330 | 0.2534 | 0.0818 | 3.0961 | 0.001960782 | 0.01360585 |
| RAP1A | 23946.0733 | 0.2534 | 0.0632 | 4.0067 | 6.16E-05 | 0.000850749 |
| CLK4 | 2922.9222 | 0.2532 | 0.0803 | 3.1533 | 0.001614322 | 0.011672851 |
| SCYL1 | 4231.1644 | 0.2530 | 0.0741 | 3.4127 | 0.0006432 | 0.005611123 |
| COPS2 | 7883.2590 | 0.2528 | 0.0852 | 2.9677 | 0.003000276 | 0.01902489 |
| TNFAIP1 | 2100.2735 | 0.2523 | 0.0971 | 2.5981 | 0.009374253 | 0.045545946 |
| CHD2 | 22654.5303 | 0.2513 | 0.0903 | 2.7822 | 0.005399162 | 0.030071107 |
| PDP2 | 3313.4245 | 0.2512 | 0.0862 | 2.9149 | 0.003558441 | 0.021705056 |
| ATG4B | 6547.9153 | 0.2505 | 0.0748 | 3.3481 | 0.000813766 | 0.006772327 |
| LRRC40 | 2366.2618 | 0.2505 | 0.0959 | 2.6112 | 0.009023317 | 0.044173369 |
| B4GALT4 | 1468.9176 | 0.2502 | 0.0968 | 2.5851 | 0.009736352 | 0.046919974 |
| WBP11 | 8923.2026 | 0.2489 | 0.0677 | 3.6766 | 0.000236366 | 0.00251442 |
| CEBPG | 4024.9438 | 0.2488 | 0.0773 | 3.2186 | 0.001288183 | 0.0097627 |
| ARID5B | 4967.5088 | 0.2487 | 0.0910 | 2.7340 | 0.006257471 | 0.033578135 |
| ARF6 | 18732.5324 | 0.2486 | 0.0711 | 3.4962 | 0.000471942 | 0.004390573 |
| ACSL4 | 9793.3168 | 0.2486 | 0.0900 | 2.7630 | 0.005727898 | 0.031325349 |
| MAN2B1 | 10788.2652 | 0.2483 | 0.0704 | 3.5288 | 0.0004175 | 0.003992115 |
| HELLS | 1765.8830 | 0.2482 | 0.0919 | 2.6994 | 0.006947192 | 0.036283412 |
| HDAC3 | 8221.7819 | 0.2479 | 0.0705 | 3.5188 | 0.00043356 | 0.004111887 |
| CLDND1 | 30848.4453 | 0.2473 | 0.0841 | 2.9416 | 0.003264741 | 0.020281615 |
| POLR3A | 2167.3067 | 0.2461 | 0.0668 | 3.6863 | 0.000227526 | 0.002449434 |
| COPS7B | 3476.7238 | 0.2456 | 0.0907 | 2.7072 | 0.006784608 | 0.035797267 |
| SRRM1 | 10055.0574 | 0.2452 | 0.0844 | 2.9038 | 0.00368722 | 0.022319561 |
| SSRP1 | 11541.9494 | 0.2450 | 0.0686 | 3.5746 | 0.000350756 | 0.003474075 |
| TBC1D22B | 2648.1734 | 0.2450 | 0.0854 | 2.8679 | 0.004131858 | 0.024443728 |
| USP32 | 4033.9620 | 0.2447 | 0.0702 | 3.4868 | 0.000488883 | 0.004509105 |
| SKIV2L2 | 4467.6609 | 0.2441 | 0.0889 | 2.7464 | 0.006024847 | 0.032526001 |
| CCDC50 | 6850.6454 | 0.2425 | 0.0771 | 3.1463 | 0.001653744 | 0.01188417 |
| SEMA4B | 2844.8839 | 0.2423 | 0.0705 | 3.4383 | 0.000585393 | 0.005208963 |
| STX16 | 18696.5130 | 0.2416 | 0.0729 | 3.3126 | 0.00092441 | 0.007505934 |
| PVRL3 | 795.1821 | 0.2413 | 0.0893 | 2.7019 | 0.00689509 | 0.036165752 |
| WHSC1 | 16243.9806 | 0.2409 | 0.0792 | 3.0397 | 0.00236848 | 0.015811118 |
| POC1B | 5217.9455 | 0.2408 | 0.0876 | 2.7486 | 0.005985774 | 0.032384409 |
| C16orf72 | 7995.0981 | 0.2408 | 0.0699 | 3.4454 | 0.000570297 | 0.005104666 |
| MTCH2 | 2636.4964 | 0.2405 | 0.0818 | 2.9413 | 0.003268367 | 0.020287481 |
| EZH2 | 10047.7547 | 0.2401 | 0.0857 | 2.8005 | 0.005102151 | 0.028917284 |
| SRGAP2 | 5977.2562 | 0.2387 | 0.0707 | 3.3772 | 0.000732332 | 0.006239435 |
| TSPYL2 | 3846.4592 | 0.2386 | 0.0922 | 2.5884 | 0.009641692 | 0.046612295 |
| RUNX3 | 17298.5052 | 0.2378 | 0.0639 | 3.7223 | 0.000197409 | 0.002189805 |
| IL27RA | 5078.4388 | 0.2378 | 0.0910 | 2.6148 | 0.008927656 | 0.043818511 |
| COPB2 | 10121.8802 | 0.2371 | 0.0752 | 3.1513 | 0.001625375 | 0.011730461 |
| UAP1 | 2527.1885 | 0.2370 | 0.0898 | 2.6379 | 0.008342981 | 0.041637833 |
| RRAS2 | 3029.0478 | 0.2368 | 0.0871 | 2.7187 | 0.00655393 | 0.034847556 |
| MTERFD1 | 2313.7764 | 0.2365 | 0.0849 | 2.7855 | 0.005343706 | 0.029893841 |
| TSPAN5 | 3227.1296 | 0.2361 | 0.0776 | 3.0445 | 0.002330652 | 0.015613719 |
| NFATC1 | 4332.1997 | 0.2355 | 0.0780 | 3.0196 | 0.002530767 | 0.016536897 |
| MFAP1 | 2968.7886 | 0.2353 | 0.0723 | 3.2561 | 0.001129659 | 0.008785783 |
| RPS6KA1 | 18703.0773 | 0.2337 | 0.0680 | 3.4350 | 0.000592551 | 0.005247966 |
| WDR67 | 3082.1364 | 0.2332 | 0.0857 | 2.7206 | 0.006516226 | 0.034720309 |
| IPO5 | 8688.5654 | 0.2328 | 0.0656 | 3.5478 | 0.000388529 | 0.003757851 |
| CASP3 | 5808.1035 | 0.2325 | 0.0784 | 2.9638 | 0.003039039 | 0.019200158 |
| SNHG1 | 6782.6078 | 0.2322 | 0.0875 | 2.6533 | 0.007971286 | 0.040194561 |
| C15orf29 | 2161.4579 | 0.2322 | 0.0856 | 2.7112 | 0.006703373 | 0.035455218 |
| LRBA | 8845.4111 | 0.2313 | 0.0885 | 2.6126 | 0.008985211 | 0.044015315 |
| SFXN3 | 10691.1248 | 0.2306 | 0.0724 | 3.1860 | 0.001442614 | 0.010692025 |
| RNF121 | 1834.4404 | 0.2305 | 0.0642 | 3.5902 | 0.000330453 | 0.003312636 |
| SAFB2 | 8997.8803 | 0.2301 | 0.0848 | 2.7120 | 0.006688834 | 0.035403078 |
| SLC4A5 | 1988.1907 | 0.2299 | 0.0859 | 2.6753 | 0.007467134 | 0.038420182 |
| YARS | 11362.8714 | 0.2298 | 0.0662 | 3.4712 | 0.000518098 | 0.004720314 |
| GYS1 | 6958.7397 | 0.2297 | 0.0887 | 2.5880 | 0.009654107 | 0.046657406 |
| PISD | 2605.6431 | 0.2290 | 0.0819 | 2.7949 | 0.005191739 | 0.029249695 |
| RAD23A | 6204.8335 | 0.2287 | 0.0831 | 2.7523 | 0.005918432 | 0.03208893 |
| USP14 | 4579.2803 | 0.2281 | 0.0873 | 2.6137 | 0.008956375 | 0.043916719 |
| ESCO1 | 4533.6023 | 0.2279 | 0.0758 | 3.0062 | 0.002645564 | 0.017153639 |
| HEMK1 | 3056.0285 | 0.2263 | 0.0727 | 3.1126 | 0.001854205 | 0.013009538 |
| LRRC59 | 3203.5777 | 0.2262 | 0.0762 | 2.9696 | 0.002981958 | 0.018968912 |
| DPM1 | 4209.5067 | 0.2261 | 0.0878 | 2.5770 | 0.00996481 | 0.047701797 |
| AVL9 | 3222.7865 | 0.2256 | 0.0769 | 2.9338 | 0.003348183 | 0.020664201 |
| TNIP1 | 25669.6768 | 0.2255 | 0.0647 | 3.4866 | 0.000489155 | 0.004509105 |
| USP48 | 13599.5701 | 0.2253 | 0.0859 | 2.6216 | 0.008752189 | 0.043181464 |
| MSMO1 | 7361.9224 | 0.2245 | 0.0789 | 2.8447 | 0.00444577 | 0.025855889 |
| ZNF526 | 1194.1151 | 0.2242 | 0.0851 | 2.6341 | 0.008435766 | 0.042003893 |
| SRP68 | 7019.1649 | 0.2236 | 0.0719 | 3.1114 | 0.001861805 | 0.013050751 |
| G3BP2 | 10460.0267 | 0.2236 | 0.0755 | 2.9619 | 0.003057967 | 0.019266645 |
| GRIPAP1 | 10901.3656 | 0.2229 | 0.0586 | 3.8047 | 0.000141996 | 0.001676792 |
| NPTN | 3293.0399 | 0.2226 | 0.0655 | 3.4006 | 0.000672384 | 0.005818737 |
| PA2G4 | 7902.3187 | 0.2216 | 0.0802 | 2.7647 | 0.005697209 | 0.031247888 |
| TYMS | 6248.4968 | 0.2209 | 0.0778 | 2.8409 | 0.004499019 | 0.026125398 |
| NUMB | 2472.8013 | 0.2205 | 0.0725 | 3.0434 | 0.002339507 | 0.015659166 |
| AAGAB | 2918.7837 | 0.2205 | 0.0768 | 2.8725 | 0.004072287 | 0.024138513 |
| DNAJB6 | 9523.6899 | 0.2192 | 0.0820 | 2.6726 | 0.007526157 | 0.038618785 |
| NUS1 | 3676.7502 | 0.2191 | 0.0558 | 3.9233 | 8.74E-05 | 0.001123633 |
| PPPDE1 | 4253.7458 | 0.2190 | 0.0693 | 3.1586 | 0.001585303 | 0.011492842 |
| MDH1 | 8388.5525 | 0.2189 | 0.0767 | 2.8553 | 0.004299836 | 0.025210704 |
| TMEM48 | 2509.5573 | 0.2187 | 0.0712 | 3.0736 | 0.002114608 | 0.014447913 |
| MAPK1IP1L | 8933.4359 | 0.2186 | 0.0585 | 3.7353 | 0.000187464 | 0.002102819 |
| MED17 | 2638.6707 | 0.2182 | 0.0835 | 2.6151 | 0.008921387 | 0.043801956 |
| UBA1 | 15059.2397 | 0.2177 | 0.0548 | 3.9704 | 7.18E-05 | 0.000962389 |
| DAP | 5849.1902 | 0.2172 | 0.0788 | 2.7550 | 0.005869498 | 0.031922043 |
| ZFP64 | 1038.1416 | 0.2171 | 0.0759 | 2.8626 | 0.004201589 | 0.024769073 |
| KCTD10 | 3736.1601 | 0.2163 | 0.0677 | 3.1936 | 0.001405138 | 0.010444482 |
| HNRNPL | 18294.1062 | 0.2157 | 0.0677 | 3.1840 | 0.001452642 | 0.010742725 |
| MKRN2 | 2900.0034 | 0.2152 | 0.0782 | 2.7530 | 0.005904864 | 0.032038334 |
| DCAF13 | 1930.3596 | 0.2145 | 0.0816 | 2.6289 | 0.008566091 | 0.04251288 |
| TPM4 | 20602.0116 | 0.2143 | 0.0713 | 3.0038 | 0.002666365 | 0.01725154 |
| FBXW11 | 3677.9671 | 0.2135 | 0.0721 | 2.9606 | 0.003070439 | 0.019320521 |
| BUB3 | 15880.1740 | 0.2133 | 0.0723 | 2.9494 | 0.003183988 | 0.019894334 |
| SPATA2 | 830.9398 | 0.2125 | 0.0786 | 2.7046 | 0.006838466 | 0.035976009 |
| CAND1 | 9539.1877 | 0.2124 | 0.0646 | 3.2899 | 0.001002263 | 0.008004871 |
| PTBP1 | 14672.9553 | 0.2123 | 0.0698 | 3.0431 | 0.002341187 | 0.015663483 |
| CTDSPL2 | 5760.4292 | 0.2122 | 0.0749 | 2.8341 | 0.004594871 | 0.026590137 |
| RDH11 | 4781.0113 | 0.2121 | 0.0738 | 2.8756 | 0.00403262 | 0.023950313 |
| TBX19 | 6163.3775 | 0.2114 | 0.0644 | 3.2819 | 0.001031104 | 0.008206245 |
| GRSF1 | 9029.8720 | 0.2113 | 0.0603 | 3.5017 | 0.000462338 | 0.004325165 |
| PSMD1 | 4273.8624 | 0.2111 | 0.0769 | 2.7448 | 0.006054934 | 0.032644172 |
| C3orf19 | 2403.5399 | 0.2110 | 0.0730 | 2.8900 | 0.003851951 | 0.023094914 |
| SELT | 4441.2498 | 0.2105 | 0.0712 | 2.9560 | 0.003116515 | 0.019577876 |
| AGFG1 | 6087.6673 | 0.2103 | 0.0603 | 3.4869 | 0.0004886 | 0.004509105 |
| ARF4 | 6418.3624 | 0.2097 | 0.0790 | 2.6523 | 0.00799425 | 0.040282816 |
| DBF4 | 3211.1739 | 0.2095 | 0.0741 | 2.8254 | 0.004722137 | 0.027212102 |
| LDHA | 114228.9909 | 0.2086 | 0.0617 | 3.3803 | 0.000724049 | 0.00618098 |
| USP15 | 6650.3919 | 0.2085 | 0.0759 | 2.7467 | 0.006019873 | 0.032522365 |
| KPNA6 | 5587.0803 | 0.2084 | 0.0582 | 3.5802 | 0.000343387 | 0.00341963 |
| SCOC | 3457.6605 | 0.2057 | 0.0786 | 2.6188 | 0.008824925 | 0.043415919 |
| FTSJD2 | 5802.7162 | 0.2041 | 0.0766 | 2.6651 | 0.007696436 | 0.039179032 |
| KCMF1 | 6573.4521 | 0.2034 | 0.0737 | 2.7585 | 0.005807434 | 0.03169158 |
| ATXN2 | 2484.4950 | 0.2026 | 0.0758 | 2.6740 | 0.007495636 | 0.038501353 |
| SRSF7 | 8907.2455 | 0.2024 | 0.0740 | 2.7347 | 0.006242829 | 0.033523346 |
| VAPB | 3463.5893 | 0.2023 | 0.0742 | 2.7258 | 0.006415406 | 0.034267601 |
| FAM193A | 4350.3065 | 0.2021 | 0.0721 | 2.8034 | 0.00505635 | 0.028689951 |
| TDG | 2478.9991 | 0.2017 | 0.0741 | 2.7219 | 0.006491011 | 0.034610351 |
| DIS3 | 5555.1215 | 0.2011 | 0.0678 | 2.9654 | 0.003022984 | 0.019125334 |
| MTMR2 | 2307.0645 | 0.2004 | 0.0734 | 2.7321 | 0.006292505 | 0.033682491 |
| PRMT5 | 2027.4254 | 0.1998 | 0.0741 | 2.6955 | 0.007027737 | 0.036607637 |
| WHSC2 | 3442.1606 | 0.1985 | 0.0628 | 3.1581 | 0.001587952 | 0.011504288 |
| XPO6 | 18038.4936 | 0.1979 | 0.0488 | 4.0590 | 4.93E-05 | 0.000706783 |
| QSOX1 | 3454.4674 | 0.1965 | 0.0679 | 2.8934 | 0.003810354 | 0.02291818 |
| DAZAP1 | 13038.8359 | 0.1964 | 0.0569 | 3.4497 | 0.000561137 | 0.005043235 |
| RAP2C | 6159.5124 | 0.1960 | 0.0674 | 2.9067 | 0.003652848 | 0.022146367 |
| OTUD5 | 7355.0362 | 0.1956 | 0.0608 | 3.2202 | 0.001280966 | 0.009712872 |
| UBR3 | 3721.8910 | 0.1953 | 0.0738 | 2.6458 | 0.008149949 | 0.040931035 |
| TBRG4 | 3357.5499 | 0.1947 | 0.0735 | 2.6475 | 0.008108905 | 0.040765505 |
| SYAP1 | 2697.3437 | 0.1946 | 0.0567 | 3.4323 | 0.000598398 | 0.005285778 |
| RSBN1 | 4361.4217 | 0.1939 | 0.0692 | 2.8041 | 0.005045215 | 0.02865902 |
| SERBP1 | 17197.5679 | 0.1934 | 0.0578 | 3.3446 | 0.00082404 | 0.006841521 |
| MAML1 | 5376.8512 | 0.1933 | 0.0739 | 2.6155 | 0.008909152 | 0.043770295 |
| WIPF2 | 4327.0144 | 0.1931 | 0.0747 | 2.5845 | 0.009753255 | 0.046986459 |
| XPO1 | 17136.5416 | 0.1924 | 0.0698 | 2.7548 | 0.005872879 | 0.031922043 |
| GTF3C1 | 6166.9150 | 0.1921 | 0.0726 | 2.6449 | 0.008172295 | 0.041029639 |
| UBA2 | 11418.3387 | 0.1920 | 0.0635 | 3.0221 | 0.002510372 | 0.016467646 |
| RCC2 | 7972.3775 | 0.1918 | 0.0742 | 2.5855 | 0.009722764 | 0.046884364 |
| RARA | 3771.8845 | 0.1895 | 0.0721 | 2.6274 | 0.008603425 | 0.042684162 |
| APBA3 | 5467.9691 | 0.1895 | 0.0685 | 2.7670 | 0.005658206 | 0.031056487 |
| RAB6A | 13970.2806 | 0.1894 | 0.0614 | 3.0835 | 0.002045576 | 0.014071593 |
| RAB27A | 9551.1974 | 0.1887 | 0.0704 | 2.6792 | 0.007380688 | 0.038079013 |
| HTT | 8921.8891 | 0.1872 | 0.0716 | 2.6145 | 0.008935531 | 0.043842469 |
| STK11 | 6945.5018 | 0.1871 | 0.0698 | 2.6804 | 0.007352729 | 0.037960657 |
| PLEKHB2 | 6569.6585 | 0.1862 | 0.0700 | 2.6600 | 0.007814467 | 0.039650907 |
| CSE1L | 4103.8391 | 0.1852 | 0.0663 | 2.7952 | 0.005186852 | 0.029233052 |
| HIAT1 | 5792.4312 | 0.1829 | 0.0686 | 2.6657 | 0.007682605 | 0.039155114 |
| RNF114 | 8004.2739 | 0.1825 | 0.0638 | 2.8629 | 0.004197645 | 0.024755469 |
| GSPT1 | 9002.5075 | 0.1809 | 0.0447 | 4.0437 | 5.26E-05 | 0.000745268 |
| GRB2 | 12059.1145 | 0.1775 | 0.0513 | 3.4585 | 0.000543292 | 0.00490357 |
| EIF2D | 5973.0013 | 0.1770 | 0.0680 | 2.6030 | 0.009241926 | 0.045039503 |
| DR1 | 11513.1806 | 0.1723 | 0.0625 | 2.7554 | 0.005862331 | 0.031910572 |
| SRSF1 | 16149.4223 | 0.1717 | 0.0544 | 3.1527 | 0.001617553 | 0.011685162 |
| TMEM209 | 2107.4503 | 0.1691 | 0.0611 | 2.7657 | 0.005680876 | 0.031169608 |
| CNOT8 | 9977.7874 | 0.1627 | 0.0598 | 2.7219 | 0.006491014 | 0.034610351 |
| MED24 | 6076.5113 | 0.1603 | 0.0614 | 2.6124 | 0.008991412 | 0.044031433 |
| COX10 | 1953.8225 | 0.1584 | 0.0569 | 2.7844 | 0.005362033 | 0.029974678 |
| PITPNB | 5341.8367 | 0.1561 | 0.0602 | 2.5908 | 0.00957486 | 0.046378132 |
| DDX24 | 12355.9580 | 0.1559 | 0.0421 | 3.6991 | 0.000216406 | 0.002355085 |
| BAG6 | 15389.1392 | 0.1527 | 0.0531 | 2.8741 | 0.004052277 | 0.024048174 |
| SRSF4 | 14505.7266 | 0.1506 | 0.0572 | 2.6335 | 0.008451254 | 0.042011869 |
| GOLGA8A | 34259.9583 | -0.1425 | 0.0534 | -2.6665 | 0.007664842 | 0.039104911 |
| SH2D3C | 6725.4269 | -0.1454 | 0.0540 | -2.6933 | 0.007073857 | 0.036771904 |
| SNX18 | 4916.6713 | -0.1473 | 0.0553 | -2.6654 | 0.007689983 | 0.039167127 |
| UNC13D | 12487.1350 | -0.1512 | 0.0586 | -2.5811 | 0.009847431 | 0.047297531 |
| ANAPC5 | 10261.2538 | -0.1529 | 0.0559 | -2.7362 | 0.006215459 | 0.033400088 |
| PSEN1 | 4829.3341 | -0.1576 | 0.0564 | -2.7968 | 0.005161583 | 0.029131426 |
| HEXIM1 | 6048.7944 | -0.1585 | 0.0613 | -2.5868 | 0.009685827 | 0.046750956 |
| CAPN1 | 8525.6821 | -0.1602 | 0.0612 | -2.6189 | 0.008822029 | 0.043415919 |
| DSCR3 | 7112.3862 | -0.1642 | 0.0554 | -2.9610 | 0.003065984 | 0.019300519 |
| HMGN4 | 8646.8694 | -0.1676 | 0.0547 | -3.0658 | 0.002170538 | 0.01478997 |
| GLG1 | 16111.5733 | -0.1695 | 0.0581 | -2.9155 | 0.003550674 | 0.021675157 |
| AHSA2 | 8288.5316 | -0.1718 | 0.0613 | -2.8036 | 0.005053442 | 0.02868421 |
| PRKAG1 | 3081.6436 | -0.1724 | 0.0644 | -2.6763 | 0.007444052 | 0.038353604 |
| VPS26B | 9177.6521 | -0.1740 | 0.0547 | -3.1832 | 0.001456721 | 0.010754427 |
| ZFAND6 | 4184.1886 | -0.1767 | 0.0663 | -2.6661 | 0.007672637 | 0.039131484 |
| ZNF33A | 4851.9752 | -0.1768 | 0.0671 | -2.6336 | 0.008447907 | 0.042011869 |
| RAB5B | 9698.3932 | -0.1779 | 0.0517 | -3.4379 | 0.000586146 | 0.005212595 |
| ELMOD2 | 1638.2193 | -0.1811 | 0.0685 | -2.6433 | 0.008209718 | 0.041203849 |
| CDR2 | 3614.9815 | -0.1813 | 0.0701 | -2.5862 | 0.009704921 | 0.046813247 |
| ZER1 | 4792.6628 | -0.1813 | 0.0649 | -2.7929 | 0.005223761 | 0.029353579 |
| HNRNPA1L2 | 2004.9024 | -0.1816 | 0.0657 | -2.7622 | 0.0057422 | 0.031392217 |
| HCLS1 | 27433.9706 | -0.1828 | 0.0635 | -2.8768 | 0.004017215 | 0.023896345 |
| STAT6 | 13177.7314 | -0.1829 | 0.0573 | -3.1922 | 0.001411727 | 0.010483647 |
| PRKRIR | 3017.5153 | -0.1839 | 0.0644 | -2.8568 | 0.004278728 | 0.025135658 |
| VPS11 | 3740.5832 | -0.1840 | 0.0575 | -3.1992 | 0.001378083 | 0.010274158 |
| TBRG1 | 6357.3716 | -0.1844 | 0.0712 | -2.5895 | 0.009611007 | 0.046523423 |
| PSAP | 18005.9897 | -0.1873 | 0.0660 | -2.8356 | 0.004573732 | 0.026488074 |
| C2orf89 | 3196.8826 | -0.1896 | 0.0718 | -2.6392 | 0.008310368 | 0.041529876 |
| GBA2 | 4089.9032 | -0.1896 | 0.0628 | -3.0190 | 0.002536034 | 0.016564157 |
| OGDH | 9791.1064 | -0.1914 | 0.0578 | -3.3085 | 0.000938 | 0.007591831 |
| CNDP2 | 5556.9138 | -0.1916 | 0.0722 | -2.6544 | 0.007944498 | 0.040125684 |
| MECP2 | 12975.8431 | -0.1924 | 0.0663 | -2.9029 | 0.003697656 | 0.022357548 |
| ARHGAP30 | 31795.6186 | -0.1933 | 0.0438 | -4.4169 | 1.00E-05 | 0.000187428 |
| MICAL1 | 12688.9840 | -0.1968 | 0.0703 | -2.7985 | 0.005133267 | 0.029043857 |
| PIP4K2A | 12215.6125 | -0.1978 | 0.0591 | -3.3485 | 0.000812547 | 0.00677212 |
| EXOC4 | 5039.5899 | -0.1979 | 0.0724 | -2.7326 | 0.006282947 | 0.033676552 |
| PPP6C | 4800.3417 | -0.1979 | 0.0663 | -2.9875 | 0.002813046 | 0.01808455 |
| HADH | 2480.0271 | -0.1982 | 0.0744 | -2.6649 | 0.007701818 | 0.039187824 |
| GTPBP3 | 1396.6791 | -0.2003 | 0.0719 | -2.7854 | 0.005345597 | 0.029893841 |
| GIT1 | 6089.8778 | -0.2010 | 0.0689 | -2.9159 | 0.003546939 | 0.021665636 |
| YWHAZ | 58507.6288 | -0.2011 | 0.0691 | -2.9082 | 0.003635148 | 0.02206797 |
| YWHAQ | 19034.7144 | -0.2019 | 0.0676 | -2.9865 | 0.002821968 | 0.018126502 |
| TFCP2 | 3291.4874 | -0.2021 | 0.0763 | -2.6497 | 0.008056506 | 0.040551038 |
| GUSBP11 | 4787.1902 | -0.2036 | 0.0672 | -3.0279 | 0.002462409 | 0.016237517 |
| DAXX | 6128.5476 | -0.2036 | 0.0767 | -2.6540 | 0.007953591 | 0.040144804 |
| TMUB2 | 3610.2757 | -0.2040 | 0.0730 | -2.7935 | 0.005214341 | 0.029322431 |
| CYTH2 | 3191.0953 | -0.2049 | 0.0671 | -3.0544 | 0.002255268 | 0.015175905 |
| AGAP2 | 5441.7544 | -0.2056 | 0.0686 | -2.9988 | 0.002710406 | 0.017499065 |
| MAVS | 9268.3372 | -0.2061 | 0.0799 | -2.5808 | 0.009856482 | 0.047302982 |
| ACOT13 | 1476.1359 | -0.2066 | 0.0797 | -2.5902 | 0.009593083 | 0.046451526 |
| NAA38 | 4870.8787 | -0.2070 | 0.0676 | -3.0621 | 0.002198265 | 0.014900797 |
| HMHA1 | 37870.4511 | -0.2072 | 0.0756 | -2.7408 | 0.006128319 | 0.03299042 |
| RTN3 | 8296.4305 | -0.2084 | 0.0686 | -3.0361 | 0.002396582 | 0.015949448 |
| GALNT11 | 2619.5130 | -0.2084 | 0.0795 | -2.6226 | 0.008726581 | 0.043080997 |
| PITPNA | 4270.1478 | -0.2085 | 0.0790 | -2.6399 | 0.008292298 | 0.041466975 |
| GNB5 | 2210.0542 | -0.2090 | 0.0697 | -3.0000 | 0.00269951 | 0.017436159 |
| RSU1 | 5120.3195 | -0.2090 | 0.0811 | -2.5771 | 0.009963289 | 0.047701797 |
| ZBED5 | 4011.6863 | -0.2097 | 0.0696 | -3.0133 | 0.002584101 | 0.016820009 |
| PHKB | 3533.5145 | -0.2107 | 0.0566 | -3.7194 | 0.000199678 | 0.002207987 |
| ABR | 9703.0512 | -0.2118 | 0.0747 | -2.8362 | 0.0045658 | 0.02645226 |
| TRAPPC6B | 4387.1894 | -0.2127 | 0.0777 | -2.7391 | 0.006160915 | 0.033154093 |
| TBC1D5 | 5837.2282 | -0.2128 | 0.0703 | -3.0287 | 0.002456245 | 0.016225161 |
| PRKCB | 17707.8115 | -0.2130 | 0.0813 | -2.6191 | 0.008816835 | 0.043415449 |
| GET4 | 5914.0312 | -0.2132 | 0.0712 | -2.9930 | 0.00276226 | 0.017826239 |
| PYGO2 | 3682.2526 | -0.2133 | 0.0632 | -3.3771 | 0.000732547 | 0.006239435 |
| PRMT2 | 12547.4450 | -0.2135 | 0.0723 | -2.9538 | 0.003138406 | 0.019674539 |
| CDV3 | 15805.9749 | -0.2141 | 0.0533 | -4.0149 | 5.95E-05 | 0.000826226 |
| FAM116A | 2839.2799 | -0.2142 | 0.0781 | -2.7412 | 0.006122426 | 0.032970428 |
| ARL4C | 34271.8958 | -0.2145 | 0.0678 | -3.1611 | 0.001571516 | 0.01141254 |
| KIAA1967 | 7385.4701 | -0.2148 | 0.0700 | -3.0707 | 0.002135221 | 0.01457558 |
| KRI1 | 3553.7000 | -0.2174 | 0.0761 | -2.8560 | 0.004289866 | 0.025181532 |
| UBXN2B | 2909.6597 | -0.2175 | 0.0840 | -2.5872 | 0.009674941 | 0.046728234 |
| ZDHHC24 | 1930.4664 | -0.2179 | 0.0788 | -2.7636 | 0.005717273 | 0.0312974 |
| N4BP2L2 | 19583.8180 | -0.2180 | 0.0676 | -3.2243 | 0.001262611 | 0.009602573 |
| MFGE8 | 2008.0157 | -0.2181 | 0.0803 | -2.7148 | 0.006631234 | 0.035135087 |
| GOLGA8B | 14470.7725 | -0.2197 | 0.0614 | -3.5766 | 0.000348074 | 0.003452662 |
| PECR | 2562.5684 | -0.2202 | 0.0780 | -2.8242 | 0.004739582 | 0.027282015 |
| NFATC3 | 20585.4993 | -0.2216 | 0.0804 | -2.7551 | 0.005866776 | 0.031922043 |
| ANAPC7 | 2263.0892 | -0.2217 | 0.0852 | -2.6019 | 0.009271091 | 0.045101427 |
| CCNDBP1 | 5967.2172 | -0.2222 | 0.0832 | -2.6698 | 0.007588842 | 0.038829681 |
| HDHD2 | 2865.0815 | -0.2225 | 0.0676 | -3.2920 | 0.000994641 | 0.007956601 |
| C15orf40 | 1927.6109 | -0.2233 | 0.0846 | -2.6393 | 0.008307303 | 0.041528281 |
| FUCA2 | 1456.6396 | -0.2252 | 0.0851 | -2.6463 | 0.008138169 | 0.040899029 |
| SLC38A9 | 1658.1924 | -0.2259 | 0.0792 | -2.8519 | 0.004345591 | 0.025410032 |
| PRNP | 6255.0290 | -0.2265 | 0.0849 | -2.6688 | 0.007612447 | 0.038914969 |
| EIF4EBP2 | 10117.6004 | -0.2267 | 0.0559 | -4.0528 | 5.06E-05 | 0.000719385 |
| GALT | 3145.0151 | -0.2269 | 0.0742 | -3.0595 | 0.0022169 | 0.014953948 |
| CUL4A | 8419.6041 | -0.2270 | 0.0643 | -3.5291 | 0.000416963 | 0.003989496 |
| ZFP90 | 3660.5284 | -0.2272 | 0.0791 | -2.8714 | 0.004086133 | 0.024201618 |
| DCAKD | 2204.9855 | -0.2274 | 0.0670 | -3.3950 | 0.000686345 | 0.005905771 |
| GEMIN8 | 1536.6215 | -0.2291 | 0.0881 | -2.5999 | 0.009324644 | 0.045340369 |
| PCBD2 | 6498.8639 | -0.2302 | 0.0788 | -2.9191 | 0.003510601 | 0.02146518 |
| ELMOD3 | 2684.2731 | -0.2303 | 0.0835 | -2.7591 | 0.005795637 | 0.031638614 |
| DRG2 | 2504.9442 | -0.2304 | 0.0831 | -2.7725 | 0.005562269 | 0.030663429 |
| ULK3 | 4619.1356 | -0.2310 | 0.0810 | -2.8522 | 0.004341492 | 0.025395881 |
| DECR1 | 4341.4012 | -0.2318 | 0.0645 | -3.5950 | 0.000324336 | 0.003259955 |
| PCBP4 | 2436.5308 | -0.2319 | 0.0780 | -2.9720 | 0.002958616 | 0.018836274 |
| MAPK9 | 3096.8007 | -0.2320 | 0.0843 | -2.7532 | 0.005900803 | 0.032036064 |
| PHF15 | 4500.7085 | -0.2326 | 0.0823 | -2.8266 | 0.004704615 | 0.027142149 |
| PASK | 9839.0134 | -0.2327 | 0.0755 | -3.0807 | 0.002065207 | 0.014187279 |
| SNX20 | 6864.1962 | -0.2339 | 0.0867 | -2.6993 | 0.006948704 | 0.036283412 |
| CD96 | 42359.4809 | -0.2340 | 0.0837 | -2.7938 | 0.005209667 | 0.029307039 |
| FAM192A | 8492.6397 | -0.2340 | 0.0685 | -3.4168 | 0.000633691 | 0.005544157 |
| ZNF263 | 2587.7105 | -0.2360 | 0.0739 | -3.1935 | 0.001405762 | 0.010444482 |
| TMC6 | 22195.8127 | -0.2367 | 0.0768 | -3.0836 | 0.002045135 | 0.014071593 |
| KLF12 | 6739.4737 | -0.2374 | 0.0825 | -2.8784 | 0.00399746 | 0.02383439 |
| BTN3A2 | 11914.1250 | -0.2375 | 0.0704 | -3.3748 | 0.000738819 | 0.006282247 |
| AOAH | 9961.9185 | -0.2379 | 0.0777 | -3.0622 | 0.002197175 | 0.014900797 |
| VEZF1 | 3841.7532 | -0.2392 | 0.0666 | -3.5902 | 0.0003304 | 0.003312636 |
| EVL | 57834.2946 | -0.2392 | 0.0649 | -3.6848 | 0.000228915 | 0.00245589 |
| RG9MTD3 | 3480.1447 | -0.2396 | 0.0628 | -3.8136 | 0.000136951 | 0.001632511 |
| OBFC2A | 4677.7421 | -0.2404 | 0.0892 | -2.6957 | 0.007024993 | 0.03660595 |
| CBR4 | 4778.2197 | -0.2405 | 0.0714 | -3.3695 | 0.00075302 | 0.006374336 |
| C18orf8 | 2532.2680 | -0.2405 | 0.0810 | -2.9675 | 0.003002077 | 0.01902489 |
| TMEM185B | 1736.9804 | -0.2407 | 0.0885 | -2.7190 | 0.006548289 | 0.034842056 |
| PLCL2 | 4808.4350 | -0.2412 | 0.0691 | -3.4878 | 0.000487034 | 0.004500527 |
| ZNF266 | 4280.1826 | -0.2413 | 0.0910 | -2.6521 | 0.008000219 | 0.040299473 |
| SNTB2 | 3552.0709 | -0.2416 | 0.0795 | -3.0385 | 0.002377632 | 0.015851229 |
| CMC1 | 2284.0994 | -0.2419 | 0.0661 | -3.6579 | 0.000254294 | 0.002665774 |
| ACBD4 | 2664.7268 | -0.2433 | 0.0794 | -3.0627 | 0.002193613 | 0.014900797 |
| YWHAB | 15428.4781 | -0.2437 | 0.0877 | -2.7807 | 0.005424818 | 0.030180664 |
| NT5DC1 | 3707.0718 | -0.2439 | 0.0725 | -3.3626 | 0.000772004 | 0.006502286 |
| LCK | 30242.3209 | -0.2440 | 0.0854 | -2.8579 | 0.004265088 | 0.025065261 |
| SENP7 | 6501.6967 | -0.2443 | 0.0894 | -2.7325 | 0.006284716 | 0.033676552 |
| HMGB1 | 20935.5405 | -0.2452 | 0.0860 | -2.8491 | 0.004384717 | 0.025569628 |
| MAP4K4 | 5893.9903 | -0.2453 | 0.0806 | -3.0429 | 0.002342981 | 0.015668554 |
| LRRC8A | 1893.7307 | -0.2461 | 0.0722 | -3.4086 | 0.000652986 | 0.005683382 |
| INF2 | 5513.2367 | -0.2463 | 0.0889 | -2.7714 | 0.005582384 | 0.030745258 |
| ITM2B | 13689.6009 | -0.2467 | 0.0709 | -3.4814 | 0.000498721 | 0.004580538 |
| DEF6 | 17178.3064 | -0.2467 | 0.0716 | -3.4471 | 0.000566567 | 0.005077285 |
| GALNT1 | 3069.1944 | -0.2468 | 0.0719 | -3.4302 | 0.000603055 | 0.005309901 |
| ZDHHC3 | 2749.2165 | -0.2469 | 0.0829 | -2.9776 | 0.002905098 | 0.018565871 |
| NOA1 | 3211.0378 | -0.2478 | 0.0734 | -3.3752 | 0.000737709 | 0.006276335 |
| MTAP | 1534.2599 | -0.2480 | 0.0799 | -3.1039 | 0.001909925 | 0.013350941 |
| KLHDC4 | 3057.1793 | -0.2490 | 0.0777 | -3.2071 | 0.001340616 | 0.010064267 |
| SLC35C2 | 4770.7996 | -0.2491 | 0.0750 | -3.3216 | 0.000895189 | 0.007331629 |
| DENND1C | 11470.2325 | -0.2499 | 0.0681 | -3.6664 | 0.000245967 | 0.002591051 |
| COG4 | 5226.0638 | -0.2500 | 0.0937 | -2.6698 | 0.007589406 | 0.038829681 |
| SMC6 | 5558.5348 | -0.2504 | 0.0875 | -2.8605 | 0.00422999 | 0.024907381 |
| AMIGO2 | 1126.1633 | -0.2510 | 0.0947 | -2.6493 | 0.008065244 | 0.040572979 |
| CNPPD1 | 4446.9777 | -0.2512 | 0.0727 | -3.4545 | 0.00055137 | 0.004967585 |
| LOC100506776 | 6567.7725 | -0.2516 | 0.0944 | -2.6651 | 0.0076975 | 0.039179032 |
| LOC401320 | 10062.5478 | -0.2517 | 0.0906 | -2.7776 | 0.005476055 | 0.030365207 |
| TOM1L2 | 4255.3868 | -0.2518 | 0.0961 | -2.6203 | 0.008785848 | 0.043276954 |
| RAB7L1 | 4925.4026 | -0.2523 | 0.0835 | -3.0238 | 0.002496421 | 0.016397463 |
| RNF123 | 2386.0681 | -0.2525 | 0.0969 | -2.6056 | 0.009172483 | 0.044773201 |
| FTX | 6104.8507 | -0.2533 | 0.0707 | -3.5851 | 0.000336913 | 0.003361796 |
| IVD | 2848.1711 | -0.2536 | 0.0956 | -2.6533 | 0.007971421 | 0.040194561 |
| ZNF277 | 2604.2469 | -0.2537 | 0.0942 | -2.6922 | 0.007097897 | 0.036866656 |
| ZBTB22 | 1569.6174 | -0.2537 | 0.0814 | -3.1148 | 0.001840537 | 0.012931633 |
| GCLC | 2198.3235 | -0.2537 | 0.0747 | -3.3972 | 0.000680828 | 0.005869989 |
| TRABD | 12002.0170 | -0.2541 | 0.0913 | -2.7835 | 0.005377336 | 0.030009875 |
| C10orf26 | 3733.5616 | -0.2551 | 0.0871 | -2.9290 | 0.003400467 | 0.02089312 |
| ARHGEF1 | 43893.5486 | -0.2552 | 0.0699 | -3.6532 | 0.000258949 | 0.002707068 |
| PLEKHM1 | 3655.7930 | -0.2558 | 0.0644 | -3.9693 | 7.21E-05 | 0.00096584 |
| CRYL1 | 1174.8410 | -0.2559 | 0.0989 | -2.5869 | 0.009684678 | 0.046750956 |
| NUPL2 | 2663.3309 | -0.2572 | 0.0888 | -2.8956 | 0.00378459 | 0.022790404 |
| TAF12 | 1795.8171 | -0.2573 | 0.0979 | -2.6265 | 0.008627012 | 0.042759111 |
| MED29 | 2874.2267 | -0.2577 | 0.0915 | -2.8158 | 0.00486593 | 0.027878944 |
| CTPS2 | 1058.3037 | -0.2578 | 0.0963 | -2.6769 | 0.007430375 | 0.038296176 |
| BTN3A3 | 7124.6485 | -0.2578 | 0.0787 | -3.2777 | 0.001046728 | 0.008289975 |
| FAM175A | 1406.6527 | -0.2580 | 0.0942 | -2.7389 | 0.006165353 | 0.033166177 |
| C10orf88 | 1189.5495 | -0.2585 | 0.0871 | -2.9678 | 0.002999797 | 0.01902489 |
| LPGAT1 | 3121.7136 | -0.2588 | 0.0989 | -2.6159 | 0.008899867 | 0.043738884 |
| ZNF611 | 1895.7302 | -0.2602 | 0.0977 | -2.6629 | 0.007745981 | 0.039386034 |
| APEH | 4227.7744 | -0.2607 | 0.0892 | -2.9211 | 0.003487868 | 0.021334805 |
| MAST3 | 4904.2299 | -0.2615 | 0.0968 | -2.7001 | 0.006931875 | 0.036283208 |
| C21orf33 | 6755.4022 | -0.2621 | 0.0945 | -2.7738 | 0.005541405 | 0.030615353 |
| C1orf63 | 15808.1225 | -0.2627 | 0.0674 | -3.8951 | 9.82E-05 | 0.001238485 |
| KLHL22 | 2261.5803 | -0.2627 | 0.0794 | -3.3085 | 0.000937827 | 0.007591831 |
| LYRM2 | 1589.2771 | -0.2635 | 0.1011 | -2.6076 | 0.009118108 | 0.044550913 |
| TSPAN3 | 3674.3381 | -0.2636 | 0.0861 | -3.0611 | 0.002205154 | 0.014931675 |
| CYBASC3 | 2291.7181 | -0.2637 | 0.0842 | -3.1307 | 0.001743923 | 0.01239677 |
| STRADA | 5970.6952 | -0.2638 | 0.0790 | -3.3378 | 0.000844404 | 0.006974939 |
| MRPL10 | 6219.1783 | -0.2642 | 0.0987 | -2.6757 | 0.007457352 | 0.038386018 |
| SNRK | 7265.9054 | -0.2642 | 0.0667 | -3.9629 | 7.41E-05 | 0.000989648 |
| HIST1H2BK | 1408.8599 | -0.2644 | 0.1024 | -2.5811 | 0.009848944 | 0.047297531 |
| FAM35A | 3659.7684 | -0.2648 | 0.0802 | -3.3027 | 0.000957579 | 0.007725494 |
| SLC38A6 | 937.6822 | -0.2651 | 0.0970 | -2.7330 | 0.006275984 | 0.033653603 |
| ATG10 | 952.6524 | -0.2667 | 0.0855 | -3.1179 | 0.001821667 | 0.012834818 |
| APEX1 | 5706.0647 | -0.2672 | 0.0980 | -2.7256 | 0.006417717 | 0.034267843 |
| RAP2B | 5894.4428 | -0.2681 | 0.0643 | -4.1669 | 3.09E-05 | 0.00047619 |
| FAM69A | 1445.9281 | -0.2681 | 0.0982 | -2.7289 | 0.006354236 | 0.033976857 |
| TMEM22 | 1332.6988 | -0.2683 | 0.0974 | -2.7539 | 0.005889772 | 0.031990875 |
| MCART6 | 1884.8046 | -0.2685 | 0.0931 | -2.8841 | 0.003925738 | 0.023506755 |
| TDRD3 | 2133.1844 | -0.2687 | 0.0637 | -4.2191 | 2.45E-05 | 0.000392703 |
| ZGPAT | 3074.0285 | -0.2688 | 0.0774 | -3.4748 | 0.000511328 | 0.004682119 |
| CRTAP | 4581.3283 | -0.2689 | 0.1020 | -2.6367 | 0.008371619 | 0.04175321 |
| SLC25A42 | 3308.4844 | -0.2689 | 0.0753 | -3.5712 | 0.000355325 | 0.003503917 |
| C6orf89 | 8510.2720 | -0.2692 | 0.0661 | -4.0733 | 4.64E-05 | 0.000672288 |
| CTSS | 6649.4994 | -0.2692 | 0.0749 | -3.5935 | 0.000326275 | 0.003277264 |
| DOCK11 | 10156.0595 | -0.2696 | 0.0657 | -4.1041 | 4.06E-05 | 0.000600894 |
| DNAJB12 | 1901.4092 | -0.2700 | 0.0784 | -3.4433 | 0.000574695 | 0.005128858 |
| ORMDL1 | 9252.6550 | -0.2704 | 0.0634 | -4.2617 | 2.03E-05 | 0.000337239 |
| GDF11 | 1235.4709 | -0.2706 | 0.1043 | -2.5936 | 0.009498837 | 0.046054139 |
| RTF1 | 5384.6907 | -0.2709 | 0.0841 | -3.2229 | 0.001268836 | 0.009645068 |
| METTL23 | 2421.3808 | -0.2716 | 0.0841 | -3.2288 | 0.001243321 | 0.009513263 |
| HEATR5B | 4852.5128 | -0.2718 | 0.0809 | -3.3578 | 0.000785735 | 0.006599565 |
| C3orf23 | 2519.8285 | -0.2726 | 0.0842 | -3.2384 | 0.001202124 | 0.009257673 |
| QARS | 12552.5480 | -0.2726 | 0.1002 | -2.7199 | 0.006530453 | 0.034759383 |
| IRAK4 | 6478.7025 | -0.2746 | 0.0927 | -2.9635 | 0.003041631 | 0.019203149 |
| SDAD1 | 2312.2593 | -0.2747 | 0.0796 | -3.4506 | 0.000559317 | 0.00503319 |
| DGKZ | 11487.5874 | -0.2749 | 0.0808 | -3.4013 | 0.000670642 | 0.005806982 |
| ARRB1 | 3094.4191 | -0.2750 | 0.0946 | -2.9055 | 0.003666686 | 0.022213041 |
| ESD | 4177.0054 | -0.2750 | 0.0894 | -3.0765 | 0.00209448 | 0.014342782 |
| GLCCI1 | 7501.3150 | -0.2755 | 0.0972 | -2.8351 | 0.004581028 | 0.02652017 |
| ABCC5 | 3800.1956 | -0.2762 | 0.0835 | -3.3065 | 0.000944826 | 0.007642988 |
| FYB | 28662.2429 | -0.2767 | 0.0818 | -3.3822 | 0.000719177 | 0.006149794 |
| TMEM128 | 1710.7422 | -0.2777 | 0.0795 | -3.4931 | 0.000477432 | 0.004433466 |
| TMEM63A | 17586.1420 | -0.2778 | 0.0826 | -3.3639 | 0.000768616 | 0.006477357 |
| CCNY | 4191.5968 | -0.2778 | 0.0785 | -3.5399 | 0.000400209 | 0.003846618 |
| ZC3H4 | 4058.5324 | -0.2779 | 0.1035 | -2.6857 | 0.00723726 | 0.037415594 |
| ARHGEF9 | 3256.7330 | -0.2781 | 0.1050 | -2.6492 | 0.008068538 | 0.040576055 |
| CCDC97 | 3620.5902 | -0.2782 | 0.0792 | -3.5117 | 0.00044518 | 0.004198407 |
| RCAN3 | 3169.8918 | -0.2782 | 0.0834 | -3.3357 | 0.000850829 | 0.007013888 |
| MAN2B2 | 4191.3413 | -0.2783 | 0.0888 | -3.1329 | 0.001730716 | 0.012326052 |
| 9-Sep | 76058.2045 | -0.2787 | 0.0653 | -4.2674 | 1.98E-05 | 0.000330156 |
| CYBRD1 | 1059.0314 | -0.2791 | 0.0953 | -2.9294 | 0.003395829 | 0.020882288 |
| APBB1IP | 9446.3393 | -0.2795 | 0.0886 | -3.1533 | 0.001614118 | 0.011672851 |
| SUMF1 | 1169.4962 | -0.2795 | 0.0929 | -3.0072 | 0.002636747 | 0.017118483 |
| SLC2A4RG | 3404.4815 | -0.2799 | 0.1016 | -2.7555 | 0.005861065 | 0.031910572 |
| FANCF | 1182.8108 | -0.2811 | 0.1035 | -2.7153 | 0.006621822 | 0.035109816 |
| ICAM2 | 2286.4404 | -0.2811 | 0.1056 | -2.6608 | 0.007794487 | 0.039579457 |
| UBLCP1 | 1842.1121 | -0.2811 | 0.0860 | -3.2674 | 0.001085218 | 0.008518802 |
| ABHD14B | 5057.0872 | -0.2813 | 0.1041 | -2.7005 | 0.006923267 | 0.03626325 |
| ZNF195 | 2501.3021 | -0.2814 | 0.1072 | -2.6244 | 0.00868079 | 0.0429272 |
| GLIPR1 | 3974.1515 | -0.2815 | 0.0785 | -3.5874 | 0.00033394 | 0.003334331 |
| WAS | 8785.2926 | -0.2817 | 0.0764 | -3.6899 | 0.000224366 | 0.002424272 |
| ACACB | 1009.2564 | -0.2819 | 0.1005 | -2.8054 | 0.005025916 | 0.028570853 |
| TMEM168 | 3802.5314 | -0.2821 | 0.0961 | -2.9368 | 0.003316338 | 0.020509505 |
| BRD3 | 1779.5477 | -0.2827 | 0.1052 | -2.6866 | 0.007218742 | 0.037358161 |
| KIFC2 | 2545.6659 | -0.2827 | 0.1043 | -2.7109 | 0.006710943 | 0.035482851 |
| C11orf2 | 7935.2406 | -0.2827 | 0.1086 | -2.6030 | 0.009241748 | 0.045039503 |
| ARL2BP | 3108.0605 | -0.2830 | 0.0867 | -3.2631 | 0.001102073 | 0.008606637 |
| MTHFSD | 2167.0892 | -0.2830 | 0.0882 | -3.2077 | 0.001337904 | 0.010048896 |
| ZNF561 | 1156.8514 | -0.2839 | 0.1033 | -2.7491 | 0.005976559 | 0.03234751 |
| FBXO25 | 1232.4118 | -0.2848 | 0.0910 | -3.1290 | 0.001754052 | 0.012457063 |
| UBA7 | 8200.2488 | -0.2850 | 0.0821 | -3.4726 | 0.000515496 | 0.004711727 |
| ZNF506 | 2578.3869 | -0.2864 | 0.0668 | -4.2877 | 1.81E-05 | 0.000307541 |
| CA5B | 1771.4752 | -0.2866 | 0.0868 | -3.3011 | 0.000962983 | 0.007756676 |
| NBPF1 | 1603.8328 | -0.2868 | 0.0948 | -3.0247 | 0.002488966 | 0.016360891 |
| ERV3-1 | 1638.0637 | -0.2876 | 0.1038 | -2.7711 | 0.005586666 | 0.030752026 |
| ATP10A | 5007.9934 | -0.2879 | 0.0658 | -4.3739 | 1.22E-05 | 0.000220616 |
| TTC39C | 3193.4898 | -0.2887 | 0.0924 | -3.1239 | 0.001784569 | 0.012614565 |
| MAP3K1 | 6368.5216 | -0.2887 | 0.0849 | -3.3989 | 0.000676532 | 0.005844607 |
| MRFAP1L1 | 5283.0033 | -0.2890 | 0.0956 | -3.0236 | 0.002498056 | 0.016401082 |
| LOC100506023 | 521.2726 | -0.2891 | 0.1067 | -2.7085 | 0.006758215 | 0.035682904 |
| ST6GALNAC6 | 10942.6052 | -0.2891 | 0.0906 | -3.1915 | 0.001415227 | 0.010504486 |
| RGL2 | 3573.6805 | -0.2902 | 0.0782 | -3.7125 | 0.000205194 | 0.002259079 |
| RPL23AP82 | 661.5099 | -0.2904 | 0.1022 | -2.8411 | 0.004495543 | 0.026115237 |
| LFNG | 2321.3634 | -0.2909 | 0.1081 | -2.6907 | 0.007130272 | 0.036988897 |
| MAP7D1 | 9975.4645 | -0.2911 | 0.0951 | -3.0603 | 0.002211308 | 0.014953948 |
| OSBPL8 | 8834.9119 | -0.2912 | 0.0831 | -3.5047 | 0.000457065 | 0.004289094 |
| SCRN3 | 1258.5328 | -0.2913 | 0.0974 | -2.9912 | 0.002778479 | 0.017892745 |
| CCM2 | 12494.4319 | -0.2917 | 0.1007 | -2.8964 | 0.003774452 | 0.022756533 |
| RNASEH2B | 5026.7046 | -0.2923 | 0.0893 | -3.2747 | 0.001057791 | 0.008360089 |
| TMEM14A | 1403.9064 | -0.2924 | 0.0899 | -3.2510 | 0.001150189 | 0.008913374 |
| ANKRD13D | 8325.3369 | -0.2926 | 0.1022 | -2.8630 | 0.004196971 | 0.024755469 |
| PNPLA2 | 4737.4114 | -0.2943 | 0.0771 | -3.8161 | 0.000135594 | 0.001620162 |
| CUX1 | 1920.7837 | -0.2944 | 0.1023 | -2.8767 | 0.004019013 | 0.023897645 |
| PRKACB | 8243.7876 | -0.2945 | 0.1062 | -2.7720 | 0.005570596 | 0.030698147 |
| SLC44A2 | 12479.0700 | -0.2947 | 0.0620 | -4.7533 | 2.00E-06 | 4.73E-05 |
| LOC254128 | 1453.4363 | -0.2959 | 0.0843 | -3.5088 | 0.000450174 | 0.004237575 |
| DENND2D | 10770.0571 | -0.2959 | 0.0661 | -4.4762 | 7.60E-06 | 0.000148027 |
| LOC100131564 | 5149.2670 | -0.2960 | 0.0982 | -3.0147 | 0.002572564 | 0.016752117 |
| USP20 | 10728.5219 | -0.2962 | 0.0852 | -3.4757 | 0.000509447 | 0.004667715 |
| C3orf75 | 1446.1958 | -0.2964 | 0.1066 | -2.7801 | 0.005434682 | 0.030213317 |
| TMEM106B | 4376.9259 | -0.2966 | 0.0860 | -3.4504 | 0.000559759 | 0.005034172 |
| TOR3A | 2036.4106 | -0.2971 | 0.1053 | -2.8217 | 0.004777129 | 0.027476667 |
| CRTC1 | 1588.0330 | -0.2972 | 0.1019 | -2.9178 | 0.003524709 | 0.021542737 |
| TMEM50B | 2234.0391 | -0.2972 | 0.0907 | -3.2783 | 0.001044202 | 0.008282981 |
| NDRG3 | 4081.1978 | -0.2973 | 0.0690 | -4.3110 | 1.63E-05 | 0.000282613 |
| RASA4 | 871.3190 | -0.2973 | 0.1140 | -2.6088 | 0.009086742 | 0.044455094 |
| ZNF814 | 1753.9052 | -0.2975 | 0.0853 | -3.4876 | 0.000487335 | 0.004500561 |
| WASH2P | 1741.1710 | -0.2976 | 0.0810 | -3.6745 | 0.000238329 | 0.002529971 |
| CCDC69 | 10066.2110 | -0.2978 | 0.0569 | -5.2346 | 1.65E-07 | 5.67E-06 |
| AMPH | 697.3348 | -0.2978 | 0.1103 | -2.7013 | 0.006907021 | 0.036215774 |
| HPS1 | 6975.1368 | -0.2979 | 0.0785 | -3.7937 | 0.000148438 | 0.001727244 |
| CRIPAK | 1238.2840 | -0.2982 | 0.1004 | -2.9691 | 0.002986689 | 0.018983041 |
| DGCR2 | 3571.0457 | -0.2983 | 0.1005 | -2.9676 | 0.003001286 | 0.01902489 |
| NUDT16 | 1638.8490 | -0.2987 | 0.0693 | -4.3086 | 1.64E-05 | 0.000285028 |
| 9-Mar | 2718.7103 | -0.2996 | 0.1092 | -2.7448 | 0.006055368 | 0.032644172 |
| PARP8 | 11075.7275 | -0.2996 | 0.0789 | -3.7956 | 0.000147263 | 0.001717344 |
| ALDH3A2 | 1662.6475 | -0.2997 | 0.0951 | -3.1532 | 0.001615077 | 0.011672851 |
| ESYT1 | 20852.6724 | -0.3006 | 0.0790 | -3.8033 | 0.000142796 | 0.001678377 |
| STARD9 | 3384.4981 | -0.3008 | 0.1164 | -2.5842 | 0.009760335 | 0.047005597 |
| RBMS1 | 4464.2944 | -0.3010 | 0.0697 | -4.3201 | 1.56E-05 | 0.000272386 |
| SLCO3A1 | 2284.8542 | -0.3022 | 0.0900 | -3.3597 | 0.000780377 | 0.006565496 |
| TMC8 | 39886.7012 | -0.3030 | 0.0867 | -3.4956 | 0.000472939 | 0.004397138 |
| COA5 | 2872.4550 | -0.3034 | 0.0795 | -3.8147 | 0.000136364 | 0.001628082 |
| ARSG | 2969.6582 | -0.3047 | 0.1171 | -2.6021 | 0.009264559 | 0.045101427 |
| BICD1 | 753.5074 | -0.3047 | 0.1126 | -2.7054 | 0.006823137 | 0.035926522 |
| TMEM62 | 1640.9032 | -0.3049 | 0.0996 | -3.0627 | 0.00219362 | 0.014900797 |
| CDK5RAP3 | 15624.1225 | -0.3052 | 0.0974 | -3.1339 | 0.001724865 | 0.012290915 |
| GRN | 2429.3497 | -0.3057 | 0.1085 | -2.8169 | 0.004848974 | 0.027815862 |
| C7orf44 | 4337.3862 | -0.3057 | 0.1022 | -2.9921 | 0.002770768 | 0.017865804 |
| C20orf72 | 3751.2651 | -0.3061 | 0.0708 | -4.3206 | 1.56E-05 | 0.000272178 |
| CYP20A1 | 1316.6975 | -0.3063 | 0.0920 | -3.3274 | 0.000876717 | 0.007196333 |
| ORAI3 | 3932.0136 | -0.3063 | 0.1000 | -3.0641 | 0.002183342 | 0.014863824 |
| FAM55C | 2681.7070 | -0.3065 | 0.0795 | -3.8570 | 0.000114781 | 0.001408187 |
| SELPLG | 28406.3592 | -0.3066 | 0.1002 | -3.0591 | 0.002219737 | 0.01496344 |
| RABL3 | 1870.3409 | -0.3070 | 0.0966 | -3.1785 | 0.001480156 | 0.010895531 |
| SIRT7 | 2920.2631 | -0.3075 | 0.1044 | -2.9450 | 0.003229341 | 0.020086451 |
| CROT | 1337.0745 | -0.3075 | 0.0886 | -3.4700 | 0.000520544 | 0.004734982 |
| MAP4K1 | 19736.5798 | -0.3078 | 0.0743 | -4.1428 | 3.43E-05 | 0.000521597 |
| CLDN15 | 2582.7081 | -0.3083 | 0.1065 | -2.8938 | 0.003805783 | 0.022899792 |
| CTSZ | 2789.2795 | -0.3084 | 0.1127 | -2.7368 | 0.006204431 | 0.033352674 |
| PATL2 | 5772.1264 | -0.3096 | 0.0935 | -3.3110 | 0.00092979 | 0.007544791 |
| ZNF276 | 2298.0484 | -0.3097 | 0.1122 | -2.7594 | 0.005791015 | 0.031624798 |
| C19orf12 | 4473.0384 | -0.3098 | 0.0775 | -3.9955 | 6.46E-05 | 0.000883153 |
| NAGK | 2115.8789 | -0.3102 | 0.0972 | -3.1903 | 0.001421354 | 0.010539615 |
| MED12L | 1663.9630 | -0.3104 | 0.0810 | -3.8333 | 0.000126439 | 0.001533791 |
| BEX4 | 947.8791 | -0.3105 | 0.1128 | -2.7532 | 0.005902328 | 0.032036064 |
| C22orf25 | 2486.2460 | -0.3106 | 0.0917 | -3.3862 | 0.000708576 | 0.006069443 |
| PLEK | 1266.5999 | -0.3108 | 0.1085 | -2.8638 | 0.004185685 | 0.024704197 |
| RMND1 | 1956.4842 | -0.3112 | 0.0741 | -4.1992 | 2.68E-05 | 0.000420834 |
| MTERFD3 | 1068.6594 | -0.3117 | 0.1008 | -3.0915 | 0.001991434 | 0.013793231 |
| CDCA7 | 10467.6728 | -0.3123 | 0.1007 | -3.1009 | 0.001929276 | 0.01343654 |
| ADD3 | 24594.1415 | -0.3124 | 0.0982 | -3.1812 | 0.001466493 | 0.010816012 |
| NUDT9 | 693.9130 | -0.3125 | 0.0835 | -3.7432 | 0.000181707 | 0.002051257 |
| PDCD4 | 19241.3459 | -0.3131 | 0.0708 | -4.4233 | 9.72E-06 | 0.000183153 |
| NLRC5 | 19046.3161 | -0.3137 | 0.1137 | -2.7599 | 0.005782532 | 0.031589874 |
| ITGB1BP1 | 4795.0417 | -0.3138 | 0.0661 | -4.7470 | 2.06E-06 | 4.86E-05 |
| ARHGAP9 | 12596.1607 | -0.3140 | 0.0844 | -3.7213 | 0.000198163 | 0.002194442 |
| PROCA1 | 1125.3650 | -0.3145 | 0.1165 | -2.6998 | 0.006937295 | 0.036283412 |
| UBIAD1 | 877.4768 | -0.3146 | 0.1111 | -2.8306 | 0.004645801 | 0.026864306 |
| ATMIN | 3518.6622 | -0.3152 | 0.0858 | -3.6753 | 0.000237589 | 0.002523885 |
| SLC10A3 | 1746.3185 | -0.3154 | 0.0821 | -3.8417 | 0.000122204 | 0.00148719 |
| CCDC53 | 5705.7857 | -0.3158 | 0.1139 | -2.7729 | 0.005555313 | 0.030636243 |
| SBK1 | 1523.9979 | -0.3164 | 0.0940 | -3.3668 | 0.000760435 | 0.006426312 |
| C17orf56 | 1992.7078 | -0.3177 | 0.0877 | -3.6237 | 0.000290445 | 0.002978683 |
| SNAP29 | 2783.8622 | -0.3177 | 0.0913 | -3.4792 | 0.000502886 | 0.004613193 |
| CPT1A | 9827.3555 | -0.3179 | 0.0904 | -3.5159 | 0.000438252 | 0.004140815 |
| C6orf70 | 3162.9441 | -0.3183 | 0.0627 | -5.0743 | 3.89E-07 | 1.17E-05 |
| RABGAP1L | 11857.9597 | -0.3187 | 0.0945 | -3.3714 | 0.000747952 | 0.006334976 |
| RNF214 | 2318.0790 | -0.3193 | 0.0759 | -4.2089 | 2.57E-05 | 0.000406983 |
| DIS3L | 2085.8364 | -0.3196 | 0.0883 | -3.6200 | 0.000294615 | 0.003015317 |
| SIT1 | 1524.1922 | -0.3198 | 0.0909 | -3.5184 | 0.000434218 | 0.004115548 |
| C5orf28 | 1946.6312 | -0.3198 | 0.0933 | -3.4269 | 0.00061055 | 0.005366376 |
| CASP4 | 5220.0100 | -0.3206 | 0.0862 | -3.7190 | 0.000199989 | 0.002208194 |
| CTSK | 999.0582 | -0.3207 | 0.1097 | -2.9242 | 0.003453509 | 0.021150294 |
| ORAI2 | 4835.0998 | -0.3215 | 0.0984 | -3.2670 | 0.00108712 | 0.008525074 |
| ANXA2 | 12832.3394 | -0.3215 | 0.1240 | -2.5923 | 0.009534137 | 0.046210473 |
| ZBTB44 | 8913.0118 | -0.3216 | 0.0842 | -3.8179 | 0.000134601 | 0.001612115 |
| ICA1L | 543.3169 | -0.3224 | 0.1108 | -2.9090 | 0.003625937 | 0.022036781 |
| ZNF581 | 4655.0558 | -0.3225 | 0.0845 | -3.8185 | 0.000134244 | 0.001610391 |
| C8orf76 | 682.4768 | -0.3227 | 0.0936 | -3.4475 | 0.000565866 | 0.005074007 |
| EHD1 | 9050.7776 | -0.3229 | 0.0739 | -4.3692 | 1.25E-05 | 0.000224108 |
| OSTM1 | 981.0050 | -0.3230 | 0.1215 | -2.6578 | 0.007865582 | 0.039806842 |
| SASH3 | 18775.4713 | -0.3233 | 0.0827 | -3.9113 | 9.18E-05 | 0.001172672 |
| MIR548AE2 | 1427.5622 | -0.3238 | 0.1057 | -3.0639 | 0.00218484 | 0.01486733 |
| PRCP | 4101.8931 | -0.3243 | 0.0793 | -4.0916 | 4.28E-05 | 0.000628013 |
| PPOX | 2134.0495 | -0.3244 | 0.1093 | -2.9676 | 0.003001502 | 0.01902489 |
| ITGAL | 84815.8125 | -0.3244 | 0.0816 | -3.9748 | 7.04E-05 | 0.000948689 |
| ASAH1 | 2542.7615 | -0.3245 | 0.1173 | -2.7672 | 0.005653264 | 0.031040623 |
| TTC12 | 1214.2037 | -0.3247 | 0.1159 | -2.8011 | 0.005093365 | 0.028878308 |
| REEP3 | 3501.0075 | -0.3248 | 0.0822 | -3.9533 | 7.71E-05 | 0.001022823 |
| BCL9L | 6700.1566 | -0.3251 | 0.1107 | -2.9366 | 0.003318199 | 0.020512624 |
| LTB4R | 2693.3412 | -0.3253 | 0.0796 | -4.0864 | 4.38E-05 | 0.000639639 |
| TFDP2 | 2484.7670 | -0.3257 | 0.0929 | -3.5066 | 0.000453801 | 0.004266407 |
| HERPUD2 | 5949.6885 | -0.3264 | 0.0642 | -5.0847 | 3.68E-07 | 1.13E-05 |
| ACSF2 | 1513.0262 | -0.3264 | 0.1158 | -2.8182 | 0.004829174 | 0.027733834 |
| SFXN5 | 1306.0230 | -0.3265 | 0.1207 | -2.7052 | 0.00682573 | 0.035926522 |
| ATXN7L1 | 3040.8186 | -0.3266 | 0.1048 | -3.1156 | 0.001835642 | 0.012909231 |
| TASP1 | 804.3883 | -0.3269 | 0.1194 | -2.7387 | 0.006168638 | 0.033172053 |
| POLR3GL | 2157.1951 | -0.3272 | 0.1249 | -2.6187 | 0.008825541 | 0.043415919 |
| ANAPC13 | 1227.3483 | -0.3281 | 0.1215 | -2.7002 | 0.006929296 | 0.036282265 |
| GNPTG | 3636.2435 | -0.3281 | 0.0774 | -4.2372 | 2.26E-05 | 0.000366517 |
| LOC100505894 | 6131.7378 | -0.3281 | 0.0998 | -3.2879 | 0.001009524 | 0.008054364 |
| MYL12A | 24857.7634 | -0.3283 | 0.1115 | -2.9454 | 0.00322489 | 0.020068453 |
| RABL2A | 1026.6467 | -0.3283 | 0.1053 | -3.1180 | 0.001820987 | 0.012834818 |
| BRP44L | 1024.0890 | -0.3288 | 0.0865 | -3.8036 | 0.0001426 | 0.001678371 |
| IL10RA | 33012.3896 | -0.3292 | 0.0652 | -5.0459 | 4.51E-07 | 1.34E-05 |
| STK10 | 30635.3159 | -0.3294 | 0.0552 | -5.9637 | 2.47E-09 | 1.35E-07 |
| MAP2K5 | 1763.1619 | -0.3298 | 0.0868 | -3.8005 | 0.00014439 | 0.001693164 |
| PLD3 | 5762.3444 | -0.3299 | 0.0856 | -3.8536 | 0.000116412 | 0.001423573 |
| TOB1 | 2754.8403 | -0.3301 | 0.0751 | -4.3964 | 1.10E-05 | 0.000202827 |
| RAP1GDS1 | 5480.5718 | -0.3307 | 0.0782 | -4.2280 | 2.36E-05 | 0.000379409 |
| IFT140 | 1598.1679 | -0.3314 | 0.0803 | -4.1276 | 3.66E-05 | 0.000549993 |
| LOC441242 | 1828.4071 | -0.3317 | 0.1141 | -2.9082 | 0.003635445 | 0.02206797 |
| PBXIP1 | 25259.5990 | -0.3321 | 0.0797 | -4.1666 | 3.09E-05 | 0.00047619 |
| RBM15B | 2629.3927 | -0.3321 | 0.1044 | -3.1795 | 0.001475177 | 0.010869461 |
| ABHD13 | 1790.8945 | -0.3321 | 0.0830 | -3.9999 | 6.34E-05 | 0.000871312 |
| SLC25A38 | 4255.9453 | -0.3326 | 0.0857 | -3.8828 | 0.000103262 | 0.001288819 |
| LYPLAL1 | 3016.5429 | -0.3328 | 0.1113 | -2.9893 | 0.002796204 | 0.017991569 |
| NMRAL1 | 2096.8035 | -0.3329 | 0.1154 | -2.8840 | 0.003926857 | 0.023506755 |
| HELQ | 1500.3633 | -0.3329 | 0.1185 | -2.8100 | 0.004954481 | 0.028228415 |
| P2RX4 | 6171.7706 | -0.3339 | 0.1175 | -2.8431 | 0.004468263 | 0.025976717 |
| TYSND1 | 1540.3432 | -0.3341 | 0.1148 | -2.9099 | 0.003615577 | 0.021982649 |
| DOK1 | 1709.3989 | -0.3343 | 0.1265 | -2.6422 | 0.00823649 | 0.041297113 |
| PPP4R1 | 2248.1051 | -0.3343 | 0.1067 | -3.1339 | 0.001724969 | 0.012290915 |
| MUL1 | 1147.7798 | -0.3365 | 0.0939 | -3.5825 | 0.000340307 | 0.003393423 |
| BTBD11 | 2556.1443 | -0.3376 | 0.1203 | -2.8054 | 0.005025792 | 0.028570853 |
| NIPSNAP3A | 712.3824 | -0.3379 | 0.1252 | -2.6995 | 0.006943525 | 0.036283412 |
| ZNF652 | 3325.6487 | -0.3382 | 0.0809 | -4.1802 | 2.91E-05 | 0.000452793 |
| LYSMD4 | 875.9325 | -0.3382 | 0.1216 | -2.7804 | 0.005428567 | 0.030190415 |
| TMEM218 | 719.4183 | -0.3383 | 0.1102 | -3.0703 | 0.002138357 | 0.014590406 |
| ATG16L1 | 5741.0164 | -0.3386 | 0.0888 | -3.8127 | 0.000137449 | 0.001637154 |
| TP53I13 | 3685.6499 | -0.3388 | 0.1089 | -3.1124 | 0.00185588 | 0.013015249 |
| CD44 | 24766.8088 | -0.3388 | 0.0985 | -3.4384 | 0.000585068 | 0.005208963 |
| PCSK7 | 10583.9879 | -0.3391 | 0.0834 | -4.0683 | 4.74E-05 | 0.000683179 |
| SH2D3A | 1177.2068 | -0.3395 | 0.0949 | -3.5786 | 0.000345402 | 0.003430708 |
| PIGM | 1104.4696 | -0.3397 | 0.0875 | -3.8832 | 0.00010307 | 0.001287491 |
| GOLM1 | 1144.3284 | -0.3398 | 0.0903 | -3.7631 | 0.000167787 | 0.00190836 |
| ZNF226 | 1516.6333 | -0.3405 | 0.0928 | -3.6686 | 0.000243886 | 0.0025763 |
| CROCCP3 | 944.5070 | -0.3411 | 0.1063 | -3.2104 | 0.001325497 | 0.009985458 |
| CTDSP2 | 16433.4678 | -0.3412 | 0.0938 | -3.6357 | 0.000277207 | 0.002870169 |
| LOC100286793 | 1211.2200 | -0.3413 | 0.1223 | -2.7916 | 0.005245135 | 0.029451801 |
| ARHGAP25 | 6206.4362 | -0.3416 | 0.0833 | -4.1012 | 4.11E-05 | 0.000606014 |
| CD300A | 5697.9484 | -0.3416 | 0.1089 | -3.1384 | 0.001698483 | 0.012159467 |
| NUDT5 | 3746.1927 | -0.3416 | 0.0855 | -3.9969 | 6.42E-05 | 0.000880143 |
| DAGLB | 1386.3935 | -0.3417 | 0.1124 | -3.0391 | 0.002372765 | 0.015832737 |
| STYX | 3030.2846 | -0.3419 | 0.0856 | -3.9960 | 6.44E-05 | 0.000881794 |
| LOC100132273 | 730.4827 | -0.3420 | 0.1108 | -3.0864 | 0.002025751 | 0.013977466 |
| CHMP7 | 4069.4532 | -0.3429 | 0.0792 | -4.3307 | 1.49E-05 | 0.000260658 |
| RASSF2 | 9101.8395 | -0.3432 | 0.0805 | -4.2637 | 2.01E-05 | 0.00033468 |
| NLRP1 | 25931.7720 | -0.3433 | 0.0833 | -4.1193 | 3.80E-05 | 0.00056746 |
| CCDC91 | 1571.6234 | -0.3434 | 0.0909 | -3.7756 | 0.000159605 | 0.001834613 |
| LITAF | 12831.6416 | -0.3436 | 0.0977 | -3.5180 | 0.000434795 | 0.004118437 |
| ZNF605 | 2039.4626 | -0.3441 | 0.0973 | -3.5362 | 0.000405867 | 0.003895657 |
| PCOLCE | 882.0120 | -0.3442 | 0.1298 | -2.6527 | 0.007984977 | 0.040249498 |
| FKBP5 | 5666.9936 | -0.3443 | 0.0901 | -3.8195 | 0.000133711 | 0.001607887 |
| TSHZ1 | 2404.6882 | -0.3449 | 0.1218 | -2.8305 | 0.00464765 | 0.026864732 |
| C7orf31 | 543.4762 | -0.3450 | 0.1263 | -2.7323 | 0.006290122 | 0.033681656 |
| FBXL17 | 2193.4994 | -0.3451 | 0.0764 | -4.5171 | 6.27E-06 | 0.000126935 |
| MLANA | 879.7645 | -0.3455 | 0.1116 | -3.0961 | 0.00196065 | 0.01360585 |
| TES | 7482.5550 | -0.3469 | 0.0725 | -4.7856 | 1.70E-06 | 4.13E-05 |
| APLF | 716.9633 | -0.3470 | 0.1185 | -2.9293 | 0.003396963 | 0.020882288 |
| C11orf67 | 942.2325 | -0.3470 | 0.1282 | -2.7064 | 0.006801725 | 0.035862563 |
| EIF4E3 | 7185.4162 | -0.3471 | 0.0766 | -4.5299 | 5.90E-06 | 0.000120785 |
| ARL16 | 1127.8027 | -0.3473 | 0.1324 | -2.6237 | 0.008697061 | 0.042993609 |
| KIAA1671 | 3954.9346 | -0.3486 | 0.0764 | -4.5607 | 5.10E-06 | 0.000107136 |
| DICER1-AS | 387.3068 | -0.3487 | 0.1279 | -2.7263 | 0.00640398 | 0.034218653 |
| ATP2A3 | 18444.8729 | -0.3491 | 0.0740 | -4.7164 | 2.40E-06 | 5.55E-05 |
| TNFSF13B | 1132.4386 | -0.3493 | 0.1286 | -2.7174 | 0.006579101 | 0.034944546 |
| FLI1 | 11664.1174 | -0.3495 | 0.0878 | -3.9832 | 6.80E-05 | 0.000919321 |
| PGLS | 2331.3810 | -0.3496 | 0.1317 | -2.6550 | 0.007930739 | 0.040096354 |
| PLP2 | 6997.9929 | -0.3496 | 0.1154 | -3.0303 | 0.002442839 | 0.01616951 |
| ARMC6 | 916.2683 | -0.3497 | 0.1355 | -2.5798 | 0.009885263 | 0.047411023 |
| UCP2 | 14918.3814 | -0.3497 | 0.1357 | -2.5781 | 0.009934748 | 0.047633259 |
| EMB | 25211.3192 | -0.3497 | 0.1029 | -3.3971 | 0.000681023 | 0.005869989 |
| PLCB2 | 13248.3279 | -0.3498 | 0.0615 | -5.6866 | 1.30E-08 | 6.09E-07 |
| VAT1 | 2066.6566 | -0.3500 | 0.0943 | -3.7098 | 0.000207383 | 0.002275857 |
| RWDD2B | 600.3551 | -0.3509 | 0.1360 | -2.5812 | 0.009846999 | 0.047297531 |
| HLA-DOA | 1365.1857 | -0.3517 | 0.0942 | -3.7318 | 0.00019009 | 0.002126842 |
| JUP | 348.8444 | -0.3518 | 0.1300 | -2.7059 | 0.006811286 | 0.035887956 |
| RG9MTD2 | 737.2504 | -0.3518 | 0.1361 | -2.5853 | 0.009728791 | 0.046898478 |
| VIM | 130891.5862 | -0.3521 | 0.1270 | -2.7730 | 0.005553678 | 0.030636243 |
| PYHIN1 | 5306.9052 | -0.3522 | 0.0678 | -5.1951 | 2.05E-07 | 6.74E-06 |
| ARL2-SNX15 | 977.0407 | -0.3527 | 0.1302 | -2.7098 | 0.006731702 | 0.035567747 |
| PDP1 | 2133.1733 | -0.3527 | 0.0947 | -3.7246 | 0.000195594 | 0.002175547 |
| LOC202781 | 689.0481 | -0.3529 | 0.1284 | -2.7491 | 0.005976816 | 0.03234751 |
| PAFAH2 | 1018.6594 | -0.3530 | 0.1122 | -3.1465 | 0.001652472 | 0.01188402 |
| ZADH2 | 3491.5494 | -0.3533 | 0.0760 | -4.6493 | 3.33E-06 | 7.34E-05 |
| HDHD1 | 864.2882 | -0.3534 | 0.1307 | -2.7034 | 0.006863307 | 0.036049044 |
| FAR2 | 704.0789 | -0.3534 | 0.1004 | -3.5198 | 0.000431843 | 0.004098175 |
| C17orf57 | 2569.9424 | -0.3534 | 0.1101 | -3.2104 | 0.001325432 | 0.009985458 |
| FLJ33630 | 1338.2352 | -0.3535 | 0.0821 | -4.3041 | 1.68E-05 | 0.000289624 |
| MGAT4A | 9372.2204 | -0.3536 | 0.0847 | -4.1740 | 2.99E-05 | 0.000463885 |
| CLYBL | 995.3682 | -0.3539 | 0.1040 | -3.4032 | 0.000666129 | 0.005783004 |
| PCBP1-AS1 | 5375.3635 | -0.3541 | 0.0890 | -3.9783 | 6.94E-05 | 0.000935896 |
| AMICA1 | 33952.6512 | -0.3541 | 0.0981 | -3.6110 | 0.000305072 | 0.00310345 |
| RPP38 | 1123.5550 | -0.3545 | 0.1290 | -2.7476 | 0.006003649 | 0.032457901 |
| GAS2L1 | 1166.7918 | -0.3546 | 0.1329 | -2.6686 | 0.007617758 | 0.038917197 |
| SWT1 | 1273.6370 | -0.3547 | 0.1044 | -3.3991 | 0.000676009 | 0.005843421 |
| NCOA5 | 2867.0893 | -0.3547 | 0.1018 | -3.4849 | 0.000492322 | 0.004530018 |
| CAPN2 | 25990.8287 | -0.3548 | 0.0538 | -6.5905 | 4.38E-11 | 3.49E-09 |
| PEX16 | 1853.5790 | -0.3553 | 0.1322 | -2.6874 | 0.007200303 | 0.037275489 |
| TRIM3 | 641.5756 | -0.3556 | 0.0921 | -3.8609 | 0.000112986 | 0.001391043 |
| C15orf33 | 1771.1011 | -0.3556 | 0.1027 | -3.4623 | 0.000535586 | 0.004850232 |
| LSS | 5496.4677 | -0.3558 | 0.0731 | -4.8647 | 1.15E-06 | 2.95E-05 |
| ULK2 | 1008.6691 | -0.3564 | 0.0996 | -3.5786 | 0.000345407 | 0.003430708 |
| NIN | 10084.0180 | -0.3570 | 0.0921 | -3.8755 | 0.000106404 | 0.001320407 |
| TBCC | 2138.3593 | -0.3588 | 0.0943 | -3.8043 | 0.000142187 | 0.001677735 |
| ARHGAP1 | 5884.3969 | -0.3588 | 0.0697 | -5.1454 | 2.67E-07 | 8.61E-06 |
| ZC4H2 | 800.8503 | -0.3595 | 0.0977 | -3.6781 | 0.000234953 | 0.002504679 |
| LMF1 | 3444.2593 | -0.3598 | 0.1075 | -3.3452 | 0.000822229 | 0.006830237 |
| NOL9 | 1591.4683 | -0.3603 | 0.1088 | -3.3108 | 0.000930193 | 0.007544791 |
| C8orf38 | 2211.5188 | -0.3609 | 0.0859 | -4.2016 | 2.65E-05 | 0.000417252 |
| CALML4 | 1668.7518 | -0.3611 | 0.0948 | -3.8077 | 0.000140291 | 0.001665764 |
| FAM113B | 11799.1891 | -0.3611 | 0.0882 | -4.0923 | 4.27E-05 | 0.000626579 |
| PPP1R3F | 1792.0067 | -0.3613 | 0.0894 | -4.0398 | 5.35E-05 | 0.000757099 |
| DOPEY2 | 1196.7362 | -0.3615 | 0.1299 | -2.7836 | 0.005375104 | 0.030009875 |
| GBAP1 | 2030.3689 | -0.3615 | 0.0936 | -3.8608 | 0.000113016 | 0.001391043 |
| ABHD15 | 2386.3891 | -0.3617 | 0.1053 | -3.4361 | 0.000590096 | 0.005232345 |
| PLCD1 | 3751.5539 | -0.3618 | 0.0750 | -4.8209 | 1.43E-06 | 3.56E-05 |
| ZNF710 | 1631.9643 | -0.3625 | 0.1270 | -2.8544 | 0.00431175 | 0.025270767 |
| CCDC65 | 662.3123 | -0.3630 | 0.1298 | -2.7958 | 0.005176497 | 0.029196448 |
| ATP8B2 | 14925.3360 | -0.3636 | 0.1114 | -3.2625 | 0.001104339 | 0.008615438 |
| BOLA2 | 4169.0592 | -0.3636 | 0.1316 | -2.7635 | 0.005718195 | 0.0312974 |
| SERPINF1 | 598.4367 | -0.3636 | 0.1398 | -2.6019 | 0.009272101 | 0.045101427 |
| TMEM175 | 920.9988 | -0.3639 | 0.1203 | -3.0257 | 0.002480516 | 0.01631692 |
| FBXO10 | 788.2718 | -0.3643 | 0.1128 | -3.2280 | 0.001246527 | 0.009523343 |
| ZNF449 | 1044.1045 | -0.3645 | 0.1349 | -2.7025 | 0.006881306 | 0.036118499 |
| WDR77 | 1003.2106 | -0.3646 | 0.1265 | -2.8811 | 0.003962912 | 0.023694453 |
| SYNE1 | 10275.2989 | -0.3647 | 0.1107 | -3.2933 | 0.000990033 | 0.007936348 |
| AGTRAP | 986.2453 | -0.3652 | 0.1297 | -2.8157 | 0.004867343 | 0.027878944 |
| KIAA1009 | 1022.9525 | -0.3653 | 0.1284 | -2.8462 | 0.004423772 | 0.025747751 |
| ZNF585B | 713.3342 | -0.3654 | 0.1160 | -3.1511 | 0.001626642 | 0.011734006 |
| ADAM22 | 693.4827 | -0.3657 | 0.1091 | -3.3519 | 0.000802592 | 0.006703924 |
| INPP5D | 23456.1102 | -0.3665 | 0.0659 | -5.5591 | 2.71E-08 | 1.19E-06 |
| CREB3L4 | 732.6227 | -0.3665 | 0.1392 | -2.6335 | 0.008450013 | 0.042011869 |
| ZNF490 | 932.6119 | -0.3676 | 0.1201 | -3.0616 | 0.002201621 | 0.014914427 |
| CALM1 | 25984.3167 | -0.3677 | 0.0858 | -4.2852 | 1.83E-05 | 0.000310337 |
| TRAF3IP3 | 29019.5758 | -0.3680 | 0.0679 | -5.4156 | 6.11E-08 | 2.38E-06 |
| SUSD3 | 4098.6109 | -0.3684 | 0.0766 | -4.8061 | 1.54E-06 | 3.79E-05 |
| EIF5A2 | 381.2255 | -0.3685 | 0.1267 | -2.9076 | 0.003642518 | 0.022102031 |
| NAGPA | 800.0670 | -0.3688 | 0.1327 | -2.7793 | 0.005447335 | 0.030252984 |
| RTN2 | 541.4766 | -0.3689 | 0.1426 | -2.5876 | 0.009664342 | 0.046691951 |
| KIAA0513 | 985.1907 | -0.3691 | 0.1156 | -3.1935 | 0.001405357 | 0.010444482 |
| TEX22 | 285.4957 | -0.3696 | 0.1405 | -2.6295 | 0.008550599 | 0.042449922 |
| C14orf93 | 1067.0259 | -0.3697 | 0.1008 | -3.6687 | 0.000243769 | 0.0025763 |
| PUS1 | 1066.5185 | -0.3698 | 0.1208 | -3.0620 | 0.002198624 | 0.014900797 |
| GPSM3 | 19651.1354 | -0.3711 | 0.1070 | -3.4692 | 0.000521941 | 0.004744829 |
| GPR155 | 5951.9805 | -0.3713 | 0.1006 | -3.6920 | 0.00022246 | 0.002407124 |
| FAM84B | 1657.0209 | -0.3714 | 0.1270 | -2.9256 | 0.003437645 | 0.021071295 |
| DCLRE1C | 2699.4633 | -0.3715 | 0.0737 | -5.0423 | 4.60E-07 | 1.36E-05 |
| CSPP1 | 1287.4564 | -0.3715 | 0.0873 | -4.2562 | 2.08E-05 | 0.000344191 |
| HSH2D | 8539.2153 | -0.3724 | 0.0920 | -4.0458 | 5.21E-05 | 0.000739332 |
| SLC27A1 | 1190.0296 | -0.3724 | 0.1141 | -3.2632 | 0.001101564 | 0.008606637 |
| ANXA5 | 15159.3291 | -0.3726 | 0.0807 | -4.6142 | 3.95E-06 | 8.51E-05 |
| SAP30L | 2187.2191 | -0.3727 | 0.0694 | -5.3662 | 8.04E-08 | 3.06E-06 |
| TCEAL1 | 362.8032 | -0.3729 | 0.1164 | -3.2041 | 0.001355035 | 0.010142311 |
| PEX6 | 3777.5722 | -0.3729 | 0.0879 | -4.2414 | 2.22E-05 | 0.000361266 |
| CENPBD1 | 477.2261 | -0.3731 | 0.1426 | -2.6165 | 0.00888448 | 0.043677457 |
| RPUSD2 | 395.7239 | -0.3732 | 0.1432 | -2.6068 | 0.009139605 | 0.04463789 |
| FBXL13 | 423.6388 | -0.3735 | 0.1337 | -2.7939 | 0.00520764 | 0.029306536 |
| PQLC2 | 625.5268 | -0.3736 | 0.1328 | -2.8127 | 0.004912468 | 0.028035971 |
| PTPN18 | 7564.8778 | -0.3739 | 0.0972 | -3.8464 | 0.000119874 | 0.001461184 |
| MFNG | 10378.5151 | -0.3746 | 0.1227 | -3.0540 | 0.002258405 | 0.01519026 |
| SRSF8 | 2599.6735 | -0.3748 | 0.0929 | -4.0335 | 5.50E-05 | 0.000773307 |
| ZNF83 | 4553.0982 | -0.3748 | 0.1072 | -3.4951 | 0.00047394 | 0.004403742 |
| TMPRSS13 | 631.4563 | -0.3754 | 0.1298 | -2.8922 | 0.003826042 | 0.022994255 |
| VCL | 8039.5589 | -0.3760 | 0.1048 | -3.5879 | 0.000333343 | 0.003330568 |
| ZSCAN5A | 506.7180 | -0.3764 | 0.1419 | -2.6533 | 0.007970561 | 0.040194561 |
| LINC00094 | 1030.8791 | -0.3771 | 0.1386 | -2.7207 | 0.006513497 | 0.034717996 |
| ZNF493 | 1758.3628 | -0.3772 | 0.1209 | -3.1186 | 0.001817123 | 0.012814741 |
| LOC339290 | 2182.7772 | -0.3778 | 0.0775 | -4.8761 | 1.08E-06 | 2.81E-05 |
| SNHG13 | 667.1646 | -0.3779 | 0.1443 | -2.6182 | 0.008838629 | 0.043466169 |
| TTYH2 | 3378.8834 | -0.3786 | 0.1299 | -2.9141 | 0.003567498 | 0.02175153 |
| PGBD1 | 285.4937 | -0.3799 | 0.1457 | -2.6084 | 0.00909628 | 0.04447299 |
| MGC72080 | 3930.4509 | -0.3803 | 0.1095 | -3.4711 | 0.000518308 | 0.004720314 |
| ST8SIA6 | 763.7688 | -0.3804 | 0.1322 | -2.8780 | 0.004002073 | 0.02383439 |
| C2orf88 | 541.1289 | -0.3804 | 0.1376 | -2.7641 | 0.005707414 | 0.031290393 |
| TMEM164 | 2563.4077 | -0.3805 | 0.0929 | -4.0948 | 4.22E-05 | 0.000621027 |
| NAPEPLD | 1030.3885 | -0.3806 | 0.1317 | -2.8907 | 0.003843243 | 0.023051837 |
| ZNF721 | 654.2316 | -0.3811 | 0.1234 | -3.0877 | 0.002017141 | 0.013939375 |
| FLOT2 | 12245.9085 | -0.3821 | 0.0843 | -4.5330 | 5.82E-06 | 0.000120269 |
| C1orf21 | 1010.4410 | -0.3822 | 0.1416 | -2.6997 | 0.006940314 | 0.036283412 |
| SLC35B3 | 2804.1985 | -0.3823 | 0.0837 | -4.5686 | 4.91E-06 | 0.000103737 |
| VPS33A | 3342.2067 | -0.3824 | 0.0799 | -4.7870 | 1.69E-06 | 4.11E-05 |
| AKD1 | 1133.9025 | -0.3824 | 0.1169 | -3.2727 | 0.001065407 | 0.008402721 |
| CYHR1 | 4196.4306 | -0.3827 | 0.0928 | -4.1233 | 3.74E-05 | 0.000558311 |
| RTKN2 | 2935.8327 | -0.3831 | 0.1381 | -2.7741 | 0.005535093 | 0.030605504 |
| CPNE8 | 760.1231 | -0.3837 | 0.1294 | -2.9659 | 0.003017641 | 0.019107515 |
| GLTPD1 | 1151.2642 | -0.3838 | 0.1231 | -3.1163 | 0.001831477 | 0.012887534 |
| NR1H3 | 926.2826 | -0.3840 | 0.1336 | -2.8737 | 0.004056405 | 0.024063228 |
| IL11RA | 6421.5242 | -0.3841 | 0.0893 | -4.3009 | 1.70E-05 | 0.000292806 |
| NLRC3 | 22284.4078 | -0.3844 | 0.1036 | -3.7097 | 0.000207471 | 0.002275857 |
| GSN | 814.4614 | -0.3852 | 0.0984 | -3.9142 | 9.07E-05 | 0.001161761 |
| VPS18 | 1053.1950 | -0.3853 | 0.1311 | -2.9394 | 0.003288918 | 0.020389942 |
| APH1B | 1782.6098 | -0.3856 | 0.1061 | -3.6358 | 0.000277136 | 0.002870169 |
| SOCS6 | 314.0807 | -0.3857 | 0.1311 | -2.9414 | 0.003267318 | 0.020287481 |
| LOC100507501 | 529.3506 | -0.3858 | 0.1469 | -2.6265 | 0.00862647 | 0.042759111 |
| CAMK2G | 7018.7085 | -0.3859 | 0.0873 | -4.4229 | 9.74E-06 | 0.000183205 |
| RNF125 | 6659.4027 | -0.3864 | 0.0879 | -4.3958 | 1.10E-05 | 0.000203059 |
| SEC22C | 5669.1602 | -0.3868 | 0.0867 | -4.4604 | 8.18E-06 | 0.00015788 |
| DOCK9 | 6829.7952 | -0.3869 | 0.1005 | -3.8490 | 0.000118577 | 0.001447716 |
| ATF7IP2 | 2799.3173 | -0.3869 | 0.0964 | -4.0155 | 5.93E-05 | 0.000825224 |
| TNRC6C | 1989.7197 | -0.3869 | 0.0950 | -4.0741 | 4.62E-05 | 0.000670571 |
| TMEM203 | 1213.1037 | -0.3882 | 0.1389 | -2.7944 | 0.005200225 | 0.029286599 |
| LOC100289019 | 896.4744 | -0.3883 | 0.1244 | -3.1226 | 0.001792699 | 0.012666116 |
| CYB5D2 | 1571.5219 | -0.3893 | 0.1207 | -3.2261 | 0.001255026 | 0.009564122 |
| ACSS1 | 7922.8006 | -0.3895 | 0.0749 | -5.2004 | 1.99E-07 | 6.60E-06 |
| VILL | 1714.4318 | -0.3896 | 0.0929 | -4.1931 | 2.75E-05 | 0.000431385 |
| TSEN54 | 3009.6881 | -0.3901 | 0.1054 | -3.7022 | 0.000213718 | 0.00233423 |
| C12orf35 | 16564.9565 | -0.3903 | 0.1197 | -3.2591 | 0.001117768 | 0.008702256 |
| SAMD3 | 4649.7974 | -0.3906 | 0.1031 | -3.7887 | 0.000151444 | 0.001752789 |
| SLC43A1 | 1909.2496 | -0.3915 | 0.1128 | -3.4714 | 0.000517806 | 0.004720314 |
| CCND3 | 14016.2347 | -0.3922 | 0.1339 | -2.9293 | 0.003397323 | 0.020882288 |
| LOC100506334 | 5109.8887 | -0.3924 | 0.1193 | -3.2904 | 0.001000509 | 0.007995088 |
| CCDC30 | 509.1019 | -0.3927 | 0.1189 | -3.3030 | 0.000956506 | 0.007720955 |
| SLC24A1 | 1454.4978 | -0.3933 | 0.1315 | -2.9909 | 0.00278131 | 0.017903352 |
| SLC22A18 | 1552.1308 | -0.3933 | 0.1103 | -3.5653 | 0.000363419 | 0.003555908 |
| PRB4 | 671.5738 | -0.3938 | 0.1207 | -3.2623 | 0.001105012 | 0.008616243 |
| PARVG | 5241.0511 | -0.3942 | 0.0912 | -4.3218 | 1.55E-05 | 0.000270995 |
| MYOM1 | 918.7616 | -0.3950 | 0.1224 | -3.2261 | 0.001254995 | 0.009564122 |
| CCR6 | 1049.0083 | -0.3956 | 0.1042 | -3.7955 | 0.00014736 | 0.001717344 |
| KIAA0240 | 6747.6601 | -0.3959 | 0.0941 | -4.2076 | 2.58E-05 | 0.000408897 |
| SLAMF6 | 7740.5823 | -0.3960 | 0.1085 | -3.6482 | 0.000264088 | 0.002753172 |
| FUZ | 1134.1956 | -0.3960 | 0.1036 | -3.8215 | 0.000132622 | 0.001598549 |
| C17orf59 | 1290.6717 | -0.3962 | 0.1492 | -2.6563 | 0.00789927 | 0.039958211 |
| RNU5E-1 | 336.9440 | -0.3962 | 0.1281 | -3.0942 | 0.00197351 | 0.013687889 |
| SLC27A5 | 566.2695 | -0.3965 | 0.1422 | -2.7886 | 0.005293574 | 0.029677811 |
| FAM134B | 1317.0178 | -0.3966 | 0.1187 | -3.3423 | 0.000830749 | 0.006878239 |
| DUSP28 | 546.4226 | -0.3971 | 0.1431 | -2.7750 | 0.005519616 | 0.030561941 |
| UPF3A | 5444.8370 | -0.3979 | 0.0710 | -5.6012 | 2.13E-08 | 9.67E-07 |
| C8orf40 | 1404.3329 | -0.3984 | 0.1014 | -3.9282 | 8.56E-05 | 0.001105451 |
| LOC253039 | 4006.4608 | -0.3986 | 0.1012 | -3.9370 | 8.25E-05 | 0.001075654 |
| ENG | 1074.7507 | -0.3989 | 0.1022 | -3.9025 | 9.52E-05 | 0.001212265 |
| LOC148413 | 803.7786 | -0.3991 | 0.1295 | -3.0813 | 0.00206089 | 0.014164052 |
| WIPI1 | 6060.5950 | -0.3993 | 0.1069 | -3.7353 | 0.000187526 | 0.002102819 |
| PROX2 | 424.5255 | -0.3996 | 0.1404 | -2.8470 | 0.004413451 | 0.025707461 |
| MGLL | 539.5238 | -0.3998 | 0.1470 | -2.7204 | 0.006519742 | 0.034723912 |
| TATDN3 | 1467.8874 | -0.4004 | 0.1063 | -3.7649 | 0.000166619 | 0.001897751 |
| MTERFD2 | 2972.2797 | -0.4014 | 0.0805 | -4.9855 | 6.18E-07 | 1.77E-05 |
| SAYSD1 | 1292.2692 | -0.4014 | 0.0884 | -4.5383 | 5.67E-06 | 0.000117809 |
| AGPAT4 | 1797.1575 | -0.4015 | 0.1156 | -3.4726 | 0.000515382 | 0.004711727 |
| RIPK1 | 4114.3488 | -0.4016 | 0.0677 | -5.9362 | 2.92E-09 | 1.56E-07 |
| RNPEPL1 | 11053.2644 | -0.4017 | 0.1032 | -3.8910 | 9.99E-05 | 0.001253478 |
| INPP5K | 3605.0593 | -0.4018 | 0.0926 | -4.3379 | 1.44E-05 | 0.000253077 |
| APBB1 | 4226.7898 | -0.4019 | 0.0822 | -4.8874 | 1.02E-06 | 2.69E-05 |
| ANKRD19P | 948.2444 | -0.4029 | 0.1195 | -3.3715 | 0.000747499 | 0.006334689 |
| PNP | 3469.2916 | -0.4030 | 0.1173 | -3.4341 | 0.000594528 | 0.005259309 |
| CEP19 | 748.8403 | -0.4035 | 0.1197 | -3.3721 | 0.000746071 | 0.006326129 |
| CDK5 | 822.1679 | -0.4039 | 0.1333 | -3.0290 | 0.002453857 | 0.016216466 |
| CCDC92 | 3244.4036 | -0.4044 | 0.0810 | -4.9927 | 5.95E-07 | 1.72E-05 |
| PIGV | 1101.2913 | -0.4044 | 0.0753 | -5.3689 | 7.92E-08 | 3.02E-06 |
| MCRS1 | 2766.7534 | -0.4053 | 0.1122 | -3.6128 | 0.00030289 | 0.003085012 |
| WDR53 | 1101.5410 | -0.4056 | 0.0955 | -4.2465 | 2.17E-05 | 0.000354689 |
| ZSCAN21 | 517.1378 | -0.4061 | 0.1533 | -2.6496 | 0.008058202 | 0.040551038 |
| ANXA1 | 17735.5506 | -0.4063 | 0.0819 | -4.9588 | 7.09E-07 | 1.97E-05 |
| B3GALT4 | 783.5726 | -0.4065 | 0.1506 | -2.6987 | 0.006960047 | 0.036330099 |
| ZNF773 | 545.0073 | -0.4069 | 0.1277 | -3.1853 | 0.001446126 | 0.010707563 |
| EFCAB2 | 397.0535 | -0.4073 | 0.1215 | -3.3522 | 0.000801711 | 0.006700268 |
| MXD4 | 3454.2547 | -0.4079 | 0.0835 | -4.8843 | 1.04E-06 | 2.72E-05 |
| LOC387723 | 365.1698 | -0.4082 | 0.1308 | -3.1220 | 0.001796054 | 0.012677979 |
| LPAR5 | 896.8829 | -0.4091 | 0.1169 | -3.4998 | 0.000465618 | 0.004347778 |
| PPP2R5C | 29463.2913 | -0.4096 | 0.0737 | -5.5602 | 2.69E-08 | 1.18E-06 |
| GALNT7 | 1846.8691 | -0.4096 | 0.1217 | -3.3655 | 0.000763956 | 0.00644167 |
| NINJ2 | 654.7555 | -0.4102 | 0.1282 | -3.2004 | 0.001372339 | 0.010236378 |
| ST6GAL1 | 11321.2108 | -0.4104 | 0.0570 | -7.1970 | 6.16E-13 | 7.11E-11 |
| PXMP4 | 929.2431 | -0.4104 | 0.1197 | -3.4277 | 0.00060864 | 0.005355965 |
| PHOSPHO2-KLHL23 | 720.7130 | -0.4104 | 0.1521 | -2.6985 | 0.006965654 | 0.036346824 |
| HENMT1 | 1660.5780 | -0.4112 | 0.1126 | -3.6533 | 0.000258882 | 0.002707068 |
| TTC13 | 3345.4372 | -0.4114 | 0.0837 | -4.9164 | 8.81E-07 | 2.38E-05 |
| CCDC109B | 5275.5090 | -0.4122 | 0.1371 | -3.0068 | 0.002640072 | 0.01713272 |
| UBTF | 6599.2793 | -0.4124 | 0.0937 | -4.4011 | 1.08E-05 | 0.000199139 |
| HS1BP3 | 1353.4062 | -0.4125 | 0.0993 | -4.1524 | 3.29E-05 | 0.000502618 |
| LINC00324 | 675.7925 | -0.4128 | 0.1425 | -2.8960 | 0.003779792 | 0.022779644 |
| PEPD | 896.9160 | -0.4133 | 0.1541 | -2.6814 | 0.00733243 | 0.037868786 |
| ADAP1 | 1709.6284 | -0.4133 | 0.1216 | -3.4003 | 0.000673213 | 0.005822583 |
| CNPY4 | 1758.8477 | -0.4138 | 0.0791 | -5.2303 | 1.69E-07 | 5.74E-06 |
| ZNF362 | 1288.8424 | -0.4139 | 0.0825 | -5.0153 | 5.30E-07 | 1.55E-05 |
| SNPH | 655.2521 | -0.4139 | 0.1261 | -3.2820 | 0.00103066 | 0.008206245 |
| FBXL16 | 2081.1184 | -0.4139 | 0.1137 | -3.6392 | 0.000273464 | 0.002841136 |
| ZNF527 | 374.1580 | -0.4143 | 0.1579 | -2.6229 | 0.008717677 | 0.043053312 |
| CENPB | 2259.0408 | -0.4148 | 0.1352 | -3.0672 | 0.002160908 | 0.014730985 |
| RBM45 | 696.9413 | -0.4148 | 0.1322 | -3.1374 | 0.001704315 | 0.012189679 |
| C21orf49 | 459.7736 | -0.4150 | 0.1565 | -2.6519 | 0.008003621 | 0.040303189 |
| LRRC39 | 486.9515 | -0.4157 | 0.1609 | -2.5833 | 0.009785993 | 0.047099176 |
| TP53TG1 | 943.0760 | -0.4162 | 0.1524 | -2.7308 | 0.006317284 | 0.033791214 |
| MICALL1 | 1171.9879 | -0.4167 | 0.1187 | -3.5096 | 0.000448805 | 0.004227315 |
| PGAP3 | 1497.1794 | -0.4167 | 0.0987 | -4.2235 | 2.41E-05 | 0.000386625 |
| LY6G5C | 247.7290 | -0.4168 | 0.1461 | -2.8532 | 0.004328457 | 0.02534904 |
| ISYNA1 | 920.1755 | -0.4172 | 0.1376 | -3.0309 | 0.002438555 | 0.016157698 |
| PTPN12 | 2372.9633 | -0.4176 | 0.1068 | -3.9110 | 9.19E-05 | 0.001173356 |
| NBPF11 | 406.4990 | -0.4177 | 0.1332 | -3.1364 | 0.001710189 | 0.0122086 |
| PDLIM2 | 1269.6370 | -0.4179 | 0.0959 | -4.3580 | 1.31E-05 | 0.000234968 |
| LOC400657 | 1031.6672 | -0.4180 | 0.1259 | -3.3203 | 0.000899321 | 0.007357506 |
| SPIN3 | 1212.4408 | -0.4182 | 0.1360 | -3.0756 | 0.002100919 | 0.014380362 |
| BBS2 | 1718.9644 | -0.4183 | 0.1135 | -3.6862 | 0.000227666 | 0.002449434 |
| C14orf149 | 1055.0840 | -0.4186 | 0.1223 | -3.4231 | 0.000619073 | 0.005431975 |
| CROCC | 1712.6555 | -0.4187 | 0.1078 | -3.8852 | 0.000102263 | 0.001280569 |
| ADCY4 | 859.6520 | -0.4195 | 0.1077 | -3.8935 | 9.88E-05 | 0.001244681 |
| ZNF37BP | 4129.7448 | -0.4196 | 0.1374 | -3.0546 | 0.002253706 | 0.015172146 |
| TMEM5 | 895.6031 | -0.4199 | 0.1386 | -3.0283 | 0.002459123 | 0.016229996 |
| DOK2 | 9505.1204 | -0.4200 | 0.1066 | -3.9381 | 8.21E-05 | 0.001073672 |
| LRRC33 | 2430.6312 | -0.4209 | 0.0972 | -4.3306 | 1.49E-05 | 0.000260658 |
| ZNF641 | 887.7408 | -0.4209 | 0.1465 | -2.8735 | 0.004059929 | 0.024074693 |
| COQ10A | 696.1860 | -0.4212 | 0.0997 | -4.2224 | 2.42E-05 | 0.000388099 |
| ARHGAP4 | 13308.5850 | -0.4213 | 0.1056 | -3.9899 | 6.61E-05 | 0.000898325 |
| ZNF517 | 567.4606 | -0.4217 | 0.1442 | -2.9239 | 0.003456919 | 0.021162613 |
| LINC00476 | 1133.1666 | -0.4224 | 0.1146 | -3.6845 | 0.000229159 | 0.002456761 |
| LOC220906 | 1540.3495 | -0.4224 | 0.1110 | -3.8062 | 0.000141139 | 0.001670399 |
| FHOD3 | 689.6730 | -0.4224 | 0.1158 | -3.6475 | 0.000264774 | 0.002758426 |
| CTDSP1 | 12605.3497 | -0.4229 | 0.0952 | -4.4435 | 8.85E-06 | 0.000169078 |
| NSUN5P2 | 818.4853 | -0.4236 | 0.1440 | -2.9419 | 0.003261585 | 0.020270336 |
| ULK4 | 644.4465 | -0.4236 | 0.1524 | -2.7798 | 0.00543963 | 0.030229713 |
| PLEKHG3 | 3899.9146 | -0.4238 | 0.1235 | -3.4309 | 0.000601638 | 0.005303601 |
| LOC282997 | 437.8744 | -0.4242 | 0.1333 | -3.1817 | 0.001464053 | 0.010803283 |
| ZNF232 | 353.8424 | -0.4245 | 0.1289 | -3.2934 | 0.000989759 | 0.007936348 |
| AKTIP | 2973.9538 | -0.4249 | 0.0994 | -4.2750 | 1.91E-05 | 0.000322321 |
| LRRC8D | 4000.8371 | -0.4252 | 0.0784 | -5.4234 | 5.85E-08 | 2.30E-06 |
| TMEM55A | 857.8726 | -0.4254 | 0.1200 | -3.5444 | 0.000393442 | 0.003795662 |
| DMAP1 | 3214.4261 | -0.4255 | 0.0947 | -4.4910 | 7.09E-06 | 0.000140001 |
| CNN2 | 22500.2251 | -0.4260 | 0.1124 | -3.7888 | 0.000151403 | 0.001752789 |
| DYRK1B | 1493.0250 | -0.4267 | 0.0973 | -4.3866 | 1.15E-05 | 0.000209631 |
| PLEKHG5 | 212.5401 | -0.4268 | 0.1512 | -2.8237 | 0.004747303 | 0.0273155 |
| ITGB7 | 22355.0863 | -0.4268 | 0.0771 | -5.5392 | 3.04E-08 | 1.32E-06 |
| CHST12 | 1541.8502 | -0.4271 | 0.1498 | -2.8509 | 0.004360251 | 0.025480561 |
| ZNF792 | 1288.7651 | -0.4274 | 0.0984 | -4.3435 | 1.40E-05 | 0.000249008 |
| AXIN1 | 632.4578 | -0.4274 | 0.1607 | -2.6597 | 0.007821664 | 0.039650907 |
| ABLIM1 | 15608.2400 | -0.4275 | 0.0828 | -5.1615 | 2.45E-07 | 8.00E-06 |
| KRCC1 | 3796.8032 | -0.4280 | 0.1036 | -4.1307 | 3.62E-05 | 0.00054428 |
| CYB561D2 | 1231.8111 | -0.4285 | 0.1341 | -3.1947 | 0.001399627 | 0.010419369 |
| LINC00339 | 1098.2696 | -0.4291 | 0.1372 | -3.1277 | 0.001761511 | 0.012492444 |
| ZNF302 | 1751.8610 | -0.4294 | 0.1002 | -4.2845 | 1.83E-05 | 0.000310673 |
| S1PR1 | 9954.4790 | -0.4297 | 0.1057 | -4.0641 | 4.82E-05 | 0.000694082 |
| ANKMY1 | 2304.0470 | -0.4307 | 0.0868 | -4.9603 | 7.04E-07 | 1.97E-05 |
| TMEM116 | 1333.2722 | -0.4310 | 0.1097 | -3.9278 | 8.57E-05 | 0.001106267 |
| ZNF557 | 681.2718 | -0.4314 | 0.1387 | -3.1095 | 0.001873783 | 0.013116484 |
| WWOX | 2291.9595 | -0.4318 | 0.1228 | -3.5164 | 0.000437366 | 0.004135026 |
| LOC646214 | 619.3334 | -0.4320 | 0.1492 | -2.8954 | 0.003786906 | 0.022795274 |
| USP46 | 1340.9395 | -0.4325 | 0.1118 | -3.8695 | 0.000109077 | 0.001345852 |
| MYO5B | 926.8888 | -0.4330 | 0.1521 | -2.8466 | 0.004418711 | 0.02572819 |
| C9orf95 | 8473.9675 | -0.4337 | 0.1153 | -3.7617 | 0.00016876 | 0.001917976 |
| EIF2C4 | 6670.9559 | -0.4340 | 0.0820 | -5.2940 | 1.20E-07 | 4.27E-06 |
| C20orf177 | 1732.2754 | -0.4348 | 0.0780 | -5.5759 | 2.46E-08 | 1.10E-06 |
| SUN2 | 39818.5298 | -0.4350 | 0.0853 | -5.0980 | 3.43E-07 | 1.06E-05 |
| ECHDC2 | 6813.0507 | -0.4356 | 0.0599 | -7.2721 | 3.54E-13 | 4.28E-11 |
| ZNF540 | 286.5815 | -0.4360 | 0.1634 | -2.6687 | 0.007614749 | 0.038914969 |
| SLC39A11 | 855.7109 | -0.4367 | 0.1087 | -4.0167 | 5.90E-05 | 0.000822203 |
| MRM1 | 570.8105 | -0.4375 | 0.1387 | -3.1536 | 0.001612489 | 0.01167087 |
| ZRANB2-AS1 | 379.8108 | -0.4376 | 0.1366 | -3.2033 | 0.001358424 | 0.010157363 |
| C4orf19 | 644.8192 | -0.4386 | 0.1305 | -3.3607 | 0.000777396 | 0.00654406 |
| ZSWIM7 | 593.6697 | -0.4390 | 0.1553 | -2.8262 | 0.004709924 | 0.027162416 |
| ATP6AP1L | 1308.6772 | -0.4393 | 0.1514 | -2.9024 | 0.003702993 | 0.022370275 |
| LOC375190 | 1625.4054 | -0.4404 | 0.1268 | -3.4719 | 0.00051677 | 0.004716153 |
| WDR52 | 1813.9855 | -0.4406 | 0.1434 | -3.0724 | 0.002123569 | 0.014502587 |
| TRANK1 | 5741.7837 | -0.4415 | 0.1478 | -2.9875 | 0.002812904 | 0.01808455 |
| DPH5 | 1571.2709 | -0.4417 | 0.1165 | -3.7914 | 0.000149816 | 0.001737938 |
| LRP2BP | 489.5930 | -0.4432 | 0.1399 | -3.1690 | 0.001529842 | 0.011185074 |
| NOXRED1 | 397.4650 | -0.4433 | 0.1647 | -2.6922 | 0.007098283 | 0.036866656 |
| ZNF595 | 352.1123 | -0.4434 | 0.1592 | -2.7841 | 0.005367194 | 0.029992446 |
| LOC100507266 | 632.9200 | -0.4442 | 0.1215 | -3.6566 | 0.000255565 | 0.002677237 |
| ZNF815 | 968.0957 | -0.4452 | 0.0917 | -4.8573 | 1.19E-06 | 3.05E-05 |
| TRPM2 | 323.4144 | -0.4452 | 0.1656 | -2.6885 | 0.007176593 | 0.037178191 |
| PIGP | 805.1900 | -0.4453 | 0.1273 | -3.4975 | 0.00046971 | 0.004377131 |
| C1RL | 3293.2864 | -0.4453 | 0.1150 | -3.8714 | 0.000108231 | 0.001337585 |
| ATP5S | 1439.3518 | -0.4459 | 0.0899 | -4.9595 | 7.07E-07 | 1.97E-05 |
| ANO9 | 9121.1437 | -0.4465 | 0.0780 | -5.7241 | 1.04E-08 | 5.01E-07 |
| ABCA3 | 1493.5962 | -0.4468 | 0.1041 | -4.2921 | 1.77E-05 | 0.000302838 |
| MOB2 | 2352.9961 | -0.4474 | 0.1329 | -3.3671 | 0.000759511 | 0.006422094 |
| MFSD1 | 1477.8324 | -0.4476 | 0.0877 | -5.1028 | 3.35E-07 | 1.04E-05 |
| C5orf63 | 457.6644 | -0.4486 | 0.1621 | -2.7679 | 0.005642733 | 0.030994053 |
| CALHM2 | 4535.8239 | -0.4486 | 0.0938 | -4.7799 | 1.75E-06 | 4.23E-05 |
| AMACR | 481.1970 | -0.4486 | 0.1175 | -3.8164 | 0.000135424 | 0.00161941 |
| HYAL3 | 704.2067 | -0.4492 | 0.1519 | -2.9570 | 0.003106079 | 0.019520423 |
| LOC100506779 | 12659.7515 | -0.4494 | 0.1121 | -4.0093 | 6.09E-05 | 0.00084299 |
| THBS4 | 499.5532 | -0.4496 | 0.1403 | -3.2054 | 0.00134872 | 0.01012008 |
| DNAI2 | 532.3001 | -0.4501 | 0.1572 | -2.8624 | 0.004204462 | 0.024776353 |
| CCDC157 | 439.4314 | -0.4502 | 0.1581 | -2.8477 | 0.004404018 | 0.025662398 |
| DDX43 | 343.4551 | -0.4503 | 0.1528 | -2.9472 | 0.003207127 | 0.019997615 |
| SMPD1 | 2510.8559 | -0.4513 | 0.1046 | -4.3140 | 1.60E-05 | 0.000279141 |
| NAGLU | 879.1095 | -0.4516 | 0.1491 | -3.0296 | 0.002448658 | 0.016196262 |
| TLE4 | 1789.4690 | -0.4518 | 0.1004 | -4.4979 | 6.86E-06 | 0.000136418 |
| PDGFRA | 1347.1236 | -0.4519 | 0.0782 | -5.7780 | 7.56E-09 | 3.75E-07 |
| CCDC85C | 597.1298 | -0.4521 | 0.1346 | -3.3587 | 0.000783168 | 0.006581658 |
| SIRT5 | 2178.6360 | -0.4524 | 0.0838 | -5.3961 | 6.81E-08 | 2.62E-06 |
| CARD14 | 3012.9141 | -0.4527 | 0.1232 | -3.6736 | 0.00023919 | 0.002536132 |
| PCYT2 | 885.1378 | -0.4530 | 0.1499 | -3.0225 | 0.002506729 | 0.016450882 |
| SLC9A9 | 2073.5213 | -0.4530 | 0.1056 | -4.2892 | 1.79E-05 | 0.000305834 |
| MAGEE1 | 378.1768 | -0.4540 | 0.1505 | -3.0175 | 0.002548484 | 0.016631112 |
| MON1A | 737.9459 | -0.4541 | 0.1003 | -4.5259 | 6.01E-06 | 0.000122423 |
| IL17RA | 5072.2905 | -0.4548 | 0.0878 | -5.1776 | 2.25E-07 | 7.39E-06 |
| IMPACT | 872.9200 | -0.4552 | 0.1344 | -3.3870 | 0.000706568 | 0.006055672 |
| LOC100132891 | 458.3356 | -0.4556 | 0.1467 | -3.1052 | 0.001901302 | 0.013296806 |
| SLC9A3R1 | 13783.4550 | -0.4558 | 0.1075 | -4.2405 | 2.23E-05 | 0.000362392 |
| PDE6C | 455.3306 | -0.4560 | 0.1086 | -4.1989 | 2.68E-05 | 0.000420914 |
| ZFP161 | 2671.8214 | -0.4563 | 0.0843 | -5.4127 | 6.21E-08 | 2.41E-06 |
| CATSPERG | 414.8234 | -0.4571 | 0.1467 | -3.1162 | 0.001831705 | 0.012887534 |
| C2orf18 | 1144.1181 | -0.4577 | 0.1271 | -3.5998 | 0.000318454 | 0.003217936 |
| RORC | 1008.6763 | -0.4577 | 0.1373 | -3.3334 | 0.000857949 | 0.007068731 |
| TPK1 | 1183.0211 | -0.4578 | 0.1016 | -4.5060 | 6.60E-06 | 0.000133211 |
| MIR3134 | 1121.3337 | -0.4580 | 0.1129 | -4.0569 | 4.97E-05 | 0.000709725 |
| AP1G2 | 3731.1737 | -0.4583 | 0.1433 | -3.1974 | 0.001386757 | 0.010333727 |
| E2F2 | 2180.8843 | -0.4588 | 0.1542 | -2.9747 | 0.002932878 | 0.018696017 |
| AGBL2 | 510.7112 | -0.4590 | 0.1622 | -2.8294 | 0.004663236 | 0.026944524 |
| FGD3 | 14756.8073 | -0.4595 | 0.0881 | -5.2179 | 1.81E-07 | 6.08E-06 |
| POPDC2 | 622.5176 | -0.4595 | 0.1120 | -4.1019 | 4.10E-05 | 0.000605258 |
| PDE6B | 1044.1940 | -0.4602 | 0.0906 | -5.0774 | 3.83E-07 | 1.16E-05 |
| FAM45B | 3048.9087 | -0.4604 | 0.0932 | -4.9405 | 7.79E-07 | 2.14E-05 |
| MSRA | 443.2621 | -0.4606 | 0.1755 | -2.6248 | 0.008669263 | 0.042896069 |
| NEIL1 | 640.4853 | -0.4609 | 0.1707 | -2.6993 | 0.006947571 | 0.036283412 |
| PTK6 | 350.9966 | -0.4612 | 0.1702 | -2.7103 | 0.006722947 | 0.035533898 |
| ABTB1 | 4857.5000 | -0.4617 | 0.1086 | -4.2499 | 2.14E-05 | 0.000351162 |
| ALDH16A1 | 1735.5908 | -0.4622 | 0.1125 | -4.1093 | 3.97E-05 | 0.000589141 |
| TRIB2 | 4723.5514 | -0.4625 | 0.1322 | -3.4977 | 0.000469336 | 0.004377131 |
| DDX4 | 246.7404 | -0.4628 | 0.1594 | -2.9028 | 0.00369793 | 0.022357548 |
| EPHX1 | 7182.4999 | -0.4635 | 0.0870 | -5.3253 | 1.01E-07 | 3.67E-06 |
| RPS6KA2 | 812.6322 | -0.4638 | 0.1269 | -3.6554 | 0.000256753 | 0.002687821 |
| LOC100507557 | 938.2570 | -0.4641 | 0.1257 | -3.6930 | 0.000221625 | 0.002401516 |
| LOC653160 | 980.3335 | -0.4643 | 0.1399 | -3.3198 | 0.00090091 | 0.007362541 |
| FAM78A | 8634.9285 | -0.4644 | 0.0826 | -5.6208 | 1.90E-08 | 8.71E-07 |
| GLB1L2 | 681.2246 | -0.4645 | 0.1042 | -4.4575 | 8.29E-06 | 0.000159377 |
| OLFM2 | 345.0197 | -0.4652 | 0.1678 | -2.7713 | 0.00558321 | 0.030745258 |
| DYNC2H1 | 928.4778 | -0.4661 | 0.1688 | -2.7613 | 0.005756789 | 0.031460604 |
| ZNF880 | 236.0095 | -0.4666 | 0.1726 | -2.7036 | 0.006858943 | 0.036038633 |
| TOR2A | 739.4096 | -0.4666 | 0.1476 | -3.1607 | 0.001573665 | 0.011422661 |
| EFCAB4A | 429.5962 | -0.4668 | 0.1755 | -2.6603 | 0.007808242 | 0.039635997 |
| LIMD2 | 17091.6527 | -0.4668 | 0.1248 | -3.7402 | 0.000183881 | 0.002071944 |
| KIAA1841 | 782.1247 | -0.4669 | 0.1360 | -3.4323 | 0.000598568 | 0.005285778 |
| HSD17B11 | 4313.7005 | -0.4670 | 0.1099 | -4.2489 | 2.15E-05 | 0.000352028 |
| C8orf44-SGK3 | 848.5113 | -0.4677 | 0.1258 | -3.7181 | 0.000200757 | 0.002215066 |
| GDPD5 | 3466.5005 | -0.4682 | 0.1193 | -3.9260 | 8.64E-05 | 0.001112751 |
| EPHB6 | 545.5031 | -0.4689 | 0.1583 | -2.9629 | 0.003047684 | 0.019230166 |
| SNX30 | 1141.8299 | -0.4690 | 0.1450 | -3.2332 | 0.001224238 | 0.00938624 |
| ACSL6 | 1711.6422 | -0.4690 | 0.1290 | -3.6357 | 0.000277183 | 0.002870169 |
| TMX4 | 5982.3607 | -0.4701 | 0.0943 | -4.9833 | 6.25E-07 | 1.78E-05 |
| SLC2A11 | 1023.3771 | -0.4708 | 0.1134 | -4.1527 | 3.29E-05 | 0.000502618 |
| PARP15 | 5848.1412 | -0.4708 | 0.1324 | -3.5552 | 0.000377693 | 0.003660069 |
| LOC100507567 | 297.2323 | -0.4712 | 0.1610 | -2.9256 | 0.003437824 | 0.021071295 |
| CLU | 2742.8157 | -0.4715 | 0.1553 | -3.0360 | 0.002397657 | 0.015949588 |
| KCTD15 | 385.6660 | -0.4717 | 0.1737 | -2.7161 | 0.006604721 | 0.03503142 |
| WIBG | 1756.3744 | -0.4724 | 0.1341 | -3.5231 | 0.000426551 | 0.004055584 |
| SRGAP3 | 274.8447 | -0.4731 | 0.1508 | -3.1371 | 0.001706317 | 0.012198228 |
| LOC100506033 | 765.8608 | -0.4732 | 0.1313 | -3.6047 | 0.000312468 | 0.003163794 |
| CASP6 | 1313.7174 | -0.4742 | 0.1093 | -4.3399 | 1.43E-05 | 0.000252183 |
| C20orf29 | 652.7621 | -0.4743 | 0.1216 | -3.9018 | 9.55E-05 | 0.001214825 |
| ZNF251 | 1957.0245 | -0.4744 | 0.0930 | -5.1021 | 3.36E-07 | 1.04E-05 |
| NUDT7 | 281.8515 | -0.4750 | 0.1621 | -2.9309 | 0.003379317 | 0.020813898 |
| OAZ3 | 254.5019 | -0.4754 | 0.1614 | -2.9464 | 0.003214752 | 0.020020401 |
| AGBL3 | 627.8431 | -0.4763 | 0.1245 | -3.8254 | 0.00013054 | 0.001575958 |
| ZNF44 | 2839.2103 | -0.4764 | 0.0923 | -5.1591 | 2.48E-07 | 8.09E-06 |
| C9orf156 | 1398.8019 | -0.4764 | 0.0897 | -5.3137 | 1.07E-07 | 3.90E-06 |
| NAIP | 361.0005 | -0.4772 | 0.1290 | -3.6983 | 0.000217023 | 0.002360101 |
| SPON2 | 1549.9418 | -0.4773 | 0.1262 | -3.7808 | 0.000156295 | 0.001803409 |
| TMEM80 | 1788.8134 | -0.4783 | 0.0940 | -5.0865 | 3.65E-07 | 1.12E-05 |
| KLRB1 | 3304.1894 | -0.4792 | 0.1614 | -2.9693 | 0.002985016 | 0.018980382 |
| BIN2 | 15411.9280 | -0.4794 | 0.0798 | -6.0089 | 1.87E-09 | 1.06E-07 |
| TMEM53 | 194.0160 | -0.4827 | 0.1785 | -2.7038 | 0.006855075 | 0.036030825 |
| C17orf103 | 1439.0188 | -0.4831 | 0.1055 | -4.5791 | 4.67E-06 | 9.93E-05 |
| ICA1 | 337.8063 | -0.4835 | 0.1672 | -2.8915 | 0.00383414 | 0.023033771 |
| ZNF429 | 623.2974 | -0.4836 | 0.1530 | -3.1605 | 0.001575087 | 0.011427505 |
| FAM76A | 630.9027 | -0.4836 | 0.1208 | -4.0026 | 6.26E-05 | 0.000863104 |
| ZNF528 | 364.5696 | -0.4840 | 0.1739 | -2.7832 | 0.005382217 | 0.030009875 |
| ACCS | 483.8477 | -0.4841 | 0.1834 | -2.6388 | 0.008319207 | 0.04156032 |
| CAPN3 | 1077.6734 | -0.4846 | 0.1231 | -3.9364 | 8.27E-05 | 0.001076726 |
| GPD1L | 1547.2995 | -0.4850 | 0.0963 | -5.0377 | 4.71E-07 | 1.39E-05 |
| LZTFL1 | 372.9756 | -0.4857 | 0.1649 | -2.9454 | 0.003225121 | 0.020068453 |
| MMP19 | 1020.6066 | -0.4864 | 0.1492 | -3.2595 | 0.001116262 | 0.008695 |
| SEMA6C | 383.9213 | -0.4874 | 0.1598 | -3.0498 | 0.00228987 | 0.015374549 |
| NEK3 | 567.2838 | -0.4874 | 0.1482 | -3.2896 | 0.001003234 | 0.008008398 |
| ORAOV1 | 1793.9953 | -0.4879 | 0.1066 | -4.5782 | 4.69E-06 | 9.96E-05 |
| KCNMB4 | 551.6099 | -0.4884 | 0.1279 | -3.8195 | 0.000133716 | 0.001607887 |
| GAS6 | 1596.4653 | -0.4886 | 0.1312 | -3.7238 | 0.000196281 | 0.002181585 |
| NCF2 | 1870.8527 | -0.4888 | 0.1338 | -3.6522 | 0.00026005 | 0.0027167 |
| LOC139201 | 140.0766 | -0.4891 | 0.1876 | -2.6067 | 0.009141812 | 0.04463789 |
| 2-Mar | 913.0237 | -0.4897 | 0.1365 | -3.5881 | 0.000333099 | 0.003330329 |
| MPP5 | 1090.8950 | -0.4902 | 0.0797 | -6.1506 | 7.72E-10 | 4.73E-08 |
| KLRG1 | 650.0848 | -0.4919 | 0.1850 | -2.6586 | 0.007847551 | 0.039765763 |
| ZSWIM1 | 461.2770 | -0.4925 | 0.1460 | -3.3737 | 0.000741763 | 0.0063002 |
| ARHGEF12 | 9477.0613 | -0.4930 | 0.0932 | -5.2899 | 1.22E-07 | 4.36E-06 |
| ADI1 | 2136.7942 | -0.4937 | 0.0991 | -4.9817 | 6.30E-07 | 1.79E-05 |
| ANPEP | 406.3287 | -0.4945 | 0.1453 | -3.4029 | 0.000666726 | 0.005783004 |
| ITIH4 | 984.6698 | -0.4947 | 0.1164 | -4.2497 | 2.14E-05 | 0.000351162 |
| CSF1R | 365.3773 | -0.4950 | 0.1887 | -2.6225 | 0.008728978 | 0.043080997 |
| PCYOX1L | 3239.9214 | -0.4953 | 0.1094 | -4.5274 | 5.97E-06 | 0.000121919 |
| ZNF587 | 427.8012 | -0.4955 | 0.1494 | -3.3173 | 0.000908936 | 0.007407339 |
| APC2 | 1158.3259 | -0.4956 | 0.1110 | -4.4630 | 8.08E-06 | 0.000156148 |
| C20orf196 | 669.9874 | -0.4960 | 0.1782 | -2.7832 | 0.005382051 | 0.030009875 |
| FRAT1 | 507.1918 | -0.4960 | 0.1532 | -3.2381 | 0.001203294 | 0.009257673 |
| CTNNBIP1 | 598.2556 | -0.4960 | 0.1331 | -3.7268 | 0.000193892 | 0.0021598 |
| AIF1 | 368.1169 | -0.4962 | 0.1855 | -2.6749 | 0.007474775 | 0.038446418 |
| LOC648987 | 328.5207 | -0.4967 | 0.1454 | -3.4163 | 0.00063469 | 0.005549689 |
| TCTA | 1566.8284 | -0.4977 | 0.0934 | -5.3281 | 9.93E-08 | 3.63E-06 |
| ALS2CR8 | 2391.7460 | -0.4989 | 0.1465 | -3.4056 | 0.000660118 | 0.005742152 |
| KIAA1984 | 451.9853 | -0.4993 | 0.1199 | -4.1647 | 3.12E-05 | 0.000479203 |
| KANK1 | 1202.9326 | -0.4995 | 0.1147 | -4.3533 | 1.34E-05 | 0.00023921 |
| SOX12 | 773.8813 | -0.5005 | 0.1314 | -3.8093 | 0.000139387 | 0.001657635 |
| KLF3 | 839.7419 | -0.5006 | 0.1420 | -3.5253 | 0.00042298 | 0.004029227 |
| GIPC1 | 1565.4261 | -0.5010 | 0.1382 | -3.6243 | 0.000289767 | 0.002973745 |
| CFL1P1 | 136.7997 | -0.5013 | 0.1886 | -2.6580 | 0.007859877 | 0.039791286 |
| TEF | 506.4723 | -0.5015 | 0.1416 | -3.5416 | 0.000397647 | 0.003826471 |
| C11orf71 | 728.7255 | -0.5022 | 0.1339 | -3.7501 | 0.00017675 | 0.002001269 |
| LOC100506233 | 323.5556 | -0.5024 | 0.1779 | -2.8242 | 0.00473968 | 0.027282015 |
| SLC25A20 | 3219.5177 | -0.5025 | 0.1123 | -4.4761 | 7.60E-06 | 0.000148027 |
| FAAH | 243.5321 | -0.5032 | 0.1761 | -2.8567 | 0.004280592 | 0.025136846 |
| ZNF181 | 508.9046 | -0.5032 | 0.1374 | -3.6631 | 0.000249225 | 0.002619186 |
| SH3TC2 | 456.4686 | -0.5035 | 0.1611 | -3.1258 | 0.001773478 | 0.012542029 |
| SLC35D2 | 2305.7384 | -0.5037 | 0.0980 | -5.1409 | 2.73E-07 | 8.76E-06 |
| EPB41 | 16706.4111 | -0.5038 | 0.0946 | -5.3267 | 1.00E-07 | 3.65E-06 |
| ANXA9 | 690.6834 | -0.5038 | 0.1248 | -4.0358 | 5.44E-05 | 0.000766379 |
| SLC29A3 | 284.9292 | -0.5044 | 0.1385 | -3.6404 | 0.000272256 | 0.00283053 |
| PRCD | 167.3538 | -0.5052 | 0.1876 | -2.6921 | 0.007099396 | 0.036866656 |
| CDKL1 | 494.7428 | -0.5055 | 0.1324 | -3.8192 | 0.000133874 | 0.001608505 |
| PROK2 | 710.9130 | -0.5067 | 0.1880 | -2.6950 | 0.007037825 | 0.036634955 |
| ZBTB20 | 1453.0477 | -0.5067 | 0.1830 | -2.7689 | 0.005624169 | 0.030925774 |
| GYG1 | 2802.4692 | -0.5069 | 0.1148 | -4.4139 | 1.02E-05 | 0.000188896 |
| GALNT6 | 1278.3699 | -0.5072 | 0.1286 | -3.9438 | 8.02E-05 | 0.00105601 |
| PIGZ | 366.5626 | -0.5077 | 0.1425 | -3.5625 | 0.000367277 | 0.003587272 |
| DHRS12 | 609.9256 | -0.5079 | 0.1201 | -4.2289 | 2.35E-05 | 0.000378277 |
| TXNIP | 47593.0642 | -0.5094 | 0.0772 | -6.5944 | 4.27E-11 | 3.43E-09 |
| C5orf56 | 1493.2414 | -0.5100 | 0.1210 | -4.2155 | 2.49E-05 | 0.000396783 |
| ARL15 | 959.7863 | -0.5100 | 0.1350 | -3.7786 | 0.000157709 | 0.001816951 |
| IL7 | 314.1288 | -0.5107 | 0.1949 | -2.6206 | 0.008778545 | 0.043256562 |
| UBL3 | 3630.5700 | -0.5108 | 0.0844 | -6.0511 | 1.44E-09 | 8.34E-08 |
| MST1P2 | 561.3191 | -0.5116 | 0.1329 | -3.8487 | 0.000118722 | 0.001448315 |
| LOC79015 | 291.9687 | -0.5116 | 0.1731 | -2.9551 | 0.003125719 | 0.019611263 |
| ENO3 | 1641.7317 | -0.5119 | 0.1203 | -4.2547 | 2.09E-05 | 0.000345421 |
| KLF8 | 1264.7434 | -0.5121 | 0.1279 | -4.0031 | 6.25E-05 | 0.000862219 |
| C2orf63 | 1791.7863 | -0.5126 | 0.1366 | -3.7532 | 0.000174577 | 0.001978152 |
| GCNT7 | 569.2493 | -0.5129 | 0.1861 | -2.7565 | 0.005842678 | 0.031837964 |
| PDK2 | 875.3301 | -0.5135 | 0.1572 | -3.2673 | 0.001085757 | 0.008518802 |
| HCG26 | 557.8693 | -0.5140 | 0.1852 | -2.7755 | 0.005511994 | 0.030530918 |
| GSTZ1 | 683.2648 | -0.5143 | 0.1641 | -3.1344 | 0.001721995 | 0.01228129 |
| LOC340544 | 200.9839 | -0.5149 | 0.1773 | -2.9035 | 0.003689637 | 0.022325259 |
| EFHC1 | 397.5009 | -0.5153 | 0.1494 | -3.4482 | 0.000564235 | 0.005065391 |
| ZNF606 | 474.2161 | -0.5156 | 0.1231 | -4.1897 | 2.79E-05 | 0.000436965 |
| NAAA | 4013.5141 | -0.5157 | 0.0819 | -6.3002 | 2.97E-10 | 1.96E-08 |
| ATPBD4 | 298.6182 | -0.5158 | 0.1834 | -2.8131 | 0.004906204 | 0.028035971 |
| C5orf58 | 1475.6815 | -0.5159 | 0.0823 | -6.2678 | 3.66E-10 | 2.37E-08 |
| HEXIM2 | 615.5525 | -0.5174 | 0.1469 | -3.5232 | 0.000426328 | 0.004055584 |
| GALNT12 | 451.2650 | -0.5181 | 0.1657 | -3.1271 | 0.001765599 | 0.012509704 |
| ATP8B1 | 378.6343 | -0.5189 | 0.1898 | -2.7333 | 0.00627008 | 0.033633866 |
| POU6F1 | 979.6559 | -0.5199 | 0.1043 | -4.9832 | 6.25E-07 | 1.78E-05 |
| PGAM2 | 1019.4273 | -0.5203 | 0.1341 | -3.8792 | 0.000104815 | 0.001305662 |
| TPM2 | 882.7545 | -0.5207 | 0.1655 | -3.1464 | 0.001652937 | 0.01188402 |
| LOC642852 | 1654.3861 | -0.5207 | 0.1383 | -3.7663 | 0.000165675 | 0.00188859 |
| RBPMS | 346.7334 | -0.5214 | 0.1449 | -3.5971 | 0.000321829 | 0.003245541 |
| C17orf66 | 1430.6355 | -0.5218 | 0.1466 | -3.5586 | 0.000372819 | 0.003624446 |
| GLT25D2 | 954.0357 | -0.5229 | 0.1452 | -3.6002 | 0.000317929 | 0.003214778 |
| ESPN | 638.4790 | -0.5232 | 0.1618 | -3.2332 | 0.001224178 | 0.00938624 |
| SYCE1L | 1853.5615 | -0.5240 | 0.1293 | -4.0525 | 5.07E-05 | 0.000719753 |
| POLN | 1147.6225 | -0.5247 | 0.1111 | -4.7223 | 2.33E-06 | 5.41E-05 |
| B3GALTL | 482.6897 | -0.5247 | 0.1124 | -4.6670 | 3.06E-06 | 6.84E-05 |
| ZNF319 | 1395.7123 | -0.5258 | 0.1234 | -4.2613 | 2.03E-05 | 0.000337481 |
| MATK | 773.7339 | -0.5269 | 0.1651 | -3.1908 | 0.001418751 | 0.010525479 |
| TMEM192 | 615.7014 | -0.5282 | 0.1156 | -4.5703 | 4.87E-06 | 0.000103038 |
| ABAT | 1165.2233 | -0.5295 | 0.1094 | -4.8381 | 1.31E-06 | 3.32E-05 |
| PNMA3 | 627.6884 | -0.5306 | 0.1622 | -3.2718 | 0.001068599 | 0.008414731 |
| TMEM42 | 832.2973 | -0.5307 | 0.1262 | -4.2037 | 2.63E-05 | 0.000413795 |
| ZNF440 | 670.1306 | -0.5313 | 0.1677 | -3.1676 | 0.001537116 | 0.011227402 |
| RASA4P | 630.4940 | -0.5318 | 0.1290 | -4.1243 | 3.72E-05 | 0.000556306 |
| LOC283070 | 548.6716 | -0.5322 | 0.1205 | -4.4180 | 9.96E-06 | 0.000186719 |
| ABCG1 | 388.7209 | -0.5333 | 0.1653 | -3.2268 | 0.001251731 | 0.009558271 |
| HMOX1 | 779.7443 | -0.5336 | 0.1998 | -2.6715 | 0.007552204 | 0.038726167 |
| SNHG11 | 498.5661 | -0.5341 | 0.1450 | -3.6824 | 0.000231042 | 0.00246994 |
| ZNF799 | 283.1516 | -0.5347 | 0.1691 | -3.1623 | 0.001565334 | 0.011378571 |
| SYTL2 | 11484.7669 | -0.5347 | 0.0997 | -5.3631 | 8.18E-08 | 3.10E-06 |
| CBLL1 | 5240.1750 | -0.5350 | 0.0970 | -5.5173 | 3.44E-08 | 1.45E-06 |
| LOC100133286 | 281.7933 | -0.5353 | 0.1686 | -3.1756 | 0.001495182 | 0.010990098 |
| HIST1H2BD | 702.7872 | -0.5356 | 0.1475 | -3.6303 | 0.00028309 | 0.00291313 |
| CACNA2D4 | 1395.7674 | -0.5359 | 0.1107 | -4.8413 | 1.29E-06 | 3.27E-05 |
| RIPK3 | 1267.6735 | -0.5371 | 0.1169 | -4.5946 | 4.34E-06 | 9.29E-05 |
| SLC44A1 | 581.4018 | -0.5375 | 0.1412 | -3.8071 | 0.000140608 | 0.001666906 |
| DDX28 | 455.3695 | -0.5383 | 0.1662 | -3.2381 | 0.001203302 | 0.009257673 |
| LOC100130855 | 855.9761 | -0.5389 | 0.1263 | -4.2681 | 1.97E-05 | 0.00032946 |
| ZNF75A | 1250.3138 | -0.5394 | 0.1358 | -3.9710 | 7.16E-05 | 0.000960481 |
| PTGDR2 | 925.8721 | -0.5395 | 0.1292 | -4.1761 | 2.97E-05 | 0.00046061 |
| ANXA4 | 4906.3520 | -0.5402 | 0.1073 | -5.0357 | 4.76E-07 | 1.40E-05 |
| SLC27A3 | 1059.2700 | -0.5403 | 0.1619 | -3.3376 | 0.000845182 | 0.006974939 |
| NR1D2 | 1564.1583 | -0.5405 | 0.0977 | -5.5353 | 3.11E-08 | 1.34E-06 |
| OPRL1 | 323.2991 | -0.5411 | 0.1514 | -3.5749 | 0.000350334 | 0.003472806 |
| C14orf64 | 1575.6198 | -0.5422 | 0.1588 | -3.4132 | 0.000642133 | 0.005608284 |
| NID2 | 269.6777 | -0.5422 | 0.1787 | -3.0350 | 0.002405472 | 0.015983653 |
| GPR135 | 436.7841 | -0.5423 | 0.1675 | -3.2378 | 0.001204547 | 0.009258728 |
| TACR1 | 173.2348 | -0.5428 | 0.1921 | -2.8254 | 0.004722106 | 0.027212102 |
| PPP2R2B | 379.8147 | -0.5430 | 0.2005 | -2.7086 | 0.006757282 | 0.035682904 |
| PNPLA7 | 3639.1683 | -0.5438 | 0.1159 | -4.6909 | 2.72E-06 | 6.20E-05 |
| ZNF831 | 3183.4826 | -0.5445 | 0.1544 | -3.5273 | 0.000419749 | 0.004003493 |
| NOSIP | 5468.4403 | -0.5448 | 0.1779 | -3.0621 | 0.002198058 | 0.014900797 |
| DIRC2 | 205.7579 | -0.5449 | 0.1675 | -3.2525 | 0.0011439 | 0.008878282 |
| RCSD1 | 21121.3914 | -0.5453 | 0.0831 | -6.5637 | 5.25E-11 | 4.05E-09 |
| ZBP1 | 2097.0988 | -0.5456 | 0.1175 | -4.6444 | 3.41E-06 | 7.49E-05 |
| LOC100271722 | 316.6686 | -0.5463 | 0.1911 | -2.8593 | 0.004245595 | 0.02497982 |
| C3orf20 | 559.6621 | -0.5473 | 0.1880 | -2.9120 | 0.003591211 | 0.021864884 |
| ZNF554 | 286.3381 | -0.5474 | 0.1634 | -3.3509 | 0.000805409 | 0.006723267 |
| LOC100128288 | 288.1395 | -0.5475 | 0.1609 | -3.4030 | 0.000666524 | 0.005783004 |
| LOC728392 | 2816.5520 | -0.5481 | 0.1600 | -3.4268 | 0.000610703 | 0.005366376 |
| CSRP2BP | 815.7661 | -0.5483 | 0.1229 | -4.4600 | 8.20E-06 | 0.000157953 |
| TRIM74 | 219.9764 | -0.5490 | 0.1543 | -3.5578 | 0.000374 | 0.003631256 |
| FAM66C | 355.2164 | -0.5494 | 0.1683 | -3.2655 | 0.00109268 | 0.008555367 |
| UPB1 | 197.7303 | -0.5496 | 0.2048 | -2.6830 | 0.007296592 | 0.037696569 |
| MST1 | 1197.1371 | -0.5497 | 0.1445 | -3.8051 | 0.000141771 | 0.001675446 |
| LOC100505746 | 1614.4236 | -0.5498 | 0.1698 | -3.2380 | 0.001203797 | 0.009257673 |
| LOC730091 | 2943.0275 | -0.5500 | 0.1370 | -4.0154 | 5.94E-05 | 0.000825224 |
| TLR1 | 426.2091 | -0.5503 | 0.1714 | -3.2109 | 0.00132338 | 0.009985458 |
| TK2 | 1582.9681 | -0.5523 | 0.1077 | -5.1270 | 2.94E-07 | 9.26E-06 |
| SH3D19 | 248.6361 | -0.5545 | 0.1794 | -3.0904 | 0.001998703 | 0.013837245 |
| MYH3 | 221.6341 | -0.5546 | 0.1972 | -2.8123 | 0.004918294 | 0.028053936 |
| PINK1 | 1617.5077 | -0.5550 | 0.0970 | -5.7228 | 1.05E-08 | 5.03E-07 |
| LLGL2 | 1586.9130 | -0.5572 | 0.1154 | -4.8271 | 1.39E-06 | 3.47E-05 |
| YPEL1 | 4060.2556 | -0.5581 | 0.1123 | -4.9681 | 6.76E-07 | 1.92E-05 |
| SIGIRR | 12289.7867 | -0.5596 | 0.1143 | -4.8964 | 9.76E-07 | 2.58E-05 |
| LOC146880 | 2991.1007 | -0.5602 | 0.0834 | -6.7182 | 1.84E-11 | 1.60E-09 |
| C9orf103 | 981.4159 | -0.5603 | 0.1143 | -4.9015 | 9.51E-07 | 2.53E-05 |
| CCDC39 | 341.0578 | -0.5604 | 0.1712 | -3.2732 | 0.001063506 | 0.008392101 |
| TMEM19 | 695.7526 | -0.5614 | 0.1367 | -4.1072 | 4.00E-05 | 0.000593364 |
| PCMTD2 | 7983.1204 | -0.5617 | 0.0929 | -6.0434 | 1.51E-09 | 8.69E-08 |
| MYO15B | 2305.6022 | -0.5636 | 0.1062 | -5.3076 | 1.11E-07 | 4.02E-06 |
| C3orf62 | 584.3245 | -0.5641 | 0.1286 | -4.3857 | 1.16E-05 | 0.000210214 |
| C17orf72 | 391.3305 | -0.5642 | 0.1889 | -2.9866 | 0.002820587 | 0.018125327 |
| LOC100505622 | 242.3669 | -0.5649 | 0.1706 | -3.3105 | 0.000931193 | 0.00754491 |
| RENBP | 1131.0179 | -0.5650 | 0.1564 | -3.6127 | 0.000303056 | 0.003085012 |
| PION | 1440.0676 | -0.5652 | 0.1087 | -5.1987 | 2.01E-07 | 6.64E-06 |
| KIAA1024 | 175.0851 | -0.5663 | 0.2126 | -2.6630 | 0.007744828 | 0.039386034 |
| RARRES3 | 4039.3943 | -0.5667 | 0.1720 | -3.2949 | 0.00098453 | 0.007908841 |
| TRPV2 | 5779.2175 | -0.5678 | 0.0928 | -6.1170 | 9.53E-10 | 5.79E-08 |
| ZNF287 | 331.7932 | -0.5685 | 0.1573 | -3.6137 | 0.000301903 | 0.003079494 |
| FLJ12334 | 1112.2994 | -0.5690 | 0.1624 | -3.5030 | 0.000459973 | 0.00431217 |
| PLEKHB1 | 420.3602 | -0.5691 | 0.1742 | -3.2664 | 0.001089179 | 0.008532375 |
| ASPRV1 | 220.3655 | -0.5695 | 0.2142 | -2.6585 | 0.007849578 | 0.039765763 |
| EML5 | 214.8411 | -0.5695 | 0.1811 | -3.1455 | 0.00165817 | 0.011910324 |
| PPM1N | 555.5933 | -0.5700 | 0.1400 | -4.0710 | 4.68E-05 | 0.000676965 |
| TIGD2 | 224.6007 | -0.5722 | 0.1861 | -3.0747 | 0.002107372 | 0.014411494 |
| CDC42EP3 | 2445.6507 | -0.5730 | 0.0859 | -6.6688 | 2.58E-11 | 2.17E-09 |
| KLF2 | 8155.0614 | -0.5746 | 0.1581 | -3.6346 | 0.000278461 | 0.002881177 |
| TLR5 | 158.0612 | -0.5749 | 0.1853 | -3.1029 | 0.001916126 | 0.013381916 |
| ZMYND10 | 327.2242 | -0.5750 | 0.1320 | -4.3559 | 1.33E-05 | 0.00023665 |
| SEC1 | 213.0165 | -0.5756 | 0.2181 | -2.6387 | 0.008323656 | 0.041568815 |
| RCBTB2 | 7395.2819 | -0.5787 | 0.1358 | -4.2604 | 2.04E-05 | 0.000338466 |
| IL16 | 16364.9088 | -0.5788 | 0.0820 | -7.0591 | 1.68E-12 | 1.78E-10 |
| MUC20 | 592.3617 | -0.5790 | 0.1318 | -4.3944 | 1.11E-05 | 0.000204181 |
| TSGA10 | 513.9571 | -0.5797 | 0.2021 | -2.8683 | 0.004126494 | 0.024421546 |
| CD93 | 348.4439 | -0.5798 | 0.1704 | -3.4026 | 0.000667423 | 0.005785737 |
| ZNF124 | 477.1651 | -0.5800 | 0.1461 | -3.9691 | 7.22E-05 | 0.000965947 |
| MEGF6 | 532.7713 | -0.5818 | 0.1529 | -3.8065 | 0.000140965 | 0.001669839 |
| LOC100131655 | 580.4466 | -0.5819 | 0.1185 | -4.9097 | 9.12E-07 | 2.45E-05 |
| MAPK8IP1 | 104.6938 | -0.5824 | 0.1880 | -3.0982 | 0.00194676 | 0.01352597 |
| FAM21B | 560.2827 | -0.5825 | 0.1801 | -3.2340 | 0.001220858 | 0.009374576 |
| PRDM11 | 229.4122 | -0.5828 | 0.1793 | -3.2512 | 0.001149238 | 0.008910567 |
| P2RX1 | 442.3840 | -0.5838 | 0.1942 | -3.0058 | 0.00264912 | 0.017169342 |
| CCDC112 | 365.7748 | -0.5838 | 0.1932 | -3.0211 | 0.002518836 | 0.016506688 |
| EPB49 | 952.5863 | -0.5844 | 0.1192 | -4.9044 | 9.37E-07 | 2.50E-05 |
| ERP27 | 424.1993 | -0.5845 | 0.1357 | -4.3060 | 1.66E-05 | 0.000287667 |
| ZNF575 | 270.6436 | -0.5847 | 0.2021 | -2.8926 | 0.003820359 | 0.022969224 |
| THEM4 | 3058.2971 | -0.5849 | 0.0872 | -6.7078 | 1.98E-11 | 1.70E-09 |
| WNT10B | 626.9077 | -0.5864 | 0.1416 | -4.1413 | 3.45E-05 | 0.000524461 |
| LOC100379224 | 1054.7584 | -0.5880 | 0.1186 | -4.9585 | 7.10E-07 | 1.97E-05 |
| ITGA6 | 3986.3387 | -0.5886 | 0.0883 | -6.6639 | 2.67E-11 | 2.23E-09 |
| ACYP2 | 571.5791 | -0.5900 | 0.1464 | -4.0299 | 5.58E-05 | 0.000781682 |
| ZNF33B | 2700.5193 | -0.5915 | 0.1173 | -5.0434 | 4.57E-07 | 1.36E-05 |
| PAQR8 | 1870.5613 | -0.5926 | 0.0909 | -6.5221 | 6.93E-11 | 5.22E-09 |
| STK38 | 26336.7944 | -0.5934 | 0.0934 | -6.3550 | 2.08E-10 | 1.43E-08 |
| MGC39372 | 813.6029 | -0.5941 | 0.1499 | -3.9642 | 7.37E-05 | 0.000985146 |
| ADHFE1 | 1714.9491 | -0.5944 | 0.1159 | -5.1277 | 2.93E-07 | 9.26E-06 |
| NMT2 | 4861.3880 | -0.5946 | 0.0743 | -8.0066 | 1.18E-15 | 2.29E-13 |
| ZNF354A | 550.9894 | -0.5950 | 0.1658 | -3.5894 | 0.000331482 | 0.003318546 |
| CAPG | 5725.2485 | -0.5960 | 0.1609 | -3.7049 | 0.0002115 | 0.002311682 |
| USP54 | 1368.1374 | -0.5971 | 0.0909 | -6.5692 | 5.06E-11 | 3.92E-09 |
| S1PR4 | 13868.9250 | -0.5978 | 0.1592 | -3.7545 | 0.000173696 | 0.001969641 |
| NOS3 | 273.6923 | -0.5983 | 0.1484 | -4.0324 | 5.52E-05 | 0.000775353 |
| ZNF487P | 237.4990 | -0.5992 | 0.1908 | -3.1399 | 0.001690217 | 0.012106021 |
| TTC23 | 188.2438 | -0.5998 | 0.1796 | -3.3403 | 0.000836867 | 0.0069197 |
| LOC153684 | 383.2474 | -0.6023 | 0.1873 | -3.2151 | 0.001303844 | 0.009864631 |
| KCNQ1 | 591.8658 | -0.6027 | 0.1338 | -4.5044 | 6.66E-06 | 0.000133951 |
| LDOC1L | 960.5777 | -0.6036 | 0.1638 | -3.6854 | 0.000228334 | 0.002453132 |
| TGFBR3 | 3123.9689 | -0.6050 | 0.0791 | -7.6454 | 2.08E-14 | 3.25E-12 |
| LEKR1 | 356.3679 | -0.6053 | 0.1803 | -3.3562 | 0.000790126 | 0.006625404 |
| C20orf118 | 2443.6813 | -0.6060 | 0.1729 | -3.5053 | 0.000456049 | 0.004284882 |
| SLC6A16 | 626.8991 | -0.6063 | 0.1698 | -3.5707 | 0.000356032 | 0.003508598 |
| ABCA7 | 7438.4663 | -0.6064 | 0.1035 | -5.8604 | 4.62E-09 | 2.38E-07 |
| UBXN10 | 256.4443 | -0.6064 | 0.2054 | -2.9526 | 0.003150717 | 0.019738407 |
| CREB3L3 | 1854.7432 | -0.6071 | 0.0901 | -6.7385 | 1.60E-11 | 1.42E-09 |
| SLFN5 | 11706.3561 | -0.6093 | 0.1633 | -3.7315 | 0.000190313 | 0.002127763 |
| IPW | 497.3931 | -0.6137 | 0.1466 | -4.1864 | 2.83E-05 | 0.000441902 |
| CCDC102B | 256.3954 | -0.6139 | 0.1706 | -3.5987 | 0.000319867 | 0.003230057 |
| ARMC2 | 1822.5693 | -0.6157 | 0.1286 | -4.7883 | 1.68E-06 | 4.10E-05 |
| VMAC | 510.6984 | -0.6161 | 0.1560 | -3.9481 | 7.88E-05 | 0.001040508 |
| GPM6B | 965.4430 | -0.6166 | 0.1767 | -3.4887 | 0.000485317 | 0.004488897 |
| ALS2CR12 | 285.9099 | -0.6168 | 0.2273 | -2.7133 | 0.006660673 | 0.035266365 |
| EMBP1 | 470.5319 | -0.6174 | 0.1390 | -4.4425 | 8.89E-06 | 0.000169619 |
| GTF2IRD2B | 644.3632 | -0.6200 | 0.2009 | -3.0862 | 0.002027273 | 0.013977466 |
| PLA2G10 | 186.5796 | -0.6214 | 0.1594 | -3.8992 | 9.65E-05 | 0.001223847 |
| PFKFB2 | 158.0883 | -0.6215 | 0.2230 | -2.7870 | 0.005319704 | 0.029793099 |
| SNN | 362.8847 | -0.6215 | 0.1847 | -3.3659 | 0.000763031 | 0.006440479 |
| DUSP23 | 387.9816 | -0.6226 | 0.2149 | -2.8966 | 0.00377285 | 0.022755942 |
| MYO1F | 30509.4227 | -0.6230 | 0.0982 | -6.3442 | 2.24E-10 | 1.53E-08 |
| SDK2 | 2582.3955 | -0.6231 | 0.0901 | -6.9195 | 4.53E-12 | 4.38E-10 |
| ZCWPW1 | 285.9165 | -0.6240 | 0.1709 | -3.6510 | 0.000261222 | 0.002725178 |
| SLC38A7 | 421.6787 | -0.6257 | 0.2194 | -2.8523 | 0.004340289 | 0.025395881 |
| LEF1 | 5360.0696 | -0.6263 | 0.1320 | -4.7444 | 2.09E-06 | 4.90E-05 |
| KCNMB3 | 306.7890 | -0.6263 | 0.1618 | -3.8710 | 0.000108395 | 0.001338518 |
| AKAP7 | 794.4189 | -0.6281 | 0.1405 | -4.4713 | 7.77E-06 | 0.000150973 |
| LOC100507032 | 180.3799 | -0.6290 | 0.1959 | -3.2102 | 0.001326557 | 0.009988465 |
| ZNF69 | 485.9328 | -0.6308 | 0.2205 | -2.8603 | 0.004232575 | 0.024912904 |
| CDC25B | 23447.7139 | -0.6315 | 0.1273 | -4.9600 | 7.05E-07 | 1.97E-05 |
| TTTY10 | 217.6173 | -0.6343 | 0.2428 | -2.6127 | 0.008982255 | 0.044015315 |
| STMN3 | 3578.0283 | -0.6348 | 0.1327 | -4.7838 | 1.72E-06 | 4.16E-05 |
| ERMAP | 1150.8796 | -0.6357 | 0.1062 | -5.9877 | 2.13E-09 | 1.19E-07 |
| GTF2IRD1 | 297.5864 | -0.6360 | 0.2034 | -3.1266 | 0.00176848 | 0.012514926 |
| TJP3 | 545.1435 | -0.6371 | 0.1824 | -3.4926 | 0.000478423 | 0.004439945 |
| ADAM1 | 381.8730 | -0.6377 | 0.1753 | -3.6386 | 0.000274112 | 0.002845911 |
| FAM78B | 94.2275 | -0.6392 | 0.2451 | -2.6078 | 0.009111962 | 0.044535264 |
| SNX21 | 272.3145 | -0.6397 | 0.1986 | -3.2213 | 0.001276006 | 0.00968011 |
| PTPLAD2 | 3537.3268 | -0.6403 | 0.1023 | -6.2569 | 3.93E-10 | 2.53E-08 |
| C1orf162 | 279.9279 | -0.6406 | 0.2327 | -2.7524 | 0.00591532 | 0.032083556 |
| AMY2B | 1777.2455 | -0.6413 | 0.0850 | -7.5437 | 4.57E-14 | 6.77E-12 |
| TPRG1L | 1757.9301 | -0.6419 | 0.1152 | -5.5710 | 2.53E-08 | 1.13E-06 |
| S100A5 | 141.6511 | -0.6420 | 0.2317 | -2.7708 | 0.005591261 | 0.030756004 |
| PRR5-ARHGAP8 | 200.5814 | -0.6431 | 0.1880 | -3.4212 | 0.000623549 | 0.005468076 |
| ALS2CL | 254.0466 | -0.6431 | 0.2088 | -3.0805 | 0.002066254 | 0.01418803 |
| C22orf23 | 356.5893 | -0.6442 | 0.1715 | -3.7573 | 0.000171766 | 0.001950682 |
| AK1 | 4671.1086 | -0.6443 | 0.1583 | -4.0693 | 4.71E-05 | 0.000681157 |
| RASSF3 | 6018.2625 | -0.6445 | 0.1091 | -5.9076 | 3.47E-09 | 1.84E-07 |
| CDHR3 | 604.8544 | -0.6475 | 0.1799 | -3.5980 | 0.000320696 | 0.003236274 |
| LYAR | 3438.4697 | -0.6477 | 0.1604 | -4.0382 | 5.39E-05 | 0.00076019 |
| HSPBAP1 | 722.9205 | -0.6479 | 0.1547 | -4.1881 | 2.81E-05 | 0.000439139 |
| ZNF846 | 648.7900 | -0.6489 | 0.1519 | -4.2729 | 1.93E-05 | 0.000324648 |
| AKR1C3 | 308.9589 | -0.6538 | 0.1906 | -3.4302 | 0.000603052 | 0.005309901 |
| PIK3R3 | 185.5094 | -0.6542 | 0.2157 | -3.0336 | 0.002416911 | 0.016035356 |
| TMEM220 | 483.9620 | -0.6543 | 0.2063 | -3.1713 | 0.001517506 | 0.011121758 |
| MYH11 | 753.2252 | -0.6550 | 0.1455 | -4.5028 | 6.70E-06 | 0.000134516 |
| TMEM136 | 286.4384 | -0.6559 | 0.1592 | -4.1190 | 3.81E-05 | 0.000567706 |
| ADCK1 | 202.6012 | -0.6561 | 0.1912 | -3.4317 | 0.000599834 | 0.005293863 |
| PTPRN2 | 1441.1075 | -0.6564 | 0.1227 | -5.3508 | 8.76E-08 | 3.27E-06 |
| ANKRD55 | 276.7022 | -0.6566 | 0.2096 | -3.1328 | 0.001731717 | 0.012327377 |
| C6orf226 | 190.9461 | -0.6567 | 0.2404 | -2.7315 | 0.006304051 | 0.033732359 |
| PRUNE2 | 893.7568 | -0.6569 | 0.1402 | -4.6858 | 2.79E-06 | 6.33E-05 |
| GRAP | 1445.4456 | -0.6586 | 0.1375 | -4.7879 | 1.69E-06 | 4.10E-05 |
| AUTS2 | 692.3230 | -0.6607 | 0.1232 | -5.3623 | 8.22E-08 | 3.11E-06 |
| LOC100131289 | 444.0817 | -0.6609 | 0.1566 | -4.2205 | 2.44E-05 | 0.000390638 |
| SNED1 | 8719.5401 | -0.6613 | 0.0689 | -9.5953 | 8.37E-22 | 3.52E-19 |
| FAM102A | 16528.7370 | -0.6614 | 0.1027 | -6.4421 | 1.18E-10 | 8.53E-09 |
| ACE | 652.3441 | -0.6615 | 0.1732 | -3.8179 | 0.000134579 | 0.001612115 |
| ERBB3 | 144.6217 | -0.6645 | 0.1997 | -3.3276 | 0.000875952 | 0.007196333 |
| TNK1 | 215.3450 | -0.6646 | 0.2004 | -3.3170 | 0.000910059 | 0.007409014 |
| LAIR1 | 880.0596 | -0.6651 | 0.1617 | -4.1126 | 3.91E-05 | 0.000581821 |
| FAM65B | 18364.8019 | -0.6659 | 0.1024 | -6.4996 | 8.05E-11 | 5.94E-09 |
| CD52 | 31579.6122 | -0.6660 | 0.2239 | -2.9742 | 0.002937461 | 0.018717346 |
| C9orf9 | 191.1013 | -0.6666 | 0.1881 | -3.5449 | 0.000392777 | 0.003791664 |
| TNFAIP8L2-SCNM1 | 124.3788 | -0.6680 | 0.2014 | -3.3168 | 0.000910515 | 0.007409014 |
| SLC46A3 | 818.3130 | -0.6685 | 0.1179 | -5.6682 | 1.44E-08 | 6.69E-07 |
| C8orf84 | 137.9732 | -0.6706 | 0.2384 | -2.8127 | 0.004913291 | 0.028035971 |
| CAPS | 585.0477 | -0.6706 | 0.1950 | -3.4383 | 0.000585282 | 0.005208963 |
| C3AR1 | 885.8837 | -0.6711 | 0.1563 | -4.2928 | 1.76E-05 | 0.0003023 |
| CCL5 | 89393.2074 | -0.6734 | 0.1387 | -4.8561 | 1.20E-06 | 3.06E-05 |
| FLJ37035 | 243.6184 | -0.6737 | 0.1960 | -3.4371 | 0.000587944 | 0.005219381 |
| LOC100129550 | 289.1456 | -0.6741 | 0.2046 | -3.2950 | 0.000984233 | 0.007908841 |
| ARSD | 426.1483 | -0.6741 | 0.1448 | -4.6561 | 3.22E-06 | 7.13E-05 |
| FRMD4A | 361.1038 | -0.6750 | 0.1581 | -4.2690 | 1.96E-05 | 0.000328497 |
| TTC18 | 233.8317 | -0.6762 | 0.2110 | -3.2045 | 0.001352882 | 0.010131964 |
| NANOS1 | 226.0336 | -0.6780 | 0.2113 | -3.2082 | 0.001335746 | 0.010041938 |
| PYROXD2 | 759.8833 | -0.6797 | 0.0964 | -7.0510 | 1.78E-12 | 1.85E-10 |
| LOC144481 | 490.4469 | -0.6804 | 0.1532 | -4.4407 | 8.97E-06 | 0.000170611 |
| MPP7 | 322.3250 | -0.6812 | 0.2098 | -3.2469 | 0.001166771 | 0.009014168 |
| ADAM23 | 405.0660 | -0.6831 | 0.1942 | -3.5171 | 0.000436342 | 0.004127917 |
| TKTL1 | 650.6993 | -0.6846 | 0.1932 | -3.5438 | 0.000394406 | 0.003800116 |
| FLJ44635 | 651.6767 | -0.6846 | 0.1625 | -4.2132 | 2.52E-05 | 0.000400544 |
| WDR81 | 5065.4899 | -0.6856 | 0.1088 | -6.3023 | 2.93E-10 | 1.95E-08 |
| ZNF688 | 714.5105 | -0.6862 | 0.2049 | -3.3493 | 0.00081013 | 0.0067557 |
| LOC100507462 | 87.4358 | -0.6886 | 0.2624 | -2.6247 | 0.008671659 | 0.042896069 |
| LSR | 364.9543 | -0.6888 | 0.1360 | -5.0658 | 4.07E-07 | 1.22E-05 |
| PPFIBP2 | 358.9805 | -0.6889 | 0.2106 | -3.2709 | 0.001071995 | 0.008437077 |
| LRTOMT | 1292.1780 | -0.6902 | 0.1317 | -5.2412 | 1.60E-07 | 5.55E-06 |
| MXRA8 | 847.5108 | -0.6915 | 0.1082 | -6.3914 | 1.64E-10 | 1.16E-08 |
| CERKL | 2273.8729 | -0.6920 | 0.1418 | -4.8816 | 1.05E-06 | 2.74E-05 |
| GLIPR2 | 2175.4984 | -0.6920 | 0.1199 | -5.7737 | 7.76E-09 | 3.82E-07 |
| ZNF35 | 134.4930 | -0.6926 | 0.2669 | -2.5949 | 0.009460849 | 0.045899377 |
| LOC100507053 | 72.9125 | -0.6934 | 0.2479 | -2.7971 | 0.005156961 | 0.029122001 |
| LOC728855 | 1261.6479 | -0.6951 | 0.1139 | -6.1012 | 1.05E-09 | 6.34E-08 |
| ASB13 | 449.7886 | -0.6951 | 0.2023 | -3.4359 | 0.000590618 | 0.005233908 |
| EEPD1 | 406.5213 | -0.6953 | 0.1802 | -3.8580 | 0.000114308 | 0.001404224 |
| CA13 | 160.6600 | -0.6956 | 0.1921 | -3.6213 | 0.000293152 | 0.003002374 |
| LDHAL6A | 390.2428 | -0.6966 | 0.1971 | -3.5339 | 0.000409435 | 0.003927412 |
| PLXND1 | 1153.5525 | -0.6970 | 0.1523 | -4.5775 | 4.71E-06 | 9.99E-05 |
| STS | 204.6782 | -0.6971 | 0.2155 | -3.2345 | 0.001218684 | 0.00936264 |
| ZMAT1 | 867.9894 | -0.6974 | 0.1789 | -3.8979 | 9.70E-05 | 0.00122847 |
| PRAM1 | 220.6663 | -0.6987 | 0.2227 | -3.1366 | 0.001709165 | 0.012207055 |
| TC2N | 24671.9693 | -0.6987 | 0.1041 | -6.7123 | 1.92E-11 | 1.66E-09 |
| CYB561 | 2237.6510 | -0.6995 | 0.1148 | -6.0926 | 1.11E-09 | 6.59E-08 |
| C1orf101 | 203.8331 | -0.7000 | 0.1708 | -4.0973 | 4.18E-05 | 0.000615707 |
| RAB40B | 119.9861 | -0.7037 | 0.2097 | -3.3558 | 0.000791304 | 0.006631606 |
| C5 | 281.1195 | -0.7038 | 0.1518 | -4.6373 | 3.53E-06 | 7.70E-05 |
| NIPAL2 | 225.9809 | -0.7042 | 0.2028 | -3.4719 | 0.000516848 | 0.004716153 |
| DNAJB13 | 447.2031 | -0.7045 | 0.1435 | -4.9078 | 9.21E-07 | 2.47E-05 |
| SCML4 | 1906.4953 | -0.7060 | 0.1221 | -5.7806 | 7.44E-09 | 3.70E-07 |
| TMEM169 | 341.5329 | -0.7075 | 0.1664 | -4.2516 | 2.12E-05 | 0.00034904 |
| SLC40A1 | 180.3969 | -0.7081 | 0.2514 | -2.8171 | 0.004845809 | 0.027808252 |
| LCN12 | 256.5643 | -0.7086 | 0.2203 | -3.2163 | 0.001298739 | 0.009832844 |
| PZP | 217.0815 | -0.7103 | 0.2099 | -3.3841 | 0.00071417 | 0.006110433 |
| BLK | 910.1287 | -0.7110 | 0.2163 | -3.2873 | 0.001011662 | 0.008067163 |
| PKIB | 318.8074 | -0.7157 | 0.2310 | -3.0985 | 0.00194529 | 0.01352597 |
| ZNF252 | 528.9402 | -0.7169 | 0.2222 | -3.2264 | 0.001253709 | 0.009564122 |
| MAP2K6 | 753.4054 | -0.7169 | 0.1504 | -4.7678 | 1.86E-06 | 4.46E-05 |
| ODZ1 | 1383.6331 | -0.7170 | 0.1602 | -4.4747 | 7.65E-06 | 0.000148803 |
| HHAT | 86.9540 | -0.7172 | 0.2731 | -2.6258 | 0.008643584 | 0.042799181 |
| SPIRE2 | 234.4858 | -0.7177 | 0.1856 | -3.8666 | 0.000110365 | 0.001360631 |
| RASGEF1A | 1521.8910 | -0.7219 | 0.1349 | -5.3499 | 8.80E-08 | 3.28E-06 |
| C16orf7 | 1472.4456 | -0.7220 | 0.1635 | -4.4157 | 1.01E-05 | 0.000187912 |
| SPNS3 | 2969.0698 | -0.7231 | 0.1448 | -4.9933 | 5.93E-07 | 1.71E-05 |
| NAALAD2 | 465.1150 | -0.7231 | 0.1592 | -4.5419 | 5.57E-06 | 0.000116142 |
| LOC339874 | 166.3542 | -0.7239 | 0.2350 | -3.0800 | 0.002070284 | 0.014209253 |
| EML6 | 148.2381 | -0.7243 | 0.1814 | -3.9931 | 6.52E-05 | 0.000889406 |
| GTF2IRD2 | 333.7587 | -0.7250 | 0.1619 | -4.4789 | 7.50E-06 | 0.000146641 |
| KIT | 323.5926 | -0.7268 | 0.1899 | -3.8268 | 0.000129835 | 0.001571242 |
| USP50 | 214.6013 | -0.7293 | 0.2028 | -3.5954 | 0.00032394 | 0.003259574 |
| C1orf54 | 205.7654 | -0.7304 | 0.2248 | -3.2487 | 0.001159259 | 0.008960708 |
| CDCP1 | 351.8600 | -0.7374 | 0.1881 | -3.9205 | 8.84E-05 | 0.001134547 |
| CHI3L2 | 146.0911 | -0.7377 | 0.2326 | -3.1711 | 0.001518825 | 0.011126039 |
| GRIP1 | 421.9714 | -0.7392 | 0.1669 | -4.4281 | 9.51E-06 | 0.000179986 |
| ARRDC5 | 436.0494 | -0.7396 | 0.1851 | -3.9961 | 6.44E-05 | 0.000881794 |
| C20orf197 | 274.3113 | -0.7398 | 0.2146 | -3.4479 | 0.00056497 | 0.005068982 |
| SLC22A15 | 80.3742 | -0.7443 | 0.2771 | -2.6864 | 0.007223489 | 0.037369945 |
| RHOU | 1779.0595 | -0.7446 | 0.1305 | -5.7076 | 1.15E-08 | 5.45E-07 |
| SEPP1 | 206.0020 | -0.7471 | 0.2112 | -3.5371 | 0.000404504 | 0.003885038 |
| KIAA1377 | 291.5929 | -0.7471 | 0.1959 | -3.8140 | 0.000136727 | 0.001631128 |
| MASP2 | 1001.5374 | -0.7485 | 0.1137 | -6.5806 | 4.68E-11 | 3.69E-09 |
| S1PR5 | 115.8939 | -0.7493 | 0.2726 | -2.7482 | 0.005993149 | 0.032412715 |
| CRYBB1 | 139.4621 | -0.7496 | 0.2702 | -2.7749 | 0.005522356 | 0.030565926 |
| PLCD3 | 223.4204 | -0.7513 | 0.2238 | -3.3575 | 0.000786462 | 0.006602006 |
| LOC219347 | 142.3267 | -0.7514 | 0.2351 | -3.1963 | 0.001392217 | 0.010369312 |
| SPON1 | 299.0789 | -0.7515 | 0.1976 | -3.8036 | 0.000142614 | 0.001678371 |
| RCAN3AS | 94.9506 | -0.7559 | 0.2651 | -2.8508 | 0.004361021 | 0.025480561 |
| LINC00426 | 6411.3249 | -0.7604 | 0.1025 | -7.4154 | 1.21E-13 | 1.61E-11 |
| CATSPER3 | 313.3377 | -0.7624 | 0.2193 | -3.4760 | 0.000508946 | 0.004665958 |
| RNF43 | 838.0934 | -0.7626 | 0.1546 | -4.9326 | 8.12E-07 | 2.22E-05 |
| PPM1L | 254.6253 | -0.7629 | 0.1547 | -4.9303 | 8.21E-07 | 2.23E-05 |
| METTL7A | 470.4804 | -0.7700 | 0.2083 | -3.6968 | 0.000218314 | 0.002367337 |
| LOC100505576 | 578.2255 | -0.7701 | 0.1679 | -4.5860 | 4.52E-06 | 9.64E-05 |
| VEPH1 | 385.8186 | -0.7717 | 0.1557 | -4.9572 | 7.15E-07 | 1.98E-05 |
| CARD6 | 578.7960 | -0.7753 | 0.1638 | -4.7335 | 2.21E-06 | 5.16E-05 |
| GNG7 | 162.2456 | -0.7759 | 0.2252 | -3.4449 | 0.000571258 | 0.005109208 |
| KIAA1683 | 338.8445 | -0.7772 | 0.2457 | -3.1630 | 0.001561282 | 0.011360036 |
| CPNE7 | 953.7869 | -0.7794 | 0.1905 | -4.0912 | 4.29E-05 | 0.000628381 |
| AKAP6 | 121.8885 | -0.7795 | 0.2981 | -2.6152 | 0.008918118 | 0.043800122 |
| LOC388692 | 966.2861 | -0.7801 | 0.1493 | -5.2247 | 1.74E-07 | 5.89E-06 |
| CECR1 | 7705.3704 | -0.7817 | 0.1346 | -5.8094 | 6.27E-09 | 3.16E-07 |
| FAM171B | 194.6551 | -0.7833 | 0.2333 | -3.3573 | 0.000787092 | 0.006603626 |
| TCF7 | 3366.4092 | -0.7858 | 0.1498 | -5.2468 | 1.55E-07 | 5.41E-06 |
| FLJ39653 | 616.0392 | -0.7862 | 0.1314 | -5.9854 | 2.16E-09 | 1.20E-07 |
| RLN1 | 81.4435 | -0.7864 | 0.2945 | -2.6699 | 0.007587869 | 0.038829681 |
| ARHGEF11 | 534.8887 | -0.7869 | 0.2073 | -3.7968 | 0.00014657 | 0.001710775 |
| AIM1L | 426.7600 | -0.7871 | 0.2550 | -3.0863 | 0.002026694 | 0.013977466 |
| PLAC9 | 220.2811 | -0.7877 | 0.2035 | -3.8715 | 0.000108182 | 0.001337585 |
| LOC100505783 | 583.8741 | -0.7917 | 0.1355 | -5.8408 | 5.20E-09 | 2.65E-07 |
| TMEM45B | 122.1113 | -0.7939 | 0.2396 | -3.3141 | 0.00091927 | 0.007468207 |
| ZNF763 | 295.8434 | -0.7941 | 0.2162 | -3.6735 | 0.000239245 | 0.002536132 |
| FAM66B | 114.2335 | -0.7952 | 0.2995 | -2.6547 | 0.007936732 | 0.040113244 |
| LOC100506012 | 143.7536 | -0.7961 | 0.2208 | -3.6065 | 0.000310389 | 0.003151174 |
| CTSF | 1399.9336 | -0.7970 | 0.1464 | -5.4454 | 5.17E-08 | 2.07E-06 |
| LOC285847 | 164.3203 | -0.7980 | 0.2563 | -3.1138 | 0.001846727 | 0.0129691 |
| FLJ31813 | 167.8044 | -0.7984 | 0.2770 | -2.8822 | 0.003949034 | 0.023630156 |
| LRRC37A4 | 206.0397 | -0.7993 | 0.2874 | -2.7814 | 0.005412294 | 0.030122065 |
| UBXN11 | 5673.6765 | -0.7993 | 0.1684 | -4.7464 | 2.07E-06 | 4.86E-05 |
| SORL1 | 29250.3409 | -0.7997 | 0.1077 | -7.4262 | 1.12E-13 | 1.50E-11 |
| GRIN2C | 130.1564 | -0.8011 | 0.2342 | -3.4206 | 0.000624861 | 0.005473232 |
| CDHR2 | 415.9769 | -0.8026 | 0.1556 | -5.1577 | 2.50E-07 | 8.13E-06 |
| ABCA10 | 405.3328 | -0.8109 | 0.2976 | -2.7245 | 0.006440536 | 0.034377555 |
| MGC16275 | 334.5111 | -0.8117 | 0.1550 | -5.2368 | 1.63E-07 | 5.64E-06 |
| MORN3 | 298.8747 | -0.8121 | 0.1338 | -6.0707 | 1.27E-09 | 7.47E-08 |
| LOC100506757 | 121.5132 | -0.8124 | 0.2834 | -2.8667 | 0.00414716 | 0.024524665 |
| GSTO2 | 204.2640 | -0.8140 | 0.1779 | -4.5753 | 4.76E-06 | 0.000100755 |
| ALDH3B1 | 49.2995 | -0.8150 | 0.3087 | -2.6402 | 0.00828631 | 0.041462619 |
| CATSPERB | 593.2532 | -0.8153 | 0.1653 | -4.9319 | 8.14E-07 | 2.22E-05 |
| GNAO1 | 298.1105 | -0.8158 | 0.1986 | -4.1078 | 3.99E-05 | 0.000592434 |
| WDR96 | 50.7494 | -0.8158 | 0.3058 | -2.6678 | 0.007634462 | 0.038963057 |
| GDF9 | 91.0334 | -0.8165 | 0.2890 | -2.8253 | 0.004724304 | 0.027214222 |
| RFX8 | 1112.5751 | -0.8201 | 0.1169 | -7.0163 | 2.28E-12 | 2.31E-10 |
| PXDNL | 215.3967 | -0.8223 | 0.3037 | -2.7076 | 0.006777694 | 0.035773265 |
| FCGRT | 500.9682 | -0.8231 | 0.2024 | -4.0664 | 4.77E-05 | 0.000687732 |
| C21orf63 | 1058.3284 | -0.8262 | 0.1547 | -5.3424 | 9.17E-08 | 3.40E-06 |
| TTLL1 | 426.8693 | -0.8310 | 0.1588 | -5.2328 | 1.67E-07 | 5.69E-06 |
| ADAMTS16 | 109.9911 | -0.8339 | 0.2629 | -3.1722 | 0.001512751 | 0.011097665 |
| SPRN | 295.3507 | -0.8340 | 0.2040 | -4.0879 | 4.35E-05 | 0.000636714 |
| LEFTY1 | 92.7898 | -0.8375 | 0.2799 | -2.9920 | 0.002771598 | 0.017865804 |
| TMEM229B | 194.1636 | -0.8389 | 0.2829 | -2.9652 | 0.003024455 | 0.019126646 |
| FMO5 | 358.1378 | -0.8392 | 0.2039 | -4.1154 | 3.86E-05 | 0.00057601 |
| ROBO3 | 197.7363 | -0.8403 | 0.2081 | -4.0382 | 5.39E-05 | 0.00076019 |
| TSPAN32 | 5462.7425 | -0.8450 | 0.1122 | -7.5303 | 5.06E-14 | 7.36E-12 |
| ALK | 97.2097 | -0.8540 | 0.2993 | -2.8530 | 0.004330879 | 0.025353409 |
| CYP4V2 | 1877.4242 | -0.8543 | 0.1394 | -6.1281 | 8.89E-10 | 5.42E-08 |
| SLC13A3 | 211.2880 | -0.8545 | 0.1680 | -5.0877 | 3.62E-07 | 1.11E-05 |
| A2M | 135.5165 | -0.8554 | 0.2697 | -3.1713 | 0.001517501 | 0.011121758 |
| HVCN1 | 574.8727 | -0.8555 | 0.1536 | -5.5686 | 2.57E-08 | 1.14E-06 |
| RASGRP2 | 13954.4540 | -0.8662 | 0.1145 | -7.5626 | 3.95E-14 | 5.92E-12 |
| CUBN | 608.5713 | -0.8741 | 0.1489 | -5.8694 | 4.37E-09 | 2.26E-07 |
| C16orf71 | 116.8281 | -0.8750 | 0.2609 | -3.3536 | 0.000797618 | 0.00667343 |
| CTSO | 442.8746 | -0.8753 | 0.1888 | -4.6363 | 3.55E-06 | 7.73E-05 |
| OGFRL1 | 864.8930 | -0.8763 | 0.1489 | -5.8864 | 3.95E-09 | 2.06E-07 |
| LOC400685 | 178.2734 | -0.8765 | 0.3369 | -2.6018 | 0.009272522 | 0.045101427 |
| NPM2 | 130.2809 | -0.8765 | 0.2892 | -3.0307 | 0.0024401 | 0.016160853 |
| C21orf91-OT1 | 148.5585 | -0.8792 | 0.3123 | -2.8148 | 0.004881368 | 0.027938119 |
| PYGM | 229.6435 | -0.8823 | 0.1724 | -5.1170 | 3.10E-07 | 9.68E-06 |
| CCDC78 | 298.6723 | -0.8833 | 0.1801 | -4.9050 | 9.34E-07 | 2.49E-05 |
| FAM19A2 | 220.5490 | -0.8892 | 0.2357 | -3.7719 | 0.000161992 | 0.00185539 |
| NHSL2 | 605.8815 | -0.8892 | 0.2012 | -4.4204 | 9.85E-06 | 0.000185127 |
| NUAK2 | 361.7663 | -0.8895 | 0.1886 | -4.7153 | 2.41E-06 | 5.57E-05 |
| LOC100132832 | 126.8078 | -0.8899 | 0.3173 | -2.8043 | 0.005043108 | 0.028657813 |
| ZNF664-FAM101A | 76.8625 | -0.8905 | 0.2958 | -3.0104 | 0.002609199 | 0.016954192 |
| C11orf21 | 5810.9765 | -0.8935 | 0.1254 | -7.1254 | 1.04E-12 | 1.15E-10 |
| TTC16 | 199.5269 | -0.8942 | 0.2032 | -4.3998 | 1.08E-05 | 0.000200086 |
| FAM194A | 97.7771 | -0.8952 | 0.2515 | -3.5597 | 0.000371252 | 0.003613367 |
| KIAA0825 | 2815.0777 | -0.8990 | 0.1458 | -6.1683 | 6.90E-10 | 4.25E-08 |
| FBLN5 | 577.7661 | -0.8993 | 0.1869 | -4.8127 | 1.49E-06 | 3.69E-05 |
| CACNA1I | 4980.0042 | -0.9027 | 0.1792 | -5.0376 | 4.71E-07 | 1.39E-05 |
| KIAA1908 | 576.7026 | -0.9057 | 0.1936 | -4.6782 | 2.89E-06 | 6.53E-05 |
| ZNF300 | 102.4238 | -0.9080 | 0.3254 | -2.7903 | 0.005266075 | 0.02953649 |
| TIPARP-AS1 | 85.3867 | -0.9103 | 0.3452 | -2.6373 | 0.008356487 | 0.041691484 |
| LOC285359 | 190.2146 | -0.9122 | 0.2394 | -3.8103 | 0.000138812 | 0.001652087 |
| TMC4 | 373.8605 | -0.9125 | 0.1977 | -4.6165 | 3.90E-06 | 8.45E-05 |
| XPNPEP2 | 152.9783 | -0.9145 | 0.3251 | -2.8127 | 0.00491309 | 0.028035971 |
| FGR | 198.4159 | -0.9195 | 0.2557 | -3.5953 | 0.000323961 | 0.003259574 |
| NMUR1 | 1815.0546 | -0.9229 | 0.1513 | -6.0995 | 1.06E-09 | 6.39E-08 |
| LOC100506207 | 93.7568 | -0.9243 | 0.2564 | -3.6055 | 0.000311595 | 0.0031576 |
| TTYH1 | 276.6905 | -0.9245 | 0.2323 | -3.9803 | 6.88E-05 | 0.000928642 |
| C12orf33 | 121.5076 | -0.9306 | 0.2707 | -3.4374 | 0.000587229 | 0.0052161 |
| WDR86 | 230.3619 | -0.9306 | 0.3153 | -2.9511 | 0.003166029 | 0.019798478 |
| DNAJC28 | 86.7029 | -0.9319 | 0.3542 | -2.6312 | 0.008507437 | 0.042263384 |
| PRR5 | 181.7786 | -0.9331 | 0.2850 | -3.2741 | 0.001059927 | 0.008372597 |
| LOC100129345 | 68.1294 | -0.9344 | 0.2791 | -3.3482 | 0.000813303 | 0.006772327 |
| CFP | 299.6003 | -0.9366 | 0.2402 | -3.8996 | 9.63E-05 | 0.0012225 |
| GML | 85.6307 | -0.9401 | 0.2968 | -3.1678 | 0.00153589 | 0.011223872 |
| MIR548H4 | 172.3758 | -0.9402 | 0.2019 | -4.6559 | 3.23E-06 | 7.13E-05 |
| HPCAL4 | 292.0901 | -0.9406 | 0.2388 | -3.9393 | 8.17E-05 | 0.00107033 |
| LRRC2 | 138.3072 | -0.9429 | 0.2565 | -3.6763 | 0.000236613 | 0.00251528 |
| RTN4R | 66.4637 | -0.9516 | 0.3554 | -2.6778 | 0.007410404 | 0.038219291 |
| ARMC12 | 116.2653 | -0.9560 | 0.2671 | -3.5789 | 0.000345022 | 0.003430708 |
| PDE6G | 493.7344 | -0.9644 | 0.1696 | -5.6860 | 1.30E-08 | 6.09E-07 |
| CTNNA3 | 103.3108 | -0.9690 | 0.2282 | -4.2463 | 2.17E-05 | 0.000354689 |
| NPTXR | 152.0128 | -0.9695 | 0.3251 | -2.9818 | 0.002865487 | 0.01837483 |
| BAIAP3 | 1269.1536 | -0.9701 | 0.1775 | -5.4643 | 4.65E-08 | 1.88E-06 |
| VNN2 | 1194.5388 | -0.9728 | 0.1610 | -6.0432 | 1.51E-09 | 8.69E-08 |
| HLA-DOB | 136.0294 | -0.9762 | 0.3296 | -2.9614 | 0.003062672 | 0.019287696 |
| FAM171A1 | 197.8536 | -0.9768 | 0.2178 | -4.4851 | 7.29E-06 | 0.000143014 |
| ZFP3 | 299.0265 | -0.9778 | 0.2653 | -3.6855 | 0.000228264 | 0.002453132 |
| ZNF583 | 120.1281 | -0.9820 | 0.2851 | -3.4448 | 0.00057148 | 0.005109208 |
| INMT | 114.3792 | -0.9831 | 0.2788 | -3.5254 | 0.000422846 | 0.004029227 |
| USP6 | 45.6427 | -0.9835 | 0.3749 | -2.6230 | 0.00871608 | 0.043053312 |
| D4S234E | 235.8036 | -0.9869 | 0.3054 | -3.2313 | 0.001232376 | 0.009439061 |
| PTGDS | 928.7261 | -0.9909 | 0.3517 | -2.8172 | 0.004843773 | 0.027807118 |
| ZNF396 | 179.4335 | -0.9909 | 0.2086 | -4.7497 | 2.04E-06 | 4.81E-05 |
| TSHR | 255.5123 | -0.9991 | 0.2171 | -4.6014 | 4.20E-06 | 9.03E-05 |
| C9orf116 | 75.1547 | -1.0027 | 0.3164 | -3.1691 | 0.001529237 | 0.011185074 |
| LOC257396 | 74.3629 | -1.0027 | 0.2693 | -3.7236 | 0.000196438 | 0.002181731 |
| C22orf26 | 91.5859 | -1.0066 | 0.3584 | -2.8085 | 0.004977154 | 0.028325589 |
| FXYD7 | 288.3719 | -1.0082 | 0.2359 | -4.2742 | 1.92E-05 | 0.000323175 |
| THBS1 | 416.1256 | -1.0125 | 0.2174 | -4.6579 | 3.19E-06 | 7.09E-05 |
| SERPINF2 | 162.8674 | -1.0161 | 0.3240 | -3.1366 | 0.001709009 | 0.012207055 |
| LY9 | 2509.8423 | -1.0171 | 0.1605 | -6.3362 | 2.35E-10 | 1.59E-08 |
| RAB37 | 12829.1319 | -1.0186 | 0.1394 | -7.3056 | 2.76E-13 | 3.42E-11 |
| CPAMD8 | 133.8520 | -1.0224 | 0.3268 | -3.1283 | 0.001757956 | 0.012473074 |
| FAIM3 | 5461.1657 | -1.0232 | 0.1384 | -7.3904 | 1.46E-13 | 1.91E-11 |
| TLE1 | 136.8523 | -1.0235 | 0.2817 | -3.6336 | 0.000279441 | 0.002889343 |
| CD101 | 609.2949 | -1.0240 | 0.2223 | -4.6070 | 4.09E-06 | 8.80E-05 |
| RHBDL2 | 180.7880 | -1.0253 | 0.2824 | -3.6309 | 0.000282389 | 0.0029079 |
| PDGFD | 154.3925 | -1.0329 | 0.1900 | -5.4361 | 5.45E-08 | 2.17E-06 |
| CCL23 | 73.0626 | -1.0366 | 0.2900 | -3.5745 | 0.000350921 | 0.003474075 |
| URGCP | 108.3003 | -1.0383 | 0.3521 | -2.9490 | 0.003187547 | 0.019908347 |
| IGSF22 | 147.6480 | -1.0414 | 0.2371 | -4.3927 | 1.12E-05 | 0.000205043 |
| FAM178B | 147.8130 | -1.0485 | 0.2631 | -3.9851 | 6.75E-05 | 0.000914458 |
| ST6GALNAC2 | 146.4804 | -1.0562 | 0.2971 | -3.5555 | 0.000377292 | 0.003658527 |
| LDLRAP1 | 1993.7118 | -1.0588 | 0.1466 | -7.2239 | 5.05E-13 | 6.02E-11 |
| GUCY1B2 | 93.1856 | -1.0591 | 0.2608 | -4.0618 | 4.87E-05 | 0.000700337 |
| VIPR1 | 716.4669 | -1.0659 | 0.1813 | -5.8793 | 4.12E-09 | 2.13E-07 |
| BFSP1 | 883.1812 | -1.0694 | 0.1840 | -5.8114 | 6.20E-09 | 3.13E-07 |
| CLEC3B | 36.4179 | -1.0731 | 0.3998 | -2.6843 | 0.007268166 | 0.037562539 |
| EPHA4 | 1023.5544 | -1.0966 | 0.2076 | -5.2815 | 1.28E-07 | 4.54E-06 |
| LHX4 | 114.2089 | -1.1010 | 0.4191 | -2.6268 | 0.008618193 | 0.042743409 |
| LOC643723 | 39.5324 | -1.1057 | 0.4043 | -2.7350 | 0.006238031 | 0.033509479 |
| FCGR3A | 222.0863 | -1.1116 | 0.4040 | -2.7518 | 0.005926517 | 0.032109752 |
| KATNAL2 | 248.9701 | -1.1197 | 0.2359 | -4.7474 | 2.06E-06 | 4.86E-05 |
| NEGR1 | 65.9114 | -1.1198 | 0.4194 | -2.6702 | 0.007580519 | 0.038829681 |
| ZNF713 | 73.4042 | -1.1211 | 0.3449 | -3.2503 | 0.001152639 | 0.00892779 |
| CYP2A7 | 88.0040 | -1.1308 | 0.4081 | -2.7710 | 0.005588505 | 0.030752026 |
| FLJ12825 | 115.9547 | -1.1331 | 0.2707 | -4.1857 | 2.84E-05 | 0.00044294 |
| CLIC3 | 206.4399 | -1.1453 | 0.3623 | -3.1616 | 0.001568882 | 0.011398884 |
| ATP6V1E2 | 142.9488 | -1.1530 | 0.2596 | -4.4417 | 8.92E-06 | 0.000170034 |
| SOAT2 | 299.8195 | -1.1546 | 0.3301 | -3.4976 | 0.000469552 | 0.004377131 |
| NKD1 | 135.3006 | -1.1550 | 0.3523 | -3.2780 | 0.001045331 | 0.008287594 |
| BSN | 138.4545 | -1.1552 | 0.2609 | -4.4278 | 9.52E-06 | 0.000180037 |
| EFHB | 76.8183 | -1.1585 | 0.3898 | -2.9723 | 0.002955836 | 0.0188265 |
| ALG1L2 | 96.9161 | -1.1627 | 0.4304 | -2.7012 | 0.006909454 | 0.036215979 |
| TLCD2 | 133.7506 | -1.1628 | 0.3481 | -3.3402 | 0.000837116 | 0.0069197 |
| DPEP2 | 4605.5621 | -1.1682 | 0.1108 | -10.5473 | 5.23E-26 | 5.65E-23 |
| AIRE | 1146.9042 | -1.1704 | 0.3415 | -3.4267 | 0.000610888 | 0.005366376 |
| NAT8L | 83.4462 | -1.1794 | 0.4198 | -2.8091 | 0.004967686 | 0.028282343 |
| TMEM72-AS1 | 81.1541 | -1.1843 | 0.4397 | -2.6935 | 0.007071013 | 0.036769752 |
| ZNF878 | 93.7097 | -1.1858 | 0.2757 | -4.3014 | 1.70E-05 | 0.000292366 |
| THSD1 | 35.9209 | -1.1942 | 0.4519 | -2.6425 | 0.008228918 | 0.041286513 |
| FGF14 | 91.2387 | -1.2034 | 0.3255 | -3.6970 | 0.00021819 | 0.002367337 |
| FBXO40 | 44.8782 | -1.2254 | 0.4113 | -2.9794 | 0.002888139 | 0.018504398 |
| MYLK4 | 173.2139 | -1.2381 | 0.3125 | -3.9622 | 7.43E-05 | 0.000991503 |
| SLC14A1 | 899.3943 | -1.2778 | 0.2311 | -5.5296 | 3.21E-08 | 1.38E-06 |
| RFPL2 | 107.2017 | -1.2886 | 0.2728 | -4.7229 | 2.33E-06 | 5.40E-05 |
| RFESD | 29.8196 | -1.2951 | 0.4655 | -2.7824 | 0.005395599 | 0.030062332 |
| WBSCR27 | 85.2486 | -1.2955 | 0.3718 | -3.4849 | 0.00049232 | 0.004530018 |
| APBA2 | 83.5031 | -1.3066 | 0.4303 | -3.0367 | 0.002391512 | 0.015922713 |
| SLFN14 | 1935.6836 | -1.3067 | 0.1680 | -7.7763 | 7.47E-15 | 1.23E-12 |
| CTRC | 28.5451 | -1.3101 | 0.5044 | -2.5976 | 0.009386993 | 0.045584927 |
| GALNT9 | 111.8817 | -1.3184 | 0.2429 | -5.4274 | 5.72E-08 | 2.25E-06 |
| ITGB6 | 36.8292 | -1.3191 | 0.4090 | -3.2249 | 0.001260345 | 0.009590159 |
| TSPAN18 | 87.7164 | -1.3378 | 0.3104 | -4.3106 | 1.63E-05 | 0.000282727 |
| AKR1C1 | 76.8932 | -1.3436 | 0.3958 | -3.3950 | 0.000686264 | 0.005905771 |
| GJA9-MYCBP | 77.4828 | -1.3471 | 0.2784 | -4.8389 | 1.31E-06 | 3.31E-05 |
| CALHM1 | 67.8471 | -1.3660 | 0.4747 | -2.8776 | 0.00400716 | 0.023855296 |
| VSIG1 | 574.7427 | -1.3762 | 0.1926 | -7.1439 | 9.07E-13 | 1.02E-10 |
| ICAM4 | 157.0673 | -1.3867 | 0.2783 | -4.9830 | 6.26E-07 | 1.78E-05 |
| CORO7-PAM16 | 100.3308 | -1.3919 | 0.2947 | -4.7230 | 2.32E-06 | 5.40E-05 |
| AQP3 | 2449.5043 | -1.3929 | 0.1897 | -7.3417 | 2.11E-13 | 2.68E-11 |
| SLC22A23 | 98.5851 | -1.4022 | 0.3519 | -3.9845 | 6.76E-05 | 0.000915731 |
| MLC1 | 53.7009 | -1.4066 | 0.4825 | -2.9152 | 0.003554246 | 0.021688215 |
| MSX2P1 | 82.8039 | -1.4081 | 0.3544 | -3.9733 | 7.09E-05 | 0.000953177 |
| RAP1GAP2 | 2701.5013 | -1.4162 | 0.1813 | -7.8125 | 5.60E-15 | 9.53E-13 |
| ARHGEF4 | 91.0523 | -1.4308 | 0.3432 | -4.1686 | 3.06E-05 | 0.000473531 |
| LY75-CD302 | 45.4084 | -1.4350 | 0.5453 | -2.6315 | 0.008501174 | 0.042246145 |
| FAM70A | 84.9650 | -1.4401 | 0.5139 | -2.8021 | 0.005077133 | 0.028797071 |
| FLJ40852 | 133.1501 | -1.4477 | 0.2327 | -6.2203 | 4.96E-10 | 3.13E-08 |
| DTX4 | 128.4628 | -1.4623 | 0.2708 | -5.4007 | 6.64E-08 | 2.56E-06 |
| CDH23 | 252.3134 | -1.4633 | 0.2115 | -6.9183 | 4.57E-12 | 4.38E-10 |
| VWC2L | 27.0932 | -1.4636 | 0.5385 | -2.7180 | 0.006567892 | 0.03489726 |
| FAM19A1 | 108.8541 | -1.4687 | 0.4268 | -3.4408 | 0.000580052 | 0.005170559 |
| CREG2 | 71.9897 | -1.4787 | 0.4458 | -3.3172 | 0.00090933 | 0.007407339 |
| CD248 | 68.4287 | -1.4844 | 0.4895 | -3.0324 | 0.002426528 | 0.016092102 |
| DSCR9 | 39.7330 | -1.5434 | 0.4534 | -3.4038 | 0.000664553 | 0.005777408 |
| AMIGO1 | 39.3455 | -1.5439 | 0.5787 | -2.6678 | 0.007634458 | 0.038963057 |
| CPLX2 | 85.4592 | -1.5831 | 0.4635 | -3.4159 | 0.000635675 | 0.005555087 |
| LINC00264 | 37.7731 | -1.5948 | 0.4472 | -3.5666 | 0.000361628 | 0.003545272 |
| TTC39A | 19.3380 | -1.5989 | 0.5981 | -2.6732 | 0.007512605 | 0.038562329 |
| C19orf18 | 24.7077 | -1.6017 | 0.5497 | -2.9139 | 0.003569713 | 0.021756264 |
| ZP1 | 48.4793 | -1.6138 | 0.5283 | -3.0549 | 0.002251475 | 0.015163875 |
| SUN3 | 32.6080 | -1.6394 | 0.5901 | -2.7780 | 0.005469282 | 0.030338773 |
| OCM | 38.7815 | -1.6435 | 0.6077 | -2.7043 | 0.006845549 | 0.035993263 |
| FLJ42280 | 66.4536 | -1.6520 | 0.4261 | -3.8771 | 0.000105688 | 0.001312591 |
| CCL16 | 40.2935 | -1.6521 | 0.5190 | -3.1833 | 0.00145592 | 0.010753759 |
| CCDC80 | 35.6694 | -1.6596 | 0.5660 | -2.9324 | 0.003363471 | 0.020741631 |
| HSD17B3 | 32.2864 | -1.6704 | 0.6080 | -2.7474 | 0.006006247 | 0.03246034 |
| TBC1D3G | 100.1425 | -1.6813 | 0.4274 | -3.9334 | 8.38E-05 | 0.001086531 |
| LOC284100 | 65.8525 | -1.6928 | 0.4296 | -3.9407 | 8.12E-05 | 0.001065899 |
| LOC100131176 | 325.1694 | -1.7205 | 0.2323 | -7.4063 | 1.30E-13 | 1.71E-11 |
| CLEC18B | 28.9477 | -1.7462 | 0.5854 | -2.9827 | 0.00285706 | 0.018328563 |
| KLKB1 | 32.3069 | -1.7695 | 0.5940 | -2.9788 | 0.002893782 | 0.018532702 |
| TCHH | 25.4725 | -1.7700 | 0.4814 | -3.6768 | 0.000236216 | 0.00251442 |
| VGF | 24.5095 | -1.7724 | 0.5429 | -3.2649 | 0.001095174 | 0.008566024 |
| BST1 | 21.9994 | -1.7880 | 0.6101 | -2.9307 | 0.003381874 | 0.020821167 |
| C11orf41 | 25.1134 | -1.8152 | 0.6975 | -2.6023 | 0.009258961 | 0.045101427 |
| LOC440131 | 281.5135 | -1.8161 | 0.3263 | -5.5665 | 2.60E-08 | 1.15E-06 |
| CXCR1 | 104.3102 | -1.8288 | 0.4215 | -4.3386 | 1.43E-05 | 0.000252827 |
| CMKLR1 | 28.7197 | -1.8383 | 0.6046 | -3.0404 | 0.00236286 | 0.015780568 |
| GLIS1 | 27.5775 | -1.8506 | 0.6711 | -2.7576 | 0.005822308 | 0.031749838 |
| KRT72 | 80.5231 | -1.8589 | 0.4443 | -4.1836 | 2.87E-05 | 0.000446541 |
| HEMGN | 43.0277 | -1.8638 | 0.4933 | -3.7786 | 0.000157701 | 0.001816951 |
| ZNF683 | 1162.0353 | -1.8763 | 0.2358 | -7.9578 | 1.75E-15 | 3.23E-13 |
| CHGB | 31.0627 | -1.9102 | 0.6735 | -2.8363 | 0.004563933 | 0.026451574 |
| FAM176A | 27.4147 | -1.9182 | 0.5909 | -3.2465 | 0.001168424 | 0.00902233 |
| LOC100144603 | 18.9040 | -1.9498 | 0.5833 | -3.3426 | 0.000830049 | 0.006876318 |
| PSCA | 27.0012 | -1.9528 | 0.6677 | -2.9248 | 0.003446702 | 0.021117158 |
| TTLL2 | 24.1075 | -1.9664 | 0.6829 | -2.8794 | 0.003983919 | 0.023780268 |
| MMP28 | 28.9913 | -2.0207 | 0.5110 | -3.9544 | 7.67E-05 | 0.001020725 |
| TYROBP | 28.6562 | -2.0375 | 0.7665 | -2.6581 | 0.007858262 | 0.039791286 |
| RNF222 | 20.6221 | -2.0411 | 0.6854 | -2.9781 | 0.002900185 | 0.018544173 |
| PLLP | 22.1221 | -2.0533 | 0.6621 | -3.1011 | 0.001928051 | 0.013434188 |
| GFPT2 | 1004.5376 | -2.0656 | 0.3380 | -6.1120 | 9.84E-10 | 5.95E-08 |
| LOC729156 | 17.7165 | -2.0822 | 0.6353 | -3.2778 | 0.001046305 | 0.008289975 |
| CES1 | 35.2472 | -2.0831 | 0.6060 | -3.4375 | 0.000587015 | 0.0052161 |
| BMPR1B | 25.1121 | -2.0857 | 0.6097 | -3.4210 | 0.000623929 | 0.005468237 |
| SCUBE2 | 17.0883 | -2.0972 | 0.6952 | -3.0168 | 0.002554594 | 0.016656615 |
| COL28A1 | 12.1784 | -2.1066 | 0.7762 | -2.7140 | 0.00664818 | 0.035212542 |
| KIAA1598 | 16.3214 | -2.1427 | 0.7676 | -2.7914 | 0.005248503 | 0.02945978 |
| LOC147646 | 30.4682 | -2.1782 | 0.6967 | -3.1266 | 0.001768436 | 0.012514926 |
| C16orf74 | 123.4142 | -2.2461 | 0.4063 | -5.5285 | 3.23E-08 | 1.38E-06 |
| HOXA1 | 16.1738 | -2.2971 | 0.8374 | -2.7431 | 0.006085838 | 0.032785065 |
| C1orf115 | 18.1416 | -2.3038 | 0.7232 | -3.1855 | 0.001445287 | 0.010706589 |
| SLC13A5 | 33.7718 | -2.3057 | 0.4028 | -5.7244 | 1.04E-08 | 5.01E-07 |
| PPP1R1C | 34.2597 | -2.3202 | 0.6930 | -3.3480 | 0.000813915 | 0.006772327 |
| PACSIN1 | 111.2498 | -2.3687 | 0.4276 | -5.5393 | 3.04E-08 | 1.32E-06 |
| SYCE3 | 15.8551 | -2.4015 | 0.8653 | -2.7755 | 0.005511679 | 0.030530918 |
| LOC100128714 | 12.6665 | -2.4620 | 0.8639 | -2.8498 | 0.00437508 | 0.025523271 |
| C1orf88 | 10.9899 | -2.4824 | 0.8919 | -2.7833 | 0.005380919 | 0.030009875 |
| CRYBA4 | 22.4112 | -2.5695 | 0.8478 | -3.0309 | 0.002437958 | 0.016157698 |
| C7orf72 | 29.7467 | -2.5763 | 0.7300 | -3.5293 | 0.000416602 | 0.003988573 |
| PTPRR | 13.3975 | -2.6253 | 0.8497 | -3.0898 | 0.002002757 | 0.013858971 |
| DUOX2 | 8.7740 | -2.7695 | 1.0511 | -2.6349 | 0.008416916 | 0.041923836 |
| SPINK2 | 33.4702 | -2.8911 | 0.5985 | -4.8309 | 1.36E-06 | 3.42E-05 |
| CCDC89 | 13.9936 | -2.9742 | 1.1266 | -2.6401 | 0.008288686 | 0.041462619 |
| C10orf55 | 5.6965 | -3.0676 | 1.0371 | -2.9579 | 0.003097745 | NA |
| CR1 | 50.3106 | -3.1604 | 0.6036 | -5.2358 | 1.64E-07 | 5.66E-06 |
| CFHR4 | 12.2538 | -3.1766 | 1.1543 | -2.7519 | 0.005924564 | 0.032109752 |
| RPS4Y2 | 14.4607 | -3.1773 | 1.1091 | -2.8649 | 0.004171976 | 0.024642513 |
| LIPN | 10.2697 | -3.1927 | 1.0612 | -3.0087 | 0.002623598 | 0.017040435 |
| CCL14-CCL15 | 13.3870 | -3.2074 | 0.9248 | -3.4683 | 0.000523842 | 0.00475925 |
| SMTNL2 | 5.9788 | -3.2397 | 1.2188 | -2.6582 | 0.007855764 | NA |
| LOC100129961 | 12.4772 | -3.4904 | 0.9166 | -3.8078 | 0.00014023 | 0.001665764 |
