## Supplementary material for "Marker genes for predicting cytokine release syndrome in vitro before CAR T cell infusion": Table S2

| gene | baseMean | log2FoldChange | lfcSE | stat | pvalue | padj |
| --- | --- | --- | --- | --- | --- | --- |
| HSPA1B | 2291.110 | -2.931 | 0.535 | -5.480 | 4.25E-08 | 0.0006 |
| CA2 | 93.410 | -1.879 | 0.396 | -4.744 | 2.10E-06 | 0.0146 |
| CCDC144B | 675.812 | 1.559 | 0.337 | 4.626 | 3.72E-06 | 0.0172 |
| OSBPL6 | 111.982 | -2.376 | 0.526 | -4.518 | 6.25E-06 | 0.0217 |
| DUSP1 | 342.811 | -1.447 | 0.325 | -4.446 | 8.73E-06 | 0.0242 |
| TCEAL4 | 372.518 | -1.498 | 0.341 | -4.389 | 1.14E-05 | 0.0264 |
| HPCAL4 | 184.526 | 2.224 | 0.514 | 4.322 | 1.54E-05 | 0.0302 |
| SIRPG | 2018.268 | 1.076 | 0.250 | 4.296 | 1.74E-05 | 0.0302 |
| LOC284801 | 149.326 | -1.713 | 0.402 | -4.264 | 2.01E-05 | 0.0309 |
| TCEAL1 | 303.061 | -1.136 | 0.273 | -4.160 | 3.18E-05 | 0.0425 |
| EPB41L2 | 2322.445 | -1.245 | 0.300 | -4.147 | 3.37E-05 | 0.0425 |
| C3orf26 | 812.428 | -1.843 | 0.460 | -4.010 | 6.08E-05 | 0.0691 |
| HSPH1 | 5786.206 | -1.194 | 0.299 | -3.995 | 6.48E-05 | 0.0691 |
| SPRED2 | 136.908 | -1.469 | 0.372 | -3.948 | 7.89E-05 | 0.0705 |
| CYB5A | 1287.696 | -2.195 | 0.557 | -3.944 | 8.02E-05 | 0.0705 |
| IL21 | 332.628 | -2.485 | 0.631 | -3.940 | 8.14E-05 | 0.0705 |
| NR4A1 | 2012.591 | -1.409 | 0.362 | -3.891 | 9.97E-05 | 0.0807 |
| FDX1 | 3843.437 | -2.430 | 0.626 | -3.879 | 0.000104805 | 0.0807 |
| LHFP | 634.453 | -1.932 | 0.502 | -3.847 | 0.000119638 | 0.0873 |
| PCOLCE2 | 45.420 | -2.729 | 0.718 | -3.802 | 0.000143739 | 0.0996 |
| CXorf69 | 52.366 | -1.744 | 0.462 | -3.778 | 0.000158186 | 0.1016 |
| AIG1 | 262.153 | -0.831 | 0.220 | -3.773 | 0.000161203 | 0.1016 |
| ASCL1 | 11.248 | -4.716 | 1.258 | -3.749 | 0.000177306 | NA |
| C5orf32 | 857.073 | -1.697 | 0.454 | -3.736 | 0.000187157 | 0.1128 |
| TCEAL3 | 137.845 | -1.170 | 0.316 | -3.705 | 0.00021163 | 0.1143 |
| PXDC1 | 168.767 | -1.723 | 0.465 | -3.702 | 0.000213703 | 0.1143 |
| MID2 | 394.768 | -1.523 | 0.412 | -3.702 | 0.000214327 | 0.1143 |
| ACN9 | 1080.419 | -1.971 | 0.534 | -3.690 | 0.000224238 | 0.1152 |
| NXT1 | 972.449 | -1.833 | 0.500 | -3.669 | 0.000243634 | 0.1206 |
| DDIT4L | 410.440 | -0.819 | 0.225 | -3.642 | 0.000270861 | 0.1295 |
| DNAJC6 | 381.390 | -1.558 | 0.433 | -3.598 | 0.000320851 | 0.1483 |
| LOC100127983 | 205.365 | -1.851 | 0.521 | -3.554 | 0.000379199 | 0.1696 |
| GPR160 | 1530.341 | -0.974 | 0.276 | -3.535 | 0.000407219 | 0.1706 |
| STK3 | 332.810 | -1.570 | 0.445 | -3.526 | 0.000421808 | 0.1706 |
| ZNF69 | 359.203 | 1.635 | 0.465 | 3.519 | 0.000433323 | 0.1706 |
| BOLA3 | 850.876 | -1.995 | 0.568 | -3.515 | 0.000440219 | 0.1706 |
| GCA | 681.040 | -0.980 | 0.279 | -3.508 | 0.000451441 | 0.1706 |
| SNRPG | 4075.851 | -1.926 | 0.549 | -3.506 | 0.000455142 | 0.1706 |
| IER3 | 2863.436 | -1.831 | 0.524 | -3.497 | 0.000470048 | 0.1715 |
| PFDN2 | 2635.672 | -1.911 | 0.547 | -3.490 | 0.000482913 | 0.1717 |
| AKAP6 | 113.056 | -2.998 | 0.862 | -3.479 | 0.000503724 | 0.1742 |
| CKS2 | 3826.263 | -1.912 | 0.551 | -3.473 | 0.000515265 | 0.1742 |
| CDC42BPB | 133.438 | -1.762 | 0.509 | -3.463 | 0.000534298 | 0.1749 |
| RND1 | 105.884 | -1.523 | 0.441 | -3.452 | 0.000555934 | 0.1749 |
| DZIP1 | 99.482 | -2.678 | 0.776 | -3.449 | 0.000563571 | 0.1749 |
| OSBPL1A | 151.665 | -1.831 | 0.531 | -3.447 | 0.000567645 | 0.1749 |
| SLC35F1 | 624.287 | -2.165 | 0.632 | -3.425 | 0.000614057 | 0.1815 |
| SPRED1 | 5.989 | -6.317 | 1.845 | -3.424 | 0.00061769 | NA |
| SIRPB1 | 176.414 | 1.618 | 0.473 | 3.423 | 0.000619399 | 0.1815 |
| UBE2T | 1810.164 | -1.907 | 0.558 | -3.419 | 0.000628291 | 0.1815 |
| RPL22L1 | 2566.073 | -1.794 | 0.527 | -3.407 | 0.000656447 | 0.1857 |
| RAN | 17093.110 | -1.549 | 0.459 | -3.375 | 0.000737586 | 0.2030 |
| HPRT1 | 5039.375 | -1.835 | 0.545 | -3.369 | 0.000753693 | 0.2030 |
| MYO6 | 899.854 | -1.180 | 0.351 | -3.365 | 0.00076496 | 0.2030 |
| CHPT1 | 1828.282 | -1.385 | 0.413 | -3.355 | 0.000794132 | 0.2030 |
| CMTM8 | 183.901 | -1.683 | 0.502 | -3.353 | 0.000798454 | 0.2030 |
| PTGR1 | 123.305 | -1.644 | 0.491 | -3.350 | 0.00080949 | 0.2030 |
| LOC339822 | 45.432 | 3.861 | 1.154 | 3.345 | 0.000823437 | 0.2030 |
| HMGN5 | 264.728 | -1.972 | 0.590 | -3.341 | 0.000834701 | 0.2030 |
| HMGB2 | 42867.843 | -1.972 | 0.593 | -3.326 | 0.000880138 | 0.2079 |
| CDK2AP1 | 2868.424 | -1.807 | 0.544 | -3.325 | 0.000884476 | 0.2079 |
| TXNDC17 | 1344.613 | -1.397 | 0.423 | -3.306 | 0.000947844 | 0.2138 |
| ULBP2 | 89.051 | -1.776 | 0.537 | -3.304 | 0.000951955 | 0.2138 |
| GSTP1 | 4912.657 | -0.985 | 0.298 | -3.300 | 0.000967343 | 0.2138 |
| SPC25 | 495.267 | -1.264 | 0.384 | -3.295 | 0.000984376 | 0.2138 |
| TXN | 9401.619 | -1.418 | 0.432 | -3.283 | 0.001025505 | 0.2138 |
| GLUL | 14502.920 | -1.126 | 0.343 | -3.281 | 0.001033859 | 0.2138 |
| TRPV1 | 855.708 | 0.783 | 0.239 | 3.277 | 0.001050945 | 0.2138 |
| SNRPF | 3583.549 | -1.682 | 0.514 | -3.270 | 0.001074495 | 0.2138 |
| ENOPH1 | 3955.428 | -1.468 | 0.449 | -3.268 | 0.001081483 | 0.2138 |
| C12orf45 | 451.843 | -1.401 | 0.429 | -3.266 | 0.001091591 | 0.2138 |
| BCAS4 | 1003.706 | -1.458 | 0.447 | -3.262 | 0.001105843 | 0.2138 |
| ZNF439 | 213.539 | 1.517 | 0.465 | 3.262 | 0.001105946 | 0.2138 |
| HNF1B | 712.037 | -3.942 | 1.209 | -3.260 | 0.001112374 | 0.2138 |
| C4orf43 | 2027.533 | -1.794 | 0.551 | -3.255 | 0.001132265 | 0.2138 |
| TMEM38B | 556.440 | -1.727 | 0.531 | -3.252 | 0.001145925 | 0.2138 |
| NUDT11 | 80.736 | -1.972 | 0.607 | -3.247 | 0.001167446 | 0.2138 |
| PYGL | 37.109 | -1.950 | 0.602 | -3.240 | 0.001194476 | 0.2138 |
| MAD2L1 | 4652.794 | -1.812 | 0.559 | -3.239 | 0.001197557 | 0.2138 |
| RBM7 | 3285.404 | -1.617 | 0.499 | -3.238 | 0.00120303 | 0.2138 |
| NLN | 408.476 | -1.660 | 0.515 | -3.225 | 0.001261928 | 0.2215 |
| C14orf109 | 270.971 | -1.696 | 0.527 | -3.220 | 0.001282201 | 0.2222 |
| TCL6 | 11.588 | 4.691 | 1.460 | 3.214 | 0.001309053 | NA |
| ITGA4 | 29844.902 | 0.712 | 0.222 | 3.212 | 0.0013201 | 0.2260 |
| CDKN3 | 1123.186 | -1.722 | 0.538 | -3.198 | 0.001381938 | 0.2337 |
| ALDH3A2 | 1446.155 | 0.851 | 0.267 | 3.190 | 0.001421129 | 0.2361 |
| CDRT15P1 | 201.452 | 1.467 | 0.460 | 3.188 | 0.001430511 | 0.2361 |
| FKBP4 | 3742.327 | -0.880 | 0.277 | -3.178 | 0.001483503 | 0.2370 |
| C6orf125 | 585.835 | -1.222 | 0.385 | -3.177 | 0.001486842 | 0.2370 |
| ANO10 | 611.848 | -0.861 | 0.272 | -3.167 | 0.001538718 | 0.2370 |
| GTF2F2 | 1252.759 | -1.370 | 0.433 | -3.167 | 0.001540078 | 0.2370 |
| VBP1 | 4243.535 | -1.630 | 0.517 | -3.156 | 0.001597261 | 0.2370 |
| DBI | 4497.472 | -1.460 | 0.463 | -3.155 | 0.001602791 | 0.2370 |
| C11orf1 | 289.306 | -0.984 | 0.312 | -3.154 | 0.00161259 | 0.2370 |
| CDC42BPA | 323.572 | -1.284 | 0.407 | -3.153 | 0.001614335 | 0.2370 |
| GRAP | 1073.400 | 0.933 | 0.296 | 3.150 | 0.001630406 | 0.2370 |
| ICAM4 | 84.273 | 1.171 | 0.372 | 3.147 | 0.001652101 | 0.2370 |
| KLHL32 | 106.751 | -1.078 | 0.343 | -3.144 | 0.001663754 | 0.2370 |
| MRPS26 | 822.096 | -1.160 | 0.370 | -3.138 | 0.00170007 | 0.2370 |
| GGCT | 2244.379 | -1.578 | 0.503 | -3.138 | 0.001701437 | 0.2370 |
| GPN3 | 1004.835 | -1.521 | 0.485 | -3.138 | 0.001703359 | 0.2370 |
| C16orf87 | 2719.409 | -1.805 | 0.575 | -3.137 | 0.00170906 | 0.2370 |
| BCL2A1 | 444.888 | -1.685 | 0.538 | -3.131 | 0.001741512 | 0.2370 |
| GFRA2 | 45.240 | 3.849 | 1.229 | 3.130 | 0.001745905 | 0.2370 |
| NUDCD2 | 3175.176 | -1.602 | 0.512 | -3.130 | 0.001750358 | 0.2370 |
| GOLGA6L5 | 341.232 | 1.074 | 0.344 | 3.124 | 0.001784823 | 0.2370 |
| GATA2 | 27.240 | -2.444 | 0.783 | -3.120 | 0.001810684 | 0.2370 |
| RNF165 | 162.762 | 1.248 | 0.400 | 3.117 | 0.001827469 | 0.2370 |
| NAMPT | 35107.888 | -1.447 | 0.465 | -3.112 | 0.001856933 | 0.2370 |
| SMNDC1 | 5702.421 | -1.528 | 0.492 | -3.108 | 0.001884028 | 0.2370 |
| C12orf5 | 1360.215 | -1.113 | 0.358 | -3.107 | 0.001886785 | 0.2370 |
| NR4A2 | 733.188 | -1.124 | 0.362 | -3.106 | 0.001894941 | 0.2370 |
| SAV1 | 697.531 | -0.971 | 0.313 | -3.106 | 0.001897495 | 0.2370 |
| TMEM237 | 1184.190 | -1.661 | 0.535 | -3.102 | 0.00192526 | 0.2370 |
| SPP1 | 24.717 | -3.426 | 1.106 | -3.099 | 0.001942593 | 0.2370 |
| MZB1 | 1833.140 | 1.031 | 0.333 | 3.097 | 0.00195694 | 0.2370 |
| LRRCC1 | 870.650 | -1.533 | 0.495 | -3.096 | 0.001960289 | 0.2370 |
| PCDH11X | 107.458 | -1.815 | 0.586 | -3.095 | 0.001965742 | 0.2370 |
| PASK | 8546.840 | 0.596 | 0.193 | 3.087 | 0.002019539 | 0.2414 |
| LAMC1 | 45.970 | -1.549 | 0.504 | -3.073 | 0.002116539 | 0.2485 |
| NDUFC1 | 1543.351 | -0.661 | 0.215 | -3.068 | 0.002153035 | 0.2485 |
| NPM3 | 744.628 | -1.742 | 0.568 | -3.068 | 0.002157525 | 0.2485 |
| BRI3 | 2328.552 | -1.455 | 0.475 | -3.065 | 0.002176023 | 0.2485 |
| COX17 | 2378.038 | -1.457 | 0.476 | -3.065 | 0.002179635 | 0.2485 |
| CKLF-CMTM1 | 123.849 | 0.887 | 0.289 | 3.064 | 0.002186492 | 0.2485 |
| PDCD10 | 6166.894 | -1.519 | 0.497 | -3.054 | 0.002260926 | 0.2525 |
| GATM | 85.292 | -1.575 | 0.516 | -3.052 | 0.002276411 | 0.2525 |
| COQ2 | 410.033 | -0.776 | 0.254 | -3.052 | 0.002276773 | 0.2525 |
| STMN1 | 11269.578 | -1.407 | 0.462 | -3.047 | 0.002312592 | 0.2536 |
| LSM6 | 2117.551 | -1.372 | 0.450 | -3.046 | 0.002322601 | 0.2536 |
| PCBD1 | 1222.678 | -0.885 | 0.291 | -3.043 | 0.002343711 | 0.2539 |
| CISD3 | 770.242 | -1.136 | 0.374 | -3.036 | 0.002394851 | 0.2568 |
| CCDC85C | 496.230 | 0.977 | 0.322 | 3.034 | 0.002411857 | 0.2568 |
| CHN1 | 409.097 | -1.569 | 0.518 | -3.029 | 0.00245572 | 0.2568 |
| TRIP10 | 2684.894 | -1.064 | 0.351 | -3.028 | 0.002460469 | 0.2568 |
| CCDC6 | 5576.749 | -1.162 | 0.384 | -3.027 | 0.002466243 | 0.2568 |
| HLX | 146.130 | 1.052 | 0.348 | 3.024 | 0.002494271 | 0.2568 |
| SKAP2 | 1595.339 | -0.866 | 0.286 | -3.023 | 0.002500881 | 0.2568 |
| SAP18 | 5055.404 | -1.389 | 0.461 | -3.015 | 0.002571086 | 0.2608 |
| NUDT8 | 209.478 | -1.088 | 0.361 | -3.011 | 0.002602868 | 0.2608 |
| SBDSP1 | 2438.452 | -1.672 | 0.556 | -3.008 | 0.002627121 | 0.2608 |
| TPMT | 945.734 | -0.701 | 0.233 | -3.008 | 0.002632017 | 0.2608 |
| SAP30 | 6084.719 | -1.633 | 0.543 | -3.007 | 0.002640876 | 0.2608 |
| TMEM51 | 6.433 | -5.611 | 1.866 | -3.007 | 0.002641378 | NA |
| SPATA5 | 673.292 | -1.096 | 0.365 | -3.005 | 0.002659383 | 0.2608 |
| PDLIM5 | 857.531 | -0.905 | 0.301 | -3.001 | 0.00268966 | 0.2608 |
| IL2 | 177.632 | -2.553 | 0.851 | -3.001 | 0.002690123 | 0.2608 |
| RIMKLA | 1304.437 | -0.662 | 0.221 | -2.997 | 0.002730882 | 0.2615 |
| MZT1 | 3957.391 | -1.735 | 0.579 | -2.996 | 0.002734365 | 0.2615 |
| BATF3 | 2067.465 | -1.337 | 0.447 | -2.988 | 0.002809796 | 0.2624 |
| COX6C | 4096.251 | -1.189 | 0.398 | -2.987 | 0.002815507 | 0.2624 |
| C3orf14 | 221.630 | -1.246 | 0.417 | -2.985 | 0.002834586 | 0.2624 |
| 10-Sep | 78.851 | -1.961 | 0.657 | -2.982 | 0.002864042 | 0.2624 |
| NAA10 | 1503.467 | -0.908 | 0.305 | -2.981 | 0.002871888 | 0.2624 |
| B4GALT4 | 1538.140 | -1.014 | 0.340 | -2.981 | 0.002873617 | 0.2624 |
| GBA3 | 112.064 | -1.795 | 0.602 | -2.980 | 0.002886158 | 0.2624 |
| CACYBP | 7503.166 | -1.162 | 0.391 | -2.976 | 0.002923352 | 0.2624 |
| GRAPL | 66.689 | 1.661 | 0.559 | 2.970 | 0.002979035 | 0.2624 |
| MRPL19 | 3063.345 | -1.201 | 0.405 | -2.968 | 0.002996354 | 0.2624 |
| LZTS1 | 521.284 | 1.838 | 0.621 | 2.962 | 0.00305349 | 0.2624 |
| C14orf2 | 2370.295 | -1.173 | 0.396 | -2.959 | 0.003081469 | 0.2624 |
| TAOK3 | 18702.608 | -1.175 | 0.397 | -2.957 | 0.003107688 | 0.2624 |
| LSM3 | 1638.176 | -1.201 | 0.406 | -2.956 | 0.003118356 | 0.2624 |
| DOCK1 | 35.963 | -2.800 | 0.947 | -2.955 | 0.003122321 | 0.2624 |
| FOS | 94.367 | -1.578 | 0.534 | -2.954 | 0.003140407 | 0.2624 |
| BZW1 | 19241.132 | -0.982 | 0.333 | -2.953 | 0.003151625 | 0.2624 |
| STRA6 | 7.397 | -6.191 | 2.097 | -2.952 | 0.003157641 | NA |
| C1orf135 | 137.637 | -1.737 | 0.588 | -2.952 | 0.003162103 | 0.2624 |
| HSPD1 | 18056.806 | -1.108 | 0.375 | -2.951 | 0.003166234 | 0.2624 |
| MKRN3 | 254.504 | 1.288 | 0.437 | 2.949 | 0.003191627 | 0.2624 |
| ING2 | 1505.788 | -1.455 | 0.494 | -2.947 | 0.003213224 | 0.2624 |
| TOMM5 | 1014.413 | -1.376 | 0.467 | -2.946 | 0.003217509 | 0.2624 |
| GUSBP3 | 124.277 | 2.385 | 0.810 | 2.945 | 0.00322456 | 0.2624 |
| CYB5RL | 746.303 | 0.563 | 0.191 | 2.944 | 0.003239961 | 0.2624 |
| MND1 | 585.042 | -1.275 | 0.433 | -2.944 | 0.003245021 | 0.2624 |
| VDAC1 | 23980.906 | -1.300 | 0.442 | -2.941 | 0.003267744 | 0.2624 |
| RHEB | 4944.709 | -1.571 | 0.534 | -2.939 | 0.003293789 | 0.2624 |
| NENF | 1317.027 | -0.834 | 0.284 | -2.939 | 0.003296133 | 0.2624 |
| ATP5G1 | 1142.722 | -1.152 | 0.392 | -2.937 | 0.003315473 | 0.2624 |
| SPDYE5 | 231.385 | 0.894 | 0.305 | 2.935 | 0.003333195 | 0.2624 |
| ST7 | 493.915 | -0.836 | 0.285 | -2.934 | 0.003350156 | 0.2624 |
| MTHFD2 | 8915.412 | -1.431 | 0.488 | -2.931 | 0.003374313 | 0.2624 |
| TYMS | 6413.463 | -1.500 | 0.512 | -2.930 | 0.003390946 | 0.2624 |
| GLRX | 3304.156 | -1.086 | 0.371 | -2.928 | 0.003409631 | 0.2624 |
| DNAJC12 | 426.682 | -2.113 | 0.722 | -2.926 | 0.003436046 | 0.2624 |
| BRP44 | 3354.333 | -1.408 | 0.481 | -2.925 | 0.003442543 | 0.2624 |
| MLF1 | 138.725 | -1.578 | 0.540 | -2.923 | 0.003469947 | 0.2624 |
| THBS4 | 404.853 | 1.121 | 0.383 | 2.923 | 0.003470179 | 0.2624 |
| PERP | 4128.325 | -1.180 | 0.404 | -2.922 | 0.003482007 | 0.2624 |
| NRTN | 12.844 | -2.326 | 0.796 | -2.921 | 0.003483569 | NA |
| RNF217 | 70.028 | -2.469 | 0.847 | -2.916 | 0.00354275 | 0.2641 |
| CXCR7 | 89.054 | 1.206 | 0.414 | 2.916 | 0.003543437 | 0.2641 |
| RMI2 | 1001.574 | -0.899 | 0.309 | -2.913 | 0.003583668 | 0.2645 |
| SRSF3 | 16055.266 | -1.293 | 0.444 | -2.912 | 0.003585966 | 0.2645 |
| SLC48A1 | 1252.106 | -1.334 | 0.459 | -2.903 | 0.003692774 | 0.2669 |
| NLRP6 | 552.715 | 1.481 | 0.511 | 2.901 | 0.003725357 | 0.2669 |
| DDX21 | 7232.614 | -1.246 | 0.430 | -2.900 | 0.003736296 | 0.2669 |
| MTCH2 | 2732.276 | -0.945 | 0.326 | -2.896 | 0.003775125 | 0.2669 |
| TEC | 612.752 | -1.326 | 0.458 | -2.894 | 0.003804222 | 0.2669 |
| DEK | 19014.610 | -1.401 | 0.484 | -2.894 | 0.003808665 | 0.2669 |
| PPA1 | 9670.574 | -1.389 | 0.480 | -2.892 | 0.003829178 | 0.2669 |
| CALD1 | 179.191 | -1.697 | 0.587 | -2.891 | 0.003842048 | 0.2669 |
| C12orf39 | 74.040 | -1.760 | 0.609 | -2.888 | 0.00387502 | 0.2669 |
| ATP5G3 | 8040.754 | -1.159 | 0.402 | -2.884 | 0.003928989 | 0.2669 |
| AZIN1 | 12602.052 | -1.385 | 0.480 | -2.884 | 0.003930954 | 0.2669 |
| WWC2 | 16.257 | -2.497 | 0.866 | -2.883 | 0.003933331 | 0.2669 |
| TNFRSF12A | 417.747 | -1.191 | 0.413 | -2.881 | 0.00396649 | 0.2669 |
| UQCRHL | 414.870 | -1.332 | 0.463 | -2.879 | 0.003995302 | 0.2669 |
| HBXIP | 5377.770 | -1.512 | 0.525 | -2.878 | 0.003998983 | 0.2669 |
| DSTNP2 | 456.627 | -0.773 | 0.268 | -2.878 | 0.004000312 | 0.2669 |
| YARS2 | 848.347 | -0.689 | 0.239 | -2.878 | 0.004000493 | 0.2669 |
| MIIP | 2661.625 | -0.744 | 0.259 | -2.878 | 0.004006784 | 0.2669 |
| SPON2 | 1257.873 | 1.134 | 0.394 | 2.878 | 0.004007132 | 0.2669 |
| COX6A1 | 3520.542 | -1.307 | 0.454 | -2.876 | 0.004023042 | 0.2669 |
| SRM | 1636.271 | -1.123 | 0.391 | -2.875 | 0.004045129 | 0.2671 |
| DYNLT3 | 1785.207 | -1.261 | 0.439 | -2.871 | 0.004094759 | 0.2681 |
| CCDC41 | 1108.472 | -1.002 | 0.349 | -2.869 | 0.004124179 | 0.2681 |
| COPG2 | 867.916 | -0.619 | 0.216 | -2.868 | 0.004127793 | 0.2681 |
| ELFN2 | 201.663 | 1.182 | 0.412 | 2.867 | 0.004137574 | 0.2681 |
| RRM2 | 14020.818 | -1.476 | 0.516 | -2.862 | 0.004211907 | 0.2690 |
| ANK2 | 223.770 | -1.186 | 0.414 | -2.862 | 0.004214575 | 0.2690 |
| BPGM | 1466.567 | -1.081 | 0.378 | -2.858 | 0.004262291 | 0.2690 |
| CCDC86 | 898.871 | -0.862 | 0.302 | -2.858 | 0.004268898 | 0.2690 |
| BMP1 | 1629.783 | 0.849 | 0.297 | 2.857 | 0.004275096 | 0.2690 |
| HMGB3 | 862.711 | -1.225 | 0.429 | -2.857 | 0.004275561 | 0.2690 |
| UBE2V2 | 3722.276 | -1.337 | 0.468 | -2.855 | 0.004299791 | 0.2690 |
| PSMA7 | 6136.545 | -1.132 | 0.396 | -2.855 | 0.004306808 | 0.2690 |
| ELL2 | 19945.395 | -1.325 | 0.464 | -2.853 | 0.004332627 | 0.2694 |
| NDUFB3 | 1817.126 | -1.245 | 0.437 | -2.850 | 0.004373287 | 0.2707 |
| TMSB4X | 205491.405 | -1.249 | 0.438 | -2.848 | 0.0043948 | 0.2708 |
| RFX8 | 762.264 | 1.040 | 0.365 | 2.845 | 0.00443514 | 0.2715 |
| RPL39L | 418.019 | -1.031 | 0.363 | -2.844 | 0.004449466 | 0.2715 |
| CENPW | 593.228 | -1.604 | 0.564 | -2.843 | 0.004464238 | 0.2715 |
| BRIX1 | 1716.279 | -1.370 | 0.483 | -2.838 | 0.004534684 | 0.2745 |
| PGAP3 | 1230.032 | 0.784 | 0.277 | 2.837 | 0.004554286 | 0.2745 |
| BAG2 | 981.918 | -1.027 | 0.362 | -2.835 | 0.004589383 | 0.2755 |
| ASPH | 568.434 | -0.687 | 0.243 | -2.832 | 0.004628437 | 0.2762 |
| YBX1 | 34886.135 | -1.332 | 0.471 | -2.831 | 0.004641886 | 0.2762 |
| TAF3 | 1685.523 | -1.047 | 0.371 | -2.826 | 0.004714432 | 0.2787 |
| ADK | 3630.262 | -1.155 | 0.409 | -2.823 | 0.004762236 | 0.2787 |
| UQCRQ | 1924.714 | -1.088 | 0.386 | -2.821 | 0.004793561 | 0.2787 |
| GGH | 820.933 | -1.069 | 0.379 | -2.820 | 0.004799094 | 0.2787 |
| CELA2B | 329.503 | 0.782 | 0.277 | 2.818 | 0.004829655 | 0.2787 |
| USMG5 | 2566.283 | -1.137 | 0.404 | -2.817 | 0.004843935 | 0.2787 |
| KIF21A | 2206.581 | -0.521 | 0.185 | -2.817 | 0.004847559 | 0.2787 |
| TMEM191B | 42.457 | 2.013 | 0.716 | 2.812 | 0.004920235 | 0.2787 |
| AOAH | 8861.821 | 0.887 | 0.315 | 2.812 | 0.004921851 | 0.2787 |
| CYTIP | 39265.551 | -1.358 | 0.483 | -2.812 | 0.00492957 | 0.2787 |
| CDCA7 | 9372.386 | -1.656 | 0.589 | -2.811 | 0.004937588 | 0.2787 |
| HSPA2 | 251.461 | -1.010 | 0.360 | -2.807 | 0.005007091 | 0.2787 |
| KIAA1875 | 193.674 | 1.584 | 0.564 | 2.806 | 0.005009438 | 0.2787 |
| PGCP | 494.164 | -0.906 | 0.323 | -2.804 | 0.005040524 | 0.2787 |
| MRPL3 | 3321.868 | -1.054 | 0.376 | -2.802 | 0.005072848 | 0.2787 |
| POLN | 905.561 | 1.057 | 0.377 | 2.801 | 0.005090077 | 0.2787 |
| RPL26L1 | 780.268 | -1.339 | 0.478 | -2.800 | 0.005107794 | 0.2787 |
| CDCA7L | 1881.958 | -0.768 | 0.274 | -2.799 | 0.005120808 | 0.2787 |
| FLJ43860 | 159.532 | -1.749 | 0.625 | -2.799 | 0.005126622 | 0.2787 |
| NDUFB9 | 3987.124 | -1.316 | 0.470 | -2.798 | 0.005135066 | 0.2787 |
| PEA15 | 2857.390 | -1.274 | 0.456 | -2.795 | 0.00518826 | 0.2787 |
| CKS1B | 2495.213 | -1.468 | 0.526 | -2.791 | 0.005252952 | 0.2787 |
| SPAG1 | 1284.101 | -1.328 | 0.476 | -2.791 | 0.005258095 | 0.2787 |
| EIF5B | 8516.087 | -1.183 | 0.424 | -2.790 | 0.005269869 | 0.2787 |
| PHF5A | 1129.546 | -1.265 | 0.454 | -2.788 | 0.005304375 | 0.2787 |
| LOC100505483 | 114.509 | -1.307 | 0.469 | -2.787 | 0.005315377 | 0.2787 |
| ATP5J | 2693.591 | -1.102 | 0.395 | -2.787 | 0.005323124 | 0.2787 |
| C17orf58 | 1109.464 | -1.362 | 0.489 | -2.786 | 0.005338738 | 0.2787 |
| PSMB3 | 3140.642 | -0.920 | 0.330 | -2.786 | 0.005341164 | 0.2787 |
| MBOAT2 | 210.524 | -0.936 | 0.336 | -2.782 | 0.00539492 | 0.2787 |
| STRAP | 5450.760 | -1.261 | 0.453 | -2.782 | 0.005399069 | 0.2787 |
| SLC25A5 | 14553.308 | -1.374 | 0.494 | -2.782 | 0.005409754 | 0.2787 |
| MTERFD2 | 2449.483 | 0.630 | 0.227 | 2.777 | 0.005483089 | 0.2787 |
| PPP1R2 | 11964.765 | -1.305 | 0.470 | -2.777 | 0.005484329 | 0.2787 |
| PDCD2L | 241.364 | -1.161 | 0.418 | -2.777 | 0.005487781 | 0.2787 |
| IPO5 | 9061.089 | -1.229 | 0.443 | -2.775 | 0.005523049 | 0.2787 |
| CSRP2 | 97.420 | -2.666 | 0.961 | -2.775 | 0.005527524 | 0.2787 |
| SNHG8 | 1143.175 | -1.406 | 0.507 | -2.774 | 0.005529363 | 0.2787 |
| CMPK1 | 15831.484 | -1.297 | 0.468 | -2.773 | 0.005554271 | 0.2787 |
| DIAPH2 | 1045.949 | -0.957 | 0.345 | -2.772 | 0.005565168 | 0.2787 |
| PAK1IP1 | 540.362 | -1.306 | 0.471 | -2.772 | 0.005574419 | 0.2787 |
| SHISA6 | 6.332 | -3.606 | 1.301 | -2.771 | 0.005589521 | NA |
| NECAB1 | 173.714 | -1.086 | 0.392 | -2.771 | 0.005589879 | 0.2787 |
| PRRG4 | 337.517 | -0.832 | 0.301 | -2.769 | 0.005614664 | 0.2787 |
| FAM54A | 976.537 | -1.059 | 0.382 | -2.769 | 0.005630709 | 0.2787 |
| FHOD3 | 569.409 | 0.551 | 0.199 | 2.768 | 0.00563252 | 0.2787 |
| CCNE1 | 602.838 | -0.865 | 0.313 | -2.767 | 0.005649961 | 0.2787 |
| PFDN4 | 2083.330 | -1.380 | 0.500 | -2.763 | 0.005727773 | 0.2787 |
| SLC16A5 | 916.799 | 0.869 | 0.315 | 2.762 | 0.005743082 | 0.2787 |
| NUDT1 | 1022.944 | -1.140 | 0.413 | -2.762 | 0.005752814 | 0.2787 |
| CHCHD10 | 630.000 | -0.870 | 0.315 | -2.761 | 0.005760807 | 0.2787 |
| RDX | 8934.876 | -1.288 | 0.467 | -2.760 | 0.005780069 | 0.2787 |
| C16orf61 | 2778.117 | -0.855 | 0.310 | -2.758 | 0.0058153 | 0.2787 |
| C10orf35 | 176.814 | -1.758 | 0.638 | -2.758 | 0.005823837 | 0.2787 |
| OBFC2B | 1912.926 | -0.877 | 0.318 | -2.755 | 0.005866858 | 0.2787 |
| NDUFAF2 | 541.302 | -1.054 | 0.382 | -2.755 | 0.00586718 | 0.2787 |
| GNG12 | 60.165 | -2.125 | 0.772 | -2.753 | 0.005902946 | 0.2787 |
| C2orf49 | 1670.606 | -1.078 | 0.392 | -2.751 | 0.005937163 | 0.2787 |
| DNAJB1 | 3961.338 | -0.692 | 0.252 | -2.749 | 0.005980141 | 0.2787 |
| PPP2R2B | 266.369 | -1.153 | 0.419 | -2.748 | 0.005998455 | 0.2787 |
| UCHL3 | 1107.077 | -1.057 | 0.385 | -2.744 | 0.006071866 | 0.2787 |
| RFC3 | 2515.309 | -1.083 | 0.395 | -2.743 | 0.006092204 | 0.2787 |
| ZNF367 | 1438.080 | -0.756 | 0.276 | -2.742 | 0.006104301 | 0.2787 |
| MIR155HG | 3598.847 | -1.205 | 0.440 | -2.740 | 0.006152035 | 0.2787 |
| PDE6A | 12.422 | -2.781 | 1.015 | -2.739 | 0.00615664 | NA |
| HAT1 | 1945.356 | -0.691 | 0.252 | -2.739 | 0.006164104 | 0.2787 |
| ISCU | 6451.559 | -1.281 | 0.468 | -2.739 | 0.006171404 | 0.2787 |
| AKIP1 | 939.523 | -0.890 | 0.325 | -2.738 | 0.006174856 | 0.2787 |
| ATP1B1 | 412.168 | -0.900 | 0.329 | -2.738 | 0.006178627 | 0.2787 |
| CHCHD3 | 3623.597 | -1.267 | 0.463 | -2.737 | 0.006203298 | 0.2787 |
| ITPKA | 292.474 | -1.982 | 0.724 | -2.736 | 0.00621122 | 0.2787 |
| C17orf89 | 586.878 | -1.034 | 0.378 | -2.736 | 0.006219559 | 0.2787 |
| ACPT | 11.030 | 2.707 | 0.990 | 2.734 | 0.006248481 | NA |
| POLR2L | 769.975 | -1.083 | 0.396 | -2.734 | 0.006258939 | 0.2787 |
| SAT1 | 7765.057 | -1.201 | 0.439 | -2.733 | 0.006282799 | 0.2787 |
| MOK | 115.809 | -0.936 | 0.343 | -2.732 | 0.006289553 | 0.2787 |
| GPX1 | 2431.901 | -0.735 | 0.269 | -2.730 | 0.006340418 | 0.2787 |
| PEX6 | 3179.916 | 0.671 | 0.246 | 2.729 | 0.006354505 | 0.2787 |
| HELLS | 1812.295 | -0.878 | 0.322 | -2.729 | 0.006357428 | 0.2787 |
| FAM174B | 347.589 | -1.317 | 0.483 | -2.728 | 0.006362796 | 0.2787 |
| COX5A | 5363.540 | -1.091 | 0.400 | -2.728 | 0.006365166 | 0.2787 |
| MIR4786 | 5.854 | 3.471 | 1.273 | 2.727 | 0.006393479 | NA |
| WASF1 | 275.119 | -1.058 | 0.388 | -2.727 | 0.006397254 | 0.2787 |
| DUT | 7007.950 | -1.181 | 0.434 | -2.723 | 0.006463944 | 0.2787 |
| ZNF844 | 62.188 | 1.557 | 0.572 | 2.723 | 0.006471808 | 0.2787 |
| GINS2 | 946.539 | -1.166 | 0.429 | -2.722 | 0.006491629 | 0.2787 |
| TMEM106C | 4461.673 | -1.059 | 0.389 | -2.721 | 0.006499844 | 0.2787 |
| FLJ35282 | 6.411 | -3.568 | 1.311 | -2.721 | 0.006507243 | NA |
| DAPK2 | 4490.861 | 1.065 | 0.392 | 2.719 | 0.006542572 | 0.2787 |
| KREMEN1 | 17.619 | -2.950 | 1.085 | -2.719 | 0.006553952 | 0.2787 |
| LSM5 | 2174.965 | -1.095 | 0.403 | -2.718 | 0.006573942 | 0.2787 |
| SRP14 | 10248.647 | -1.078 | 0.397 | -2.717 | 0.006583207 | 0.2787 |
| PSMB7 | 2725.488 | -0.847 | 0.312 | -2.717 | 0.006587019 | 0.2787 |
| CCDC124 | 1750.801 | -0.737 | 0.271 | -2.716 | 0.006609758 | 0.2787 |
| RGS2 | 1665.410 | -1.117 | 0.411 | -2.716 | 0.006615988 | 0.2787 |
| RPSAP9 | 78.905 | -1.625 | 0.599 | -2.714 | 0.006638309 | 0.2787 |
| ATP1B3 | 6816.042 | -0.749 | 0.276 | -2.714 | 0.006644767 | 0.2787 |
| LPAL2 | 433.328 | 1.122 | 0.414 | 2.714 | 0.006654828 | 0.2787 |
| POMP | 4360.049 | -1.000 | 0.369 | -2.713 | 0.006660893 | 0.2787 |
| LINC00161 | 11.185 | 3.687 | 1.359 | 2.712 | 0.006687992 | NA |
| PARK7 | 6562.916 | -0.950 | 0.350 | -2.711 | 0.006698504 | 0.2787 |
| PPARG | 1237.568 | -1.730 | 0.638 | -2.711 | 0.006699192 | 0.2787 |
| JUNB | 11520.216 | -0.974 | 0.359 | -2.711 | 0.006701766 | 0.2787 |
| TUBA1B | 13990.902 | -1.043 | 0.385 | -2.711 | 0.006707449 | 0.2787 |
| ACOT7 | 1428.497 | -1.165 | 0.430 | -2.711 | 0.006716615 | 0.2787 |
| TIPIN | 528.399 | -1.245 | 0.460 | -2.708 | 0.00675959 | 0.2787 |
| CEP55 | 4671.045 | -1.299 | 0.480 | -2.707 | 0.006792612 | 0.2787 |
| NDUFAF4 | 408.536 | -1.027 | 0.379 | -2.707 | 0.006797881 | 0.2787 |
| NETO1 | 5.044 | -4.999 | 1.847 | -2.706 | 0.00680988 | NA |
| FAM72D | 406.085 | -1.024 | 0.379 | -2.705 | 0.006840343 | 0.2787 |
| TESC | 964.804 | -0.936 | 0.346 | -2.704 | 0.006841535 | 0.2787 |
| IFT57 | 764.423 | -0.679 | 0.251 | -2.702 | 0.006902555 | 0.2787 |
| ESCO2 | 784.645 | -1.200 | 0.444 | -2.700 | 0.006926813 | 0.2787 |
| IER3IP1 | 2477.459 | -1.199 | 0.444 | -2.700 | 0.006930059 | 0.2787 |
| RPL12 | 40325.157 | -0.986 | 0.365 | -2.700 | 0.006932022 | 0.2787 |
| C14orf166 | 6634.389 | -1.096 | 0.406 | -2.698 | 0.00697286 | 0.2787 |
| PTPN14 | 715.204 | -1.558 | 0.577 | -2.698 | 0.006977144 | 0.2787 |
| TNFAIP8 | 7399.856 | -0.927 | 0.344 | -2.696 | 0.00701838 | 0.2787 |
| PSMG3 | 632.124 | -0.990 | 0.367 | -2.695 | 0.007034912 | 0.2787 |
| MRPL11 | 1660.526 | -1.004 | 0.373 | -2.694 | 0.007061293 | 0.2787 |
| COX7A2 | 3820.674 | -1.012 | 0.376 | -2.693 | 0.007089485 | 0.2787 |
| AHSA1 | 2810.911 | -0.752 | 0.279 | -2.691 | 0.007122417 | 0.2787 |
| C3orf33 | 201.918 | -1.265 | 0.470 | -2.690 | 0.007136508 | 0.2787 |
| SMS | 5432.939 | -1.164 | 0.433 | -2.689 | 0.007156244 | 0.2787 |
| CCNA2 | 3866.460 | -1.272 | 0.473 | -2.689 | 0.007175998 | 0.2787 |
| PTPRG | 232.117 | -1.154 | 0.430 | -2.687 | 0.007212407 | 0.2787 |
| DSEL | 18.203 | -3.491 | 1.299 | -2.687 | 0.007217246 | 0.2787 |
| MTHFD1L | 3529.981 | -1.014 | 0.378 | -2.686 | 0.007239776 | 0.2787 |
| TCOF1 | 3470.098 | -1.099 | 0.409 | -2.685 | 0.007243978 | 0.2787 |
| METTL2A | 592.513 | -0.564 | 0.210 | -2.685 | 0.007248577 | 0.2787 |
| NPM1 | 56518.167 | -1.080 | 0.402 | -2.685 | 0.007250652 | 0.2787 |
| ACE | 503.034 | 1.732 | 0.645 | 2.685 | 0.007255845 | 0.2787 |
| HSPA4L | 197.057 | -1.307 | 0.487 | -2.682 | 0.007315785 | 0.2787 |
| NDUFA1 | 3370.469 | -0.993 | 0.370 | -2.681 | 0.00732968 | 0.2787 |
| CCT5 | 5782.247 | -0.876 | 0.327 | -2.679 | 0.007383185 | 0.2787 |
| CD93 | 269.205 | 1.451 | 0.542 | 2.678 | 0.007410424 | 0.2787 |
| NAALADL2 | 36.415 | -1.993 | 0.744 | -2.677 | 0.007418473 | 0.2787 |
| MRPS35 | 2381.695 | -0.981 | 0.366 | -2.677 | 0.007421554 | 0.2787 |
| STARD3NL | 2792.453 | -1.251 | 0.467 | -2.677 | 0.007431825 | 0.2787 |
| WDR76 | 1865.934 | -1.006 | 0.376 | -2.675 | 0.007462272 | 0.2787 |
| SOD1 | 7056.439 | -1.031 | 0.385 | -2.675 | 0.00747759 | 0.2787 |
| CDK8 | 1343.908 | -0.741 | 0.277 | -2.675 | 0.007478033 | 0.2787 |
| NAP1L1 | 62033.465 | -1.039 | 0.389 | -2.673 | 0.007524482 | 0.2787 |
| ZP3 | 496.681 | 0.842 | 0.315 | 2.672 | 0.007533414 | 0.2787 |
| HSP90AA1 | 30735.568 | -1.014 | 0.380 | -2.671 | 0.007553314 | 0.2787 |
| CCT7 | 8970.562 | -0.842 | 0.316 | -2.669 | 0.007607521 | 0.2787 |
| C15orf24 | 3104.157 | -1.078 | 0.404 | -2.668 | 0.007629026 | 0.2787 |
| HINT1 | 22653.220 | -1.210 | 0.453 | -2.668 | 0.007632907 | 0.2787 |
| GBE1 | 11649.528 | -0.788 | 0.295 | -2.666 | 0.007670909 | 0.2787 |
| LOC644100 | 262.009 | -1.392 | 0.522 | -2.665 | 0.007694022 | 0.2787 |
| RTKN2 | 2351.306 | 0.917 | 0.344 | 2.663 | 0.007737406 | 0.2787 |
| DNAJC1 | 2666.322 | -0.806 | 0.303 | -2.663 | 0.00774834 | 0.2787 |
| H2AFZ | 13545.576 | -1.110 | 0.417 | -2.663 | 0.007755588 | 0.2787 |
| CNIH | 6931.192 | -1.347 | 0.506 | -2.662 | 0.007768072 | 0.2787 |
| TAF9 | 5152.869 | -1.282 | 0.482 | -2.658 | 0.007849504 | 0.2787 |
| AHCY | 4557.486 | -1.306 | 0.491 | -2.658 | 0.007855877 | 0.2787 |
| RWDD1 | 2362.211 | -1.127 | 0.424 | -2.658 | 0.007867464 | 0.2787 |
| CAMLG | 5266.252 | -1.305 | 0.491 | -2.657 | 0.007879241 | 0.2787 |
| GMDS | 1500.099 | -0.832 | 0.313 | -2.656 | 0.00791422 | 0.2787 |
| MECOM | 145.772 | -1.722 | 0.649 | -2.654 | 0.007960816 | 0.2787 |
| DNAJA1 | 6281.056 | -0.791 | 0.298 | -2.654 | 0.007966067 | 0.2787 |
| BANF1 | 1999.606 | -1.031 | 0.389 | -2.653 | 0.00798261 | 0.2787 |
| TSPAN33 | 508.792 | -1.361 | 0.513 | -2.652 | 0.007999563 | 0.2787 |
| ZFR2 | 52.815 | 1.456 | 0.549 | 2.652 | 0.008012511 | 0.2787 |
| C17orf76-AS1 | 19989.802 | -1.186 | 0.447 | -2.651 | 0.008033546 | 0.2787 |
| TMEM170B | 269.353 | -0.760 | 0.287 | -2.649 | 0.008064608 | 0.2787 |
| ARID5A | 7760.713 | -1.046 | 0.395 | -2.649 | 0.008068827 | 0.2787 |
| KIAA1524 | 2040.256 | -0.842 | 0.318 | -2.649 | 0.0080697 | 0.2787 |
| NDUFS5 | 3000.348 | -0.984 | 0.372 | -2.648 | 0.008095381 | 0.2787 |
| SRP19 | 3114.747 | -1.188 | 0.449 | -2.648 | 0.008100319 | 0.2787 |
| FAM177A1 | 2055.820 | -1.219 | 0.460 | -2.648 | 0.008101577 | 0.2787 |
| IGFALS | 13.481 | 1.893 | 0.715 | 2.646 | 0.008136745 | NA |
| HSP90AB1 | 45954.803 | -1.049 | 0.396 | -2.646 | 0.008152494 | 0.2787 |
| ATP5L | 7325.284 | -1.146 | 0.433 | -2.645 | 0.008163804 | 0.2787 |
| RPL37A | 31852.669 | -1.084 | 0.410 | -2.645 | 0.008178358 | 0.2787 |
| RNF138 | 6037.718 | -1.278 | 0.483 | -2.644 | 0.008188432 | 0.2787 |
| BRK1 | 9677.429 | -0.796 | 0.301 | -2.644 | 0.008189574 | 0.2787 |
| RPL9 | 33585.892 | -1.133 | 0.429 | -2.642 | 0.008231822 | 0.2787 |
| PTP4A1 | 12212.817 | -1.381 | 0.523 | -2.642 | 0.008235326 | 0.2787 |
| PVRL3 | 825.316 | -0.846 | 0.320 | -2.642 | 0.008237079 | 0.2787 |
| HAGH | 1657.023 | -0.799 | 0.303 | -2.641 | 0.008265517 | 0.2787 |
| CCDC72 | 5283.547 | -1.136 | 0.430 | -2.638 | 0.008333017 | 0.2787 |
| MRPL33 | 2093.541 | -0.916 | 0.347 | -2.638 | 0.008343386 | 0.2787 |
| HMSD | 901.926 | -2.144 | 0.813 | -2.636 | 0.008377721 | 0.2787 |
| HIST1H1E | 21.679 | -2.750 | 1.043 | -2.636 | 0.008378739 | 0.2787 |
| MTHFD1 | 5835.946 | -0.655 | 0.249 | -2.634 | 0.008426266 | 0.2787 |
| CBX3 | 16165.273 | -1.160 | 0.440 | -2.634 | 0.008438805 | 0.2787 |
| CAGE1 | 127.307 | -1.725 | 0.655 | -2.633 | 0.008455121 | 0.2787 |
| VMAC | 406.094 | 1.063 | 0.404 | 2.633 | 0.008469028 | 0.2787 |
| IL1R2 | 44.269 | -2.387 | 0.907 | -2.633 | 0.008473551 | 0.2787 |
| BNIP1 | 782.239 | -1.066 | 0.405 | -2.632 | 0.008482606 | 0.2787 |
| RPL35 | 15850.059 | -1.151 | 0.437 | -2.632 | 0.008487063 | 0.2787 |
| HIST1H4E | 8.837 | -4.141 | 1.574 | -2.632 | 0.008499316 | NA |
| UBL5 | 2937.277 | -0.892 | 0.339 | -2.631 | 0.008518124 | 0.2787 |
| MRPL47 | 1223.110 | -0.832 | 0.316 | -2.630 | 0.008549541 | 0.2787 |
| ATP10A | 4356.260 | 0.917 | 0.349 | 2.629 | 0.008575621 | 0.2787 |
| CTNNAL1 | 1570.819 | -1.369 | 0.521 | -2.628 | 0.008578297 | 0.2787 |
| RPS21 | 20617.507 | -1.162 | 0.442 | -2.628 | 0.008583886 | 0.2787 |
| NR1D1 | 642.183 | -1.057 | 0.402 | -2.628 | 0.008589101 | 0.2787 |
| OSTF1 | 12445.093 | -1.205 | 0.459 | -2.627 | 0.008603013 | 0.2787 |
| PSORS1C3 | 18.418 | 3.046 | 1.159 | 2.627 | 0.008608756 | 0.2787 |
| SNRPD2 | 7279.168 | -1.009 | 0.384 | -2.626 | 0.008640003 | 0.2787 |
| C3orf38 | 1869.025 | -0.885 | 0.337 | -2.624 | 0.008681691 | 0.2787 |
| QPCT | 24.630 | -3.443 | 1.313 | -2.623 | 0.008716626 | 0.2787 |
| RNASEH2A | 949.261 | -1.132 | 0.432 | -2.622 | 0.008729462 | 0.2787 |
| POF1B | 37.619 | -2.353 | 0.898 | -2.621 | 0.008758382 | 0.2787 |
| PAK1 | 1837.731 | -0.818 | 0.312 | -2.620 | 0.008788777 | 0.2787 |
| RAB3IP | 1041.082 | -0.813 | 0.310 | -2.620 | 0.008790111 | 0.2787 |
| RNF14 | 1919.105 | -0.744 | 0.284 | -2.619 | 0.008823974 | 0.2787 |
| SET | 22620.641 | -1.123 | 0.429 | -2.617 | 0.008862448 | 0.2787 |
| NUTF2 | 2178.027 | -0.929 | 0.355 | -2.616 | 0.008890831 | 0.2787 |
| MRPL14 | 1013.295 | -0.922 | 0.352 | -2.616 | 0.008899661 | 0.2787 |
| LOC338817 | 175.189 | 1.098 | 0.420 | 2.615 | 0.008913723 | 0.2787 |
| LACC1 | 413.776 | -1.173 | 0.448 | -2.615 | 0.008916658 | 0.2787 |
| MEA1 | 1987.126 | -1.025 | 0.392 | -2.614 | 0.008952099 | 0.2787 |
| RYBP | 13147.152 | -1.062 | 0.406 | -2.614 | 0.008953984 | 0.2787 |
| HSF5 | 78.874 | -1.490 | 0.570 | -2.613 | 0.008965795 | 0.2787 |
| GHITM | 12525.940 | -1.098 | 0.420 | -2.612 | 0.008990274 | 0.2787 |
| DENND5A | 2491.962 | -0.993 | 0.380 | -2.612 | 0.00899877 | 0.2787 |
| C12orf33 | 80.705 | 1.443 | 0.553 | 2.612 | 0.009005003 | 0.2787 |
| NUDT15 | 911.989 | -1.141 | 0.437 | -2.612 | 0.009005455 | 0.2787 |
| ASB2 | 20826.008 | 0.701 | 0.268 | 2.612 | 0.009007045 | 0.2787 |
| PDCD5 | 3884.209 | -1.016 | 0.389 | -2.611 | 0.009033507 | 0.2787 |
| BEX5 | 143.463 | -2.028 | 0.777 | -2.610 | 0.009046801 | 0.2787 |
| EIF4A3 | 3684.442 | -0.831 | 0.318 | -2.610 | 0.009058741 | 0.2787 |
| CRIP1 | 6689.449 | -0.885 | 0.339 | -2.610 | 0.009067309 | 0.2787 |
| CDH12 | 62.275 | -1.565 | 0.600 | -2.609 | 0.009088783 | 0.2787 |
| HMGN1 | 16797.148 | -1.147 | 0.440 | -2.605 | 0.009197424 | 0.2787 |
| CUEDC1 | 11.572 | -1.927 | 0.740 | -2.604 | 0.009202576 | NA |
| HIST1H3J | 7.281 | -3.809 | 1.463 | -2.604 | 0.009209439 | NA |
| GNPTG | 3079.372 | 0.578 | 0.222 | 2.603 | 0.00924149 | 0.2787 |
| KCMF1 | 6915.443 | -1.104 | 0.424 | -2.603 | 0.009248698 | 0.2787 |
| SDHB | 3472.264 | -1.036 | 0.398 | -2.601 | 0.009282684 | 0.2787 |
| ING3 | 4037.545 | -1.166 | 0.449 | -2.600 | 0.009332504 | 0.2787 |
| SNHG6 | 4960.579 | -1.150 | 0.443 | -2.599 | 0.009358711 | 0.2787 |
| MGST1 | 34.495 | -2.037 | 0.784 | -2.599 | 0.009361327 | 0.2787 |
| NBPF11 | 332.191 | 0.668 | 0.257 | 2.598 | 0.009365506 | 0.2787 |
| NDFIP1 | 11192.099 | -0.967 | 0.372 | -2.598 | 0.009366697 | 0.2787 |
| DOK5 | 249.751 | -2.075 | 0.799 | -2.598 | 0.009389905 | 0.2787 |
| CABP4 | 233.372 | 1.199 | 0.462 | 2.597 | 0.009404518 | 0.2787 |
| MYL12B | 13559.751 | -1.012 | 0.390 | -2.597 | 0.009412273 | 0.2787 |
| GPX4 | 6743.014 | -0.991 | 0.381 | -2.597 | 0.009417824 | 0.2787 |
| C15orf23 | 1027.605 | -1.020 | 0.393 | -2.596 | 0.009420574 | 0.2787 |
| RALB | 5448.485 | -1.079 | 0.416 | -2.596 | 0.009440269 | 0.2787 |
| ZFAND1 | 3181.644 | -1.093 | 0.421 | -2.595 | 0.009458481 | 0.2787 |
| UPRT | 2713.183 | -0.980 | 0.378 | -2.593 | 0.009516576 | 0.2787 |
| LMNA | 10892.253 | -1.117 | 0.431 | -2.592 | 0.009551832 | 0.2787 |
| RANBP1 | 3222.672 | -1.081 | 0.417 | -2.591 | 0.009559306 | 0.2787 |
| SPCS1 | 3858.383 | -1.078 | 0.416 | -2.591 | 0.009562432 | 0.2787 |
| OLA1 | 5870.697 | -1.093 | 0.422 | -2.591 | 0.009576439 | 0.2787 |
| FAM174A | 501.692 | -0.934 | 0.361 | -2.588 | 0.009640828 | 0.2787 |
| MYL6 | 15590.736 | -0.787 | 0.304 | -2.588 | 0.009665057 | 0.2787 |
| PHF16 | 92.487 | -1.598 | 0.618 | -2.587 | 0.009691877 | 0.2787 |
| PCBP1 | 47633.612 | -1.264 | 0.489 | -2.587 | 0.009693966 | 0.2787 |
| RPS9 | 38043.456 | -1.052 | 0.407 | -2.586 | 0.009709832 | 0.2787 |
| SRP9 | 12845.773 | -1.269 | 0.491 | -2.585 | 0.009739122 | 0.2787 |
| SNRPB | 7421.692 | -1.009 | 0.391 | -2.584 | 0.009758959 | 0.2787 |
| TCEB2 | 3613.325 | -0.843 | 0.326 | -2.584 | 0.009779585 | 0.2787 |
| CDK1 | 5263.095 | -1.393 | 0.540 | -2.581 | 0.009852404 | 0.2787 |
| STK39 | 5426.431 | -1.076 | 0.417 | -2.581 | 0.009865634 | 0.2787 |
| HMMR | 1161.577 | -0.781 | 0.303 | -2.579 | 0.00989925 | 0.2787 |
| MCU | 1229.218 | -0.658 | 0.255 | -2.579 | 0.009909865 | 0.2787 |
| VAV3 | 150.736 | -0.832 | 0.323 | -2.578 | 0.009937626 | 0.2787 |
| EIF4A2 | 25467.407 | -1.143 | 0.443 | -2.578 | 0.009942827 | 0.2787 |
| LINC00116 | 283.950 | -0.911 | 0.353 | -2.577 | 0.009953482 | 0.2787 |
| TPGS2 | 3594.833 | -0.859 | 0.333 | -2.577 | 0.009959651 | 0.2787 |
| MIR548N | 250.988 | 0.688 | 0.267 | 2.577 | 0.009967781 | 0.2787 |
| NR4A3 | 2264.040 | -1.082 | 0.420 | -2.577 | 0.009970235 | 0.2787 |
| POLR3G | 557.419 | -0.882 | 0.342 | -2.577 | 0.009975945 | 0.2787 |
| SRI | 5433.643 | -0.924 | 0.359 | -2.577 | 0.009976601 | 0.2787 |
