## Supplementary material for "Marker genes for predicting cytokine release syndrome in vitro before CAR T cell infusion": Table S3

| gene | baseMean | log2FoldChange | lfcSE | stat | pvalue | padj |
| --- | --- | --- | --- | --- | --- | --- |
| BCL11A | 17.170 | -7.888 | 2.997 | -2.632 | 0.00849 | NA |
| CYP39A1 | 9.568 | -7.037 | 2.715 | -2.591 | 0.00956 | NA |
| UNC80 | 9.050 | -6.542 | 2.305 | -2.838 | 0.00454 | NA |
| MYH7 | 10.609 | -6.351 | 1.865 | -3.405 | 0.00066 | NA |
| FAM13C | 7.695 | -6.316 | 2.180 | -2.898 | 0.00376 | NA |
| SAMD5 | 5.564 | -6.270 | 2.262 | -2.772 | 0.00558 | NA |
| AZGP1P1 | 5.142 | -6.164 | 1.928 | -3.196 | 0.00139 | NA |
| GULP1 | 6.724 | -6.104 | 2.312 | -2.640 | 0.00829 | NA |
| CLEC2A | 4.942 | -6.103 | 2.253 | -2.709 | 0.00674 | NA |
| WBSCR17 | 27.967 | -6.037 | 1.539 | -3.923 | 0.00009 | NA |
| DMBT1 | 5.942 | -5.944 | 2.061 | -2.885 | 0.00392 | NA |
| LEPREL1 | 7.898 | -5.929 | 2.220 | -2.670 | 0.00758 | NA |
| COL22A1 | 7.808 | -5.905 | 2.014 | -2.931 | 0.00337 | NA |
| GPR84 | 5.641 | -5.861 | 2.044 | -2.868 | 0.00414 | NA |
| KCND2 | 9.792 | -5.827 | 2.072 | -2.812 | 0.00492 | NA |
| MEG3 | 6.104 | -5.556 | 1.866 | -2.977 | 0.00291 | NA |
| ZNF385B | 11.823 | -5.458 | 1.564 | -3.489 | 0.00048 | NA |
| NLRP11 | 6.405 | -5.242 | 1.955 | -2.681 | 0.00735 | NA |
| CDH8 | 9.250 | -5.234 | 2.012 | -2.602 | 0.00928 | NA |
| OPRM1 | 8.200 | -5.182 | 1.769 | -2.930 | 0.00339 | NA |
| IL17A | 15.210 | -5.128 | 1.988 | -2.579 | 0.00992 | NA |
| CAMK2A | 7.698 | -5.062 | 1.943 | -2.605 | 0.00918 | NA |
| RAB6C | 13.173 | -5.046 | 1.694 | -2.978 | 0.00290 | NA |
| SLC16A12 | 10.325 | -4.973 | 1.877 | -2.650 | 0.00805 | NA |
| TSPAN12 | 8.772 | -4.931 | 1.710 | -2.883 | 0.00394 | NA |
| VWA3B | 7.814 | -4.885 | 1.859 | -2.628 | 0.00860 | NA |
| PPP2R2C | 9.842 | -4.762 | 1.812 | -2.628 | 0.00859 | NA |
| PKHD1 | 21.293 | -4.685 | 1.114 | -4.205 | 0.00003 | NA |
| ABCC9 | 6.828 | -4.644 | 1.751 | -2.653 | 0.00799 | NA |
| HNF1B | 425.686 | -4.465 | 1.503 | -2.972 | 0.00296 | 0.0467 |
| KLF17 | 17.107 | -4.335 | 1.672 | -2.592 | 0.00954 | NA |
| CDO1 | 10.052 | -4.245 | 1.464 | -2.899 | 0.00374 | NA |
| MYL4 | 8.727 | -4.244 | 1.633 | -2.599 | 0.00935 | NA |
| GLT1D1 | 13.622 | -4.014 | 1.387 | -2.894 | 0.00381 | NA |
| SYT1 | 22.943 | -4.014 | 1.497 | -2.681 | 0.00734 | NA |
| HIST1H1E | 33.151 | -4.012 | 0.987 | -4.065 | 0.00005 | 0.0100 |
| ARHGAP42 | 24.124 | -3.994 | 1.157 | -3.453 | 0.00055 | NA |
| PRDM16 | 19.488 | -3.863 | 1.437 | -2.688 | 0.00719 | NA |
| KPNA7 | 7.857 | -3.834 | 1.384 | -2.770 | 0.00560 | NA |
| TBC1D9 | 15.374 | -3.820 | 1.329 | -2.874 | 0.00406 | NA |
| PPP1R3C | 22.782 | -3.742 | 1.222 | -3.063 | 0.00219 | NA |
| PDE1A | 21.600 | -3.736 | 1.388 | -2.691 | 0.00712 | NA |
| SHISA6 | 17.206 | -3.655 | 1.397 | -2.616 | 0.00891 | NA |
| RBFOX1 | 92.677 | -3.653 | 0.939 | -3.889 | 0.00010 | 0.0131 |
| LRRC4C | 77.522 | -3.637 | 1.259 | -2.888 | 0.00387 | 0.0535 |
| LUZP2 | 23.352 | -3.576 | 1.180 | -3.030 | 0.00245 | NA |
| KLHL1 | 20.663 | -3.524 | 1.249 | -2.821 | 0.00478 | NA |
| NCKAP5 | 30.674 | -3.482 | 1.094 | -3.184 | 0.00145 | 0.0332 |
| KIF6 | 17.559 | -3.480 | 1.272 | -2.735 | 0.00623 | NA |
| GRM7 | 31.990 | -3.418 | 1.297 | -2.636 | 0.00839 | 0.0808 |
| ME1 | 53.360 | -3.387 | 0.866 | -3.912 | 0.00009 | 0.0128 |
| ASCL1 | 8.866 | -3.384 | 1.286 | -2.631 | 0.00850 | NA |
| KHDRBS2 | 24.036 | -3.383 | 1.271 | -2.662 | 0.00777 | NA |
| CTNNA2 | 49.770 | -3.356 | 1.038 | -3.235 | 0.00122 | 0.0305 |
| DOCK4 | 31.723 | -3.330 | 1.068 | -3.118 | 0.00182 | 0.0372 |
| ATP5EP2 | 17.342 | -3.277 | 1.163 | -2.818 | 0.00483 | NA |
| HSPA1B | 2084.603 | -3.263 | 0.550 | -5.931 | 0.00000 | 0.0000 |
| GXYLT2 | 27.880 | -3.254 | 1.176 | -2.768 | 0.00564 | NA |
| MT1G | 63.442 | -3.225 | 0.953 | -3.383 | 0.00072 | 0.0246 |
| MACROD2 | 51.436 | -3.202 | 1.179 | -2.716 | 0.00661 | 0.0707 |
| CAP2 | 43.804 | -3.173 | 1.111 | -2.857 | 0.00428 | 0.0562 |
| NRXN1 | 43.843 | -3.119 | 1.127 | -2.767 | 0.00566 | 0.0652 |
| SNAI1 | 50.443 | -2.981 | 1.003 | -2.973 | 0.00295 | 0.0467 |
| CA2 | 81.890 | -2.947 | 0.693 | -4.254 | 0.00002 | 0.0077 |
| CAGE1 | 204.428 | -2.883 | 0.624 | -4.620 | 0.00000 | 0.0044 |
| SLC25A21 | 24.670 | -2.871 | 0.862 | -3.330 | 0.00087 | NA |
| HSPA7 | 115.547 | -2.847 | 0.969 | -2.939 | 0.00329 | 0.0493 |
| RASD1 | 41.130 | -2.828 | 0.918 | -3.081 | 0.00207 | 0.0396 |
| GPC3 | 105.462 | -2.819 | 0.907 | -3.108 | 0.00189 | 0.0378 |
| LRP1B | 82.044 | -2.813 | 0.951 | -2.957 | 0.00311 | 0.0478 |
| ROBO2 | 67.439 | -2.806 | 0.900 | -3.117 | 0.00183 | 0.0372 |
| CCDC34 | 876.739 | -2.731 | 0.619 | -4.411 | 0.00001 | 0.0059 |
| DLG2 | 75784.427 | -2.680 | 1.008 | -2.660 | 0.00782 | 0.0774 |
| FAM190A | 2885.501 | -2.669 | 0.798 | -3.346 | 0.00082 | 0.0260 |
| PTPRD | 130.213 | -2.649 | 0.863 | -3.071 | 0.00213 | 0.0399 |
| KCNH1 | 39.586 | -2.628 | 0.994 | -2.644 | 0.00820 | 0.0795 |
| SLC35F1 | 718.348 | -2.626 | 0.621 | -4.227 | 0.00002 | 0.0082 |
| OSBPL6 | 62.526 | -2.609 | 0.762 | -3.424 | 0.00062 | 0.0236 |
| H3F3C | 49.577 | -2.607 | 0.921 | -2.830 | 0.00466 | 0.0587 |
| CSMD1 | 87.023 | -2.596 | 0.813 | -3.192 | 0.00141 | 0.0330 |
| SLC16A10 | 89.554 | -2.581 | 0.639 | -4.036 | 0.00005 | 0.0106 |
| EPS8 | 77.745 | -2.554 | 0.747 | -3.419 | 0.00063 | 0.0236 |
| HMGN5 | 285.627 | -2.544 | 0.614 | -4.145 | 0.00003 | 0.0086 |
| BEX5 | 119.230 | -2.515 | 0.861 | -2.922 | 0.00348 | 0.0510 |
| KLHDC8A | 167.535 | -2.485 | 0.686 | -3.622 | 0.00029 | 0.0181 |
| TJP1 | 131.520 | -2.455 | 0.679 | -3.617 | 0.00030 | 0.0181 |
| SEMA6D | 72.107 | -2.442 | 0.883 | -2.764 | 0.00571 | 0.0652 |
| KCNK7 | 29.357 | -2.373 | 0.857 | -2.767 | 0.00565 | NA |
| HMSD | 520.722 | -2.371 | 0.709 | -3.343 | 0.00083 | 0.0260 |
| FDX1 | 3182.465 | -2.365 | 0.565 | -4.183 | 0.00003 | 0.0083 |
| CDH13 | 72.271 | -2.361 | 0.630 | -3.749 | 0.00018 | 0.0152 |
| IER3 | 1234.444 | -2.343 | 0.550 | -4.260 | 0.00002 | 0.0077 |
| C16orf87 | 2440.578 | -2.338 | 0.596 | -3.924 | 0.00009 | 0.0127 |
| ALYREF | 3140.049 | -2.333 | 0.526 | -4.432 | 0.00001 | 0.0057 |
| SMOC1 | 470.953 | -2.330 | 0.636 | -3.667 | 0.00025 | 0.0176 |
| UNC13B | 79.478 | -2.318 | 0.620 | -3.740 | 0.00018 | 0.0153 |
| RPS17 | 428.331 | -2.316 | 0.651 | -3.560 | 0.00037 | 0.0193 |
| PCDH11X | 117.345 | -2.299 | 0.703 | -3.270 | 0.00107 | 0.0287 |
| TUBB2A | 60.908 | -2.275 | 0.617 | -3.685 | 0.00023 | 0.0170 |
| DZIP1 | 161.415 | -2.265 | 0.750 | -3.021 | 0.00252 | 0.0431 |
| 10-Sep | 55.049 | -2.262 | 0.714 | -3.169 | 0.00153 | 0.0345 |
| FRAS1 | 60.696 | -2.260 | 0.827 | -2.734 | 0.00627 | 0.0687 |
| FAM59A | 251.816 | -2.239 | 0.660 | -3.393 | 0.00069 | 0.0246 |
| OSM | 275.244 | -2.238 | 0.601 | -3.721 | 0.00020 | 0.0161 |
| CDH12 | 86.008 | -2.225 | 0.816 | -2.728 | 0.00638 | 0.0694 |
| BFSP2 | 48.848 | -2.222 | 0.760 | -2.925 | 0.00345 | 0.0507 |
| FLJ43860 | 217.532 | -2.216 | 0.657 | -3.373 | 0.00074 | 0.0251 |
| GPR176 | 18.471 | -2.215 | 0.702 | -3.157 | 0.00160 | NA |
| A4GALT | 32.747 | -2.214 | 0.769 | -2.878 | 0.00401 | 0.0545 |
| TMEM38B | 609.494 | -2.203 | 0.550 | -4.009 | 0.00006 | 0.0115 |
| HSP90AB3P | 61.323 | -2.198 | 0.839 | -2.620 | 0.00879 | 0.0832 |
| HPGDS | 177.596 | -2.187 | 0.837 | -2.611 | 0.00903 | 0.0843 |
| LOC643339 | 97.941 | -2.182 | 0.685 | -3.183 | 0.00146 | 0.0332 |
| UBE2T | 1684.909 | -2.175 | 0.562 | -3.871 | 0.00011 | 0.0132 |
| SNRPG | 3254.746 | -2.171 | 0.462 | -4.698 | 0.00000 | 0.0043 |
| AKAP6 | 132.989 | -2.171 | 0.746 | -2.910 | 0.00361 | 0.0519 |
| GADD45A | 4525.842 | -2.159 | 0.716 | -3.013 | 0.00258 | 0.0434 |
| PDE3A | 3363.925 | -2.145 | 0.564 | -3.806 | 0.00014 | 0.0137 |
| FLJ39080 | 103.741 | -2.134 | 0.741 | -2.882 | 0.00395 | 0.0541 |
| IL23R | 382.978 | -2.134 | 0.649 | -3.290 | 0.00100 | 0.0279 |
| CCL3 | 1569.646 | -2.120 | 0.740 | -2.863 | 0.00420 | 0.0560 |
| MECOM | 192.943 | -2.111 | 0.724 | -2.916 | 0.00355 | 0.0515 |
| SAP30 | 4984.978 | -2.109 | 0.525 | -4.020 | 0.00006 | 0.0111 |
| PLOD2 | 52.476 | -2.100 | 0.637 | -3.299 | 0.00097 | 0.0275 |
| FOS | 96.765 | -2.096 | 0.807 | -2.597 | 0.00940 | 0.0862 |
| ID3 | 71.542 | -2.079 | 0.679 | -3.061 | 0.00221 | 0.0406 |
| IL13 | 218.164 | -2.067 | 0.447 | -4.627 | 0.00000 | 0.0044 |
| DNAJC12 | 364.463 | -2.054 | 0.746 | -2.753 | 0.00590 | 0.0667 |
| RGS16 | 162.663 | -2.051 | 0.678 | -3.026 | 0.00248 | 0.0429 |
| FGD6 | 79.264 | -2.015 | 0.691 | -2.916 | 0.00355 | 0.0515 |
| HS3ST4 | 99.819 | -2.012 | 0.600 | -3.356 | 0.00079 | 0.0257 |
| RPL22L1 | 2103.835 | -2.004 | 0.520 | -3.852 | 0.00012 | 0.0132 |
| NPM3 | 614.600 | -2.003 | 0.618 | -3.243 | 0.00118 | 0.0301 |
| ANKRD30BL | 4137.650 | -2.002 | 0.527 | -3.799 | 0.00015 | 0.0137 |
| LOC644100 | 375.380 | -2.001 | 0.573 | -3.491 | 0.00048 | 0.0213 |
| RPSAP58 | 2025.417 | -1.995 | 0.509 | -3.917 | 0.00009 | 0.0128 |
| TANC1 | 46.116 | -1.989 | 0.758 | -2.623 | 0.00870 | 0.0827 |
| C3orf26 | 581.194 | -1.972 | 0.467 | -4.222 | 0.00002 | 0.0082 |
| ROMO1 | 954.746 | -1.969 | 0.426 | -4.627 | 0.00000 | 0.0044 |
| GALNTL4 | 113.602 | -1.964 | 0.648 | -3.031 | 0.00244 | 0.0428 |
| MLF1 | 122.239 | -1.957 | 0.591 | -3.314 | 0.00092 | 0.0268 |
| KCNMA1 | 68.154 | -1.928 | 0.705 | -2.735 | 0.00624 | 0.0686 |
| PFDN2 | 2104.373 | -1.924 | 0.492 | -3.912 | 0.00009 | 0.0128 |
| NUDT11 | 65.959 | -1.905 | 0.685 | -2.782 | 0.00541 | 0.0638 |
| KRT7 | 367.795 | -1.905 | 0.480 | -3.965 | 0.00007 | 0.0116 |
| CDKN3 | 1191.771 | -1.902 | 0.488 | -3.900 | 0.00010 | 0.0130 |
| CYB5A | 958.405 | -1.900 | 0.539 | -3.524 | 0.00043 | 0.0203 |
| SEMA5A | 56.685 | -1.894 | 0.675 | -2.808 | 0.00499 | 0.0607 |
| SNRPF | 3687.773 | -1.889 | 0.444 | -4.254 | 0.00002 | 0.0077 |
| NXT1 | 648.803 | -1.871 | 0.533 | -3.514 | 0.00044 | 0.0204 |
| UNC5D | 60.066 | -1.865 | 0.653 | -2.857 | 0.00428 | 0.0562 |
| DAPK1 | 527.483 | -1.859 | 0.566 | -3.282 | 0.00103 | 0.0284 |
| C17orf79 | 347.893 | -1.856 | 0.469 | -3.958 | 0.00008 | 0.0117 |
| SCARNA17 | 111.503 | -1.856 | 0.718 | -2.586 | 0.00971 | 0.0881 |
| TCEB1 | 4002.598 | -1.851 | 0.502 | -3.686 | 0.00023 | 0.0170 |
| RPL35 | 20346.163 | -1.850 | 0.415 | -4.463 | 0.00001 | 0.0056 |
| DIAPH3 | 1306.951 | -1.845 | 0.501 | -3.681 | 0.00023 | 0.0171 |
| PPARG | 748.509 | -1.843 | 0.651 | -2.831 | 0.00464 | 0.0587 |
| MZT1 | 3503.478 | -1.833 | 0.541 | -3.390 | 0.00070 | 0.0246 |
| HBXIP | 4661.669 | -1.833 | 0.459 | -3.993 | 0.00007 | 0.0115 |
| ANKRD37 | 1845.269 | -1.832 | 0.497 | -3.684 | 0.00023 | 0.0170 |
| NR4A1 | 566.739 | -1.827 | 0.511 | -3.576 | 0.00035 | 0.0191 |
| C15orf48 | 159.219 | -1.826 | 0.702 | -2.600 | 0.00933 | 0.0859 |
| NR4A2 | 215.368 | -1.818 | 0.561 | -3.239 | 0.00120 | 0.0303 |
| CKS1B | 2220.916 | -1.816 | 0.489 | -3.713 | 0.00020 | 0.0164 |
| ANKRD6 | 75.123 | -1.802 | 0.652 | -2.766 | 0.00568 | 0.0652 |
| GPN3 | 824.056 | -1.799 | 0.523 | -3.440 | 0.00058 | 0.0230 |
| OSBPL1A | 191.200 | -1.798 | 0.574 | -3.130 | 0.00175 | 0.0365 |
| ASS1 | 168.242 | -1.798 | 0.646 | -2.783 | 0.00539 | 0.0638 |
| POLR2L | 1107.578 | -1.796 | 0.477 | -3.769 | 0.00016 | 0.0146 |
| CDK2AP1 | 2254.578 | -1.791 | 0.506 | -3.542 | 0.00040 | 0.0199 |
| C5orf32 | 768.368 | -1.790 | 0.453 | -3.951 | 0.00008 | 0.0117 |
| CRIP1 | 11790.988 | -1.778 | 0.425 | -4.182 | 0.00003 | 0.0083 |
| IQCK | 45.976 | -1.773 | 0.565 | -3.135 | 0.00172 | 0.0362 |
| LGMN | 777.243 | -1.769 | 0.555 | -3.190 | 0.00142 | 0.0331 |
| C14orf142 | 898.972 | -1.758 | 0.458 | -3.836 | 0.00012 | 0.0132 |
| RPL9 | 43320.420 | -1.755 | 0.415 | -4.226 | 0.00002 | 0.0082 |
| CTNNAL1 | 1605.697 | -1.755 | 0.524 | -3.348 | 0.00081 | 0.0260 |
| HSPA2 | 222.467 | -1.751 | 0.425 | -4.126 | 0.00004 | 0.0086 |
| STK3 | 217.246 | -1.749 | 0.533 | -3.283 | 0.00103 | 0.0284 |
| C10orf35 | 156.854 | -1.746 | 0.577 | -3.028 | 0.00246 | 0.0429 |
| EIF5B | 11203.633 | -1.744 | 0.485 | -3.594 | 0.00033 | 0.0187 |
| SNHG8 | 1184.236 | -1.741 | 0.449 | -3.878 | 0.00011 | 0.0132 |
| GGCT | 2361.742 | -1.734 | 0.515 | -3.371 | 0.00075 | 0.0252 |
| ACOT7 | 1476.168 | -1.725 | 0.410 | -4.202 | 0.00003 | 0.0083 |
| PTPRG | 308.869 | -1.724 | 0.544 | -3.169 | 0.00153 | 0.0345 |
| TBCA | 6388.497 | -1.723 | 0.474 | -3.635 | 0.00028 | 0.0177 |
| ARHGEF37 | 80.850 | -1.722 | 0.451 | -3.821 | 0.00013 | 0.0132 |
| RRM2 | 12660.322 | -1.721 | 0.473 | -3.642 | 0.00027 | 0.0177 |
| TXNDC17 | 1126.202 | -1.720 | 0.412 | -4.173 | 0.00003 | 0.0084 |
| SNX10 | 3962.496 | -1.718 | 0.503 | -3.416 | 0.00064 | 0.0237 |
| RGS2 | 2052.515 | -1.716 | 0.425 | -4.040 | 0.00005 | 0.0106 |
| IL4I1 | 260.281 | -1.711 | 0.485 | -3.527 | 0.00042 | 0.0203 |
| NUDCD2 | 3042.060 | -1.709 | 0.488 | -3.499 | 0.00047 | 0.0210 |
| AHCY | 4813.148 | -1.700 | 0.455 | -3.734 | 0.00019 | 0.0156 |
| RAN | 13944.540 | -1.699 | 0.422 | -4.029 | 0.00006 | 0.0108 |
| SNHG15 | 4254.505 | -1.699 | 0.488 | -3.481 | 0.00050 | 0.0216 |
| COX6A1 | 3894.586 | -1.696 | 0.390 | -4.353 | 0.00001 | 0.0064 |
| TMEM237 | 965.694 | -1.694 | 0.507 | -3.345 | 0.00082 | 0.0260 |
| TSPAN33 | 570.961 | -1.694 | 0.614 | -2.760 | 0.00578 | 0.0658 |
| SNHG5 | 9320.383 | -1.689 | 0.483 | -3.495 | 0.00047 | 0.0211 |
| BRP44 | 3638.701 | -1.689 | 0.458 | -3.684 | 0.00023 | 0.0170 |
| WBP5 | 57.593 | -1.686 | 0.610 | -2.765 | 0.00570 | 0.0652 |
| BATF3 | 1066.123 | -1.676 | 0.487 | -3.443 | 0.00058 | 0.0230 |
| RPL39L | 511.173 | -1.676 | 0.437 | -3.834 | 0.00013 | 0.0132 |
| HLF | 176.728 | -1.670 | 0.604 | -2.764 | 0.00572 | 0.0652 |
| NUDT1 | 1242.440 | -1.668 | 0.385 | -4.329 | 0.00001 | 0.0068 |
| LSM6 | 1984.656 | -1.664 | 0.429 | -3.878 | 0.00011 | 0.0132 |
| NGFRAP1 | 920.261 | -1.662 | 0.453 | -3.672 | 0.00024 | 0.0174 |
| ST20 | 44.300 | -1.660 | 0.453 | -3.662 | 0.00025 | 0.0176 |
| C12orf45 | 403.460 | -1.660 | 0.465 | -3.567 | 0.00036 | 0.0193 |
| BCAS4 | 1227.481 | -1.659 | 0.460 | -3.606 | 0.00031 | 0.0183 |
| PTP4A1 | 10925.762 | -1.659 | 0.504 | -3.292 | 0.00099 | 0.0278 |
| GADD45GIP1 | 1409.839 | -1.654 | 0.366 | -4.517 | 0.00001 | 0.0052 |
| RPS27L | 8540.777 | -1.646 | 0.416 | -3.961 | 0.00007 | 0.0117 |
| LSMD1 | 1059.334 | -1.641 | 0.381 | -4.304 | 0.00002 | 0.0070 |
| SMNDC1 | 5058.250 | -1.641 | 0.485 | -3.384 | 0.00071 | 0.0246 |
| UQCRHL | 434.640 | -1.640 | 0.466 | -3.515 | 0.00044 | 0.0204 |
| CWF19L2 | 2037.611 | -1.636 | 0.442 | -3.700 | 0.00022 | 0.0167 |
| ING2 | 1306.312 | -1.635 | 0.491 | -3.331 | 0.00087 | 0.0264 |
| SCD5 | 211.538 | -1.635 | 0.593 | -2.756 | 0.00585 | 0.0664 |
| MND1 | 704.528 | -1.635 | 0.456 | -3.583 | 0.00034 | 0.0190 |
| TAF3 | 2020.619 | -1.633 | 0.427 | -3.820 | 0.00013 | 0.0132 |
| RPS21 | 26645.327 | -1.627 | 0.425 | -3.825 | 0.00013 | 0.0132 |
| RPL36 | 20469.452 | -1.626 | 0.395 | -4.119 | 0.00004 | 0.0086 |
| SNRPD1 | 2607.757 | -1.624 | 0.461 | -3.523 | 0.00043 | 0.0203 |
| ATF3 | 1871.674 | -1.624 | 0.605 | -2.683 | 0.00730 | 0.0744 |
| DTWD2 | 1205.178 | -1.613 | 0.475 | -3.395 | 0.00069 | 0.0246 |
| RPS9 | 46931.594 | -1.612 | 0.390 | -4.131 | 0.00004 | 0.0086 |
| NDFIP2 | 6037.631 | -1.610 | 0.560 | -2.877 | 0.00402 | 0.0546 |
| NLN | 363.890 | -1.608 | 0.474 | -3.389 | 0.00070 | 0.0246 |
| JDP2 | 78.874 | -1.607 | 0.532 | -3.019 | 0.00254 | 0.0432 |
| PLEKHA7 | 1061.633 | -1.606 | 0.572 | -2.807 | 0.00500 | 0.0608 |
| UBE2C | 2480.984 | -1.604 | 0.439 | -3.653 | 0.00026 | 0.0176 |
| MBOAT2 | 216.787 | -1.603 | 0.475 | -3.378 | 0.00073 | 0.0250 |
| PBK | 836.416 | -1.601 | 0.524 | -3.056 | 0.00224 | 0.0407 |
| GTF3C6 | 2086.970 | -1.600 | 0.462 | -3.460 | 0.00054 | 0.0223 |
| DGKI | 1002.050 | -1.599 | 0.609 | -2.627 | 0.00860 | 0.0821 |
| PPA1 | 9388.608 | -1.594 | 0.453 | -3.518 | 0.00043 | 0.0204 |
| C4orf43 | 1714.150 | -1.593 | 0.537 | -2.964 | 0.00304 | 0.0474 |
| C1orf135 | 127.206 | -1.589 | 0.526 | -3.020 | 0.00252 | 0.0431 |
| COX17 | 2116.361 | -1.588 | 0.419 | -3.787 | 0.00015 | 0.0141 |
| CNIH | 7087.524 | -1.587 | 0.471 | -3.369 | 0.00076 | 0.0252 |
| MT1F | 279.987 | -1.586 | 0.473 | -3.356 | 0.00079 | 0.0257 |
| CDC20 | 2427.524 | -1.586 | 0.346 | -4.580 | 0.00000 | 0.0047 |
| GMPR | 30.777 | -1.581 | 0.489 | -3.231 | 0.00123 | 0.0305 |
| LOC100288778 | 1769.942 | -1.577 | 0.513 | -3.073 | 0.00212 | 0.0399 |
| DBI | 4615.466 | -1.577 | 0.399 | -3.954 | 0.00008 | 0.0117 |
| BRIX1 | 1464.672 | -1.573 | 0.466 | -3.377 | 0.00073 | 0.0250 |
| LINC00115 | 216.285 | -1.569 | 0.523 | -3.000 | 0.00270 | 0.0445 |
| TYMS | 6145.376 | -1.566 | 0.515 | -3.039 | 0.00237 | 0.0422 |
| SUMO2 | 16298.712 | -1.565 | 0.480 | -3.259 | 0.00112 | 0.0292 |
| ARMC9 | 184.852 | -1.562 | 0.581 | -2.689 | 0.00716 | 0.0738 |
| C17orf89 | 623.217 | -1.561 | 0.372 | -4.192 | 0.00003 | 0.0083 |
| IL4 | 80.316 | -1.554 | 0.391 | -3.975 | 0.00007 | 0.0115 |
| C17orf58 | 1071.041 | -1.554 | 0.516 | -3.010 | 0.00261 | 0.0436 |
| ECE2 | 195.715 | -1.547 | 0.349 | -4.427 | 0.00001 | 0.0057 |
| CDCA7 | 11799.210 | -1.547 | 0.542 | -2.854 | 0.00431 | 0.0565 |
| RBM7 | 2751.782 | -1.547 | 0.531 | -2.910 | 0.00361 | 0.0519 |
| C16orf59 | 182.698 | -1.546 | 0.359 | -4.300 | 0.00002 | 0.0070 |
| TCEAL4 | 549.199 | -1.544 | 0.387 | -3.990 | 0.00007 | 0.0115 |
| MYEOV2 | 1311.056 | -1.541 | 0.445 | -3.466 | 0.00053 | 0.0222 |
| CDK1 | 4351.270 | -1.538 | 0.481 | -3.198 | 0.00138 | 0.0328 |
| MTHFD2 | 5541.168 | -1.537 | 0.578 | -2.661 | 0.00780 | 0.0773 |
| SKA1 | 1218.010 | -1.536 | 0.480 | -3.202 | 0.00137 | 0.0325 |
| BRI3 | 2819.302 | -1.534 | 0.438 | -3.503 | 0.00046 | 0.0210 |
| TMSB10 | 90284.558 | -1.533 | 0.415 | -3.696 | 0.00022 | 0.0168 |
| TXN | 8913.895 | -1.531 | 0.405 | -3.778 | 0.00016 | 0.0144 |
| CHN1 | 405.033 | -1.530 | 0.559 | -2.736 | 0.00623 | 0.0686 |
| MYBL2 | 2937.672 | -1.527 | 0.428 | -3.567 | 0.00036 | 0.0193 |
| TOMM5 | 870.243 | -1.524 | 0.440 | -3.464 | 0.00053 | 0.0222 |
| NEK2 | 1960.486 | -1.523 | 0.464 | -3.285 | 0.00102 | 0.0282 |
| RNASEH2A | 963.334 | -1.522 | 0.402 | -3.785 | 0.00015 | 0.0141 |
| ATP5I | 2452.153 | -1.522 | 0.431 | -3.528 | 0.00042 | 0.0203 |
| COX6C | 5000.322 | -1.521 | 0.398 | -3.825 | 0.00013 | 0.0132 |
| C3orf14 | 239.880 | -1.515 | 0.381 | -3.974 | 0.00007 | 0.0115 |
| STMN1 | 12774.585 | -1.515 | 0.398 | -3.803 | 0.00014 | 0.0137 |
| GPX1 | 3101.618 | -1.512 | 0.346 | -4.366 | 0.00001 | 0.0064 |
| FAM174A | 470.397 | -1.510 | 0.441 | -3.424 | 0.00062 | 0.0236 |
| ODC1 | 5324.158 | -1.509 | 0.446 | -3.385 | 0.00071 | 0.0246 |
| PTTG1 | 3815.076 | -1.507 | 0.426 | -3.542 | 0.00040 | 0.0199 |
| DNAJC6 | 340.797 | -1.507 | 0.554 | -2.718 | 0.00656 | 0.0704 |
| GPX4 | 6020.706 | -1.504 | 0.377 | -3.987 | 0.00007 | 0.0115 |
| PSMA7 | 5993.238 | -1.502 | 0.361 | -4.157 | 0.00003 | 0.0085 |
| LRRCC1 | 934.671 | -1.502 | 0.521 | -2.883 | 0.00394 | 0.0540 |
| ZNF593 | 295.175 | -1.501 | 0.480 | -3.126 | 0.00177 | 0.0366 |
| PPP1R2 | 12005.912 | -1.499 | 0.478 | -3.135 | 0.00172 | 0.0362 |
| TCEA1 | 18684.085 | -1.496 | 0.514 | -2.908 | 0.00363 | 0.0520 |
| SNRPD2 | 8501.101 | -1.492 | 0.387 | -3.856 | 0.00012 | 0.0132 |
| USMG5 | 2898.205 | -1.490 | 0.427 | -3.487 | 0.00049 | 0.0214 |
| RPL37A | 40553.848 | -1.490 | 0.411 | -3.621 | 0.00029 | 0.0181 |
| DUSP10 | 2925.050 | -1.489 | 0.388 | -3.832 | 0.00013 | 0.0132 |
| YBX1 | 31009.869 | -1.488 | 0.433 | -3.438 | 0.00059 | 0.0230 |
| TEC | 503.309 | -1.486 | 0.433 | -3.434 | 0.00060 | 0.0232 |
| MYL6 | 21770.839 | -1.485 | 0.407 | -3.646 | 0.00027 | 0.0177 |
| GTF2A2 | 2705.738 | -1.484 | 0.461 | -3.219 | 0.00128 | 0.0310 |
| C15orf24 | 3133.519 | -1.482 | 0.411 | -3.608 | 0.00031 | 0.0183 |
| TRIAP1 | 1935.074 | -1.479 | 0.468 | -3.163 | 0.00156 | 0.0348 |
| CEP55 | 4724.839 | -1.476 | 0.478 | -3.086 | 0.00203 | 0.0394 |
| TMSB4X | 248581.251 | -1.475 | 0.412 | -3.580 | 0.00034 | 0.0191 |
| ENOPH1 | 3407.728 | -1.474 | 0.462 | -3.193 | 0.00141 | 0.0330 |
| GTF2F2 | 1302.129 | -1.471 | 0.393 | -3.743 | 0.00018 | 0.0153 |
| CCDC72 | 5716.102 | -1.470 | 0.402 | -3.654 | 0.00026 | 0.0176 |
| GSTP1 | 5731.734 | -1.470 | 0.311 | -4.728 | 0.00000 | 0.0043 |
| SERPINE1 | 704.350 | -1.469 | 0.471 | -3.120 | 0.00181 | 0.0370 |
| SF3B14 | 3451.222 | -1.467 | 0.431 | -3.404 | 0.00066 | 0.0243 |
| CCNB2 | 2522.534 | -1.466 | 0.379 | -3.865 | 0.00011 | 0.0132 |
| LTA | 867.272 | -1.465 | 0.356 | -4.120 | 0.00004 | 0.0086 |
| SV2C | 183.794 | -1.465 | 0.565 | -2.594 | 0.00950 | 0.0868 |
| SCML2 | 59.390 | -1.464 | 0.503 | -2.909 | 0.00363 | 0.0520 |
| HINT1 | 24064.725 | -1.463 | 0.402 | -3.640 | 0.00027 | 0.0177 |
| CRIP2 | 123.011 | -1.463 | 0.529 | -2.766 | 0.00568 | 0.0652 |
| C6orf225 | 89.963 | -1.463 | 0.469 | -3.121 | 0.00180 | 0.0370 |
| C12orf62 | 1381.565 | -1.461 | 0.371 | -3.942 | 0.00008 | 0.0119 |
| AIG1 | 301.706 | -1.460 | 0.330 | -4.430 | 0.00001 | 0.0057 |
| SNAPC1 | 1956.366 | -1.460 | 0.561 | -2.603 | 0.00924 | 0.0855 |
| HOXB7 | 58.107 | -1.458 | 0.565 | -2.579 | 0.00991 | 0.0888 |
| C6orf115 | 8654.394 | -1.457 | 0.483 | -3.015 | 0.00257 | 0.0434 |
| MED19 | 587.704 | -1.454 | 0.453 | -3.208 | 0.00134 | 0.0320 |
| ISG15 | 2381.918 | -1.451 | 0.494 | -2.938 | 0.00330 | 0.0494 |
| WDYHV1 | 737.115 | -1.449 | 0.433 | -3.344 | 0.00083 | 0.0260 |
| MT1X | 532.550 | -1.449 | 0.507 | -2.860 | 0.00424 | 0.0561 |
| ATP5G3 | 8050.206 | -1.448 | 0.389 | -3.718 | 0.00020 | 0.0162 |
| LOC100127983 | 191.819 | -1.446 | 0.520 | -2.783 | 0.00539 | 0.0638 |
| CHEK1 | 2698.213 | -1.446 | 0.446 | -3.241 | 0.00119 | 0.0301 |
| RRAGD | 3319.288 | -1.445 | 0.343 | -4.216 | 0.00002 | 0.0082 |
| PIM3 | 4885.494 | -1.445 | 0.430 | -3.358 | 0.00079 | 0.0257 |
| GINS2 | 987.682 | -1.443 | 0.426 | -3.387 | 0.00071 | 0.0246 |
| PPP2R2B | 502.266 | -1.442 | 0.548 | -2.633 | 0.00847 | 0.0814 |
| CD70 | 4159.692 | -1.442 | 0.332 | -4.348 | 0.00001 | 0.0064 |
| SLC48A1 | 857.141 | -1.439 | 0.404 | -3.560 | 0.00037 | 0.0193 |
| PHF5A | 1277.007 | -1.436 | 0.459 | -3.132 | 0.00173 | 0.0364 |
| RPS3A | 76720.461 | -1.436 | 0.372 | -3.862 | 0.00011 | 0.0132 |
| LRRC42 | 3402.180 | -1.436 | 0.474 | -3.028 | 0.00247 | 0.0429 |
| POLR2I | 722.313 | -1.434 | 0.415 | -3.457 | 0.00055 | 0.0225 |
| COX7A2L | 8835.298 | -1.434 | 0.432 | -3.317 | 0.00091 | 0.0267 |
| RPL18A | 35940.432 | -1.433 | 0.382 | -3.751 | 0.00018 | 0.0152 |
| RPL38 | 19636.224 | -1.432 | 0.405 | -3.539 | 0.00040 | 0.0199 |
| SNHG6 | 6206.523 | -1.432 | 0.391 | -3.664 | 0.00025 | 0.0176 |
| NAMPT | 16165.383 | -1.430 | 0.498 | -2.873 | 0.00406 | 0.0549 |
| C5orf55 | 164.604 | -1.429 | 0.412 | -3.470 | 0.00052 | 0.0221 |
| TIMM10 | 643.529 | -1.429 | 0.430 | -3.323 | 0.00089 | 0.0266 |
| TCEB2 | 4267.425 | -1.429 | 0.402 | -3.555 | 0.00038 | 0.0194 |
| CETN2 | 394.950 | -1.428 | 0.429 | -3.326 | 0.00088 | 0.0265 |
| UBE2S | 2073.142 | -1.427 | 0.317 | -4.500 | 0.00001 | 0.0053 |
| C14orf109 | 287.735 | -1.426 | 0.526 | -2.712 | 0.00668 | 0.0709 |
| PPP1R15A | 9110.211 | -1.425 | 0.465 | -3.067 | 0.00216 | 0.0401 |
| RPS19 | 72947.105 | -1.422 | 0.369 | -3.855 | 0.00012 | 0.0132 |
| GLUL | 12373.326 | -1.421 | 0.360 | -3.949 | 0.00008 | 0.0117 |
| MLF1IP | 3239.492 | -1.421 | 0.471 | -3.018 | 0.00254 | 0.0432 |
| CHCHD3 | 4119.240 | -1.418 | 0.425 | -3.336 | 0.00085 | 0.0261 |
| AZIN1 | 11898.389 | -1.417 | 0.502 | -2.822 | 0.00477 | 0.0593 |
| SLC25A5 | 15456.568 | -1.415 | 0.393 | -3.603 | 0.00031 | 0.0184 |
| GNA15 | 11284.306 | -1.415 | 0.468 | -3.023 | 0.00251 | 0.0431 |
| PSMB6 | 2427.860 | -1.414 | 0.390 | -3.630 | 0.00028 | 0.0179 |
| RPL26L1 | 842.953 | -1.414 | 0.460 | -3.071 | 0.00213 | 0.0399 |
| SAP18 | 5672.265 | -1.412 | 0.417 | -3.385 | 0.00071 | 0.0246 |
| C12orf57 | 1459.707 | -1.412 | 0.405 | -3.489 | 0.00048 | 0.0214 |
| RPS28 | 23277.036 | -1.411 | 0.422 | -3.343 | 0.00083 | 0.0260 |
| CCNA2 | 3589.412 | -1.411 | 0.455 | -3.105 | 0.00190 | 0.0378 |
| LINC00116 | 308.739 | -1.411 | 0.433 | -3.256 | 0.00113 | 0.0292 |
| LMNA | 10348.715 | -1.411 | 0.449 | -3.145 | 0.00166 | 0.0357 |
| NDUFA3 | 984.047 | -1.410 | 0.353 | -3.999 | 0.00006 | 0.0115 |
| COX7A2 | 4446.169 | -1.409 | 0.367 | -3.840 | 0.00012 | 0.0132 |
| BIRC5 | 1917.364 | -1.409 | 0.354 | -3.979 | 0.00007 | 0.0115 |
| LOC728554 | 980.216 | -1.409 | 0.306 | -4.612 | 0.00000 | 0.0044 |
| RPS7 | 32414.240 | -1.409 | 0.367 | -3.841 | 0.00012 | 0.0132 |
| NPC2 | 3577.261 | -1.404 | 0.407 | -3.452 | 0.00056 | 0.0226 |
| TCEAL3 | 160.023 | -1.403 | 0.496 | -2.830 | 0.00466 | 0.0587 |
| BLOC1S2 | 5288.383 | -1.402 | 0.419 | -3.347 | 0.00082 | 0.0260 |
| MRPL36 | 709.353 | -1.400 | 0.422 | -3.315 | 0.00091 | 0.0268 |
| CHCHD1 | 1530.215 | -1.399 | 0.368 | -3.802 | 0.00014 | 0.0137 |
| NDUFB9 | 4847.743 | -1.398 | 0.401 | -3.485 | 0.00049 | 0.0215 |
| CYTIP | 24880.668 | -1.396 | 0.467 | -2.988 | 0.00280 | 0.0457 |
| MRPL54 | 1321.840 | -1.394 | 0.461 | -3.021 | 0.00252 | 0.0431 |
| FLJ20021 | 250.845 | -1.394 | 0.464 | -3.003 | 0.00268 | 0.0442 |
| CDT1 | 4505.945 | -1.393 | 0.466 | -2.986 | 0.00283 | 0.0458 |
| CENPP | 592.650 | -1.390 | 0.410 | -3.393 | 0.00069 | 0.0246 |
| RWDD1 | 2651.750 | -1.389 | 0.397 | -3.498 | 0.00047 | 0.0210 |
| DEPDC1 | 1427.832 | -1.388 | 0.478 | -2.904 | 0.00369 | 0.0521 |
| NDUFA1 | 4181.352 | -1.387 | 0.398 | -3.480 | 0.00050 | 0.0216 |
| EIF4EBP1 | 1455.345 | -1.385 | 0.391 | -3.540 | 0.00040 | 0.0199 |
| SPC25 | 476.845 | -1.385 | 0.374 | -3.701 | 0.00021 | 0.0167 |
| PRADC1 | 248.499 | -1.384 | 0.477 | -2.905 | 0.00368 | 0.0521 |
| C17orf76-AS1 | 23727.739 | -1.382 | 0.404 | -3.417 | 0.00063 | 0.0237 |
| C14orf2 | 2972.447 | -1.382 | 0.390 | -3.542 | 0.00040 | 0.0199 |
| MEA1 | 1786.645 | -1.381 | 0.353 | -3.912 | 0.00009 | 0.0128 |
| NUDT8 | 218.711 | -1.379 | 0.463 | -2.975 | 0.00293 | 0.0465 |
| NDUFS5 | 3446.665 | -1.378 | 0.410 | -3.364 | 0.00077 | 0.0254 |
| CDCA5 | 3583.619 | -1.376 | 0.422 | -3.263 | 0.00110 | 0.0290 |
| DPM3 | 608.430 | -1.376 | 0.400 | -3.445 | 0.00057 | 0.0230 |
| MRPL27 | 1106.793 | -1.371 | 0.328 | -4.181 | 0.00003 | 0.0083 |
| BEX2 | 268.387 | -1.370 | 0.420 | -3.266 | 0.00109 | 0.0289 |
| VAMP8 | 3755.394 | -1.368 | 0.379 | -3.607 | 0.00031 | 0.0183 |
| SH2D1A | 12714.561 | -1.367 | 0.434 | -3.146 | 0.00165 | 0.0357 |
| PSMG3 | 709.009 | -1.365 | 0.395 | -3.456 | 0.00055 | 0.0225 |
| COX5A | 5729.621 | -1.362 | 0.327 | -4.164 | 0.00003 | 0.0085 |
| ATP5L | 9034.687 | -1.360 | 0.401 | -3.388 | 0.00070 | 0.0246 |
| POLR3K | 590.025 | -1.359 | 0.409 | -3.327 | 0.00088 | 0.0265 |
| RPL12 | 46848.107 | -1.359 | 0.348 | -3.902 | 0.00010 | 0.0130 |
| MBIP | 2463.693 | -1.358 | 0.432 | -3.142 | 0.00168 | 0.0358 |
| LOC100505483 | 142.837 | -1.358 | 0.456 | -2.978 | 0.00291 | 0.0462 |
| AP2S1 | 2531.984 | -1.357 | 0.378 | -3.587 | 0.00033 | 0.0188 |
| IFNGR2 | 169.637 | -1.356 | 0.508 | -2.668 | 0.00764 | 0.0761 |
| CHCHD10 | 692.325 | -1.355 | 0.340 | -3.986 | 0.00007 | 0.0115 |
| PFDN4 | 2198.186 | -1.355 | 0.434 | -3.121 | 0.00180 | 0.0370 |
| GTF2IRD1 | 371.326 | -1.354 | 0.338 | -4.003 | 0.00006 | 0.0115 |
| RANBP1 | 2486.461 | -1.353 | 0.371 | -3.649 | 0.00026 | 0.0177 |
| PCNA | 8625.621 | -1.352 | 0.373 | -3.627 | 0.00029 | 0.0180 |
| CCDC112 | 437.638 | -1.351 | 0.501 | -2.699 | 0.00695 | 0.0722 |
| UQCRQ | 2130.576 | -1.350 | 0.368 | -3.674 | 0.00024 | 0.0174 |
| MRPL34 | 1805.232 | -1.350 | 0.416 | -3.246 | 0.00117 | 0.0299 |
| RPL28 | 53397.428 | -1.350 | 0.369 | -3.663 | 0.00025 | 0.0176 |
| GAS5 | 13982.606 | -1.347 | 0.363 | -3.709 | 0.00021 | 0.0165 |
| RPS6 | 107065.655 | -1.345 | 0.379 | -3.550 | 0.00039 | 0.0196 |
| NDUFA2 | 1766.401 | -1.344 | 0.394 | -3.415 | 0.00064 | 0.0237 |
| LSM3 | 1910.584 | -1.344 | 0.427 | -3.150 | 0.00163 | 0.0356 |
| POC1A | 814.050 | -1.341 | 0.349 | -3.846 | 0.00012 | 0.0132 |
| DUSP1 | 414.560 | -1.339 | 0.370 | -3.617 | 0.00030 | 0.0181 |
| FOSL1 | 182.127 | -1.338 | 0.480 | -2.785 | 0.00535 | 0.0637 |
| C19orf70 | 525.179 | -1.336 | 0.450 | -2.970 | 0.00298 | 0.0468 |
| AURKB | 2334.466 | -1.336 | 0.323 | -4.133 | 0.00004 | 0.0086 |
| BCL2A1 | 286.379 | -1.331 | 0.494 | -2.693 | 0.00708 | 0.0732 |
| RPL23 | 27051.413 | -1.330 | 0.377 | -3.530 | 0.00042 | 0.0203 |
| TTK | 2117.816 | -1.329 | 0.465 | -2.857 | 0.00427 | 0.0562 |
| RPL8 | 62094.704 | -1.329 | 0.365 | -3.642 | 0.00027 | 0.0177 |
| GGH | 816.739 | -1.327 | 0.435 | -3.048 | 0.00230 | 0.0416 |
| TK1 | 3785.889 | -1.327 | 0.345 | -3.847 | 0.00012 | 0.0132 |
| CISD3 | 651.954 | -1.326 | 0.402 | -3.300 | 0.00097 | 0.0275 |
| PSMB3 | 3492.684 | -1.326 | 0.379 | -3.501 | 0.00046 | 0.0210 |
| S100A4 | 49264.691 | -1.325 | 0.490 | -2.703 | 0.00687 | 0.0718 |
| LOC100506844 | 740.173 | -1.324 | 0.435 | -3.042 | 0.00235 | 0.0421 |
| RPS5 | 29649.039 | -1.324 | 0.360 | -3.677 | 0.00024 | 0.0172 |
| SNRPB | 7340.838 | -1.324 | 0.359 | -3.683 | 0.00023 | 0.0170 |
| CACYBP | 7430.220 | -1.323 | 0.395 | -3.348 | 0.00081 | 0.0260 |
| MRPS15 | 3029.888 | -1.322 | 0.333 | -3.973 | 0.00007 | 0.0115 |
| TBC1D12 | 96.858 | -1.319 | 0.507 | -2.601 | 0.00928 | 0.0856 |
| H2AFZ | 15437.409 | -1.319 | 0.394 | -3.344 | 0.00083 | 0.0260 |
| TUBA1B | 13199.120 | -1.319 | 0.339 | -3.890 | 0.00010 | 0.0131 |
| RBX1 | 1558.678 | -1.315 | 0.404 | -3.256 | 0.00113 | 0.0292 |
| COX8A | 5399.961 | -1.314 | 0.317 | -4.143 | 0.00003 | 0.0086 |
| MAD2L2 | 1542.906 | -1.314 | 0.369 | -3.562 | 0.00037 | 0.0193 |
| COX7B | 3940.488 | -1.313 | 0.325 | -4.038 | 0.00005 | 0.0106 |
| BNIP1 | 829.186 | -1.312 | 0.390 | -3.366 | 0.00076 | 0.0253 |
| TMEM14C | 2780.807 | -1.311 | 0.405 | -3.234 | 0.00122 | 0.0305 |
| COX4I1 | 14707.377 | -1.309 | 0.361 | -3.631 | 0.00028 | 0.0179 |
| SAT1 | 7952.931 | -1.308 | 0.394 | -3.321 | 0.00090 | 0.0266 |
| HSPB1 | 2344.390 | -1.307 | 0.396 | -3.298 | 0.00097 | 0.0275 |
| POLR2J | 2008.985 | -1.305 | 0.377 | -3.466 | 0.00053 | 0.0222 |
| STRAP | 5269.141 | -1.303 | 0.415 | -3.142 | 0.00168 | 0.0358 |
| DUT | 7471.148 | -1.302 | 0.363 | -3.590 | 0.00033 | 0.0188 |
| CTSH | 3910.193 | -1.300 | 0.422 | -3.080 | 0.00207 | 0.0396 |
| NUDCD1 | 1221.607 | -1.300 | 0.393 | -3.308 | 0.00094 | 0.0272 |
| CAMLG | 5453.133 | -1.300 | 0.450 | -2.886 | 0.00391 | 0.0538 |
| DNAJC19 | 1453.425 | -1.299 | 0.404 | -3.218 | 0.00129 | 0.0311 |
| RPS16 | 42880.503 | -1.298 | 0.381 | -3.404 | 0.00067 | 0.0243 |
| SCCPDH | 1627.588 | -1.297 | 0.465 | -2.787 | 0.00532 | 0.0637 |
| TIMM13 | 1233.051 | -1.297 | 0.426 | -3.041 | 0.00236 | 0.0421 |
| C19orf48 | 2574.644 | -1.296 | 0.380 | -3.410 | 0.00065 | 0.0241 |
| ATP5G1 | 1055.053 | -1.296 | 0.402 | -3.228 | 0.00124 | 0.0307 |
| NDUFB3 | 2143.224 | -1.296 | 0.427 | -3.034 | 0.00242 | 0.0426 |
| CHPT1 | 1916.842 | -1.295 | 0.413 | -3.137 | 0.00171 | 0.0361 |
| IFI27L2 | 2805.992 | -1.295 | 0.413 | -3.135 | 0.00172 | 0.0362 |
| ANAPC11 | 1717.791 | -1.294 | 0.373 | -3.473 | 0.00052 | 0.0221 |
| MRPL17 | 2229.154 | -1.294 | 0.373 | -3.466 | 0.00053 | 0.0222 |
| PERP | 4588.083 | -1.293 | 0.393 | -3.290 | 0.00100 | 0.0279 |
| MIIP | 2299.128 | -1.291 | 0.227 | -5.689 | 0.00000 | 0.0001 |
| EXT1 | 1730.729 | -1.291 | 0.377 | -3.421 | 0.00062 | 0.0236 |
| NUF2 | 2914.298 | -1.288 | 0.443 | -2.907 | 0.00364 | 0.0520 |
| IMPDH2 | 8317.064 | -1.288 | 0.397 | -3.247 | 0.00117 | 0.0298 |
| EDF1 | 8538.429 | -1.287 | 0.406 | -3.170 | 0.00153 | 0.0344 |
| RPL21 | 43118.924 | -1.284 | 0.389 | -3.298 | 0.00097 | 0.0275 |
| EHD4 | 1643.898 | -1.283 | 0.467 | -2.749 | 0.00597 | 0.0672 |
| RPL34 | 34848.719 | -1.283 | 0.383 | -3.350 | 0.00081 | 0.0260 |
| C10orf58 | 616.057 | -1.282 | 0.377 | -3.401 | 0.00067 | 0.0244 |
| BANF1 | 2113.402 | -1.282 | 0.334 | -3.841 | 0.00012 | 0.0132 |
| PPM1E | 63.138 | -1.281 | 0.433 | -2.961 | 0.00307 | 0.0476 |
| RPS20 | 53089.544 | -1.280 | 0.394 | -3.248 | 0.00116 | 0.0298 |
| FTH1 | 48472.044 | -1.280 | 0.419 | -3.056 | 0.00224 | 0.0407 |
| TRIP13 | 716.342 | -1.277 | 0.464 | -2.753 | 0.00591 | 0.0667 |
| NQO1 | 954.275 | -1.275 | 0.403 | -3.161 | 0.00157 | 0.0348 |
| RPL37 | 41404.869 | -1.274 | 0.404 | -3.155 | 0.00160 | 0.0354 |
| CCDC141 | 396.108 | -1.273 | 0.438 | -2.903 | 0.00369 | 0.0521 |
| SPAG5 | 1771.436 | -1.272 | 0.418 | -3.041 | 0.00236 | 0.0421 |
| MTCP1NB | 563.405 | -1.270 | 0.360 | -3.528 | 0.00042 | 0.0203 |
| NFU1 | 1111.522 | -1.270 | 0.363 | -3.498 | 0.00047 | 0.0210 |
| NUDT15 | 918.742 | -1.269 | 0.423 | -3.002 | 0.00268 | 0.0442 |
| POP7 | 758.882 | -1.268 | 0.348 | -3.640 | 0.00027 | 0.0177 |
| POLD2 | 2537.904 | -1.267 | 0.347 | -3.654 | 0.00026 | 0.0176 |
| SLC2A13 | 284.923 | -1.265 | 0.427 | -2.964 | 0.00304 | 0.0474 |
| RFC4 | 2079.990 | -1.264 | 0.329 | -3.842 | 0.00012 | 0.0132 |
| GCSH | 92.874 | -1.264 | 0.439 | -2.881 | 0.00397 | 0.0542 |
| C14orf166 | 7244.891 | -1.264 | 0.351 | -3.600 | 0.00032 | 0.0184 |
| FTL | 80540.862 | -1.263 | 0.375 | -3.368 | 0.00076 | 0.0252 |
| NET1 | 2254.145 | -1.262 | 0.448 | -2.817 | 0.00485 | 0.0599 |
| RPS8 | 67850.002 | -1.260 | 0.363 | -3.470 | 0.00052 | 0.0221 |
| PGD | 3192.638 | -1.259 | 0.406 | -3.105 | 0.00190 | 0.0378 |
| JUNB | 4868.934 | -1.259 | 0.395 | -3.186 | 0.00144 | 0.0331 |
| C7orf11 | 1259.400 | -1.257 | 0.367 | -3.423 | 0.00062 | 0.0236 |
| PAFAH1B3 | 777.915 | -1.256 | 0.349 | -3.600 | 0.00032 | 0.0184 |
| SRP19 | 3168.133 | -1.255 | 0.397 | -3.161 | 0.00157 | 0.0348 |
| COX6B1 | 6325.790 | -1.254 | 0.393 | -3.193 | 0.00141 | 0.0330 |
| TCOF1 | 3976.372 | -1.252 | 0.414 | -3.027 | 0.00247 | 0.0429 |
| GLRX2 | 580.260 | -1.252 | 0.451 | -2.778 | 0.00547 | 0.0642 |
| RPL10A | 35388.841 | -1.249 | 0.361 | -3.461 | 0.00054 | 0.0223 |
| MRPL14 | 1140.296 | -1.247 | 0.410 | -3.039 | 0.00237 | 0.0422 |
| RMI2 | 1158.172 | -1.247 | 0.341 | -3.659 | 0.00025 | 0.0176 |
| COX7C | 12164.131 | -1.246 | 0.396 | -3.148 | 0.00164 | 0.0357 |
| RND1 | 81.194 | -1.246 | 0.421 | -2.957 | 0.00310 | 0.0478 |
| EEF1B2 | 26167.932 | -1.245 | 0.377 | -3.305 | 0.00095 | 0.0272 |
| ENY2 | 3706.522 | -1.244 | 0.374 | -3.328 | 0.00088 | 0.0264 |
| RDX | 7058.919 | -1.243 | 0.471 | -2.641 | 0.00826 | 0.0799 |
| LSM5 | 2347.777 | -1.242 | 0.357 | -3.483 | 0.00050 | 0.0216 |
| NDUFA12 | 3760.528 | -1.240 | 0.477 | -2.598 | 0.00936 | 0.0861 |
| SMS | 4862.615 | -1.239 | 0.409 | -3.028 | 0.00246 | 0.0429 |
| UBL5 | 3440.511 | -1.238 | 0.358 | -3.462 | 0.00054 | 0.0223 |
| VAMP5 | 2350.984 | -1.238 | 0.402 | -3.079 | 0.00208 | 0.0396 |
| OSGIN2 | 2249.838 | -1.237 | 0.437 | -2.835 | 0.00459 | 0.0582 |
| SNRPB2 | 4546.711 | -1.237 | 0.378 | -3.274 | 0.00106 | 0.0287 |
| RPL32 | 51184.947 | -1.237 | 0.379 | -3.262 | 0.00111 | 0.0290 |
| DTL | 2386.216 | -1.236 | 0.474 | -2.611 | 0.00903 | 0.0843 |
| CDC6 | 1335.299 | -1.236 | 0.425 | -2.906 | 0.00366 | 0.0521 |
| DNAJC9 | 2607.503 | -1.235 | 0.391 | -3.162 | 0.00156 | 0.0348 |
| NPM1 | 60235.535 | -1.235 | 0.386 | -3.199 | 0.00138 | 0.0327 |
| HSPH1 | 4656.018 | -1.235 | 0.289 | -4.279 | 0.00002 | 0.0075 |
| BOD1 | 1100.091 | -1.235 | 0.397 | -3.107 | 0.00189 | 0.0378 |
| PNO1 | 495.466 | -1.233 | 0.422 | -2.923 | 0.00347 | 0.0508 |
| RPL27A | 54526.370 | -1.231 | 0.381 | -3.234 | 0.00122 | 0.0305 |
| PPP1R14B | 2699.477 | -1.231 | 0.400 | -3.079 | 0.00208 | 0.0396 |
| NUP37 | 1893.524 | -1.231 | 0.430 | -2.859 | 0.00425 | 0.0562 |
| C14orf80 | 537.841 | -1.230 | 0.300 | -4.106 | 0.00004 | 0.0090 |
| SOD1 | 8066.070 | -1.230 | 0.385 | -3.195 | 0.00140 | 0.0329 |
| ADK | 4082.463 | -1.229 | 0.379 | -3.242 | 0.00119 | 0.0301 |
| C19orf79 | 1341.220 | -1.229 | 0.404 | -3.044 | 0.00233 | 0.0419 |
| RPL31 | 46227.250 | -1.229 | 0.365 | -3.362 | 0.00077 | 0.0254 |
| PSMC3 | 3467.929 | -1.226 | 0.343 | -3.578 | 0.00035 | 0.0191 |
| GMNN | 2767.531 | -1.226 | 0.377 | -3.255 | 0.00113 | 0.0292 |
| SNRPE | 3682.029 | -1.226 | 0.363 | -3.375 | 0.00074 | 0.0250 |
| FH | 2247.884 | -1.224 | 0.344 | -3.562 | 0.00037 | 0.0193 |
| CCNB1 | 2664.716 | -1.224 | 0.346 | -3.541 | 0.00040 | 0.0199 |
| TNFSF9 | 469.425 | -1.224 | 0.367 | -3.336 | 0.00085 | 0.0261 |
| ING3 | 3618.734 | -1.219 | 0.465 | -2.620 | 0.00879 | 0.0832 |
| MCM10 | 874.112 | -1.219 | 0.445 | -2.738 | 0.00619 | 0.0686 |
| LSM10 | 1270.908 | -1.216 | 0.371 | -3.281 | 0.00103 | 0.0284 |
| HSD17B10 | 1943.079 | -1.216 | 0.377 | -3.226 | 0.00126 | 0.0307 |
| LAMTOR2 | 1634.035 | -1.214 | 0.355 | -3.420 | 0.00063 | 0.0236 |
| RPL24 | 30529.939 | -1.214 | 0.380 | -3.193 | 0.00141 | 0.0330 |
| RPSA | 91324.857 | -1.214 | 0.357 | -3.398 | 0.00068 | 0.0245 |
| ZWINT | 2605.738 | -1.213 | 0.402 | -3.015 | 0.00257 | 0.0434 |
| NDUFA4 | 4881.065 | -1.212 | 0.336 | -3.609 | 0.00031 | 0.0183 |
| TUBG1 | 784.478 | -1.211 | 0.325 | -3.727 | 0.00019 | 0.0159 |
| CCDC167 | 1382.511 | -1.211 | 0.360 | -3.363 | 0.00077 | 0.0254 |
| MED10 | 5114.914 | -1.211 | 0.427 | -2.837 | 0.00456 | 0.0580 |
| IKBIP | 2065.217 | -1.210 | 0.451 | -2.683 | 0.00729 | 0.0744 |
| RPS3 | 104064.219 | -1.209 | 0.350 | -3.453 | 0.00055 | 0.0226 |
| ERH | 4522.890 | -1.209 | 0.392 | -3.086 | 0.00203 | 0.0394 |
| RPA3 | 1726.403 | -1.207 | 0.386 | -3.128 | 0.00176 | 0.0366 |
| NHP2 | 1853.978 | -1.204 | 0.381 | -3.162 | 0.00157 | 0.0348 |
| UQCRFS1 | 3661.435 | -1.204 | 0.361 | -3.336 | 0.00085 | 0.0261 |
| RPL18 | 44601.615 | -1.204 | 0.361 | -3.334 | 0.00086 | 0.0262 |
| MGMT | 1026.871 | -1.203 | 0.393 | -3.061 | 0.00220 | 0.0406 |
| SKA3 | 956.973 | -1.203 | 0.397 | -3.030 | 0.00244 | 0.0428 |
| RBM3 | 15628.976 | -1.203 | 0.432 | -2.787 | 0.00533 | 0.0637 |
| PRDX5 | 6010.504 | -1.201 | 0.362 | -3.315 | 0.00092 | 0.0268 |
| C6orf226 | 246.913 | -1.200 | 0.269 | -4.464 | 0.00001 | 0.0056 |
| KCNK1 | 482.747 | -1.199 | 0.410 | -2.923 | 0.00346 | 0.0508 |
| RGS10 | 7855.271 | -1.199 | 0.425 | -2.820 | 0.00480 | 0.0595 |
| MTFP1 | 4550.092 | -1.197 | 0.341 | -3.509 | 0.00045 | 0.0207 |
| C9orf16 | 2630.960 | -1.197 | 0.310 | -3.863 | 0.00011 | 0.0132 |
| GMDS | 1249.821 | -1.197 | 0.331 | -3.611 | 0.00030 | 0.0183 |
| NSMCE2 | 2582.940 | -1.197 | 0.380 | -3.147 | 0.00165 | 0.0357 |
| MRPL11 | 1896.551 | -1.194 | 0.355 | -3.363 | 0.00077 | 0.0254 |
| AVEN | 401.686 | -1.194 | 0.400 | -2.984 | 0.00284 | 0.0458 |
| NFKBIA | 3495.488 | -1.193 | 0.341 | -3.503 | 0.00046 | 0.0210 |
| C6orf125 | 553.704 | -1.193 | 0.362 | -3.294 | 0.00099 | 0.0277 |
| MRPL51 | 3140.930 | -1.193 | 0.364 | -3.276 | 0.00105 | 0.0286 |
| RPS15 | 24045.647 | -1.193 | 0.350 | -3.407 | 0.00066 | 0.0243 |
| TMEM106C | 3561.715 | -1.193 | 0.338 | -3.532 | 0.00041 | 0.0203 |
| RPLP2 | 45817.799 | -1.192 | 0.369 | -3.232 | 0.00123 | 0.0305 |
| ZNFX1-AS1 | 8255.997 | -1.192 | 0.366 | -3.259 | 0.00112 | 0.0292 |
| RPF2 | 1148.211 | -1.191 | 0.419 | -2.844 | 0.00446 | 0.0575 |
| HRAS | 1412.078 | -1.190 | 0.310 | -3.841 | 0.00012 | 0.0132 |
| C6orf48 | 7078.357 | -1.190 | 0.383 | -3.104 | 0.00191 | 0.0378 |
| TCEAL1 | 428.196 | -1.190 | 0.272 | -4.373 | 0.00001 | 0.0064 |
| C1orf31 | 1264.207 | -1.189 | 0.398 | -2.986 | 0.00283 | 0.0458 |
| TIPIN | 512.671 | -1.188 | 0.405 | -2.933 | 0.00336 | 0.0500 |
| EPB41L2 | 1553.418 | -1.187 | 0.342 | -3.472 | 0.00052 | 0.0221 |
| EBP | 4501.299 | -1.187 | 0.320 | -3.708 | 0.00021 | 0.0165 |
| GSTCD | 872.376 | -1.186 | 0.429 | -2.767 | 0.00566 | 0.0652 |
| HSPD1 | 16266.589 | -1.186 | 0.390 | -3.045 | 0.00233 | 0.0418 |
| IPO5 | 8354.761 | -1.185 | 0.440 | -2.696 | 0.00702 | 0.0729 |
| NDUFAF4 | 483.203 | -1.185 | 0.387 | -3.061 | 0.00221 | 0.0406 |
| GAMT | 758.745 | -1.184 | 0.386 | -3.070 | 0.00214 | 0.0399 |
| AURKAIP1 | 2516.083 | -1.183 | 0.367 | -3.226 | 0.00126 | 0.0307 |
| MRPL23 | 2159.712 | -1.182 | 0.335 | -3.524 | 0.00043 | 0.0203 |
| RPL27 | 31505.985 | -1.181 | 0.355 | -3.322 | 0.00089 | 0.0266 |
| RPL41 | 20952.610 | -1.179 | 0.353 | -3.338 | 0.00084 | 0.0261 |
| RPL13 | 77535.044 | -1.179 | 0.330 | -3.567 | 0.00036 | 0.0193 |
| TUBB | 49851.033 | -1.178 | 0.373 | -3.163 | 0.00156 | 0.0348 |
| FAU | 22576.861 | -1.178 | 0.346 | -3.409 | 0.00065 | 0.0241 |
| UQCRB | 16385.223 | -1.177 | 0.387 | -3.041 | 0.00236 | 0.0421 |
| SLIRP | 950.544 | -1.177 | 0.323 | -3.638 | 0.00027 | 0.0177 |
| COPS3 | 3959.794 | -1.176 | 0.370 | -3.181 | 0.00147 | 0.0333 |
| UQCRH | 8214.439 | -1.176 | 0.374 | -3.142 | 0.00168 | 0.0358 |
| MIS18A | 636.132 | -1.176 | 0.438 | -2.688 | 0.00720 | 0.0739 |
| PSMD9 | 2638.104 | -1.174 | 0.347 | -3.387 | 0.00071 | 0.0246 |
| NOP10 | 2566.547 | -1.174 | 0.374 | -3.141 | 0.00169 | 0.0358 |
| CDCA8 | 2518.028 | -1.173 | 0.378 | -3.101 | 0.00193 | 0.0382 |
| HSP90AA1 | 31235.240 | -1.173 | 0.403 | -2.907 | 0.00365 | 0.0520 |
| EIF1B | 2181.956 | -1.173 | 0.374 | -3.137 | 0.00171 | 0.0361 |
| SGOL1 | 1970.775 | -1.172 | 0.415 | -2.823 | 0.00476 | 0.0593 |
| NR1D1 | 647.840 | -1.170 | 0.404 | -2.895 | 0.00379 | 0.0529 |
| ECI1 | 1181.530 | -1.170 | 0.320 | -3.659 | 0.00025 | 0.0176 |
| CDCA3 | 1043.332 | -1.169 | 0.310 | -3.774 | 0.00016 | 0.0145 |
| LSM2 | 2632.477 | -1.169 | 0.347 | -3.370 | 0.00075 | 0.0252 |
| COPS2 | 7651.062 | -1.168 | 0.436 | -2.680 | 0.00736 | 0.0747 |
| UBE2D2 | 9523.401 | -1.168 | 0.419 | -2.785 | 0.00535 | 0.0637 |
| NENF | 1530.323 | -1.167 | 0.422 | -2.768 | 0.00564 | 0.0652 |
| C19orf53 | 2930.623 | -1.165 | 0.334 | -3.494 | 0.00048 | 0.0211 |
| PIBF1 | 1895.698 | -1.165 | 0.406 | -2.870 | 0.00410 | 0.0553 |
| SEC61G | 6929.337 | -1.164 | 0.355 | -3.276 | 0.00105 | 0.0286 |
| RAD54B | 1344.660 | -1.164 | 0.431 | -2.700 | 0.00692 | 0.0721 |
| ATP5J | 3104.963 | -1.164 | 0.406 | -2.866 | 0.00416 | 0.0557 |
| POLE4 | 959.378 | -1.162 | 0.318 | -3.656 | 0.00026 | 0.0176 |
| TRMT112 | 3521.517 | -1.162 | 0.350 | -3.323 | 0.00089 | 0.0266 |
| HLA-DRA | 22525.817 | -1.160 | 0.355 | -3.267 | 0.00109 | 0.0288 |
| RPL7A | 78297.813 | -1.160 | 0.347 | -3.346 | 0.00082 | 0.0260 |
| MTHFD1L | 2751.094 | -1.159 | 0.397 | -2.921 | 0.00349 | 0.0510 |
| CENPA | 1052.330 | -1.157 | 0.376 | -3.074 | 0.00211 | 0.0398 |
| UBE2B | 7084.878 | -1.155 | 0.403 | -2.868 | 0.00413 | 0.0555 |
| S100A6 | 23094.957 | -1.154 | 0.368 | -3.139 | 0.00170 | 0.0360 |
| HMGB3 | 925.374 | -1.154 | 0.405 | -2.848 | 0.00440 | 0.0570 |
| TAOK3 | 21038.567 | -1.152 | 0.365 | -3.161 | 0.00157 | 0.0348 |
| RNASEK-C17ORF49 | 7477.579 | -1.152 | 0.323 | -3.563 | 0.00037 | 0.0193 |
| CRY1 | 1497.127 | -1.150 | 0.378 | -3.042 | 0.00235 | 0.0421 |
| GLRX5 | 1864.125 | -1.150 | 0.415 | -2.772 | 0.00557 | 0.0649 |
| RELB | 1670.230 | -1.150 | 0.370 | -3.107 | 0.00189 | 0.0378 |
| C15orf23 | 893.378 | -1.149 | 0.335 | -3.433 | 0.00060 | 0.0232 |
| FAM96B | 3101.033 | -1.149 | 0.330 | -3.487 | 0.00049 | 0.0214 |
| SRM | 1600.863 | -1.149 | 0.364 | -3.152 | 0.00162 | 0.0355 |
| FAM158A | 1124.433 | -1.146 | 0.342 | -3.347 | 0.00082 | 0.0260 |
| STOML2 | 3422.512 | -1.146 | 0.337 | -3.398 | 0.00068 | 0.0245 |
| SQRDL | 9881.931 | -1.146 | 0.395 | -2.901 | 0.00372 | 0.0524 |
| TALDO1 | 6613.118 | -1.146 | 0.334 | -3.430 | 0.00060 | 0.0232 |
| IGFBP4 | 2594.579 | -1.145 | 0.339 | -3.378 | 0.00073 | 0.0250 |
| POMP | 4105.985 | -1.144 | 0.302 | -3.784 | 0.00015 | 0.0141 |
| HIST1H1C | 1238.622 | -1.141 | 0.328 | -3.480 | 0.00050 | 0.0216 |
| CHCHD5 | 1036.228 | -1.141 | 0.405 | -2.819 | 0.00482 | 0.0597 |
| DRAP1 | 3866.222 | -1.140 | 0.320 | -3.561 | 0.00037 | 0.0193 |
| FBL | 10594.602 | -1.140 | 0.331 | -3.446 | 0.00057 | 0.0230 |
| RPL29 | 41356.715 | -1.138 | 0.338 | -3.371 | 0.00075 | 0.0252 |
| CD74 | 75812.309 | -1.138 | 0.304 | -3.747 | 0.00018 | 0.0152 |
| NINJ1 | 2690.041 | -1.138 | 0.381 | -2.987 | 0.00281 | 0.0458 |
| STX8 | 2179.702 | -1.138 | 0.386 | -2.949 | 0.00319 | 0.0482 |
| WDR34 | 914.844 | -1.137 | 0.291 | -3.904 | 0.00009 | 0.0130 |
| RPS18 | 84635.196 | -1.137 | 0.335 | -3.394 | 0.00069 | 0.0246 |
| OLA1 | 6001.628 | -1.136 | 0.395 | -2.873 | 0.00406 | 0.0549 |
| UBE2D1 | 1029.578 | -1.135 | 0.417 | -2.719 | 0.00655 | 0.0704 |
| LSM1 | 1679.317 | -1.135 | 0.393 | -2.889 | 0.00386 | 0.0535 |
| WDR76 | 1934.059 | -1.134 | 0.390 | -2.910 | 0.00361 | 0.0519 |
| NEIL3 | 1174.254 | -1.132 | 0.424 | -2.674 | 0.00750 | 0.0756 |
| UPF3B | 2614.731 | -1.131 | 0.407 | -2.777 | 0.00548 | 0.0642 |
| ITGAE | 3159.086 | -1.130 | 0.294 | -3.849 | 0.00012 | 0.0132 |
| PSMD14 | 4821.832 | -1.130 | 0.367 | -3.080 | 0.00207 | 0.0396 |
| KPNA2 | 4982.367 | -1.129 | 0.314 | -3.592 | 0.00033 | 0.0187 |
| RPL39 | 18718.508 | -1.129 | 0.378 | -2.984 | 0.00284 | 0.0458 |
| BLVRA | 1118.551 | -1.129 | 0.299 | -3.772 | 0.00016 | 0.0145 |
| HLA-DPB1 | 11271.640 | -1.128 | 0.373 | -3.021 | 0.00252 | 0.0431 |
| PSMB7 | 2816.890 | -1.127 | 0.328 | -3.432 | 0.00060 | 0.0232 |
| YWHAE | 10664.700 | -1.127 | 0.368 | -3.062 | 0.00220 | 0.0405 |
| C6orf1 | 1413.703 | -1.125 | 0.365 | -3.082 | 0.00206 | 0.0396 |
| TOMM22 | 2236.930 | -1.124 | 0.381 | -2.952 | 0.00315 | 0.0480 |
| H3F3B | 30477.520 | -1.123 | 0.380 | -2.953 | 0.00315 | 0.0480 |
| SMYD3 | 2389.322 | -1.123 | 0.364 | -3.083 | 0.00205 | 0.0396 |
| NAA10 | 1681.870 | -1.123 | 0.311 | -3.608 | 0.00031 | 0.0183 |
| RPS23 | 55296.248 | -1.122 | 0.376 | -2.984 | 0.00285 | 0.0458 |
| GRHPR | 2435.835 | -1.122 | 0.295 | -3.797 | 0.00015 | 0.0138 |
| KIF2C | 3743.969 | -1.122 | 0.410 | -2.734 | 0.00627 | 0.0687 |
| AURKA | 2433.509 | -1.121 | 0.402 | -2.786 | 0.00534 | 0.0637 |
| FBXO6 | 566.242 | -1.121 | 0.293 | -3.831 | 0.00013 | 0.0132 |
| CAMTA1 | 1348.476 | -1.121 | 0.316 | -3.552 | 0.00038 | 0.0195 |
| STK39 | 5864.071 | -1.120 | 0.404 | -2.770 | 0.00561 | 0.0652 |
| PKMYT1 | 1592.683 | -1.119 | 0.391 | -2.862 | 0.00421 | 0.0560 |
| HNRNPA1 | 73135.244 | -1.119 | 0.410 | -2.731 | 0.00631 | 0.0689 |
| NDUFAB1 | 2069.925 | -1.118 | 0.351 | -3.185 | 0.00145 | 0.0331 |
| SRP14 | 10741.906 | -1.118 | 0.362 | -3.089 | 0.00201 | 0.0392 |
| C11orf51 | 572.922 | -1.116 | 0.336 | -3.317 | 0.00091 | 0.0267 |
| POLR2F | 807.462 | -1.116 | 0.399 | -2.796 | 0.00517 | 0.0623 |
| FAM177A1 | 1944.400 | -1.115 | 0.412 | -2.709 | 0.00676 | 0.0712 |
| SPATS2L | 1795.269 | -1.115 | 0.396 | -2.818 | 0.00483 | 0.0598 |
| MRPS23 | 1125.501 | -1.113 | 0.382 | -2.912 | 0.00359 | 0.0518 |
| C14orf1 | 3043.087 | -1.113 | 0.410 | -2.712 | 0.00668 | 0.0709 |
| APOA1BP | 1308.747 | -1.112 | 0.353 | -3.149 | 0.00164 | 0.0356 |
| RPS12 | 55803.378 | -1.110 | 0.348 | -3.186 | 0.00144 | 0.0331 |
| PTS | 945.536 | -1.109 | 0.390 | -2.843 | 0.00446 | 0.0575 |
| TMEM126A | 1302.601 | -1.109 | 0.352 | -3.150 | 0.00163 | 0.0356 |
| TIMM17B | 1832.745 | -1.108 | 0.268 | -4.137 | 0.00004 | 0.0086 |
| NT5C3 | 416.307 | -1.107 | 0.334 | -3.320 | 0.00090 | 0.0266 |
| RPS19BP1 | 3429.174 | -1.106 | 0.285 | -3.882 | 0.00010 | 0.0132 |
| LOC100287482 | 151.946 | -1.106 | 0.421 | -2.626 | 0.00863 | 0.0824 |
| NUTF2 | 2298.744 | -1.106 | 0.327 | -3.376 | 0.00073 | 0.0250 |
| SNRPC | 3212.758 | -1.105 | 0.326 | -3.389 | 0.00070 | 0.0246 |
| TIMM8B | 589.314 | -1.105 | 0.360 | -3.065 | 0.00218 | 0.0403 |
| ARL6IP4 | 7028.262 | -1.102 | 0.301 | -3.660 | 0.00025 | 0.0176 |
| CCDC109B | 6154.926 | -1.102 | 0.413 | -2.669 | 0.00760 | 0.0761 |
| B2M | 482465.570 | -1.101 | 0.421 | -2.617 | 0.00887 | 0.0836 |
| NDUFAF2 | 536.722 | -1.101 | 0.379 | -2.903 | 0.00369 | 0.0521 |
| UQCR10 | 2251.957 | -1.101 | 0.387 | -2.844 | 0.00445 | 0.0575 |
| UBA52 | 35305.692 | -1.101 | 0.378 | -2.915 | 0.00355 | 0.0515 |
| FAM100B | 5747.159 | -1.100 | 0.293 | -3.760 | 0.00017 | 0.0148 |
| HINT2 | 1497.369 | -1.098 | 0.374 | -2.939 | 0.00330 | 0.0493 |
| NDUFA6 | 2014.774 | -1.097 | 0.364 | -3.010 | 0.00261 | 0.0436 |
| DTD1 | 2471.910 | -1.096 | 0.353 | -3.106 | 0.00190 | 0.0378 |
| RRS1 | 432.925 | -1.095 | 0.423 | -2.591 | 0.00957 | 0.0872 |
| PRDX1 | 7381.348 | -1.095 | 0.348 | -3.145 | 0.00166 | 0.0357 |
| PSMB9 | 6170.048 | -1.094 | 0.330 | -3.317 | 0.00091 | 0.0267 |
| SEC61B | 5678.713 | -1.094 | 0.340 | -3.219 | 0.00128 | 0.0310 |
| DDIT3 | 1031.263 | -1.094 | 0.374 | -2.927 | 0.00343 | 0.0505 |
| CDK4 | 2743.585 | -1.094 | 0.334 | -3.279 | 0.00104 | 0.0286 |
| RNF7 | 4257.721 | -1.094 | 0.356 | -3.072 | 0.00212 | 0.0399 |
| PFDN5 | 11085.528 | -1.092 | 0.375 | -2.914 | 0.00357 | 0.0517 |
| SKP1 | 8160.146 | -1.089 | 0.356 | -3.057 | 0.00224 | 0.0407 |
| GSTO1 | 3866.824 | -1.087 | 0.349 | -3.119 | 0.00181 | 0.0370 |
| SPATA5 | 591.418 | -1.085 | 0.298 | -3.643 | 0.00027 | 0.0177 |
| GLRX | 3603.560 | -1.084 | 0.363 | -2.987 | 0.00282 | 0.0458 |
| KCMF1 | 6248.356 | -1.084 | 0.372 | -2.913 | 0.00358 | 0.0518 |
| RBMX2 | 2559.120 | -1.082 | 0.325 | -3.329 | 0.00087 | 0.0264 |
| ILF2 | 5346.828 | -1.082 | 0.332 | -3.263 | 0.00110 | 0.0290 |
| HSP90AB1 | 34684.909 | -1.079 | 0.406 | -2.659 | 0.00783 | 0.0774 |
| BATF | 3674.318 | -1.077 | 0.364 | -2.959 | 0.00308 | 0.0477 |
| PNKD | 1193.906 | -1.076 | 0.337 | -3.196 | 0.00139 | 0.0329 |
| LGALS1 | 48905.365 | -1.072 | 0.333 | -3.217 | 0.00129 | 0.0312 |
| C16orf80 | 1365.585 | -1.071 | 0.362 | -2.958 | 0.00310 | 0.0478 |
| NRBF2 | 2280.834 | -1.071 | 0.394 | -2.714 | 0.00664 | 0.0708 |
| NAP1L1 | 63580.701 | -1.070 | 0.360 | -2.975 | 0.00293 | 0.0465 |
| MRPL47 | 1250.772 | -1.068 | 0.323 | -3.308 | 0.00094 | 0.0272 |
| FDXR | 4119.806 | -1.068 | 0.389 | -2.742 | 0.00610 | 0.0679 |
| FAH | 2476.280 | -1.068 | 0.376 | -2.838 | 0.00453 | 0.0579 |
| OBFC2B | 2176.155 | -1.068 | 0.316 | -3.384 | 0.00072 | 0.0246 |
| SUMO3 | 4679.626 | -1.068 | 0.373 | -2.860 | 0.00423 | 0.0561 |
| PMVK | 1682.869 | -1.066 | 0.338 | -3.151 | 0.00163 | 0.0356 |
| C11orf1 | 419.220 | -1.066 | 0.374 | -2.850 | 0.00437 | 0.0568 |
| RPLP0 | 151927.831 | -1.065 | 0.331 | -3.221 | 0.00128 | 0.0310 |
| CUTA | 8713.479 | -1.065 | 0.309 | -3.442 | 0.00058 | 0.0230 |
| MDK | 112.116 | -1.063 | 0.391 | -2.720 | 0.00652 | 0.0703 |
| RAB11A | 5751.106 | -1.062 | 0.357 | -2.975 | 0.00293 | 0.0465 |
| CLEC2B | 8760.050 | -1.062 | 0.328 | -3.238 | 0.00120 | 0.0303 |
| UQCR11 | 2788.963 | -1.062 | 0.364 | -2.921 | 0.00348 | 0.0510 |
| KATNA1 | 1307.732 | -1.062 | 0.260 | -4.079 | 0.00005 | 0.0098 |
| NME4 | 628.801 | -1.061 | 0.373 | -2.844 | 0.00446 | 0.0575 |
| RPS13 | 26055.024 | -1.061 | 0.355 | -2.993 | 0.00276 | 0.0451 |
| TIPRL | 5003.265 | -1.061 | 0.384 | -2.761 | 0.00575 | 0.0655 |
| CDC45 | 1613.729 | -1.061 | 0.394 | -2.696 | 0.00701 | 0.0729 |
| NDUFS8 | 1782.673 | -1.060 | 0.317 | -3.340 | 0.00084 | 0.0260 |
| MRPL15 | 1271.332 | -1.060 | 0.350 | -3.027 | 0.00247 | 0.0429 |
| LINC00493 | 1537.924 | -1.059 | 0.390 | -2.714 | 0.00665 | 0.0708 |
| DRG1 | 2475.774 | -1.057 | 0.300 | -3.520 | 0.00043 | 0.0203 |
| TPX2 | 6223.742 | -1.056 | 0.380 | -2.779 | 0.00546 | 0.0642 |
| RPL26 | 47573.324 | -1.055 | 0.354 | -2.979 | 0.00289 | 0.0461 |
| EIF4E | 2347.497 | -1.055 | 0.398 | -2.652 | 0.00801 | 0.0783 |
| MRPS26 | 845.636 | -1.054 | 0.282 | -3.740 | 0.00018 | 0.0153 |
| TRAPPC2L | 1585.449 | -1.054 | 0.350 | -3.015 | 0.00257 | 0.0434 |
| RPS24 | 48771.617 | -1.054 | 0.373 | -2.822 | 0.00477 | 0.0593 |
| NDUFA7 | 1761.327 | -1.053 | 0.305 | -3.449 | 0.00056 | 0.0228 |
| EEF1D | 24079.229 | -1.052 | 0.342 | -3.078 | 0.00209 | 0.0396 |
| MRPS35 | 2525.930 | -1.052 | 0.360 | -2.927 | 0.00343 | 0.0505 |
| ATP5D | 3175.317 | -1.050 | 0.339 | -3.102 | 0.00192 | 0.0380 |
| RAD51 | 1054.814 | -1.050 | 0.388 | -2.706 | 0.00682 | 0.0716 |
| RPLP1 | 82968.585 | -1.050 | 0.329 | -3.188 | 0.00143 | 0.0331 |
| NDFIP1 | 9258.570 | -1.049 | 0.355 | -2.953 | 0.00315 | 0.0480 |
| RPL14 | 51172.839 | -1.049 | 0.345 | -3.037 | 0.00239 | 0.0423 |
| A1BG | 587.967 | -1.048 | 0.392 | -2.671 | 0.00756 | 0.0758 |
| DCXR | 1255.872 | -1.048 | 0.325 | -3.226 | 0.00126 | 0.0307 |
| CDC25C | 586.379 | -1.048 | 0.365 | -2.869 | 0.00412 | 0.0554 |
| CXCR4 | 12883.864 | -1.047 | 0.292 | -3.590 | 0.00033 | 0.0188 |
| AP3S1 | 9844.790 | -1.047 | 0.402 | -2.604 | 0.00922 | 0.0854 |
| RPL7 | 71323.611 | -1.046 | 0.366 | -2.857 | 0.00428 | 0.0562 |
| EXOC6B | 598.398 | -1.046 | 0.243 | -4.311 | 0.00002 | 0.0070 |
| RNF181 | 4387.734 | -1.046 | 0.328 | -3.188 | 0.00143 | 0.0331 |
| EIF2B3 | 637.507 | -1.046 | 0.324 | -3.228 | 0.00125 | 0.0307 |
| CCDC59 | 1811.053 | -1.046 | 0.405 | -2.582 | 0.00983 | 0.0884 |
| MYL12B | 15510.926 | -1.045 | 0.337 | -3.097 | 0.00195 | 0.0385 |
| COQ3 | 316.048 | -1.044 | 0.286 | -3.656 | 0.00026 | 0.0176 |
| SET | 22275.067 | -1.044 | 0.398 | -2.622 | 0.00875 | 0.0830 |
| COX5B | 4076.119 | -1.043 | 0.307 | -3.397 | 0.00068 | 0.0245 |
| RPS15A | 63643.325 | -1.042 | 0.333 | -3.131 | 0.00174 | 0.0364 |
| SUB1 | 15044.750 | -1.042 | 0.381 | -2.734 | 0.00625 | 0.0686 |
| MZT2B | 3753.587 | -1.042 | 0.256 | -4.065 | 0.00005 | 0.0100 |
| SHFM1 | 5362.864 | -1.042 | 0.335 | -3.113 | 0.00185 | 0.0375 |
| RPL11 | 63056.781 | -1.042 | 0.351 | -2.968 | 0.00300 | 0.0469 |
| CWC15 | 2996.789 | -1.042 | 0.329 | -3.166 | 0.00155 | 0.0347 |
| LOC100129250 | 459.254 | -1.041 | 0.383 | -2.721 | 0.00651 | 0.0703 |
| RUVBL2 | 2462.034 | -1.041 | 0.291 | -3.574 | 0.00035 | 0.0191 |
| LOC550643 | 1436.118 | -1.041 | 0.324 | -3.209 | 0.00133 | 0.0319 |
| PFDN6 | 1040.840 | -1.040 | 0.367 | -2.832 | 0.00462 | 0.0586 |
| CYC1 | 2892.977 | -1.039 | 0.307 | -3.384 | 0.00071 | 0.0246 |
| TAF10 | 2100.856 | -1.038 | 0.321 | -3.232 | 0.00123 | 0.0305 |
| EIF4A2 | 25141.772 | -1.038 | 0.403 | -2.577 | 0.00997 | 0.0890 |
| TESC | 1056.278 | -1.038 | 0.376 | -2.764 | 0.00571 | 0.0652 |
| PEBP1 | 6488.299 | -1.038 | 0.311 | -3.340 | 0.00084 | 0.0260 |
| CCDC6 | 4402.501 | -1.036 | 0.401 | -2.582 | 0.00982 | 0.0884 |
| PRDX2 | 7094.294 | -1.033 | 0.290 | -3.557 | 0.00038 | 0.0193 |
| TRIP10 | 2053.559 | -1.032 | 0.312 | -3.307 | 0.00094 | 0.0272 |
| CCDC107 | 501.515 | -1.032 | 0.245 | -4.211 | 0.00003 | 0.0082 |
| UQCRC1 | 6841.548 | -1.032 | 0.288 | -3.581 | 0.00034 | 0.0190 |
| MCM5 | 5995.651 | -1.032 | 0.343 | -3.005 | 0.00265 | 0.0439 |
| SAC3D1 | 531.378 | -1.030 | 0.330 | -3.118 | 0.00182 | 0.0371 |
| C6orf108 | 759.544 | -1.030 | 0.333 | -3.096 | 0.00196 | 0.0385 |
| PSME1 | 12240.674 | -1.029 | 0.317 | -3.249 | 0.00116 | 0.0298 |
| CFL1 | 28377.522 | -1.029 | 0.319 | -3.222 | 0.00127 | 0.0309 |
| CLIC1 | 28593.372 | -1.028 | 0.329 | -3.129 | 0.00176 | 0.0366 |
| SRI | 5763.373 | -1.028 | 0.343 | -2.997 | 0.00273 | 0.0447 |
| PARK7 | 6227.408 | -1.026 | 0.323 | -3.174 | 0.00151 | 0.0341 |
| AKIP1 | 1135.859 | -1.026 | 0.361 | -2.844 | 0.00446 | 0.0575 |
| SAE1 | 5904.253 | -1.025 | 0.306 | -3.350 | 0.00081 | 0.0260 |
| VDAC2 | 8100.095 | -1.024 | 0.368 | -2.782 | 0.00540 | 0.0638 |
| RP9 | 943.841 | -1.024 | 0.325 | -3.147 | 0.00165 | 0.0357 |
| FKBP4 | 3156.538 | -1.020 | 0.262 | -3.892 | 0.00010 | 0.0131 |
| CHMP2A | 4407.299 | -1.017 | 0.309 | -3.288 | 0.00101 | 0.0280 |
| UBXN1 | 8415.693 | -1.016 | 0.331 | -3.069 | 0.00215 | 0.0400 |
| SSSCA1 | 491.730 | -1.016 | 0.302 | -3.363 | 0.00077 | 0.0254 |
| IAH1 | 3795.364 | -1.015 | 0.362 | -2.804 | 0.00505 | 0.0612 |
| PRDX6 | 7981.515 | -1.013 | 0.346 | -2.931 | 0.00338 | 0.0501 |
| ATP6V0B | 3275.138 | -1.010 | 0.273 | -3.699 | 0.00022 | 0.0167 |
| ACP1 | 7836.346 | -1.010 | 0.369 | -2.733 | 0.00627 | 0.0687 |
| TTC7B | 147.619 | -1.009 | 0.294 | -3.430 | 0.00060 | 0.0232 |
| RNF19B | 2785.432 | -1.006 | 0.351 | -2.865 | 0.00417 | 0.0558 |
| CLPP | 3115.133 | -1.006 | 0.314 | -3.205 | 0.00135 | 0.0322 |
| RPL19 | 71872.056 | -1.006 | 0.334 | -3.009 | 0.00262 | 0.0437 |
| SF3B5 | 3533.651 | -1.006 | 0.360 | -2.791 | 0.00526 | 0.0631 |
| IFT52 | 1309.029 | -1.005 | 0.389 | -2.582 | 0.00984 | 0.0884 |
| CHTF18 | 1660.355 | -1.005 | 0.343 | -2.930 | 0.00339 | 0.0501 |
| POLR2H | 3185.616 | -1.005 | 0.328 | -3.060 | 0.00221 | 0.0406 |
| RPL3 | 146108.587 | -1.004 | 0.310 | -3.237 | 0.00121 | 0.0304 |
| RPL13A | 152421.144 | -1.004 | 0.349 | -2.875 | 0.00404 | 0.0548 |
| RPS27A | 53936.194 | -1.003 | 0.367 | -2.729 | 0.00635 | 0.0692 |
| RAD54L | 1680.282 | -1.003 | 0.354 | -2.835 | 0.00458 | 0.0582 |
| RRM1 | 8159.218 | -1.002 | 0.373 | -2.685 | 0.00726 | 0.0742 |
| MRPS33 | 1410.832 | -1.002 | 0.372 | -2.691 | 0.00712 | 0.0735 |
| C1orf122 | 727.873 | -1.000 | 0.337 | -2.970 | 0.00298 | 0.0468 |
| BUD31 | 2565.211 | -1.000 | 0.325 | -3.073 | 0.00212 | 0.0399 |
| SRGN | 25021.186 | -0.999 | 0.356 | -2.806 | 0.00501 | 0.0608 |
| DLGAP5 | 2503.262 | -0.999 | 0.367 | -2.724 | 0.00645 | 0.0700 |
| ZNHIT3 | 2729.307 | -0.999 | 0.338 | -2.954 | 0.00314 | 0.0480 |
| CCDC124 | 1650.647 | -0.999 | 0.275 | -3.626 | 0.00029 | 0.0180 |
| COMMD1 | 693.056 | -0.998 | 0.360 | -2.770 | 0.00561 | 0.0652 |
| TSPO | 4625.016 | -0.998 | 0.283 | -3.523 | 0.00043 | 0.0203 |
| PIM1 | 11760.751 | -0.998 | 0.351 | -2.841 | 0.00450 | 0.0577 |
| CCT8 | 16593.674 | -0.997 | 0.344 | -2.901 | 0.00372 | 0.0524 |
| TMED3 | 3836.472 | -0.997 | 0.307 | -3.254 | 0.00114 | 0.0294 |
| MRPL13 | 2061.276 | -0.997 | 0.366 | -2.725 | 0.00642 | 0.0698 |
| TACO1 | 1026.222 | -0.996 | 0.345 | -2.884 | 0.00393 | 0.0540 |
| MRPL33 | 2045.141 | -0.995 | 0.332 | -3.002 | 0.00268 | 0.0442 |
| RANGAP1 | 4262.986 | -0.994 | 0.264 | -3.767 | 0.00017 | 0.0146 |
| PIN4 | 916.181 | -0.994 | 0.355 | -2.799 | 0.00513 | 0.0619 |
| GPR160 | 1420.782 | -0.994 | 0.292 | -3.399 | 0.00068 | 0.0245 |
| ZCRB1 | 3635.601 | -0.993 | 0.385 | -2.578 | 0.00995 | 0.0890 |
| PRKCE | 765.257 | -0.993 | 0.319 | -3.113 | 0.00185 | 0.0375 |
| BZW1 | 13198.488 | -0.993 | 0.358 | -2.776 | 0.00550 | 0.0644 |
| SSRP1 | 11132.687 | -0.992 | 0.359 | -2.765 | 0.00569 | 0.0652 |
| TOMM40 | 1372.499 | -0.991 | 0.330 | -3.008 | 0.00263 | 0.0437 |
| MTCH2 | 2544.718 | -0.991 | 0.338 | -2.931 | 0.00338 | 0.0501 |
| STAM | 3695.232 | -0.990 | 0.332 | -2.982 | 0.00286 | 0.0458 |
| RPL23A | 48367.219 | -0.990 | 0.360 | -2.748 | 0.00599 | 0.0672 |
| C20orf20 | 2927.453 | -0.990 | 0.379 | -2.610 | 0.00906 | 0.0845 |
| TXNL4A | 3626.829 | -0.989 | 0.368 | -2.686 | 0.00723 | 0.0741 |
| RPS25 | 46264.830 | -0.989 | 0.360 | -2.749 | 0.00599 | 0.0672 |
| METTL21A | 724.814 | -0.988 | 0.375 | -2.635 | 0.00842 | 0.0809 |
| RFC2 | 1431.715 | -0.987 | 0.360 | -2.742 | 0.00610 | 0.0679 |
| MLST8 | 1878.749 | -0.987 | 0.309 | -3.188 | 0.00143 | 0.0331 |
| RPS14 | 47254.124 | -0.986 | 0.330 | -2.985 | 0.00284 | 0.0458 |
| MRPS30 | 2394.235 | -0.986 | 0.319 | -3.088 | 0.00202 | 0.0394 |
| ST13 | 16824.764 | -0.985 | 0.372 | -2.651 | 0.00802 | 0.0783 |
| FAF1 | 2498.260 | -0.985 | 0.302 | -3.258 | 0.00112 | 0.0292 |
| CDC25A | 416.875 | -0.984 | 0.376 | -2.620 | 0.00879 | 0.0832 |
| CCDC86 | 983.452 | -0.984 | 0.278 | -3.540 | 0.00040 | 0.0199 |
| PPIA | 42363.849 | -0.983 | 0.294 | -3.340 | 0.00084 | 0.0260 |
| SDHB | 3460.109 | -0.983 | 0.333 | -2.954 | 0.00314 | 0.0480 |
| CISD1 | 1578.015 | -0.982 | 0.345 | -2.852 | 0.00435 | 0.0567 |
| CCT3 | 8563.864 | -0.982 | 0.338 | -2.905 | 0.00367 | 0.0521 |
| HN1 | 5952.149 | -0.981 | 0.296 | -3.312 | 0.00093 | 0.0270 |
| RHBDD2 | 3336.136 | -0.981 | 0.321 | -3.058 | 0.00223 | 0.0407 |
| CISH | 3919.715 | -0.980 | 0.315 | -3.108 | 0.00188 | 0.0378 |
| ACAT1 | 3210.289 | -0.979 | 0.331 | -2.962 | 0.00305 | 0.0475 |
| CDKN2AIPNL | 1061.465 | -0.978 | 0.329 | -2.970 | 0.00297 | 0.0468 |
| ATPIF1 | 3764.526 | -0.977 | 0.287 | -3.406 | 0.00066 | 0.0243 |
| ATP5O | 1895.694 | -0.977 | 0.277 | -3.530 | 0.00041 | 0.0203 |
| SGMS1 | 2404.162 | -0.977 | 0.246 | -3.968 | 0.00007 | 0.0116 |
| C12orf5 | 1336.895 | -0.976 | 0.325 | -2.999 | 0.00271 | 0.0446 |
| FAM26F | 453.830 | -0.975 | 0.378 | -2.581 | 0.00985 | 0.0885 |
| TRAF4 | 1780.336 | -0.975 | 0.319 | -3.053 | 0.00227 | 0.0411 |
| PQBP1 | 3022.817 | -0.975 | 0.338 | -2.885 | 0.00391 | 0.0539 |
| RBBP7 | 8181.881 | -0.974 | 0.329 | -2.959 | 0.00308 | 0.0477 |
| ZNRD1 | 1628.262 | -0.973 | 0.339 | -2.869 | 0.00411 | 0.0554 |
| ZNF524 | 1434.040 | -0.973 | 0.327 | -2.979 | 0.00289 | 0.0461 |
| NDUFB6 | 1589.561 | -0.973 | 0.339 | -2.873 | 0.00407 | 0.0549 |
| FIBP | 2745.018 | -0.973 | 0.294 | -3.303 | 0.00096 | 0.0273 |
| YWHAH | 6206.619 | -0.972 | 0.339 | -2.864 | 0.00418 | 0.0558 |
| SSNA1 | 2276.374 | -0.972 | 0.330 | -2.949 | 0.00319 | 0.0483 |
| EIF3I | 8179.984 | -0.972 | 0.298 | -3.265 | 0.00110 | 0.0290 |
| APOO | 301.680 | -0.972 | 0.335 | -2.899 | 0.00374 | 0.0524 |
| TSTA3 | 1187.385 | -0.971 | 0.341 | -2.850 | 0.00437 | 0.0568 |
| PUSL1 | 250.638 | -0.970 | 0.366 | -2.651 | 0.00802 | 0.0783 |
| ASF1B | 2204.812 | -0.970 | 0.329 | -2.947 | 0.00321 | 0.0485 |
| FARS2 | 1592.157 | -0.969 | 0.342 | -2.833 | 0.00462 | 0.0585 |
| AEN | 2631.812 | -0.968 | 0.315 | -3.067 | 0.00216 | 0.0401 |
| SENP3-EIF4A1 | 29817.714 | -0.966 | 0.297 | -3.255 | 0.00113 | 0.0292 |
| ADRM1 | 4134.307 | -0.964 | 0.292 | -3.299 | 0.00097 | 0.0275 |
| RDBP | 2082.284 | -0.963 | 0.313 | -3.080 | 0.00207 | 0.0396 |
| GLRX3 | 4416.199 | -0.962 | 0.299 | -3.213 | 0.00131 | 0.0315 |
| CLNS1A | 4733.472 | -0.962 | 0.336 | -2.862 | 0.00421 | 0.0560 |
| PHYH | 932.830 | -0.959 | 0.323 | -2.972 | 0.00296 | 0.0467 |
| NAPA | 1159.646 | -0.959 | 0.287 | -3.339 | 0.00084 | 0.0260 |
| RHOA | 35383.823 | -0.959 | 0.341 | -2.809 | 0.00498 | 0.0607 |
| TUBA1C | 4588.259 | -0.958 | 0.255 | -3.762 | 0.00017 | 0.0148 |
| IMPA2 | 350.315 | -0.957 | 0.368 | -2.599 | 0.00936 | 0.0861 |
| AIP | 6172.086 | -0.956 | 0.280 | -3.419 | 0.00063 | 0.0236 |
| PSMC5 | 6046.710 | -0.955 | 0.303 | -3.149 | 0.00164 | 0.0356 |
| SELS | 2198.257 | -0.955 | 0.349 | -2.736 | 0.00621 | 0.0686 |
| ANXA7 | 6046.153 | -0.954 | 0.339 | -2.811 | 0.00493 | 0.0606 |
| CCT4 | 8447.513 | -0.953 | 0.296 | -3.225 | 0.00126 | 0.0307 |
| MRPL37 | 4931.471 | -0.953 | 0.320 | -2.978 | 0.00290 | 0.0462 |
| PCBD1 | 1307.379 | -0.952 | 0.299 | -3.181 | 0.00147 | 0.0333 |
| CCNH | 2168.283 | -0.952 | 0.314 | -3.032 | 0.00243 | 0.0428 |
| NCAPH2 | 2825.632 | -0.952 | 0.211 | -4.520 | 0.00001 | 0.0052 |
| EPS8L2 | 1549.352 | -0.952 | 0.246 | -3.872 | 0.00011 | 0.0132 |
| CENPN | 1737.159 | -0.949 | 0.357 | -2.657 | 0.00789 | 0.0778 |
| CBX8 | 304.456 | -0.949 | 0.294 | -3.222 | 0.00127 | 0.0309 |
| EIF4A3 | 2974.983 | -0.949 | 0.260 | -3.645 | 0.00027 | 0.0177 |
| PGCP | 613.529 | -0.949 | 0.326 | -2.909 | 0.00363 | 0.0520 |
| BAG4 | 1641.428 | -0.948 | 0.340 | -2.787 | 0.00531 | 0.0636 |
| METTL5 | 1819.928 | -0.948 | 0.318 | -2.983 | 0.00285 | 0.0458 |
| RAD51C | 1650.533 | -0.948 | 0.338 | -2.804 | 0.00504 | 0.0611 |
| MRPL28 | 3245.092 | -0.947 | 0.344 | -2.755 | 0.00587 | 0.0665 |
| GOT2 | 3961.177 | -0.946 | 0.325 | -2.907 | 0.00365 | 0.0520 |
| PSMD8 | 6024.446 | -0.945 | 0.297 | -3.185 | 0.00145 | 0.0331 |
| NASP | 5913.571 | -0.945 | 0.261 | -3.624 | 0.00029 | 0.0181 |
| LDHB | 52898.167 | -0.944 | 0.338 | -2.794 | 0.00521 | 0.0627 |
| RPL22 | 23019.352 | -0.943 | 0.361 | -2.615 | 0.00892 | 0.0839 |
| TSSC4 | 1161.319 | -0.943 | 0.327 | -2.886 | 0.00390 | 0.0538 |
| HMGN2 | 14353.882 | -0.943 | 0.305 | -3.091 | 0.00200 | 0.0391 |
| RPL30 | 43733.055 | -0.940 | 0.351 | -2.682 | 0.00732 | 0.0745 |
| HSD17B8 | 449.587 | -0.938 | 0.311 | -3.015 | 0.00257 | 0.0434 |
| RPS4X | 85996.012 | -0.938 | 0.322 | -2.916 | 0.00354 | 0.0515 |
| DIAPH2 | 1240.478 | -0.938 | 0.326 | -2.876 | 0.00403 | 0.0547 |
| CCT7 | 8640.240 | -0.937 | 0.306 | -3.061 | 0.00220 | 0.0406 |
| GCHFR | 1536.872 | -0.936 | 0.303 | -3.086 | 0.00203 | 0.0394 |
| ELOF1 | 2216.871 | -0.934 | 0.299 | -3.125 | 0.00178 | 0.0368 |
| TRAPPC1 | 6531.588 | -0.934 | 0.310 | -3.016 | 0.00256 | 0.0434 |
| UNC119 | 3982.716 | -0.934 | 0.351 | -2.663 | 0.00774 | 0.0768 |
| SNRNP25 | 1033.810 | -0.934 | 0.332 | -2.810 | 0.00496 | 0.0607 |
| PEMT | 592.275 | -0.932 | 0.340 | -2.744 | 0.00607 | 0.0678 |
| NDUFB11 | 4090.576 | -0.932 | 0.340 | -2.743 | 0.00609 | 0.0679 |
| C5orf4 | 195.505 | -0.931 | 0.344 | -2.709 | 0.00676 | 0.0712 |
| MYO6 | 844.641 | -0.931 | 0.326 | -2.860 | 0.00424 | 0.0562 |
| RPS11 | 90253.844 | -0.931 | 0.360 | -2.582 | 0.00981 | 0.0884 |
| SCAND1 | 2221.379 | -0.929 | 0.309 | -3.010 | 0.00261 | 0.0436 |
| ALKBH7 | 1570.290 | -0.929 | 0.351 | -2.645 | 0.00816 | 0.0793 |
| SUMO1 | 6107.266 | -0.929 | 0.333 | -2.793 | 0.00522 | 0.0627 |
| FDFT1 | 14936.031 | -0.928 | 0.343 | -2.709 | 0.00675 | 0.0712 |
| PSME2 | 4720.649 | -0.928 | 0.284 | -3.274 | 0.00106 | 0.0287 |
| MDH2 | 9492.146 | -0.927 | 0.281 | -3.301 | 0.00096 | 0.0275 |
| C1orf35 | 955.701 | -0.927 | 0.275 | -3.370 | 0.00075 | 0.0252 |
| ARHGAP15 | 10875.694 | -0.927 | 0.329 | -2.815 | 0.00488 | 0.0602 |
| TIMM9 | 1586.567 | -0.925 | 0.320 | -2.895 | 0.00380 | 0.0529 |
| MRP63 | 1433.334 | -0.925 | 0.310 | -2.981 | 0.00288 | 0.0460 |
| C15orf29 | 2059.255 | -0.924 | 0.339 | -2.726 | 0.00640 | 0.0696 |
| PDRG1 | 917.679 | -0.924 | 0.276 | -3.346 | 0.00082 | 0.0260 |
| PLEKHJ1 | 1904.766 | -0.923 | 0.323 | -2.859 | 0.00426 | 0.0562 |
| NDUFB5 | 4492.474 | -0.923 | 0.346 | -2.668 | 0.00764 | 0.0761 |
| NDUFV2 | 11537.044 | -0.923 | 0.324 | -2.848 | 0.00440 | 0.0570 |
| KARS | 7455.925 | -0.921 | 0.311 | -2.966 | 0.00302 | 0.0471 |
| NDUFB7 | 1787.975 | -0.920 | 0.293 | -3.141 | 0.00168 | 0.0358 |
| PRMT1 | 4334.557 | -0.920 | 0.255 | -3.616 | 0.00030 | 0.0181 |
| HIST3H2A | 387.443 | -0.920 | 0.316 | -2.908 | 0.00364 | 0.0520 |
| PTGES3 | 18154.597 | -0.920 | 0.347 | -2.653 | 0.00798 | 0.0781 |
| ELMO1 | 9489.762 | -0.920 | 0.324 | -2.838 | 0.00454 | 0.0579 |
| CIB1 | 7320.205 | -0.919 | 0.253 | -3.640 | 0.00027 | 0.0177 |
| CAPZA2 | 5163.758 | -0.918 | 0.356 | -2.583 | 0.00980 | 0.0884 |
| MRPL55 | 1286.286 | -0.918 | 0.298 | -3.082 | 0.00206 | 0.0396 |
| DYNLRB1 | 3770.862 | -0.918 | 0.314 | -2.921 | 0.00349 | 0.0510 |
| ATP5F1 | 8003.555 | -0.917 | 0.331 | -2.769 | 0.00562 | 0.0652 |
| CTPS | 791.643 | -0.916 | 0.349 | -2.624 | 0.00868 | 0.0827 |
| PMM1 | 2968.311 | -0.913 | 0.279 | -3.272 | 0.00107 | 0.0287 |
| FAM50A | 3748.510 | -0.913 | 0.314 | -2.905 | 0.00367 | 0.0521 |
| SLC25A11 | 2486.475 | -0.913 | 0.278 | -3.290 | 0.00100 | 0.0279 |
| TNFAIP8 | 7527.358 | -0.913 | 0.333 | -2.745 | 0.00605 | 0.0676 |
| SLC27A4 | 1691.573 | -0.912 | 0.333 | -2.740 | 0.00615 | 0.0683 |
| H2AFJ | 1769.683 | -0.912 | 0.274 | -3.321 | 0.00090 | 0.0266 |
| NCAPG | 3705.331 | -0.910 | 0.338 | -2.693 | 0.00708 | 0.0732 |
| RRAS | 718.432 | -0.910 | 0.296 | -3.076 | 0.00210 | 0.0397 |
| USE1 | 1357.282 | -0.910 | 0.352 | -2.586 | 0.00970 | 0.0880 |
| SYF2 | 5875.474 | -0.909 | 0.348 | -2.614 | 0.00895 | 0.0840 |
| PDCD5 | 4095.336 | -0.908 | 0.325 | -2.797 | 0.00516 | 0.0623 |
| SUGT1 | 3643.447 | -0.905 | 0.294 | -3.082 | 0.00206 | 0.0396 |
| GNG5 | 4088.570 | -0.905 | 0.299 | -3.023 | 0.00250 | 0.0431 |
| NDUFS3 | 2479.291 | -0.905 | 0.301 | -3.006 | 0.00265 | 0.0439 |
| TMEM208 | 977.892 | -0.904 | 0.333 | -2.717 | 0.00658 | 0.0706 |
| ATP5G2 | 14022.904 | -0.904 | 0.314 | -2.882 | 0.00396 | 0.0541 |
| ARPC3 | 18663.067 | -0.903 | 0.295 | -3.056 | 0.00224 | 0.0407 |
| PSMB4 | 9917.327 | -0.903 | 0.275 | -3.278 | 0.00105 | 0.0286 |
| C12orf10 | 1846.810 | -0.900 | 0.287 | -3.141 | 0.00168 | 0.0358 |
| PSTPIP2 | 1935.043 | -0.900 | 0.320 | -2.813 | 0.00491 | 0.0605 |
| PA2G4 | 7588.737 | -0.899 | 0.334 | -2.694 | 0.00706 | 0.0730 |
| ANP32B | 25075.767 | -0.899 | 0.331 | -2.716 | 0.00661 | 0.0707 |
| PSMB10 | 6948.813 | -0.897 | 0.317 | -2.830 | 0.00465 | 0.0587 |
| SNHG12 | 1874.866 | -0.897 | 0.234 | -3.826 | 0.00013 | 0.0132 |
| YIF1A | 923.064 | -0.897 | 0.269 | -3.333 | 0.00086 | 0.0262 |
| THYN1 | 3628.006 | -0.896 | 0.279 | -3.213 | 0.00131 | 0.0315 |
| ST7 | 459.605 | -0.896 | 0.287 | -3.120 | 0.00181 | 0.0370 |
| COMMD7 | 2970.306 | -0.895 | 0.302 | -2.964 | 0.00304 | 0.0474 |
| MRPS34 | 2586.848 | -0.894 | 0.314 | -2.852 | 0.00434 | 0.0567 |
| CCNE1 | 632.527 | -0.893 | 0.273 | -3.276 | 0.00105 | 0.0286 |
| C11orf10 | 5123.239 | -0.892 | 0.316 | -2.826 | 0.00472 | 0.0591 |
| DUSP12 | 1285.762 | -0.892 | 0.341 | -2.618 | 0.00884 | 0.0835 |
| CCT5 | 5839.731 | -0.891 | 0.296 | -3.014 | 0.00258 | 0.0434 |
| UBTD1 | 225.811 | -0.889 | 0.318 | -2.801 | 0.00510 | 0.0617 |
| EAF2 | 772.196 | -0.889 | 0.305 | -2.911 | 0.00360 | 0.0519 |
| UBB | 18665.557 | -0.889 | 0.249 | -3.575 | 0.00035 | 0.0191 |
| MIEN1 | 1165.158 | -0.888 | 0.313 | -2.837 | 0.00455 | 0.0580 |
| AKR1A1 | 3666.136 | -0.886 | 0.289 | -3.070 | 0.00214 | 0.0399 |
| EEF1G | 110835.960 | -0.886 | 0.298 | -2.973 | 0.00295 | 0.0466 |
| COTL1 | 60620.963 | -0.885 | 0.312 | -2.840 | 0.00452 | 0.0577 |
| MRPL3 | 3221.395 | -0.885 | 0.343 | -2.578 | 0.00993 | 0.0889 |
| COPE | 5055.909 | -0.885 | 0.266 | -3.320 | 0.00090 | 0.0266 |
| HCFC1R1 | 2694.693 | -0.883 | 0.319 | -2.766 | 0.00567 | 0.0652 |
| THOC3 | 1019.321 | -0.883 | 0.301 | -2.932 | 0.00337 | 0.0500 |
| VPS28 | 7016.953 | -0.883 | 0.300 | -2.944 | 0.00324 | 0.0487 |
| EIF3K | 10087.798 | -0.883 | 0.322 | -2.742 | 0.00610 | 0.0679 |
| EIF3G | 8367.433 | -0.882 | 0.302 | -2.918 | 0.00353 | 0.0514 |
| MRPL22 | 1565.237 | -0.881 | 0.310 | -2.841 | 0.00450 | 0.0577 |
| PHB | 2017.077 | -0.879 | 0.317 | -2.778 | 0.00547 | 0.0642 |
| RPL4 | 123761.836 | -0.879 | 0.332 | -2.648 | 0.00810 | 0.0790 |
| C16orf13 | 2257.355 | -0.879 | 0.323 | -2.718 | 0.00656 | 0.0704 |
| ELAVL1 | 5756.192 | -0.878 | 0.323 | -2.721 | 0.00652 | 0.0703 |
| OST4 | 6050.363 | -0.877 | 0.308 | -2.851 | 0.00436 | 0.0567 |
| PSMD4 | 7253.253 | -0.875 | 0.284 | -3.082 | 0.00206 | 0.0396 |
| PSMB5 | 1888.537 | -0.875 | 0.325 | -2.693 | 0.00709 | 0.0732 |
| HAUS4 | 3994.213 | -0.873 | 0.333 | -2.619 | 0.00882 | 0.0834 |
| DENND5A | 2317.458 | -0.873 | 0.330 | -2.645 | 0.00816 | 0.0793 |
| AIFM1 | 2428.096 | -0.871 | 0.264 | -3.296 | 0.00098 | 0.0277 |
| WDR54 | 6342.073 | -0.869 | 0.300 | -2.898 | 0.00375 | 0.0525 |
| MAGOH | 1616.963 | -0.869 | 0.322 | -2.702 | 0.00689 | 0.0720 |
| BRK1 | 9632.601 | -0.868 | 0.279 | -3.113 | 0.00185 | 0.0375 |
| SEC11C | 8262.270 | -0.868 | 0.323 | -2.690 | 0.00715 | 0.0737 |
| GTSF1 | 352.971 | -0.867 | 0.299 | -2.899 | 0.00375 | 0.0524 |
| MRTO4 | 868.211 | -0.866 | 0.318 | -2.720 | 0.00653 | 0.0703 |
| ACBD6 | 2118.789 | -0.866 | 0.296 | -2.931 | 0.00338 | 0.0501 |
| NRGN | 203.315 | -0.866 | 0.318 | -2.721 | 0.00651 | 0.0703 |
| GABARAPL2 | 5799.309 | -0.865 | 0.329 | -2.630 | 0.00853 | 0.0815 |
| BAD | 1819.991 | -0.864 | 0.288 | -2.996 | 0.00274 | 0.0448 |
| POLR2E | 5607.391 | -0.863 | 0.292 | -2.951 | 0.00317 | 0.0482 |
| TXNL1 | 6259.228 | -0.861 | 0.305 | -2.826 | 0.00471 | 0.0591 |
| GMFG | 20197.177 | -0.861 | 0.330 | -2.613 | 0.00898 | 0.0842 |
| MNAT1 | 2070.429 | -0.860 | 0.322 | -2.666 | 0.00767 | 0.0763 |
| HNRPDL | 18158.722 | -0.860 | 0.308 | -2.789 | 0.00528 | 0.0634 |
| HIGD2A | 4022.191 | -0.858 | 0.321 | -2.672 | 0.00754 | 0.0758 |
| C22orf28 | 3065.726 | -0.858 | 0.286 | -2.996 | 0.00273 | 0.0448 |
| UBE2M | 3243.535 | -0.858 | 0.254 | -3.384 | 0.00071 | 0.0246 |
| NME3 | 1492.724 | -0.858 | 0.321 | -2.676 | 0.00744 | 0.0752 |
| DCTN3 | 2604.152 | -0.856 | 0.305 | -2.809 | 0.00498 | 0.0607 |
| ETFA | 3687.698 | -0.854 | 0.323 | -2.647 | 0.00812 | 0.0792 |
| DAP | 5719.539 | -0.854 | 0.318 | -2.685 | 0.00726 | 0.0742 |
| PSMG2 | 4949.654 | -0.853 | 0.286 | -2.980 | 0.00288 | 0.0460 |
| EEF1A1 | 546038.514 | -0.852 | 0.319 | -2.669 | 0.00761 | 0.0761 |
| MRPL21 | 1380.969 | -0.851 | 0.305 | -2.790 | 0.00527 | 0.0633 |
| FLJ43663 | 3750.895 | -0.850 | 0.324 | -2.624 | 0.00870 | 0.0827 |
| NDUFA8 | 1470.849 | -0.848 | 0.316 | -2.686 | 0.00723 | 0.0741 |
| GMPS | 5148.948 | -0.846 | 0.309 | -2.737 | 0.00620 | 0.0686 |
| SNRPA | 5485.469 | -0.846 | 0.287 | -2.949 | 0.00318 | 0.0482 |
| TAGLN2 | 38886.521 | -0.846 | 0.287 | -2.946 | 0.00322 | 0.0486 |
| RACGAP1 | 4800.937 | -0.845 | 0.309 | -2.740 | 0.00614 | 0.0682 |
| FARSB | 1857.815 | -0.845 | 0.272 | -3.110 | 0.00187 | 0.0376 |
| GPS1 | 2320.518 | -0.844 | 0.311 | -2.715 | 0.00664 | 0.0708 |
| C9orf123 | 1234.128 | -0.844 | 0.322 | -2.618 | 0.00883 | 0.0835 |
| RPL6 | 34693.194 | -0.842 | 0.325 | -2.593 | 0.00951 | 0.0868 |
| PCCB | 1840.005 | -0.842 | 0.247 | -3.405 | 0.00066 | 0.0243 |
| HAUS1 | 1914.035 | -0.841 | 0.317 | -2.657 | 0.00788 | 0.0778 |
| NSA2 | 6156.051 | -0.841 | 0.307 | -2.738 | 0.00618 | 0.0685 |
| CDC34 | 2367.144 | -0.841 | 0.235 | -3.578 | 0.00035 | 0.0191 |
| G6PC3 | 653.317 | -0.840 | 0.202 | -4.161 | 0.00003 | 0.0085 |
| ATP6V1H | 2606.882 | -0.837 | 0.287 | -2.920 | 0.00350 | 0.0511 |
| PSRC1 | 410.531 | -0.836 | 0.289 | -2.889 | 0.00386 | 0.0535 |
| TUFT1 | 318.323 | -0.836 | 0.298 | -2.809 | 0.00497 | 0.0607 |
| GNB2L1 | 95620.795 | -0.835 | 0.306 | -2.733 | 0.00628 | 0.0687 |
| SIVA1 | 1079.548 | -0.833 | 0.306 | -2.722 | 0.00649 | 0.0703 |
| RNF113A | 2703.231 | -0.830 | 0.242 | -3.437 | 0.00059 | 0.0230 |
| PSMC6 | 5389.874 | -0.830 | 0.276 | -3.013 | 0.00259 | 0.0435 |
| TSFM | 912.644 | -0.829 | 0.294 | -2.823 | 0.00476 | 0.0593 |
| UQCRC2 | 13588.924 | -0.828 | 0.306 | -2.708 | 0.00678 | 0.0714 |
| THOP1 | 1923.244 | -0.827 | 0.270 | -3.056 | 0.00224 | 0.0407 |
| APRT | 3488.041 | -0.826 | 0.306 | -2.702 | 0.00689 | 0.0720 |
| SEC11A | 9031.276 | -0.826 | 0.316 | -2.611 | 0.00903 | 0.0843 |
| DNAJA1 | 5130.362 | -0.824 | 0.259 | -3.179 | 0.00148 | 0.0335 |
| IDH2 | 8972.567 | -0.823 | 0.234 | -3.518 | 0.00044 | 0.0204 |
| CHAF1A | 2476.972 | -0.822 | 0.261 | -3.154 | 0.00161 | 0.0355 |
| GLO1 | 10548.276 | -0.821 | 0.318 | -2.585 | 0.00975 | 0.0882 |
| HEXB | 3394.042 | -0.820 | 0.306 | -2.678 | 0.00741 | 0.0748 |
| PAK1 | 1752.799 | -0.819 | 0.309 | -2.652 | 0.00800 | 0.0783 |
| HAX1 | 4475.441 | -0.818 | 0.258 | -3.167 | 0.00154 | 0.0346 |
| PHB2 | 10767.867 | -0.818 | 0.292 | -2.804 | 0.00504 | 0.0611 |
| RPP30 | 2641.869 | -0.818 | 0.291 | -2.809 | 0.00496 | 0.0607 |
| CUEDC2 | 3222.986 | -0.817 | 0.297 | -2.755 | 0.00588 | 0.0665 |
| MRPL18 | 1756.928 | -0.817 | 0.256 | -3.188 | 0.00143 | 0.0331 |
| PITHD1 | 2414.496 | -0.815 | 0.287 | -2.840 | 0.00451 | 0.0577 |
| ALCAM | 2371.619 | -0.815 | 0.312 | -2.611 | 0.00902 | 0.0843 |
| CNKSR2 | 175.590 | -0.813 | 0.314 | -2.586 | 0.00970 | 0.0880 |
| C4orf46 | 1168.634 | -0.812 | 0.297 | -2.735 | 0.00624 | 0.0686 |
| STUB1 | 1107.161 | -0.811 | 0.272 | -2.983 | 0.00285 | 0.0458 |
| ING1 | 1975.616 | -0.810 | 0.293 | -2.768 | 0.00564 | 0.0652 |
| THAP3 | 1464.206 | -0.809 | 0.266 | -3.035 | 0.00240 | 0.0425 |
| RAD23A | 5968.367 | -0.807 | 0.291 | -2.775 | 0.00552 | 0.0646 |
| UGP2 | 7118.457 | -0.806 | 0.281 | -2.867 | 0.00414 | 0.0556 |
| NDUFS7 | 1655.562 | -0.805 | 0.226 | -3.563 | 0.00037 | 0.0193 |
| MRPL32 | 2206.988 | -0.805 | 0.289 | -2.785 | 0.00534 | 0.0637 |
| IMP4 | 1319.900 | -0.805 | 0.249 | -3.234 | 0.00122 | 0.0305 |
| NDUFA9 | 2249.659 | -0.804 | 0.249 | -3.231 | 0.00123 | 0.0305 |
| MRPL4 | 1697.450 | -0.803 | 0.261 | -3.078 | 0.00208 | 0.0396 |
| ZNF367 | 1616.282 | -0.803 | 0.258 | -3.111 | 0.00186 | 0.0375 |
| GUK1 | 12884.623 | -0.802 | 0.241 | -3.330 | 0.00087 | 0.0264 |
| MRPS18B | 3196.141 | -0.801 | 0.285 | -2.808 | 0.00498 | 0.0607 |
| ANO10 | 852.747 | -0.800 | 0.293 | -2.732 | 0.00630 | 0.0689 |
| TMEM11 | 791.919 | -0.799 | 0.287 | -2.788 | 0.00531 | 0.0636 |
| PRDX4 | 1657.521 | -0.799 | 0.295 | -2.713 | 0.00667 | 0.0708 |
| NFKBIE | 1010.630 | -0.799 | 0.257 | -3.111 | 0.00186 | 0.0375 |
| RSRC1 | 1166.359 | -0.798 | 0.305 | -2.614 | 0.00894 | 0.0839 |
| ADA | 4325.856 | -0.794 | 0.263 | -3.024 | 0.00250 | 0.0431 |
| COPS4 | 2523.445 | -0.794 | 0.266 | -2.987 | 0.00282 | 0.0458 |
| SSBP1 | 4327.375 | -0.794 | 0.278 | -2.854 | 0.00431 | 0.0565 |
| RHOG | 7639.828 | -0.792 | 0.263 | -3.011 | 0.00260 | 0.0436 |
| TMEM101 | 1986.333 | -0.791 | 0.256 | -3.094 | 0.00197 | 0.0387 |
| DNAJC1 | 2630.620 | -0.791 | 0.283 | -2.796 | 0.00517 | 0.0623 |
| ETHE1 | 1928.287 | -0.791 | 0.288 | -2.748 | 0.00599 | 0.0672 |
| GAPDH | 529052.768 | -0.791 | 0.281 | -2.817 | 0.00485 | 0.0599 |
| SSR4 | 6633.764 | -0.790 | 0.303 | -2.609 | 0.00907 | 0.0845 |
| VIM | 148775.223 | -0.789 | 0.304 | -2.597 | 0.00940 | 0.0862 |
| MRPL9 | 2867.988 | -0.788 | 0.279 | -2.824 | 0.00474 | 0.0592 |
| PFN1 | 59636.531 | -0.788 | 0.292 | -2.701 | 0.00690 | 0.0720 |
| COX4NB | 1029.225 | -0.786 | 0.258 | -3.047 | 0.00231 | 0.0417 |
| GTF3A | 11254.416 | -0.785 | 0.304 | -2.584 | 0.00978 | 0.0884 |
| CSNK2B | 9073.064 | -0.782 | 0.288 | -2.711 | 0.00671 | 0.0710 |
| KIFC1 | 4212.525 | -0.781 | 0.293 | -2.669 | 0.00761 | 0.0761 |
| RIT1 | 2452.818 | -0.781 | 0.285 | -2.739 | 0.00617 | 0.0685 |
| MRPS18C | 1212.685 | -0.779 | 0.233 | -3.339 | 0.00084 | 0.0260 |
| DNAJB1 | 4926.292 | -0.777 | 0.272 | -2.853 | 0.00434 | 0.0567 |
| ARF5 | 4950.003 | -0.774 | 0.256 | -3.026 | 0.00247 | 0.0429 |
| RTN4IP1 | 421.091 | -0.774 | 0.300 | -2.580 | 0.00989 | 0.0887 |
| FAM89B | 2160.792 | -0.773 | 0.271 | -2.850 | 0.00437 | 0.0568 |
| THAP4 | 2738.611 | -0.773 | 0.257 | -3.008 | 0.00263 | 0.0437 |
| PCGF1 | 981.512 | -0.772 | 0.255 | -3.022 | 0.00251 | 0.0431 |
| MED27 | 1080.046 | -0.772 | 0.207 | -3.726 | 0.00019 | 0.0159 |
| FNDC3B | 1221.945 | -0.770 | 0.247 | -3.114 | 0.00185 | 0.0375 |
| PSMB1 | 8427.094 | -0.769 | 0.277 | -2.773 | 0.00555 | 0.0648 |
| FRG1 | 4452.521 | -0.768 | 0.282 | -2.721 | 0.00650 | 0.0703 |
| PSMC4 | 3309.226 | -0.768 | 0.231 | -3.329 | 0.00087 | 0.0264 |
| WDR62 | 1328.384 | -0.767 | 0.278 | -2.763 | 0.00572 | 0.0652 |
| EIF3F | 17519.778 | -0.765 | 0.287 | -2.663 | 0.00775 | 0.0768 |
| CRYL1 | 1342.814 | -0.763 | 0.282 | -2.705 | 0.00684 | 0.0717 |
| PGP | 974.730 | -0.761 | 0.263 | -2.900 | 0.00374 | 0.0524 |
| ZDHHC12 | 2002.756 | -0.761 | 0.221 | -3.438 | 0.00059 | 0.0230 |
| ECH1 | 6821.106 | -0.759 | 0.250 | -3.037 | 0.00239 | 0.0423 |
| CDKAL1 | 2012.376 | -0.758 | 0.263 | -2.884 | 0.00393 | 0.0540 |
| DNAJC7 | 4639.142 | -0.758 | 0.258 | -2.942 | 0.00326 | 0.0490 |
| PPDPF | 4783.987 | -0.756 | 0.241 | -3.144 | 0.00167 | 0.0358 |
| MZT2A | 3797.404 | -0.756 | 0.242 | -3.128 | 0.00176 | 0.0366 |
| CDKN2C | 715.207 | -0.755 | 0.261 | -2.899 | 0.00374 | 0.0524 |
| FLOT1 | 9547.442 | -0.754 | 0.281 | -2.682 | 0.00732 | 0.0745 |
| EIF6 | 4166.299 | -0.752 | 0.287 | -2.621 | 0.00877 | 0.0831 |
| ENSA | 7806.918 | -0.749 | 0.288 | -2.599 | 0.00935 | 0.0860 |
| KRTCAP2 | 3261.525 | -0.746 | 0.274 | -2.725 | 0.00643 | 0.0698 |
| ATP6V1E1 | 4925.016 | -0.745 | 0.288 | -2.584 | 0.00975 | 0.0882 |
| GALE | 805.057 | -0.743 | 0.238 | -3.121 | 0.00180 | 0.0370 |
| GAR1 | 954.413 | -0.743 | 0.253 | -2.940 | 0.00328 | 0.0492 |
| ACD | 1485.983 | -0.743 | 0.250 | -2.969 | 0.00299 | 0.0469 |
| PPM1G | 8228.572 | -0.741 | 0.269 | -2.759 | 0.00580 | 0.0659 |
| WBSCR22 | 2466.166 | -0.740 | 0.271 | -2.730 | 0.00634 | 0.0691 |
| PSMD13 | 10109.332 | -0.740 | 0.241 | -3.070 | 0.00214 | 0.0399 |
| STK32C | 1514.861 | -0.739 | 0.262 | -2.826 | 0.00471 | 0.0591 |
| MCU | 1191.067 | -0.739 | 0.244 | -3.024 | 0.00250 | 0.0431 |
| DPP7 | 10822.808 | -0.738 | 0.286 | -2.577 | 0.00996 | 0.0890 |
| PSMB2 | 5353.880 | -0.737 | 0.276 | -2.672 | 0.00753 | 0.0757 |
| GALK1 | 1028.101 | -0.736 | 0.283 | -2.604 | 0.00920 | 0.0854 |
| UBAC1 | 1941.544 | -0.735 | 0.273 | -2.689 | 0.00718 | 0.0738 |
| SERTAD3 | 550.166 | -0.735 | 0.276 | -2.661 | 0.00778 | 0.0771 |
| PSMB8 | 10070.255 | -0.730 | 0.241 | -3.028 | 0.00247 | 0.0429 |
| CREG1 | 1570.808 | -0.727 | 0.269 | -2.700 | 0.00692 | 0.0721 |
| GNB2 | 10094.198 | -0.724 | 0.250 | -2.901 | 0.00372 | 0.0524 |
| MDH1 | 8152.318 | -0.724 | 0.268 | -2.703 | 0.00686 | 0.0718 |
| AP1S1 | 2133.037 | -0.723 | 0.245 | -2.950 | 0.00318 | 0.0482 |
| NDUFV1 | 6860.989 | -0.721 | 0.218 | -3.316 | 0.00091 | 0.0267 |
| SLC25A1 | 3673.486 | -0.721 | 0.226 | -3.186 | 0.00144 | 0.0331 |
| UFD1L | 4355.561 | -0.721 | 0.200 | -3.610 | 0.00031 | 0.0183 |
| GSS | 2989.412 | -0.713 | 0.240 | -2.966 | 0.00302 | 0.0471 |
| PPP1R35 | 1377.911 | -0.710 | 0.267 | -2.660 | 0.00782 | 0.0774 |
| TPI1 | 83557.143 | -0.709 | 0.247 | -2.866 | 0.00416 | 0.0557 |
| ATP5A1 | 21441.896 | -0.702 | 0.239 | -2.932 | 0.00337 | 0.0500 |
| UBE2I | 8722.190 | -0.700 | 0.249 | -2.812 | 0.00492 | 0.0606 |
| TXN2 | 4894.734 | -0.698 | 0.257 | -2.715 | 0.00663 | 0.0708 |
| SCPEP1 | 2381.284 | -0.696 | 0.257 | -2.707 | 0.00679 | 0.0715 |
| MRPL1 | 1522.742 | -0.695 | 0.246 | -2.830 | 0.00466 | 0.0587 |
| CD83 | 417.366 | -0.694 | 0.242 | -2.864 | 0.00418 | 0.0558 |
| NDC80 | 3234.968 | -0.694 | 0.268 | -2.590 | 0.00959 | 0.0872 |
| NFKBIB | 2248.029 | -0.692 | 0.226 | -3.057 | 0.00224 | 0.0407 |
| ATP5B | 26311.405 | -0.689 | 0.235 | -2.933 | 0.00336 | 0.0500 |
| UBC | 59123.146 | -0.688 | 0.227 | -3.038 | 0.00238 | 0.0423 |
| MRPL16 | 2078.373 | -0.687 | 0.233 | -2.946 | 0.00322 | 0.0486 |
| BAX | 12949.296 | -0.685 | 0.250 | -2.736 | 0.00622 | 0.0686 |
| MLF2 | 5498.501 | -0.684 | 0.235 | -2.916 | 0.00355 | 0.0515 |
| SYNGR2 | 4687.579 | -0.684 | 0.259 | -2.643 | 0.00821 | 0.0796 |
| IDH3B | 3849.959 | -0.683 | 0.248 | -2.753 | 0.00590 | 0.0667 |
| MCM7 | 8929.362 | -0.682 | 0.248 | -2.754 | 0.00588 | 0.0665 |
| NRM | 2420.348 | -0.679 | 0.235 | -2.888 | 0.00387 | 0.0535 |
| ZFAND2A | 1399.433 | -0.679 | 0.257 | -2.637 | 0.00836 | 0.0806 |
| CDC37 | 8600.206 | -0.679 | 0.221 | -3.075 | 0.00211 | 0.0398 |
| EIF5A | 10549.968 | -0.678 | 0.261 | -2.596 | 0.00942 | 0.0863 |
| UTP11L | 1040.025 | -0.677 | 0.255 | -2.653 | 0.00797 | 0.0781 |
| DDX39A | 5556.053 | -0.677 | 0.223 | -3.035 | 0.00241 | 0.0425 |
| XRCC6 | 14157.155 | -0.676 | 0.241 | -2.808 | 0.00499 | 0.0607 |
| RPS6KB2 | 2864.058 | -0.675 | 0.196 | -3.439 | 0.00058 | 0.0230 |
| SRA1 | 1919.627 | -0.675 | 0.209 | -3.224 | 0.00126 | 0.0308 |
| IFT57 | 920.390 | -0.674 | 0.248 | -2.717 | 0.00659 | 0.0706 |
| ARHGDIB | 56183.537 | -0.673 | 0.258 | -2.613 | 0.00897 | 0.0841 |
| PSMA4 | 5152.307 | -0.672 | 0.258 | -2.603 | 0.00925 | 0.0856 |
| ZFAND2B | 2636.982 | -0.667 | 0.253 | -2.635 | 0.00841 | 0.0809 |
| PLK1 | 2551.448 | -0.666 | 0.251 | -2.654 | 0.00795 | 0.0781 |
| ATG3 | 3098.058 | -0.665 | 0.240 | -2.766 | 0.00567 | 0.0652 |
| MRPL49 | 2758.476 | -0.658 | 0.255 | -2.582 | 0.00983 | 0.0884 |
| AKR7A2 | 2489.986 | -0.657 | 0.254 | -2.585 | 0.00974 | 0.0882 |
| RBM42 | 2939.049 | -0.657 | 0.239 | -2.750 | 0.00595 | 0.0671 |
| CAPZB | 20179.114 | -0.656 | 0.236 | -2.778 | 0.00547 | 0.0642 |
| USP39 | 4071.822 | -0.655 | 0.248 | -2.637 | 0.00836 | 0.0806 |
| PRIM2 | 1431.322 | -0.652 | 0.234 | -2.783 | 0.00539 | 0.0638 |
| MRPL52 | 2356.133 | -0.645 | 0.248 | -2.598 | 0.00937 | 0.0861 |
| VARS | 3293.964 | -0.637 | 0.224 | -2.840 | 0.00451 | 0.0577 |
| XAB2 | 4789.602 | -0.632 | 0.219 | -2.880 | 0.00398 | 0.0542 |
| BSG | 10301.286 | -0.631 | 0.168 | -3.747 | 0.00018 | 0.0152 |
| STK16 | 2616.946 | -0.628 | 0.234 | -2.682 | 0.00731 | 0.0745 |
| CDK2AP2 | 4415.170 | -0.627 | 0.202 | -3.100 | 0.00193 | 0.0382 |
| KATNB1 | 1889.501 | -0.626 | 0.228 | -2.747 | 0.00601 | 0.0673 |
| HDGFRP2 | 3386.957 | -0.625 | 0.221 | -2.827 | 0.00470 | 0.0591 |
| TP53TG1 | 1130.142 | -0.612 | 0.236 | -2.589 | 0.00962 | 0.0874 |
| IER5 | 3878.470 | -0.611 | 0.232 | -2.631 | 0.00851 | 0.0814 |
| AKIRIN2 | 5100.849 | -0.610 | 0.225 | -2.710 | 0.00672 | 0.0710 |
| ARPC1A | 2889.205 | -0.608 | 0.227 | -2.672 | 0.00754 | 0.0757 |
| ANKRD54 | 1260.321 | -0.601 | 0.195 | -3.075 | 0.00211 | 0.0398 |
| FAM127A | 1939.056 | -0.595 | 0.230 | -2.582 | 0.00981 | 0.0884 |
| SF3B2 | 15890.243 | -0.584 | 0.222 | -2.632 | 0.00849 | 0.0814 |
| PSMC1 | 4192.947 | -0.584 | 0.215 | -2.720 | 0.00653 | 0.0703 |
| GPHN | 2431.952 | -0.576 | 0.191 | -3.012 | 0.00259 | 0.0435 |
| CD69 | 3143.228 | -0.571 | 0.213 | -2.679 | 0.00738 | 0.0747 |
| UBL7 | 3434.168 | -0.571 | 0.219 | -2.607 | 0.00914 | 0.0850 |
| PPP2R1A | 11192.119 | -0.569 | 0.212 | -2.680 | 0.00737 | 0.0747 |
| HTRA2 | 2125.711 | -0.557 | 0.208 | -2.682 | 0.00732 | 0.0745 |
| EI24 | 2407.504 | -0.551 | 0.211 | -2.615 | 0.00892 | 0.0839 |
| SLC2A4RG | 3925.125 | -0.548 | 0.197 | -2.780 | 0.00544 | 0.0641 |
| NUBP2 | 2087.835 | -0.534 | 0.206 | -2.592 | 0.00955 | 0.0871 |
| HSF1 | 5132.961 | -0.534 | 0.197 | -2.704 | 0.00685 | 0.0717 |
| SGTA | 3554.948 | -0.525 | 0.178 | -2.954 | 0.00314 | 0.0480 |
| SELT | 4321.797 | -0.519 | 0.188 | -2.755 | 0.00586 | 0.0665 |
| RNH1 | 5941.222 | -0.516 | 0.174 | -2.961 | 0.00307 | 0.0476 |
| TMTC2 | 1281.785 | -0.512 | 0.193 | -2.653 | 0.00797 | 0.0781 |
| AZI1 | 1030.272 | -0.509 | 0.183 | -2.783 | 0.00539 | 0.0638 |
| IPO9 | 5448.770 | 0.507 | 0.179 | 2.830 | 0.00466 | 0.0587 |
| TMEM79 | 426.238 | 0.509 | 0.195 | 2.607 | 0.00913 | 0.0850 |
| PAN3 | 7045.876 | 0.510 | 0.178 | 2.857 | 0.00427 | 0.0562 |
| ZMYM6 | 5494.995 | 0.520 | 0.168 | 3.105 | 0.00190 | 0.0378 |
| TM9SF1 | 3552.800 | 0.524 | 0.184 | 2.852 | 0.00435 | 0.0567 |
| C4orf29 | 3884.881 | 0.528 | 0.176 | 3.004 | 0.00266 | 0.0440 |
| ANGEL2 | 4360.193 | 0.533 | 0.188 | 2.842 | 0.00449 | 0.0577 |
| GOLGB1 | 15054.521 | 0.533 | 0.197 | 2.711 | 0.00671 | 0.0710 |
| TSC1 | 5521.822 | 0.541 | 0.198 | 2.729 | 0.00636 | 0.0692 |
| UBE3B | 5046.216 | 0.561 | 0.204 | 2.750 | 0.00595 | 0.0671 |
| CD96 | 47296.793 | 0.564 | 0.179 | 3.145 | 0.00166 | 0.0357 |
| PPWD1 | 6795.028 | 0.566 | 0.215 | 2.636 | 0.00839 | 0.0808 |
| KRIT1 | 4620.858 | 0.571 | 0.203 | 2.817 | 0.00485 | 0.0599 |
| KIAA2018 | 2685.770 | 0.572 | 0.213 | 2.684 | 0.00726 | 0.0742 |
| CSTF3 | 1532.555 | 0.574 | 0.203 | 2.836 | 0.00457 | 0.0581 |
| APPL2 | 5721.803 | 0.578 | 0.220 | 2.625 | 0.00866 | 0.0825 |
| PDE4A | 5066.288 | 0.580 | 0.212 | 2.733 | 0.00628 | 0.0687 |
| SECISBP2 | 5352.668 | 0.582 | 0.205 | 2.842 | 0.00448 | 0.0577 |
| DNAJC16 | 1962.035 | 0.583 | 0.212 | 2.749 | 0.00599 | 0.0672 |
| IMMP1L | 1251.970 | 0.584 | 0.203 | 2.883 | 0.00394 | 0.0540 |
| HLCS | 1979.221 | 0.585 | 0.180 | 3.248 | 0.00116 | 0.0298 |
| WDR55 | 2926.445 | 0.586 | 0.219 | 2.669 | 0.00761 | 0.0761 |
| THAP6 | 3898.813 | 0.591 | 0.187 | 3.162 | 0.00157 | 0.0348 |
| GABPB2 | 1125.494 | 0.592 | 0.214 | 2.764 | 0.00571 | 0.0652 |
| ATR | 2761.493 | 0.594 | 0.209 | 2.838 | 0.00454 | 0.0579 |
| LOC254128 | 1658.609 | 0.595 | 0.169 | 3.510 | 0.00045 | 0.0207 |
| NFAT5 | 10896.060 | 0.595 | 0.218 | 2.736 | 0.00622 | 0.0686 |
| TMEM154 | 1411.674 | 0.597 | 0.206 | 2.890 | 0.00386 | 0.0535 |
| NAPEPLD | 1198.433 | 0.603 | 0.210 | 2.877 | 0.00401 | 0.0545 |
| C15orf33 | 2048.257 | 0.609 | 0.227 | 2.685 | 0.00726 | 0.0742 |
| ZNF785 | 1553.862 | 0.613 | 0.237 | 2.590 | 0.00960 | 0.0874 |
| SETX | 11408.914 | 0.613 | 0.233 | 2.632 | 0.00849 | 0.0814 |
| EXOC7 | 5665.515 | 0.621 | 0.234 | 2.655 | 0.00793 | 0.0780 |
| WDR35 | 799.978 | 0.624 | 0.235 | 2.654 | 0.00797 | 0.0781 |
| CARD8 | 20217.486 | 0.627 | 0.192 | 3.263 | 0.00110 | 0.0290 |
| DDX26B | 6860.800 | 0.628 | 0.239 | 2.632 | 0.00849 | 0.0814 |
| MLH3 | 2269.624 | 0.631 | 0.241 | 2.617 | 0.00886 | 0.0836 |
| PLBD2 | 1069.525 | 0.633 | 0.243 | 2.602 | 0.00927 | 0.0856 |
| ZNF397 | 2819.495 | 0.636 | 0.244 | 2.606 | 0.00917 | 0.0851 |
| LYPLAL1 | 3494.721 | 0.655 | 0.226 | 2.899 | 0.00374 | 0.0524 |
| LOC340037 | 409.446 | 0.657 | 0.242 | 2.712 | 0.00668 | 0.0709 |
| ALG13 | 6384.279 | 0.659 | 0.203 | 3.245 | 0.00118 | 0.0300 |
| FAM208A | 10022.349 | 0.659 | 0.247 | 2.672 | 0.00753 | 0.0757 |
| ZNF434 | 1651.751 | 0.670 | 0.208 | 3.229 | 0.00124 | 0.0306 |
| ALDH3A2 | 1889.937 | 0.676 | 0.256 | 2.640 | 0.00829 | 0.0801 |
| ZNF189 | 1834.496 | 0.680 | 0.242 | 2.808 | 0.00499 | 0.0607 |
| LOC550112 | 3291.630 | 0.681 | 0.190 | 3.582 | 0.00034 | 0.0190 |
| FBXO38 | 5092.335 | 0.684 | 0.214 | 3.190 | 0.00142 | 0.0331 |
| SNX30 | 1376.515 | 0.688 | 0.234 | 2.943 | 0.00325 | 0.0489 |
| SYTL2 | 14009.297 | 0.695 | 0.267 | 2.602 | 0.00927 | 0.0856 |
| ZNF780A | 2169.036 | 0.701 | 0.269 | 2.600 | 0.00931 | 0.0859 |
| GLS2 | 426.080 | 0.709 | 0.261 | 2.711 | 0.00670 | 0.0710 |
| DFFB | 1575.304 | 0.712 | 0.275 | 2.584 | 0.00976 | 0.0882 |
| SCAND2 | 2645.851 | 0.713 | 0.187 | 3.821 | 0.00013 | 0.0132 |
| ZFP62 | 1421.313 | 0.715 | 0.184 | 3.884 | 0.00010 | 0.0132 |
| LINC00426 | 8208.469 | 0.715 | 0.269 | 2.656 | 0.00791 | 0.0779 |
| PCBP1-AS1 | 6206.773 | 0.717 | 0.209 | 3.439 | 0.00058 | 0.0230 |
| ZNF37A | 3050.070 | 0.719 | 0.249 | 2.882 | 0.00395 | 0.0541 |
| CHD9 | 4600.322 | 0.720 | 0.238 | 3.021 | 0.00252 | 0.0431 |
| TMEM164 | 3046.398 | 0.725 | 0.257 | 2.825 | 0.00472 | 0.0591 |
| ZNF83 | 5394.522 | 0.725 | 0.276 | 2.625 | 0.00866 | 0.0825 |
| PARP8 | 12859.273 | 0.730 | 0.282 | 2.592 | 0.00956 | 0.0871 |
| ZNF175 | 649.771 | 0.731 | 0.259 | 2.826 | 0.00472 | 0.0591 |
| ACPL2 | 2480.871 | 0.732 | 0.271 | 2.706 | 0.00682 | 0.0716 |
| LUZP1 | 2690.200 | 0.739 | 0.267 | 2.764 | 0.00571 | 0.0652 |
| CCDC85C | 703.698 | 0.740 | 0.283 | 2.615 | 0.00893 | 0.0839 |
| MFSD9 | 872.617 | 0.742 | 0.268 | 2.766 | 0.00568 | 0.0652 |
| SAMD9 | 7458.844 | 0.747 | 0.242 | 3.083 | 0.00205 | 0.0396 |
| ZNF41 | 992.603 | 0.748 | 0.280 | 2.672 | 0.00754 | 0.0757 |
| MTERFD2 | 3528.267 | 0.749 | 0.216 | 3.474 | 0.00051 | 0.0221 |
| ZNF619 | 599.255 | 0.751 | 0.271 | 2.774 | 0.00553 | 0.0647 |
| ZNF655 | 22446.413 | 0.754 | 0.231 | 3.269 | 0.00108 | 0.0288 |
| ENTPD7 | 846.777 | 0.764 | 0.275 | 2.782 | 0.00540 | 0.0638 |
| SETD4 | 1396.397 | 0.765 | 0.201 | 3.802 | 0.00014 | 0.0137 |
| ZNF791 | 1326.536 | 0.768 | 0.243 | 3.153 | 0.00162 | 0.0355 |
| C1QTNF3 | 404.771 | 0.770 | 0.261 | 2.957 | 0.00311 | 0.0478 |
| CBY1 | 787.065 | 0.773 | 0.254 | 3.046 | 0.00232 | 0.0417 |
| ODF2L | 7281.748 | 0.773 | 0.267 | 2.891 | 0.00383 | 0.0533 |
| ZNF283 | 1219.146 | 0.783 | 0.244 | 3.207 | 0.00134 | 0.0320 |
| ZNF611 | 2145.204 | 0.788 | 0.283 | 2.784 | 0.00536 | 0.0638 |
| MGA | 3438.811 | 0.789 | 0.280 | 2.811 | 0.00493 | 0.0606 |
| N4BP2L2 | 21903.862 | 0.791 | 0.274 | 2.889 | 0.00386 | 0.0535 |
| ZNF831 | 3728.238 | 0.791 | 0.303 | 2.608 | 0.00911 | 0.0849 |
| MLXIP | 3170.829 | 0.792 | 0.293 | 2.704 | 0.00684 | 0.0717 |
| CCDC146 | 1899.370 | 0.794 | 0.290 | 2.737 | 0.00620 | 0.0686 |
| LOC100506334 | 5991.685 | 0.797 | 0.297 | 2.689 | 0.00717 | 0.0738 |
| LOC375190 | 1929.473 | 0.799 | 0.263 | 3.034 | 0.00242 | 0.0426 |
| C12orf26 | 793.060 | 0.802 | 0.271 | 2.958 | 0.00310 | 0.0478 |
| C12orf35 | 19657.943 | 0.804 | 0.272 | 2.957 | 0.00310 | 0.0478 |
| IFT80 | 4891.793 | 0.806 | 0.244 | 3.296 | 0.00098 | 0.0277 |
| AKD1 | 1299.076 | 0.806 | 0.277 | 2.908 | 0.00364 | 0.0520 |
| ZNF180 | 787.184 | 0.810 | 0.232 | 3.496 | 0.00047 | 0.0211 |
| PHC1 | 1500.097 | 0.810 | 0.277 | 2.930 | 0.00339 | 0.0501 |
| BTN3A1 | 15604.893 | 0.815 | 0.304 | 2.679 | 0.00739 | 0.0747 |
| ZNF169 | 1248.566 | 0.815 | 0.300 | 2.714 | 0.00665 | 0.0708 |
| SLFN12 | 1120.950 | 0.816 | 0.227 | 3.594 | 0.00033 | 0.0187 |
| ZSCAN29 | 2893.565 | 0.816 | 0.303 | 2.695 | 0.00704 | 0.0730 |
| OBFC2A | 5294.250 | 0.819 | 0.180 | 4.553 | 0.00001 | 0.0050 |
| LOC100190939 | 6640.222 | 0.820 | 0.315 | 2.606 | 0.00916 | 0.0851 |
| ZNF195 | 2846.761 | 0.822 | 0.188 | 4.364 | 0.00001 | 0.0064 |
| ENOSF1 | 6254.146 | 0.829 | 0.281 | 2.950 | 0.00318 | 0.0482 |
| C14orf182 | 1537.735 | 0.839 | 0.294 | 2.857 | 0.00427 | 0.0562 |
| ATP6AP1L | 1521.718 | 0.842 | 0.289 | 2.911 | 0.00360 | 0.0519 |
| ANKAR | 563.456 | 0.844 | 0.258 | 3.271 | 0.00107 | 0.0287 |
| SLC16A5 | 1197.970 | 0.849 | 0.282 | 3.006 | 0.00265 | 0.0439 |
| PASK | 11208.162 | 0.855 | 0.219 | 3.898 | 0.00010 | 0.0130 |
| GVINP1 | 12307.335 | 0.858 | 0.251 | 3.419 | 0.00063 | 0.0236 |
| JAKMIP2 | 290.336 | 0.859 | 0.333 | 2.581 | 0.00986 | 0.0885 |
| DET1 | 427.639 | 0.863 | 0.317 | 2.720 | 0.00653 | 0.0703 |
| ITPRIPL2 | 1830.550 | 0.869 | 0.318 | 2.737 | 0.00619 | 0.0686 |
| ZNF599 | 388.937 | 0.871 | 0.279 | 3.122 | 0.00180 | 0.0370 |
| HERC1 | 14078.968 | 0.871 | 0.319 | 2.735 | 0.00624 | 0.0686 |
| ZNF682 | 426.152 | 0.874 | 0.331 | 2.645 | 0.00817 | 0.0793 |
| ATP10A | 5694.299 | 0.875 | 0.329 | 2.655 | 0.00792 | 0.0780 |
| LOC285456 | 647.919 | 0.875 | 0.293 | 2.990 | 0.00279 | 0.0455 |
| AGAP4 | 376.806 | 0.879 | 0.325 | 2.707 | 0.00680 | 0.0715 |
| C6orf170 | 1881.088 | 0.882 | 0.250 | 3.521 | 0.00043 | 0.0203 |
| CTIF | 3046.728 | 0.893 | 0.288 | 3.097 | 0.00195 | 0.0385 |
| SLC22A5 | 1018.309 | 0.894 | 0.236 | 3.790 | 0.00015 | 0.0141 |
| ASB14 | 482.093 | 0.904 | 0.327 | 2.762 | 0.00574 | 0.0653 |
| LOC653160 | 1178.505 | 0.926 | 0.359 | 2.583 | 0.00979 | 0.0884 |
| LOC100130557 | 351.866 | 0.928 | 0.284 | 3.271 | 0.00107 | 0.0287 |
| ZNRD1-AS1 | 1749.134 | 0.941 | 0.334 | 2.819 | 0.00481 | 0.0597 |
| PLEKHM1P | 3526.550 | 0.960 | 0.370 | 2.597 | 0.00942 | 0.0863 |
| FAM161B | 186.894 | 0.964 | 0.346 | 2.783 | 0.00538 | 0.0638 |
| HCG18 | 3301.781 | 0.966 | 0.336 | 2.879 | 0.00399 | 0.0543 |
| ARSG | 3366.982 | 0.968 | 0.357 | 2.714 | 0.00664 | 0.0708 |
| LOC100505894 | 7175.496 | 0.991 | 0.361 | 2.747 | 0.00601 | 0.0673 |
| BMP1 | 1976.504 | 0.995 | 0.274 | 3.635 | 0.00028 | 0.0177 |
| LOC100630923 | 201.103 | 0.997 | 0.368 | 2.713 | 0.00667 | 0.0708 |
| TCP11L2 | 3194.091 | 0.998 | 0.345 | 2.894 | 0.00380 | 0.0529 |
| ZNF235 | 1215.730 | 1.001 | 0.339 | 2.955 | 0.00313 | 0.0480 |
| LOC202181 | 2019.427 | 1.004 | 0.333 | 3.019 | 0.00253 | 0.0432 |
| C14orf149 | 1260.152 | 1.007 | 0.364 | 2.766 | 0.00567 | 0.0652 |
| ITGA4 | 42768.087 | 1.011 | 0.292 | 3.461 | 0.00054 | 0.0223 |
| TMEM220 | 596.928 | 1.027 | 0.374 | 2.747 | 0.00601 | 0.0673 |
| CXorf23 | 1170.174 | 1.028 | 0.284 | 3.617 | 0.00030 | 0.0181 |
| CD84 | 10188.296 | 1.028 | 0.335 | 3.067 | 0.00216 | 0.0401 |
| LRTOMT | 1613.112 | 1.035 | 0.325 | 3.182 | 0.00146 | 0.0333 |
| PILRA | 289.579 | 1.043 | 0.345 | 3.023 | 0.00250 | 0.0431 |
| TUBA3D | 480.483 | 1.048 | 0.401 | 2.611 | 0.00902 | 0.0843 |
| HIST1H4H | 315.537 | 1.068 | 0.344 | 3.104 | 0.00191 | 0.0378 |
| ALOX12P2 | 1080.129 | 1.069 | 0.335 | 3.189 | 0.00143 | 0.0331 |
| MKNK1 | 4364.565 | 1.070 | 0.385 | 2.782 | 0.00541 | 0.0638 |
| C10orf32-AS3MT | 216.968 | 1.072 | 0.402 | 2.668 | 0.00764 | 0.0761 |
| TRIM66 | 7466.939 | 1.091 | 0.406 | 2.688 | 0.00719 | 0.0739 |
| PITPNM2 | 1173.762 | 1.092 | 0.370 | 2.953 | 0.00315 | 0.0480 |
| PCMTD2 | 9882.534 | 1.100 | 0.317 | 3.468 | 0.00052 | 0.0221 |
| KIAA1024 | 217.064 | 1.108 | 0.360 | 3.081 | 0.00206 | 0.0396 |
| LPAL2 | 300.318 | 1.109 | 0.323 | 3.430 | 0.00060 | 0.0232 |
| GIMAP8 | 3686.561 | 1.109 | 0.379 | 2.927 | 0.00342 | 0.0505 |
| DAPK2 | 5807.019 | 1.122 | 0.387 | 2.897 | 0.00377 | 0.0527 |
| FLJ12334 | 1360.492 | 1.124 | 0.436 | 2.579 | 0.00991 | 0.0888 |
| SEPP1 | 267.911 | 1.126 | 0.374 | 3.014 | 0.00258 | 0.0434 |
| CYP4V2 | 2431.980 | 1.127 | 0.401 | 2.809 | 0.00498 | 0.0607 |
| KCNC3 | 3702.732 | 1.136 | 0.339 | 3.355 | 0.00079 | 0.0258 |
| RNF43 | 1093.263 | 1.137 | 0.430 | 2.646 | 0.00814 | 0.0792 |
| SPNS2 | 297.132 | 1.142 | 0.412 | 2.770 | 0.00560 | 0.0652 |
| ZNF493 | 2079.766 | 1.144 | 0.363 | 3.152 | 0.00162 | 0.0355 |
| CABP4 | 351.037 | 1.152 | 0.430 | 2.679 | 0.00738 | 0.0747 |
| ZFP3 | 372.961 | 1.166 | 0.353 | 3.306 | 0.00095 | 0.0272 |
| ZNF44 | 3431.663 | 1.172 | 0.358 | 3.272 | 0.00107 | 0.0287 |
| KCNAB3 | 386.246 | 1.176 | 0.335 | 3.516 | 0.00044 | 0.0204 |
| KCNG2 | 54.780 | 1.181 | 0.443 | 2.667 | 0.00765 | 0.0761 |
| KRBA2 | 412.314 | 1.183 | 0.307 | 3.846 | 0.00012 | 0.0132 |
| CCDC48 | 147.997 | 1.184 | 0.366 | 3.233 | 0.00123 | 0.0305 |
| MGC72080 | 4632.339 | 1.186 | 0.363 | 3.268 | 0.00108 | 0.0288 |
| SYS1-DBNDD2 | 151.661 | 1.193 | 0.458 | 2.607 | 0.00914 | 0.0850 |
| ADAMTS16 | 144.022 | 1.193 | 0.361 | 3.308 | 0.00094 | 0.0272 |
| EMR2 | 84.054 | 1.209 | 0.422 | 2.862 | 0.00421 | 0.0560 |
| LOC100616668 | 3666.160 | 1.214 | 0.342 | 3.553 | 0.00038 | 0.0195 |
| LDHD | 51.756 | 1.221 | 0.442 | 2.763 | 0.00572 | 0.0652 |
| BCO2 | 488.354 | 1.225 | 0.374 | 3.278 | 0.00104 | 0.0286 |
| THBS4 | 599.962 | 1.234 | 0.379 | 3.258 | 0.00112 | 0.0292 |
| ZNF594 | 781.594 | 1.235 | 0.465 | 2.657 | 0.00789 | 0.0778 |
| POLN | 1403.647 | 1.240 | 0.326 | 3.810 | 0.00014 | 0.0137 |
| CLDN14 | 502.071 | 1.243 | 0.397 | 3.132 | 0.00174 | 0.0364 |
| LOC100132111 | 855.467 | 1.250 | 0.414 | 3.018 | 0.00255 | 0.0432 |
| ZNF780B | 3856.800 | 1.262 | 0.386 | 3.273 | 0.00107 | 0.0287 |
| ZNF37BP | 4891.323 | 1.263 | 0.404 | 3.128 | 0.00176 | 0.0366 |
| GIPC3 | 143.717 | 1.265 | 0.469 | 2.696 | 0.00703 | 0.0729 |
| ZNF70 | 403.763 | 1.268 | 0.423 | 2.997 | 0.00273 | 0.0447 |
| ZNF284 | 378.543 | 1.274 | 0.444 | 2.871 | 0.00409 | 0.0552 |
| LDHAL6A | 475.575 | 1.275 | 0.454 | 2.811 | 0.00494 | 0.0607 |
| MIR3916 | 466.855 | 1.277 | 0.491 | 2.604 | 0.00922 | 0.0854 |
| LOC283624 | 1467.326 | 1.291 | 0.423 | 3.050 | 0.00229 | 0.0414 |
| KIAA1257 | 250.845 | 1.308 | 0.487 | 2.687 | 0.00721 | 0.0739 |
| RFX8 | 1488.238 | 1.314 | 0.341 | 3.849 | 0.00012 | 0.0132 |
| PCDH12 | 304.208 | 1.315 | 0.429 | 3.067 | 0.00216 | 0.0401 |
| FAM179A | 2386.861 | 1.320 | 0.402 | 3.286 | 0.00102 | 0.0282 |
| CXCL13 | 147.724 | 1.327 | 0.512 | 2.592 | 0.00954 | 0.0871 |
| RTKN2 | 3554.770 | 1.334 | 0.362 | 3.685 | 0.00023 | 0.0170 |
| LRRC2 | 190.023 | 1.347 | 0.509 | 2.646 | 0.00813 | 0.0792 |
| PAX8 | 851.377 | 1.348 | 0.510 | 2.641 | 0.00827 | 0.0799 |
| ARL10 | 893.719 | 1.379 | 0.441 | 3.127 | 0.00177 | 0.0366 |
| RBM26-AS1 | 328.216 | 1.390 | 0.291 | 4.775 | 0.00000 | 0.0043 |
| CNTNAP1 | 347.303 | 1.395 | 0.378 | 3.691 | 0.00022 | 0.0170 |
| LOC553103 | 801.719 | 1.397 | 0.481 | 2.904 | 0.00369 | 0.0521 |
| MKRN3 | 344.486 | 1.405 | 0.401 | 3.507 | 0.00045 | 0.0208 |
| GOLGA6L5 | 443.386 | 1.417 | 0.398 | 3.559 | 0.00037 | 0.0193 |
| TNS1 | 1587.994 | 1.420 | 0.533 | 2.664 | 0.00772 | 0.0767 |
| IL7R | 20435.585 | 1.422 | 0.420 | 3.389 | 0.00070 | 0.0246 |
| AVPR2 | 152.221 | 1.428 | 0.540 | 2.644 | 0.00820 | 0.0795 |
| GOLGA7B | 329.636 | 1.429 | 0.386 | 3.699 | 0.00022 | 0.0167 |
| GABRR2 | 200.113 | 1.432 | 0.547 | 2.617 | 0.00887 | 0.0836 |
| LOC644714 | 71.553 | 1.447 | 0.501 | 2.886 | 0.00391 | 0.0538 |
| LOXL4 | 38.311 | 1.460 | 0.542 | 2.694 | 0.00706 | 0.0730 |
| WNT9A | 645.877 | 1.472 | 0.532 | 2.766 | 0.00567 | 0.0652 |
| C3orf20 | 656.563 | 1.486 | 0.433 | 3.431 | 0.00060 | 0.0232 |
| ATP10B | 745.756 | 1.496 | 0.370 | 4.039 | 0.00005 | 0.0106 |
| CCDC144B | 689.138 | 1.500 | 0.318 | 4.721 | 0.00000 | 0.0043 |
| LOC100506274 | 402.492 | 1.549 | 0.433 | 3.573 | 0.00035 | 0.0191 |
| GPX3 | 244.101 | 1.556 | 0.531 | 2.932 | 0.00336 | 0.0500 |
| C15orf52 | 205.276 | 1.586 | 0.520 | 3.052 | 0.00227 | 0.0412 |
| SDK2 | 3214.512 | 1.591 | 0.559 | 2.844 | 0.00445 | 0.0575 |
| KLRC4-KLRK1 | 2148.503 | 1.671 | 0.601 | 2.779 | 0.00545 | 0.0641 |
| SULT1B1 | 149.222 | 1.698 | 0.648 | 2.622 | 0.00875 | 0.0830 |
| ZNF844 | 71.621 | 1.698 | 0.414 | 4.102 | 0.00004 | 0.0090 |
| LOC441009 | 101.728 | 1.707 | 0.597 | 2.861 | 0.00422 | 0.0560 |
| ZNF439 | 319.814 | 1.718 | 0.430 | 3.996 | 0.00006 | 0.0115 |
| ACE | 810.698 | 1.761 | 0.581 | 3.030 | 0.00244 | 0.0428 |
| HPCAL4 | 406.717 | 1.779 | 0.605 | 2.938 | 0.00331 | 0.0494 |
| PPT2-EGFL8 | 141.836 | 1.796 | 0.522 | 3.440 | 0.00058 | 0.0230 |
| LOC400685 | 224.785 | 1.838 | 0.381 | 4.827 | 0.00000 | 0.0043 |
| LOC100288842 | 59.792 | 1.859 | 0.711 | 2.615 | 0.00894 | 0.0839 |
| SIRPB1 | 253.024 | 1.860 | 0.659 | 2.823 | 0.00476 | 0.0593 |
| ZNF300 | 135.047 | 1.946 | 0.652 | 2.984 | 0.00284 | 0.0458 |
| TSPY26P | 37.303 | 2.004 | 0.751 | 2.669 | 0.00760 | 0.0761 |
| OVCH1 | 385.931 | 2.059 | 0.744 | 2.767 | 0.00566 | 0.0652 |
| ZNF69 | 620.617 | 2.069 | 0.397 | 5.206 | 0.00000 | 0.0008 |
| ALDOB | 31.956 | 2.249 | 0.745 | 3.021 | 0.00252 | 0.0431 |
| PLGLB2 | 43.899 | 2.370 | 0.741 | 3.197 | 0.00139 | 0.0328 |
| BDAG1 | 29.688 | 2.488 | 0.783 | 3.178 | 0.00148 | NA |
| CYP11B1 | 61.639 | 2.515 | 0.797 | 3.156 | 0.00160 | 0.0353 |
| GUSBP3 | 153.381 | 2.671 | 0.895 | 2.982 | 0.00286 | 0.0458 |
| CYP2A6 | 38.527 | 2.923 | 0.949 | 3.078 | 0.00208 | 0.0396 |
| REG4 | 43.723 | 2.995 | 0.968 | 3.095 | 0.00197 | 0.0386 |
| LOC339822 | 63.025 | 3.120 | 1.202 | 2.596 | 0.00944 | 0.0864 |
| USP32P1 | 7064.655 | 3.162 | 0.897 | 3.527 | 0.00042 | 0.0203 |
| MIR4737 | 14.611 | 3.300 | 1.088 | 3.032 | 0.00243 | NA |
| NBPF9 | 7.039 | 3.991 | 1.457 | 2.739 | 0.00617 | NA |
| NEUROD2 | 6.950 | 4.874 | 1.610 | 3.028 | 0.00246 | NA |
| CCDC144NL | 6.990 | 4.885 | 1.720 | 2.840 | 0.00451 | NA |
| TAS2R40 | 6.059 | 5.197 | 1.871 | 2.778 | 0.00547 | NA |
| FAM181A | 5.973 | 5.683 | 2.018 | 2.816 | 0.00486 | NA |
| MTRNR2L4 | 9.826 | 5.912 | 1.792 | 3.299 | 0.00097 | NA |
